# Biosynthesis of 14-membered cyclopeptide alkaloids via non-heme-iron- and 2-oxoglutarate-dependent oxidative decarboxylation

**DOI:** 10.64898/2026.01.20.700549

**Authors:** Jordan Hungerford, Lisa S. Mydy, Xiaofeng Wang, Lorena Mendoza-Perez, Derrick A. Ousley, Khadija Shafiq, Kali M. McDonough, Wenjie Li, Gabrielle May, Desnor N. Chigumba, Shengrui Yao, Roland D. Kersten

## Abstract

Cyclopeptide alkaloids are an expanding class of plant peptide natural products defined by a macrocyclic ether-crosslink via a tyrosine-derived phenol. Classical cyclopeptide alkaloids are characterized by strained 13-to 15-membered cyclophanes and terminal modifications such as *N*-methylation and C-terminal styrylamine moieties. While synthetic access to many classical cyclopeptide alkaloids has been established, no biosynthetic route has been reported. Here, we elucidate the biosynthetic pathway of a 14-membered cyclopeptide alkaloid, lotusine A, from Chinese date tree (*Ziziphus jujuba*) which features peptide cyclization on a ribosomal precursor peptide by a split burpitide cyclase, non-heme-iron and 2-oxoglutarate-dependent oxidative decarboxylation affording the C-terminal hydroxystyrylamine, and SAM-dependent N-terminal α-*N,N*-dimethylation. We apply discovered *Z. jujuba* enzymes in combination with a clubmoss cyclopeptide alkaloid cyclase for biosynthesis and diversification of analgesic adouetine X and anxiolytic sanjoinine A by combining *in planta* and *in vitro* reactions. Our work expands the biocatalytic repertoire of non-heme-iron- and 2-oxoglutarate-dependent enzymology to oxidative peptide decarboxylation and primes scaled metabolic engineering and chemoenzymatic synthesis of 14-membered cyclopeptide alkaloids with terminal posttranslational modifications.

## Introduction

Cyclopeptide alkaloids (CPAs) are a diverse class of plant peptides, which were structurally first characterized in the 1960s as small cyclic peptides featuring 13-to 15-membered macrocycles from buckthorn (Rhamnaceae) plants^1^. CPAs have medicinal activities such as analgesic adouetine X^2,3^, anxiolytic sanjoinine A^4^, and antiviral jubanines^5^ and have therefore been leads for medicinal chemistry (**Figure 1A, Figure S1**). Most CPAs are structurally characterized by an ether-crosslink between a C-terminal hydroxystyrylamine, tyrosine, or octopamine to an unactivated β-carbon of another amino acid such as leucine (33% of known CPAs, **Data S1**), proline (40% of known CPAs), or phenylalanine (21% of known CPAs)^6,7^. In addition, classical CPAs feature terminal modifications such as α-*N,N*-dimethylation (54% of known CPAs) and a C-terminal hydroxystyrylamine (87% of known CPAs, **Data S1**)^6,7^. Despite their promising bioactivities and drug-like properties, exploration of CPAs for drug discovery has been limited by low isolation yields from source plants and synthetic challenges of the strained macrocycles, the C-terminal enamide and peptide diversification^8,9^.

**Figure 1.**
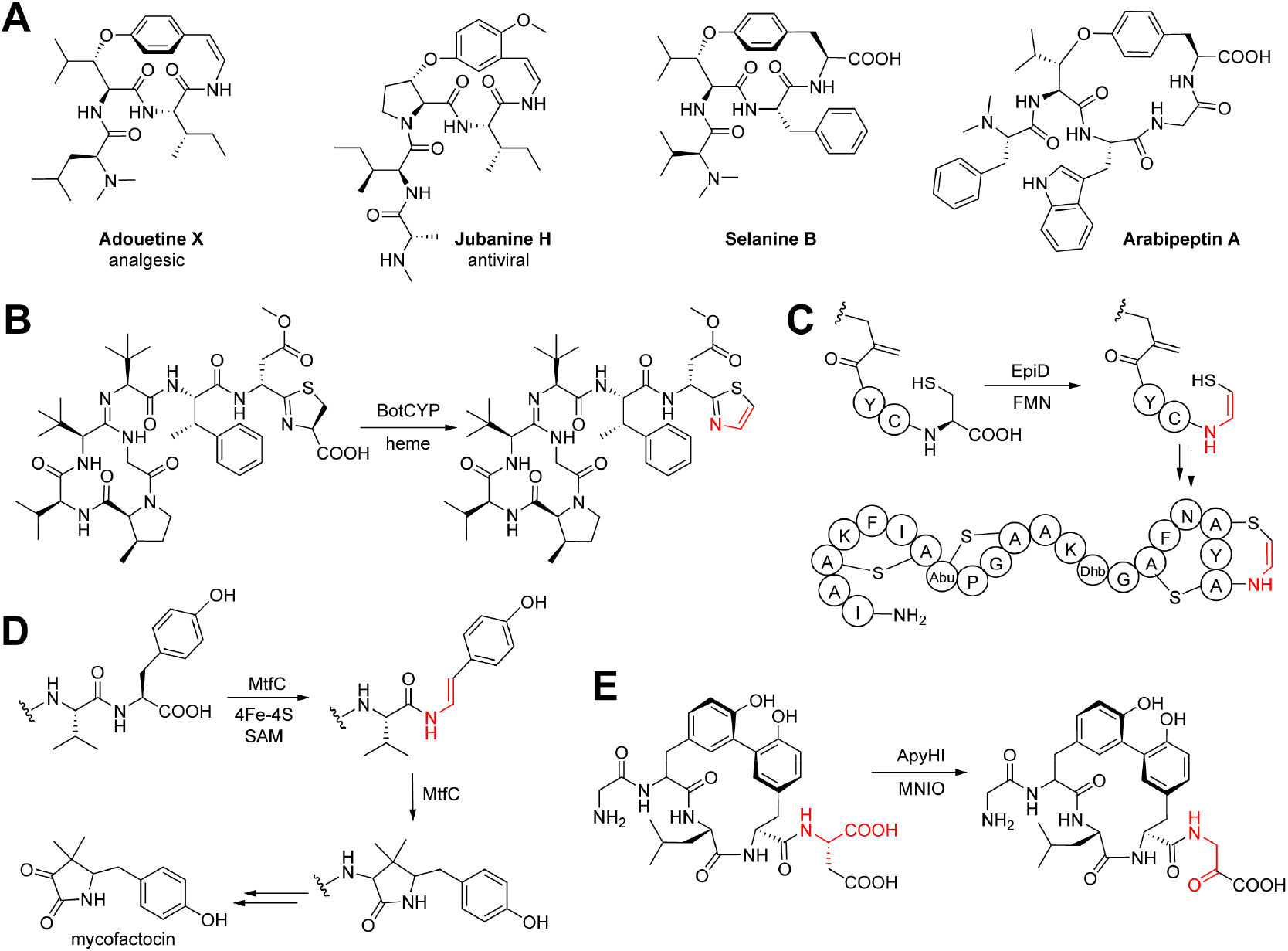
C-terminal RiPP decarboxylation. (**A**) Plant cyclopeptide alkaloid structures. Adouetine X and jubanine H are representatives of decarboxylated (classical) CPAs with a C-terminal hydroxystyrylamine. Selanine B and arabipeptin A represent characterized CPA-burpitide RiPPs with a C-terminal tyrosine. (**B**) Oxidative decarboxylation in bottomycin A2 biosynthesis via CYP450 enzyme BotCYP. (**C**) C-terminal oxidative decarboxylation of epidermin precursor peptide via flavoenzyme EpiD and subsequent aminovinyl-cysteine formation. (**D**) Oxidative decarboxylation of a C-terminal tyrosine during mycofactocin biosynthesis catalyzed by radical SAM enzyme MftC. (**E**) Oxidative decarboxylation by MNIO enzyme ApyH with RRE-containing protein ApyI towards C-terminal β-amino-α-keto acid formation. Decarboxylation modifications are highlighted in red in B-D. Abbreviations: FMN - flavin mononucleotide, SAM - S-adenosylmethionine, MNIO - multinuclear non-heme-iron oxidative enzyme, RRE - RiPP recognition element^86^.

Recently, plant cyclopeptides with tyrosine-derived C-O-crosslinks such as 14-membered selanine B from *Selaginella kraussiana* were biosynthetically defined as ribosomally-synthesized and posttranslationally modified peptides (RiPP)^10^ called burptides^7,11,12^, which are derived from copper-dependent peptide cyclases with BURP domains^11–15^. Selanine B is macrocyclized by ether-crosslinking of a C-terminal tyrosine-phenol-hydroxy to a leucine-β-carbon in an autocatalytic copper-dependent reaction by burpitide cyclase SkrBURP that is fused to its core peptide substrates and also generates bicyclic cyclopeptide alkaloids with an additional 17-membered C-terminal macrocycle such as selanine A via a second C-terminal tyrosine-crosslink (**Figure S1**)^11^. Another example of a CPA-burpitide with a C-O-macrocyclic bond found is arabipeptin A, which is a 17-membered cyclopeptide. Arabipeptin A core peptide is cyclized by ArbB, a burpitide cyclase which is split from its core peptide substrates in a separate precursor peptide^12,15^ co-localized on the genome with its burpitide cyclase. Both arabipeptin A and selanine B lack the enamine moeity of classical CPAs as they have a C-terminal carboxy group. Based on the formation of CPA-type ether crosslinks formed by burpitide cyclases such as SkrBURP, it has been proposed to re-classify CPAs on the basis of biosynthesis as burpitides with tyrosine-derived ether crosslinks^7^.

Burpitide cyclases contain a dicopper center defined by a 4xCysHis-motif within the BURP-domain fold^14^. Burpitide cyclases such as legumenin cyclase AhyBURP catalyze crosslinking of an aromatic residue such as tryptophan or tyrosine with a C(*sp*^*3*^) carbon in another amino acid of a core peptide motif by a radical, dioxygen-dependent reaction^14,16^. Decarboxylated peptide C-termini as observed in classical CPAs occur in several bacterial RiPPs such as lanthipeptides, linaridins, and bottromycins (**Figure S1**)^17–22^. In lanthipeptides and linaridins, which feature a C-terminal aminovinyl-cysteine posttranslational modification, a C-terminal cysteine is oxidatively decarboxylated by a flavoenzyme (**Figure 1B**)^17–22^. Bottromycins contain a C-terminal thiazole formed from a thiazoline through oxidative decarboxylation by a CYP450 enzyme (**Figure 1C**)^23^. Other C-terminal decarboxylation reactions in RiPP biosynthesis were reported in mycofactocin biosynthesis via a radical *S*-adenosyl-L-methionine (SAM) enzyme mechanism (**Figure 1D**)^24,25^ and for the formation of a *Burkholderia* RiPP with a C-terminal β-amino-α-keto acid generated by a multinuclear non-heme-iron oxidative enzyme (**Figure 1E**)^26^. Similarly, multiple RiPP classes such as lanthipeptides^27–30^, linaridins^21^, phomopsins^31,32^, linear azol(in)e-containing peptides^33^ have N-terminal α-*N*-methylations that are often generated by SAM-dependent methyltransferases (**Figure S1**)^34^. Burpitides selanine B and arabipeptin A also contain α-*N,N*-dimethylation but the biosynthetic basis of this modification is unknown.

Here, we establish the biosynthetic pathway to 14-membered CPAs with N-terminal α-*N*-methylation and C-terminal oxidative decarboxylation from *Ziziphus jujuba* and utilize the discovered enzymes including a non-heme-iron and 2-oxoglutarate-dependent decarboxylase for the biosynthesis of proline- and leucine-macrocyclic CPAs such as lotusine A and analgesic adouetine X, respectively.

## Results

### Characterization of 14-membered CPA-burpitide cyclase ZjuBURP

To elucidate a classical CPA biosynthetic pathway, we targeted Chinese date tree (*Ziziphus jujuba*, Rhamnaceae), a common source plant of medicinal CPA chemistry including adouetine X, sanjoinine A and jubanines. Chekan and colleagues identified through bioinformatics candidate split precursor genes of 13- and 14-membered CPAs in the *Z. jujuba* genome (**Figure 2A**), which co-localize with BURP-domain-encoding genes of the BNM2 subfamily^12,35^. These proposed burpitide precursor genes constitute an entry point to biosynthetic pathway elucidation of classical CPAs featuring both ether-crosslinks, alkylated N-termini and decarboxylated C-termini, which are highly diversified in *Ziziphus* plants^6,7^. To characterize a burpitide cyclase, we chose a BURP-domain-containing gene co-localized with a candidate precursor gene on chromosome 9 of *Z. jujuba* cultivar Dongzao^36^ which had eleven predicted core peptides of FPIY and three predicted core peptides of FLIY (**Figure 2A**). FPIY matched previously identified cyclopeptide alkaloids from *Ziziphus sp*. such as 14-membered lotusine A and 13-membered daechuine S26 (**Figure S1**)37,38.

**Figure 2.**
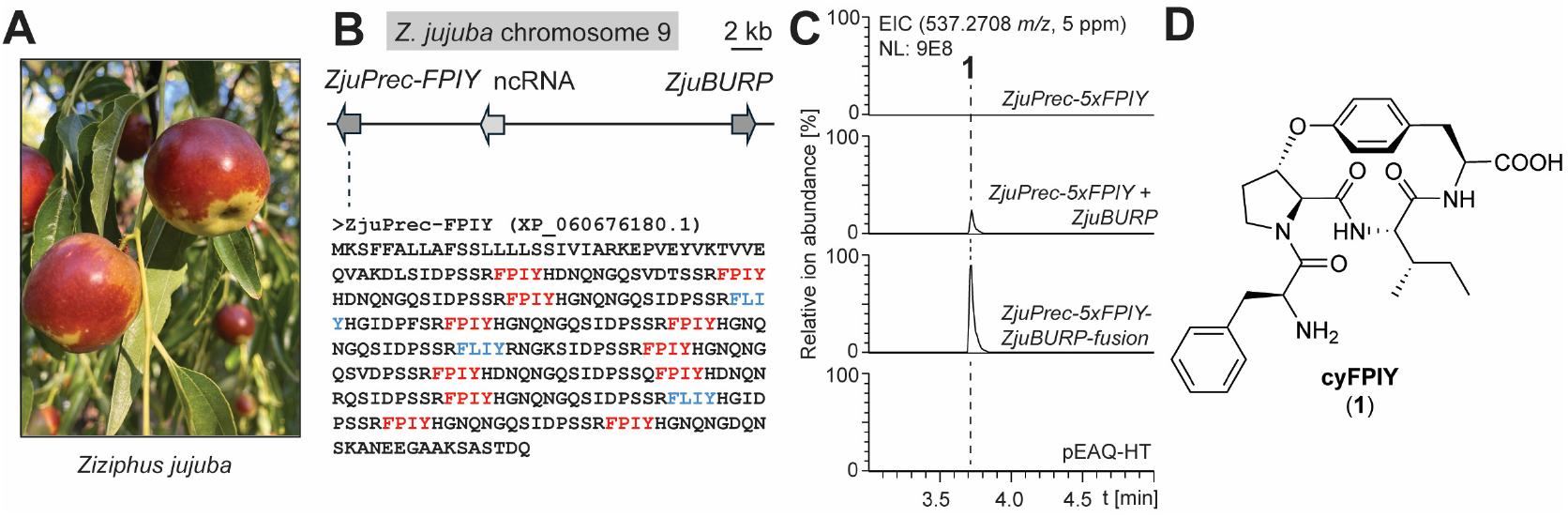
Characterization of 14-membered CPA-burpitide cyclase ZjuBURP from *Ziziphus jujuba*. (**A**) *Ziziphus jujuba* cultivar Dongzao. (**B**) *ZjuBURP* locus in the *Z. jujuba* genome with co-localized *ZjuPrec-FPIY* gene. Predicted core peptides are highlighted in blue and red color. (**C**) LCMS detection of cyFPIY peptide in transgenic *N. benthamiana* after 7 days of coexpression of ZjuPrec-5xFPIY and ZjuBURP or a ZjuPrec-5xFPIY-ZjuBURP-fusion construct. (**D**) cyFPIY structure isolated from transgenic *N. benthamiana* after scaled expression of *ZjuPrec-5xFPIY-ZjuBURP-fusion*. Abbreviations: ncRNA - non-coding RNA, EIC extracted ion chromatogram.

We tested if the BURP-domain-containing gene encodes a split burpitide cyclase such as ArbB that is involved in macrocyclization of core peptides such as FPIY motifs encoded in the co-localized precursor gene. A truncated version of the target precursor gene with five FPIY core peptide repeats, named ZjuPrec-5xFPIY, was synthesized, cloned and expressed with the pEAQ-HT plasmid in transgenic *Nicotiana benthamiana* via agroinfiltration for seven days^39^. Methanol extracts of the transgenic *N. benthamiana* leaves showed no analytes corresponding to a monocyclic FPIY peptide (**Figure 2B**). Co-expression of pEAQ-HT-*ZjuPrec-5xFPIY* and the candidate burpitide cyclase gene *ZjuBURP* led to the detection of an analyte with an *m/z* value of 537.271 corresponding to a monocyclic FPIY peptide with unmodified peptide termini ([M+H]^+^, **Figure 2B**). To determine the structure of analyte cyFPIY (**1**), we applied a scaled expression of the cyclase-precursor peptide pair in *N. benthamiana*. To optimize yields, we tested if a fused version of the split burpitide cyclase and the truncated precursor peptide would improve cyFPIY yields in transgenic *N. benthamiana*. Expression of ZjuPrec-5xFPIY-ZjuBURP-fusion yielded the same cyFPIY product from the split system while increasing yields by 2.9-fold from agroinfiltration based on cyFPIY EIC area-under-the-curve values from infiltrated tobacco leaf extracts (**Figure 2B, Figure S2**). Scaled expression of the fusion gene enabled reproducible production of 5 mg pure cyFPIY (**1**) per 1 kg of tobacco wet weight. 1D and 2D NMR analysis of **1** revealed a 14-membered FPIY peptide in which the macrocyclic C-O-crosslink is between the C-terminal tyrosine-phenol-hydroxy group and the β-carbon of the proline (**Figure 2C, Figures S3-S10, Table S1**). Marfey’s analysis showed L-Phe, L-Pro, L-Ile, and L-Tyr (**Figure S11**). In the ROESY spectrum, a correlation is observed between Pro2-Hα and Trp4-H5, as well as between Pro2-Hβ and Trp4-H3. Together with the *J*-value of 7 Hz for the Cα–Cβ bond in Pro2, this indicates an L-erythro stereochemistry, corresponding to an S-configuration at the Pro2 Cβ position (**Figure S6, Table S1**)^40^.

*In vitro* reconstitution of ZjuBURP was challenged by low protein yields. The production of a cyclic 14-membered CPA from transgenic *N. benthamiana* expressing ZjuBURP in the presence of a split predicted CPA precursor peptide such as ZjuPrec-FPIY suggests that ZjuBURP is a burpitide cyclase installing the ether-crosslink on the core peptide of the precursor peptide, which is further processed proteolytically to cyFPIY during CPA biosynthesis in *Z. jujuba. In vitro* precedence for split 14-membered CPA cyclization was recently provided by ArbB and a ZjuBURP-homologous split burpitide cyclase, CamBURP1 from *Ceanothus americanus*, which were reconstituted as a copper-dependent peptide cyclases^15^.

### Discovery of a non-heme-iron and 2-oxoglutarate-dependent CPA-decarboxylase from jujube

Next, we explored jujube for biosynthetic genes involved in the terminal modifications of CPAs after we established a CPA-burpitide cyclase. A *Ziziphus jujuba* var. Dongzao cultivar was obtained and analyzed for the presence of CPAs in its mature leaf and stem tissues. While no CPAs such as lotusine A matching an FPIY core peptide could be detected in jujube samples, a candidate CPA analyte with an *m/z* value of 696.3756 was detected in high abundance in mature stem tissue via CPA-targeted molecular networking (**Figure S12**). This CPA was isolated and structurally elucidated by 1D and 2D NMR, Marfey’s analysis and PGME derivatization (**Figure S13-21, Table S2**)^41^ to be a 13-membered CPA with a FIPFY core peptide, α-*N,N*-dimethylation on its N-terminus, and a meta-crosslinked hydroxystyrylamine moeity with an ortho-methoxy group (**Figure S22**). This peptide is a new CPA named jubanine K, which showed that the target *Ziziphus jujuba* plants have the metabolic capacity for oxidative decarboxylation and N-methylation of CPAs.

To identify the enzymes catalyzing the N-terminal modifications of jujube CPAs, we used a multi-omic approach to find gene candidates upregulated in tissues with increased CPA content. Young stem and leaf tissues of the same cultivar used for jubanine K isolation showed low levels of jubanine K in its stem tissue and no jubanine K mass signals in the young leaves. However, a putative biosynthetic intermediate for jubanine K without the methoxy group was detected in high ion abundance in young stems compared to young leaves (**Figure S22**). To identify candidate genes for terminal CPA modifications, young stem and young leaf mRNA were sequenced by RNA-seq. Using ZjuPrec-FPIY as a query, a candidate jubanine K precursor peptide was identified by BLAST search^42^ in the assembled transcriptome which features a core peptide of FIPFY matching jubanine K structure and was coexpressed with ZjuBURP (**Table S3**). Co-expression of this precursor peptide with ZjuBURP yielded a monocyclic FIPFY analyte based on MS/MS analysis indicating that ZjuBURP is part of the jubanine K biosynthetic pathway (**Figure S23**). We hypothesized that either *N*-methylation or decarboxylation could occur after the cyclization and a proteolytic processing step to yield the cyclic core peptide. Both routes were supported based on the detection of low intensity analytes corresponding to *N,N*-dimethylated cyclic FIPFY and decarboxylated cyclic FIPFY in young *Z. jujuba* stem extracts (**Figure S22**). Using ZjuBURP as a bait gene, we searched for oxidase genes, which were upregulated in young stems compared to young leaves. Several CYP450 and non-heme-iron dioxygenase genes with similar expression patterns as ZjuBURP (**Table S3**) were synthesized, cloned into pEAQ-HT, and co-expressed with ZjuPrec-5xFPIY-ZjuBURP-fusion in *N. benthamiana*. One dioxygenase gene, named ZjuDC, yielded an analyte in *N. benthamiana* co-expression experiments, which had a mass loss corresponding to oxidative decarboxylation, i.e. a mass loss of two hydrogens and CO_2_, compared to cyFPIY (**Figure 3A**). Bioinformatic analysis predicted *ZjuDC* to encode a member of the non-heme-iron and 2-oxoglutarate (2OG)-dependent dioxygenase family, as the sequence and AlphaFold3^43^ model of ZjuDC revealed the characteristic iron-coordinating His-X-(Asp/Glu)-X_48-153_-His motif in comparison to experimentally determined structures of homologous plant dioxygenases (**Figure S24**)^44–47^.

**Figure 3.**
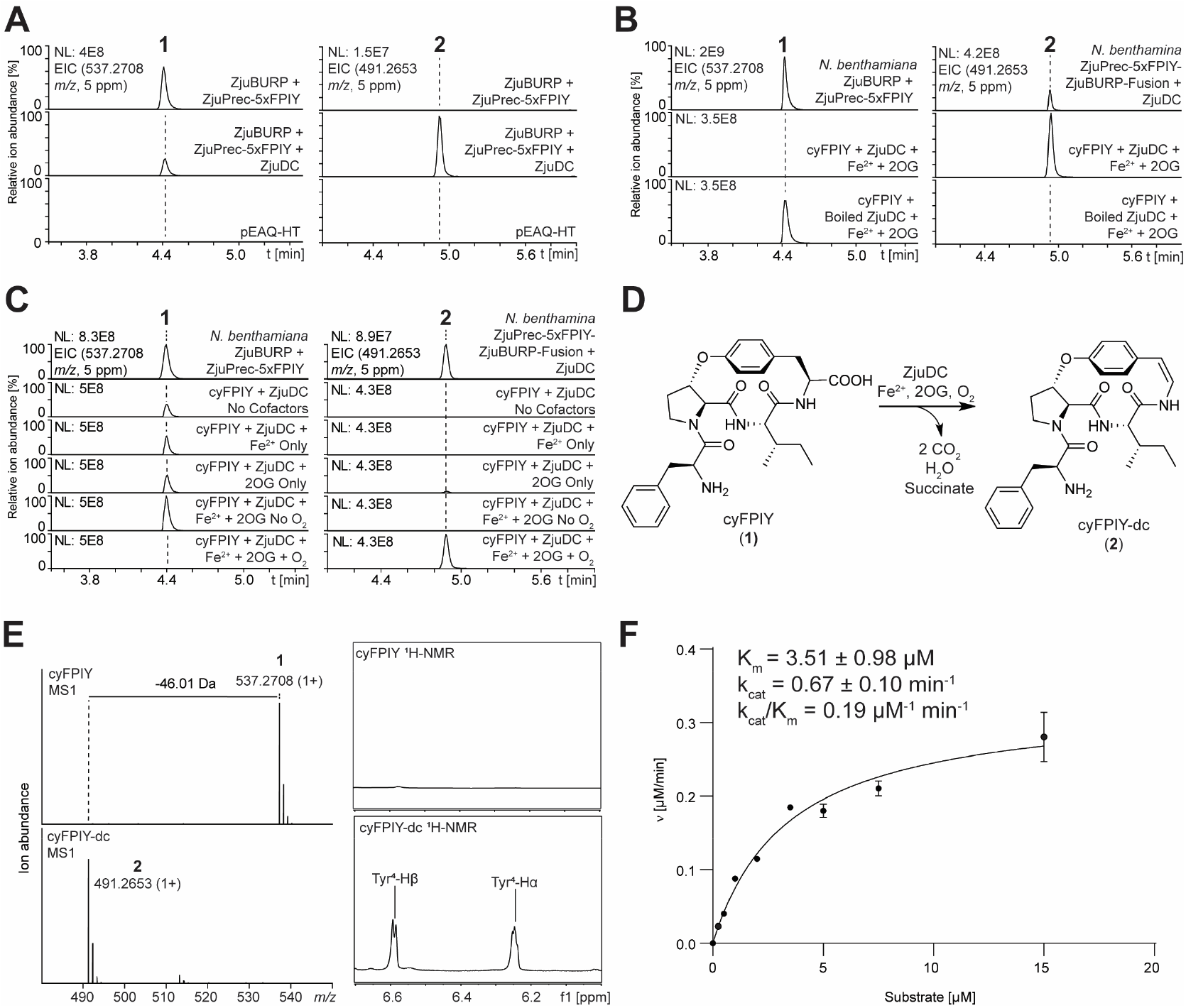
Discovery and characterization of a non-heme-iron and 2-oxoglutarate-dependent enzyme for oxidative CPA decarboxylation. (**A**) EIC traces of cyFPIY and decarboxylated cyFPIY in methanolic extracts of transgenic *N. benthamiana* leaves expressing ZjuPrec-3xFPIY-ZjuBURP with or without candidate decarboxylase ZjuDC for 7 days. (**B**) *In vitro* reconstitution of ZjuDC activity in the presence of Fe(II) and 2OG. (**C**) In vitro cofactor and substrate screening assays of ZjuDC. (**D**) ZjuDC reaction. (**E**) Characterization of oxidative decarboxylation in cyFPIY by ZjuDC reaction. (**F**) Kinetic characterization of ZjuDC with cyFPIY substrate. Abbreviations: 2OG - 2-oxoglutarate, EIC – extracted ion chromatogram.

To test ZjuDC decarboxylation of cyFPIY, ZjuDC was expressed heterologously in *E. coli* BL21(DE3) and purified via immobilized metal affinity chromatography and size exclusion chromatography (**Figure S25**). *In vitro* incubation of ZjuDC with cyFPIY in the presence of 2-oxoglutarate (2-OG), O_2_ and Fe(II) yielded the decarboxylated analyte cyFPIY-dc (**2**) observed *in planta* (**Figure 3B**). Further *in vitro* enzyme assays showed that the ZjuDC reaction requires 2OG, O_2_ and Fe(II) for the conversion of cyFPIY into **2** (**Figure 3C**) while generating succinate (**Figure S26**). We tried to generate analyte **2** at scale in *N. benthamiana*, however its purification was challenging from transgenic leaves to reach milligram quantities. Therefore, we purified 5.5 mg of cyFPIY from transgenic *N. benthamiana* via ZjuPrec-5xFPIY-ZjuBURP-fusion expression and converted it in a scaled *in vitro* ZjuDC reaction to **2** for further purification and structure elucidation. MS and MS/MS analysis of **2** showed the loss of 46.01 Da corresponding to C-terminal oxidative decarboxylation compared to **1** and 1D and 2D NMR analysis of purified **2** confirmed the presence of a *p*-hydroxystyrylamine at the C-terminus of **2** (**Figures 3D & 3E, Figure S27-34, Table S4**). The stereochemical analysis of **2** also showed the same L-configuration of Phe1, Pro2, and Ile3 compared to cyFPIY based on Marfey’s analysis, while the C-terminal alkene is in a *cis*-configuration given its *J*-value of 7 Hz (**Figure 3E, Figures S35, Table S4**). Since this is the first example of a 2OG-dependent non-heme iron enzyme catalyzing styrylamine formation, we used steady state kinetics as an indicator for affinity and turnover values for the cyclic peptide substrate. Kinetic analysis of ZjuDC with purified cyFPIY had a K_m_ of 3.51±0.98 μM, k_cat_ = 0.67±0.10 min^-1^, and k_cat_/K_m_ = 0.19 μM^-1^ min^-1^ (**Figure 3F**). In summary, RNA-seq analysis enabled us to discover a non-heme-iron and 2OG-dependent decarboxylase catalyzing the characteristic C-terminal modification of classical CPAs and brought us one step closer to a complete biosynthesis of a classical CPA.

### Characterization of a CPA-burpitide *N*-methyltransferase from jujube

Having identified the biosynthetic enzyme responsible for styrylamine formation, we applied differential gene expression analysis of young jujube shoot tissues to identify an *N*-methyltransferase to complete the biosynthetic pathway for select 14-membered CPAs. ZjuDC was used as a bait gene for the identification of candidate methyltransferases with high correlation coefficients (**Table S3**). In addition to annotated *N*-methyltransferases, *O-*methyltransferases were used as candidate genes as they could have evolved to methylate amines or possess generalist activity allowing them to methylate both amines and alcohols^48^. Nine candidate genes encoding five *N-*methyltransferases and four *O-*methyltransferases were co-expressed with ZjuPrec-5xFPIY-ZjuBURP-fusion and ZjuDC as pEAQ-HT constructs in *Nicotiana benthamania* via agroinfiltration. Infiltrated tobacco leaves were collected seven days later, extracted with 80% methanol and analyzed by LC-MS. The leaf extracts revealed an analyte with a mass shift corresponding to the *N,N-*dimethylated product of cyFPIY-dc in the presence of candidate methyltransferase ZjuNMT (**Figure 4A**). Bioinformatic analysis predicted ZjuNMT being a SAM-dependent *O*-methyltransferase (**Table S3**). ZjuNMT was recombinantly expressed and purified from *E. coli* BL21(DE3) to validate its function (**Figure S25**). Incubation of ZjuNMT with cyFPIY-dc and SAM as a methyl donor resulted in formation of a dimethylated analyte by LC-MS/MS analysis (**Figure 4B**). Interestingly, ZjuNMT also generated mono- and trimethylated CPA products *in planta* and *in vitro* (**Figure S36**). A time course for the ZjuNMT reaction with cyFPIY substrate revealed that the dimethylation product is highest in 10 min reaction time in the presence of 1 mM SAM as this condition showed lowest amounts of other methylation side products (**Figure S36**). To structurally confirm a ZjuNMT product, cyFPIY-dc (**2**) was reacted *in vitro* at milligram-scale by ZjuNMT reaction to its *N,N*-dimethylated analog. The resulting CPA was purified and structurally characterized (**Figure 4C, Figure S37-44, Table S5**). NMR and Marfey’s analysis and PGME derivatization of the reaction product characterized the 3D structure of lotusine A with L-configuration at the amino acid α-carbons S-configuration at the crosslink-β-carbon (**Figure S45**). ZjuNMT was able to methylate both cyFPIY to *N,N*-Me_2_-cyFPIY (**4**) and cyFPIY-dc (**2**) to lotusine A, respectively, indicating its substrate promiscuity within *Ziziphus* CPA biosynthesis (**Figure 4B, 4D, & Figure S36**). A cofactor screen of ZjuNMT revealed its SAM-dependent product formation (**Figure 4D**). Therefore, ZjuNMT discovery enabled the full reconstitution of 14-membered classical CPA biosynthesis.

**Figure 4.**
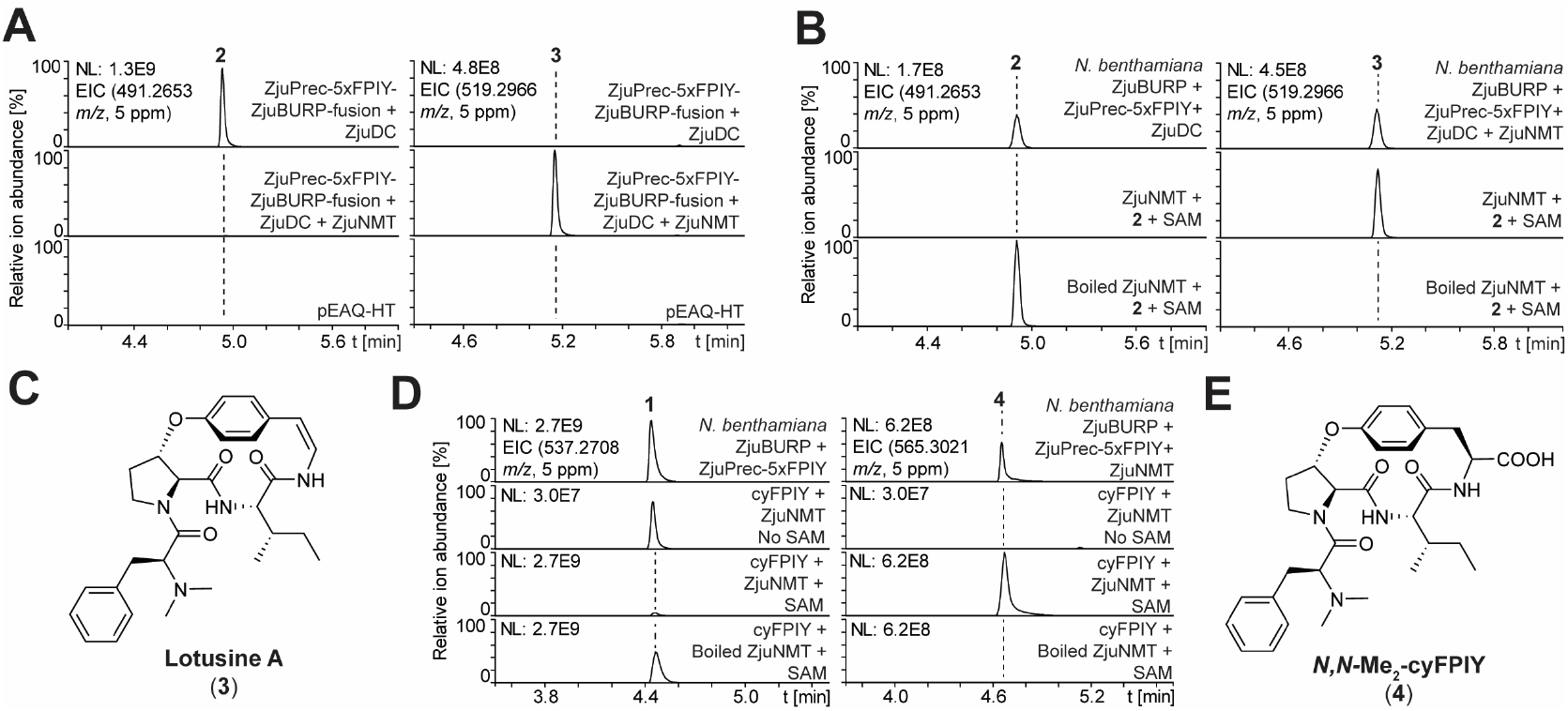
Discovery and characterization of a SAM-dependent *N*-methyltransferase in CPA biosynthesis from jujube. (**A**) EIC traces of decarboxylated cyFPIY and *N,N*-dimethylated cyFPIY-dc (lotusine A) in methanolic extracts of transgenic *N. benthamiana* leaves expressing ZjuPrec-3xFPIY-ZjuBURP and ZjuDC with or without candidate N-methyltransferase ZjuNMT for 7 days. (**B**) *In vitro* reconstitution of ZjuNMT activity in the presence of SAM. (**C**) Lotusine A structure with detected N-methyl NMR correlations. (**D**) Substrate promiscuity assay of ZjuNMT with cyFPIY as a substrate including SAM-dependence assay. (**E**) *N,N*-Me_2_-cyFPIY structure predicted by tandem mass spectrometry (**Figure S36**). Abbreviations: SAM - S-adenosylmethionine.

### Biosynthesis of adouetine X and sanjoinine A with ZjuDC and ZjuNMT

With two enzymes for terminal CPA modifications discovered, we tested if they can be applied for biosynthesis of medicinal CPAs. Two target CPAs with interesting medicinal properties are adouetine X and sanjoinine A (**Figure 1A**): adouetine X shows analgesic properties in a mouse model and sanjoinine A has anxiolytic activity by targeting GABA signaling^3,4^. To enable drug development of these CPAs, a biosynthetic route with potential for diversification is desired. Adouetine X and sanjoinine A are Leu-CPAs, i.e. their ether-crosslink connects the C-terminal phenol with a leucine-β-carbon. To enable biosynthesis of Leu-CPAs, we applied SkrBURP which is a fused burpitide cyclase from African clubmoss (*Selaginella kraussiana*). SkrBURP produces both macrocycles for mono- and bicyclic CPAs such as selanine B and selanine A, and it generates a 14-membered macrocycle in both CPA products by crosslinking a tyrosine-phenol-ring to a leucine-Cβ via an ether bond (**Figure 1A, Figure S1**). Based on their structures, adouetine X and sanjoinine A could be derived from cyLLIY (**5**) and cyFLLY (**6**) peptides by sequential C-terminal decarboxylation and *N,N*-dimethylation (**Figure 5A**). To establish biosynthetic routes to adouetine X or sanjoinine A, we used SkrBURP mutants with either LLIY or FLLY core peptides (**Figure 1, Figure S46**). To optimize cyclic peptide yields from transient expression of SkrBURP in *N. benthamiana*, we tested constructs with one or five FLLY core peptides with either a C-terminal tyrosine (FLLYxxY) or phenylalanine (FLLYxxF) of the core peptide of bicyclic selanine A (core: xLxYxxY). The core peptide FLLYxxF was designed to prohibit bicyclization and increase monocyclic product formation, i.e. the 14-membered ring of the selanine scaffold (**Figure S46**). Among tested SkrBURP constructs, SkrBURP-5xFLLYxxF generated the most cyFLLY *in planta* and both FLLYxxF constructs increased monocyclic yields compared to FLLYxxY constructs by 7-9-fold (**Figure S47**). Therefore, we designed and synthesized two *SkrBURP* variants with five core peptides of LLIYxxF or five core peptides of FLLYxxF for biosynthesis of cyLLIY and cyFLLY, respectively (**Figure S46**). Transient expression of *SkrBURP-5xLLIYxxF* and *SkrBURP-5xFLLYxxF* in *N. benthamiana* yielded analytes with the expected *m/z* values of cyLLIY and cyFLLY (**Figure 5B**). To confirm their structures, both peptides were purified from 1-1.2 kg of transgenic *N. benthamiana*. NMR and Marfey’s analysis showed the predicted CPA intermediates towards sanjoinine A and adouetine X (**Figures S48-S56 & S57-S65, Tables S6 & S7**). Co-expression of *SkrBURP-5xLLIYxxF* or *SkrBURP-5xFLLYxxF* with both decarboxylase ZjuDC and *N*-methyltransferase ZjuNMT in *N. benthamiana* yielded analytes in methanol extracts of leaves seven days after agroinfiltration which matched authentic standards of sanjoinine A isolated from *Ziziphus cambodiana* and adouetine X isolated from *Ceanothus americanus* (**Figure 5C, Figures S66-S73 & Figures S74-S83, Tables S8 & S9**). We further tested if ZjuDC and ZjuNMT could be applied for *in vitro* biosynthesis of our target CPAs. Using cyLLIY or cyFLLY isolated from transgenic *N. benthamiana* as a starting point, ZjuDC and ZjuNMT could convert the substrates to the desired products in sequential reactions or in a one-pot-reaction (**Figure 5C**) indicating that the discovered enzymes from *Z. jujuba* have potential for biocatalytic applications towards the production of other CPAs than lotusine A. Finally, we tested ZjuDC and ZjuNMT for diversification of sanjoinine A and adouetine X *in planta*. Transient expression of SkrBURP constructs with all proteinogenic analogs of either FLxY or LlxY cores distributed in four 5xcore constructs resulted in the detection of 17 cyFLxY and 18 cyLLxY analogs in methanol extracts of transgenic *N. benthamiana* leaves (**Figure 5D, Figures S84 & S85**). These cyclic CPA intermediates could then be intermediates for ZjuDC and ZjuNMT via *in planta* co-expression experiments. Seven sanjoinine analogs with hydrophobic residues (Ala, Ile, Leu, Val), threonine or serine at position 3 and four adouetine X analogs with hydrophobic residues (Ile, Leu, Val) or threonine at position 3 were generated (**Figure 5D, Figures S86 & S87**). Therefore, a total of ∼30% of possible analogs could be generated via the *Ziziphus* CPA biosynthetic enzymes in *N. benthamiana*.

**Figure 5.**
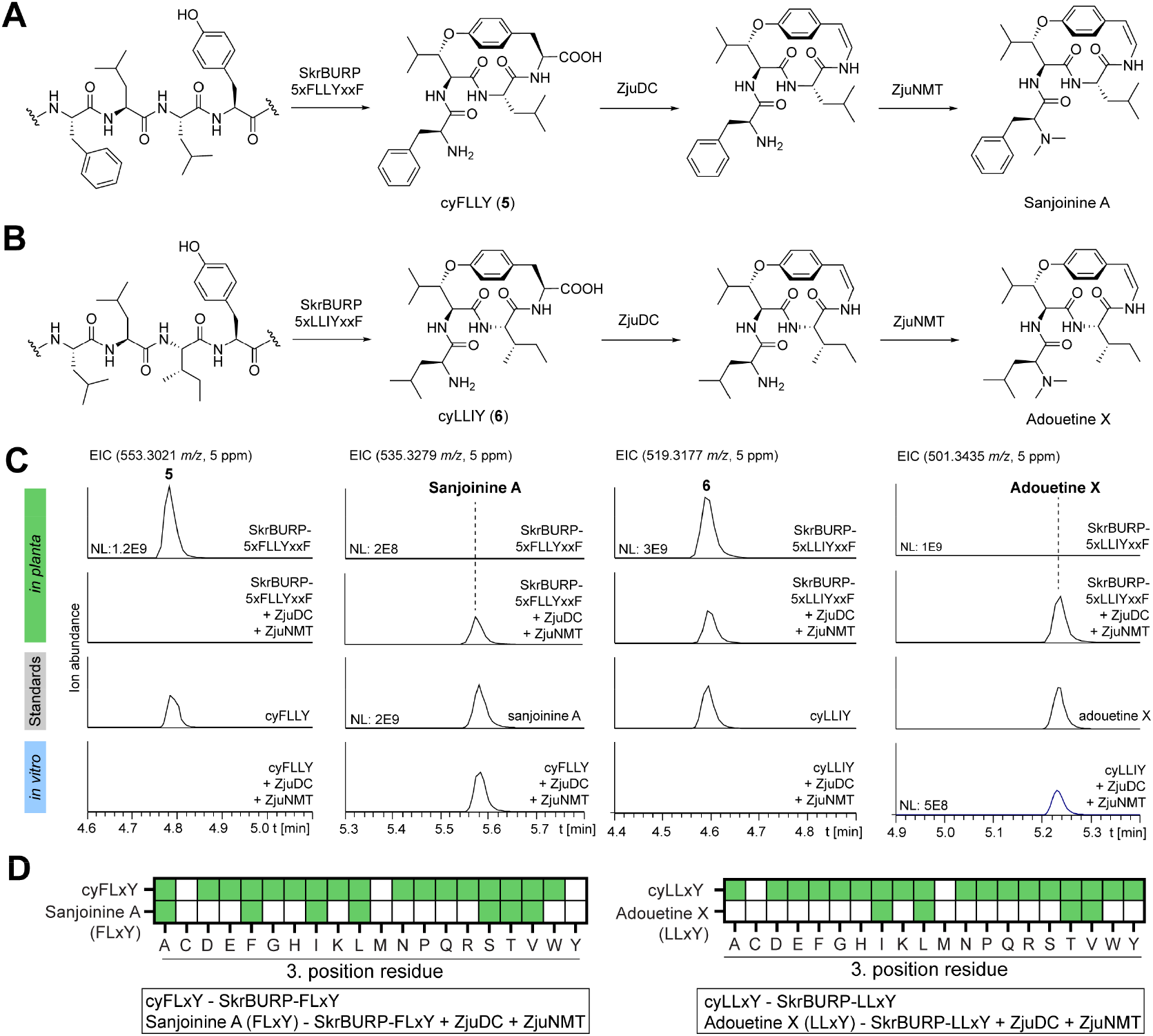
Biosynthesis of medicinal CPAs via ZjuDC and ZjuNMT. (**A**) Biosynthetic route to sanjoinine A. (**B**) Biosynthetic route to adouetine X. (**C**) In planta and in vitro biosynthesis of sanjoinine A and adouetine X from SkrBURP-5xFLLY or SkrBURP-5xLLIY products, respectively. (**D**) *In planta* diversification of sanjoinine A and adouetine X at position 3 via SkrBURP, ZjuDC and ZjuNMT in *Nicotiana benthamiana*. Green color indicates detected analytes, white color indicates no detected analytes.

## Discussion

Over the last decades, multiple synthetic strategies have been developed to generate strained macrocycles and C-terminal enamides for the production of classical 14-membered cyclopeptide alkaloids^8,9^. Here, we present a biosynthetic route to classical cyclopeptide alkaloids with leucine and proline ether crosslinks including analgesic adouetine X and anxiolytic sanjoinine A. The elucidated CPA biosynthetic pathway from *Ziziphus jujuba* features ribosomal synthesis of a repetitive precursor peptide, a split burpitide cyclase for 14-membered core peptide macrocyclization, one or more putative proteases for core peptide proteolysis, a non-heme-iron and 2OG-dependent enzyme for C-terminal oxidative decarboxylation and a SAM-dependent methyltransferase for α-*N*-methylation (**Figure S88**). Compared to synthetic routes, which can be up to 20 steps long such as for sanjoinine A^49^, the elucidated biosynthetic pathway includes only five steps and enables diversification at the non-aromatic crosslinking residue by exchange of cyclases with different substrate specificity and at non-crosslinked residues by core peptide mutagenesis.

The first step in CPA biosynthesis is the formation of the ether crosslink by a burpitide cyclase (**Figure S88**). As a member of the BNM2 BURP-domain-containing proteins, ZjuBURP is a split burpitide cyclase with a separate precursor peptide. Interestingly, ZjuBURP has a precursor gene co-localized in the *Z. jujuba* genome on chromosome 9 for lotusine A as predicted previously^12^ but it also appears to be active on precursor peptide ZjuPrec-FIPFY which is encoded on chromosome 4 without a co-localized BURP domain-containing gene. This is an example of a split burpitide precursor gene not being co-localized with a burpitide cyclase gene but being co-expressed (**Table S3**). Fusion of ZjuBURP with truncated ZjuPrec-FPIY generated higher cyclic peptide yields in transgenic *N. benthamiana* which could be due to higher catalytic efficiency because of substrate proximity to the burpitide cyclase active site, less proteolytic degradation of the core peptide domain in *N. benthamiana*, or higher precursor peptide expression as a fused construct given higher relative OD_600_ during agroinfiltration. Challenges of split burpitide cyclases as biocatalysts for the formation of CPA biosynthetic intermediates are their low protein yields for *in vitro* reactions. As exemplified in our work, this issue can be addressed by three metabolic engineering strategies *in planta*^50^: (a) identification of other CPA-burpitide cyclases such as SkrBURP by mining plant omics data^11,51,52^, (b) by optimization of core peptide repeats on precursor peptides^53^, and (c) reduction of shunt products such as bicyclic CPAs in SkrBURP. The second step in CPA biosynthesis is proteolysis of cyclic core peptides from modified precursor peptides. The identity of proteases involved in either CPA source plants such as *Ziziphus jujuba* or heterologous host plant *Nicotiana benthamiana* is unknown but their identification and co-expression with CPA biosynthetic genes could further boost the yields of CPAs *in planta* and enable their production from precursor peptides *in vitro*^54–57^.

The third step in CPA biosynthesis is the oxidative decarboxylation of cyclic peptide intermediates by non-heme-iron and 2OG-dependent enzyme ZjuDC. This enzyme is an example of plant non-heme-iron enzymes as a potential source for new biocatalysts^58,59^ as it introduces a new desaturation reaction to this enzyme family which involves the loss of a C-terminal peptide carboxy functionality to yield a C-terminal cyclopeptide enamide^60^. Non-heme-iron and 2OG-dependent enzymes have been described for the oxidative decarboxylation of amino acid-derived substrates such as tyrosine isonitrile to yield phenol vinyl isonitrile during rhabduscin biosynthesis^61,62^. Another example of a non-heme-iron and 2OG-dependent enzyme catalyzing decarboxylation-assisted olefin formation is ethylene-forming enzyme which generates ethylene from 2OG^63^. ZjuDC represents a new enzyme family for peptide oxidative decarboxylation in RiPP biosynthesis (**Figure 1B**). Although ZjuDC shows a low k_cat_, it could offer a starting point for biocatalytic decarboxylation of cyclic peptides given its solubility from *E. coli* expression and a potential wider substrate scope given its catalytic activity with cyFPIY, and 11 analogs of cyFLxY and cyLLxY substrates. Structural and mechanistic studies of ZjuDC can inform improvements for a broader substrate scope beyond 14-membered macrocycles. The oxidative decarboxylation mechanism of ZjuDC could begin with the activation of dioxygen by the iron cofactor to form a high-valent Fe(IV)-oxo intermediate^64^, similar to other enzymes in this superfamily^65^. Three candidate mechanisms for ZjuDC are presented in **Figure S89** based on different Fe(IV)-oxo intermediates: radical hydroxylation followed by dehydration and decarboxylation, single electron transfer to the substrate leading to a cation, and diradical formation proceeding homolytic cleavage^66^.

The final step in the presented biosynthesis of 14-membered CPAs is the α-*N*-methylation of the decarboxylated CPA intermediate by SAM-dependent ZjuNMT. This methyltransferase also offers biocatalytic access to *N*-methylated CPAs with reaction products including mono-, di- and tri-*N*-methylated peptides. The reaction time of ZjuNMT was optimized to reach the highest yields of the desired dimethylation product. The detection of trimethylated products as a ZjuNMT side product analyte in *in planta* biosynthesis suggests that *in vitro* reactions might be a more controllable applications of ZjuNMT towards a desired *N*-methylated CPA. Interestingly, trimethylated N-termini have been reported before in lanthipeptide biosynthesis from marine symbiotic bacteria^27^. ZjuDC and ZjuNMT are the first characterized enzymes involved in terminal posttranslational modifications of burpitides and represent two diversification strategies of these RiPPs in plants beyond the macrocyclization step^52^ as they accept core peptide substrates with different crosslinking and non-crosslinking residues. A putative *N*-methyltransferase in CPA biosynthesis was proposed based on *in planta* activity in jujube tissue, which was co-localized with a BURP-domain protein on chromosome 6^67^. Our characterized ZjuNMT is localized on chromosome 4 whereas ZjuDC is localized on chromosome 1. With ZjuBURP and ZjuPrec-FPIY localized on chromosome 9, the described CPA biosynthetic pathway is not entirely clustered on the jujube genome. Further research is needed into the nature and genome location of missing enzymes to complete biosynthesis of 13-membered CPAs such as jubanine K.

The characterization of a 14-membered CPA biosynthetic pathway sets up more scalable biosynthesis or chemoenzymatic synthesis of bioactive CPAs such as adouetine X and sanjoinine A for drug development. The introduced enzymes ZjuDC and ZjuNMT further could be explored for biocatalytic decarboxylation and methylation, respectively, of cyclic peptide substrates beyond CPAs^68,69^. They could also serve as starting points for bioinformatic discovery of homologs in the plant kingdom with different substrate scopes^52,70^. Ultimately, this work provides a biosynthetic alternative to CPA synthesis for further chemical and biological exploration of these RiPP natural products.

## Data availability

*Ziziphus jujuba* RNA-seq data has been deposited in the NCBI sequence read archive^71^ under Bioproject accession PRJNA1276874. Gene sequences were deposited in GenBank (ZjuBURP - PV929269, ZjuNMT PV929270, ZjuPrec-FIPFY - PV929271). LC-MS data was deposited to GNPS-MassIVE^72^ under accession MSV000098504 (Password: 4aUgs78Bsvd1V0ar). Synthetic gene sequences used in this study are listed in **Data S2**.

## Materials and Methods Materials

All chemicals and reagents were purchased from Thermo Fisher Scientific Inc. if not otherwise noted. LC-MS solvents were Optima LC-MS-grade solvents (Fisher Scientific Inc.). Sanjoinine A isolated from *Ziziphus cambodiana* Pierre. (Cat# HY-119637) was purchased from MedChemExpress. Synthetic gene fragments were obtained from Twist Biosciences Inc. *Ceanothus americanus* dried roots were purchased from Starwest Botanicals. Deoxynucleotide primers were purchased from Integrated DNA Technologies Inc. LC-MS analysis was performed on a Thermo QExactive Orbitrap mass spectrometer coupled to a Vanquish Ultra high performance liquid chromatography instrument. LC-MS analysis was performed on a Thermo QExactive Orbitrap mass spectrometer coupled to a Vanquish Ultra high performance liquid chromatography instrument. LC-MS data was analyzed with QualBrowser from Thermo Xcalibur software package (v4.3.73.11, Thermo Scientific). Preparative and semi-preparative high performance liquid chromatography (HPLC) was performed on a Shimadzu HPLC with two binary LC20-AP pumps, a DGU-403 Degasser, an SPD-20A ultraviolet-visible light detector, and a FRC-10A fraction collector. NMR analysis was done on a Bruker Ascend 800 MHz NMR spectrometer with a 5 mm Triple Resonance Inverse Detection TCI CryoProbe. NMR data was analyzed with MestReNova (v14.0.0, Mestrelab Research).

### Plant cultivation

*Nicotiana benthamiana* were grown from seeds in Premier Horticulture Pro Mix HP High Porosity with Mycorrhizae for plant growth under plant growth lights with a 16 h light/8 h dark cycle for 4-12 weeks at room temperature. *N. benthamiana* seedlings were transferred after one week to larger pots with the same soil and added Osmocote slow-release fertilizer. Young stem and leaves of *Ziziphus jujuba* Mill. var. Dongzao for RNA-seq and metabolomics were collected on June 6^th^, 2023 from New Mexico State University Sustainable Agriculture Science Center at Alcalde, NM 87511. The trees are part of the jujube cultivar trial planted in 2015 at Alcalde, NM (lat. 36°05’27.94” N, long. 106°03’24.56” W, elevation, 1730 m).

### Transcriptomics

Young stem and leaf tissue from shoots of three different *Ziziphus jujuba* trees were frozen in liquid nitrogen and ground with mortar and pestle. Total RNA was isolated from ground plant tissue samples with the QIAGEN RNeasy Plant Mini kit with highest RNA yields in the RLC buffer. RNA-seq library was prepared with NEBNext Poly(A) mRNA Magnetic Isolation Module and the NEBNext Ultra II RNA Library Prep Kit for Illumina. RNA-seq data was generated by Illumina sequencing in paired-end format (2×150 bp) on a NovaSeq 6000 Instrument S4 flow cell (Azenta Genewiz Inc).

RNA-seq data analysis was performed on the Great Lakes High-Performance Computing Cluster at the University of Michigan, Ann Arbor. Forward and reverse reads of raw fastq-files were trimmed by Trimgalore (v0.6.7) with default settings and forward and reverse reads of all trimmed files were combined respectively. Combined paired fastq-files were *de novo* assembled by SPAdes (v3.15.5)^73,74^. The resulting *Ziziphus jujuba de novo* transcriptome fasta file was formatted for BLAST database conversion (truncation of fasta headers <51 letters) and uploaded to an internal Sequenceserver (v3.1.0)^75^ for BLAST+ analysis (v2.16.0)^42^. Target protein sequences such as ZjuPrec-FPIY were queried in the *de novo* transcriptome by tblastn search (v2.16.0) on Sequenceserver (v3.1.0) with the following BLAST parameters: e-value 1e-5, matrix BLOSUM62, gap-open 11, gap-extend 1, filter L. For transcript quantification, kallisto (v0.46.0)^76^ was used to (1) index the combined *de novo Z. jujuba* transcriptome and (2) count transcripts in each RNA-seq dataset. TPM (transcripts per million reads) values were combined for all six samples in a table. For identification of co-expressed genes to a given bait gene, the transcript of a bait gene was determined by tblastn search on Sequenceserver and the identified transcript was used as a bait transcript to calculate Pearson coefficients for TPM values of all other transcripts in the *de novo* transcriptome relative to the TPM values of the bait transcript in Microsoft Excel (2023). Transcripts were then sorted by decreasing Pearson coefficient and the top 131,551 transcripts were generated in a fasta file with seqkit2^77^ and annotated with EnTAP (v1.0.0)^78^ using UniProt/SwissProt^79^ as a database. For burpitide cyclase gene search, we searched BURP-domain-containing proteins among the top co-expressed genes relative to *ZjuPrec-FIPFY* transcript. For decarboxylase gene search, we used ZjuBURP as a bait gene and searched for co-expressed transcripts encoding annotated oxidative genes such as CYP450s, type III peroxidases, laccases, BURP-domain-containing proteins, and non-heme-iron and 2OG-dependent oxygenases. For *N*-methyltransferase gene discovery, we used ZjuDC as a bait gene and searched the top co-expressed transcripts for genes encoding annotated *O*- and *N*-methyltransferases.

### Pathway elucidation in *Nicotiana benthamiana*

Young stem total RNA of *Z. jujuba* was used as a template to generate cDNA with the Thermo Scientific™ Maxima H Minus First Strand cDNA Synthesis Kit. ZjuPrec-FIPFY was amplified from the young stem *Z. jujuba* cDNA by cloning PCR with forward primer tgcccaaattcgcgaccggtATGAAAAGTTTCTTTGCGCTTC and with reverse primer ccagagttaaaggcctcgagTTACTGATGATCAGGAGAAACAAC and cloned via Gibson cloning into pEAQ-HT vector after it was linearized with BshTI (Fisher Scientific) and XhoI (Fisher Scientific). Resulting plasmids were sequenced by Nanopore sequencing (Plasmidsaurus Inc.).

Synthetic genes (*ZjuPrec-5xFPIY, ZjuBURP, ZjuDC* (GenBank: XM_048464910.1), *ZjuNMT, ZjuPrec-5xFPIY-ZjuBURP, SkrBURP-1xFLLY, SkrBURP-1xFLLY*..*Y, SkrBURP-5xFLLY, SkrBURP-5xFLLY*..*Y, SkrBURP-5xLLIY*..*F, SkrBURP-5xLLxY*..*F-ACDEF, SkrBURP-5xLLxY*..*F-GHIKL, SkrBURP-5xLLxY*..*F-MNPQR, SkrBURP-5xLLxY*..*F-STVWY, SkrBURP-5xFLxY*..*F-ACDEF, SkrBURP-5xFLxY*..*F-GHIKL, SkrBURP-5xFLxY*..*F-MNPQR, SkrBURP-5xFLxY*..*F-STVWY*) and cloned genes (*ZjuPrec-FIPFY*) (**Data S2**) were transformed into electrocompetent *Agrobacterium tumefaciens* LBA4404 and clones were selected on sterile YM-agar medium (0.4 g yeast extract, 0.2 g magnesium sulfate heptahydrate, 0.1 g sodium chloride, 0.5 g potassium phosphate (dibasic), adjusted to 1 liter with deionized water) with 50 µg/mL rifampicin, 50 µg/mL kanamycin, and 100 µg/mL streptomycin. For small-scale expression experiments, a colony of each construct was transferred into a 5 mL culture of liquid YM medium containing 50 µg/mL rifampicin, 50 µg/mL kanamycin, and 100 µg/mL streptomycin. Each culture was grown 24-36 h at 30 °C and 220 rpm and then transferred to a 25 mL culture of liquid YM medium with 50 µg/mL rifampicin, 50 µg/mL kanamycin, and 100 µg/mL streptomycin in a 50 mL Bioreactor tube (Cell treat, Cat# 229475) and incubated for 24 h at 30 °C and 220 rpm. *A. tumefaciens* cells were then centrifugated at 3000 x *g* and 25 °C and resuspended in MMA medium (10 mM MES (pH 5.6), 10 mM magnesium chloride, 150 µM acetosyringone) to a final OD_600_ of 0.8. For pathway elucidation experiments, resuspensions of *A. tumefaciens* cultures of a single gene were combined in equal ratios. Each resuspension or resuspension mixture was incubated for 30 min at 25 °C and then infiltrated with a 1 mL syringe into the bottom of leaves of 4-6 week-old *N. benthamiana* plants. Seven days after infiltration, *N. benthamiana* leaves were collected. For qualitative experiments, 0.3 g of fresh weight transgenic *N. benthamiana* leaves were transferred to an MP Biomedicals Tissuelyzer tube (2 mL, Cat#: 5076200, 5068002) with 10-15 2.3 mm Zirconia silica beads (Cat#: NC0419764), frozen at -80 °C for at least 30 min, and ground in a MP Biomedicals FastPrep-24-5G Tissuelyzer for 60 s at 6 m/s. 1 mL of 80% methanol was added to the ground plant material in the Tissuelyzer tube and ground another 20 s in the Tissuelyzer at 6 m/s. The tubes were then incubated at 60 °C for 10 min in a waterbath, centrifugated at 16000 x *g* at room temperature for 5 min, and the supernatant was filtered with Cytvia syringeless LC-MS filters (UN503NPEORG). The plant methanol extracts were then transferred to LC-MS vials analyzed by LC-MS/MS with the following parameters: LC - Phenomenex Kinetex® 2.6 μm C18 reverse phase 100 Å 150 x 3 mm LC column; LC gradient, solvent A, 0.1% formic acid; solvent B, acetonitrile (0.1% formic acid); 0 min, 10% B; 5 min, 60% B; 5.1 min, 95% B; 6 min, 95% B; 6.1 min, 10% B; 9.9 min, 10% B; 0.5 mL/min, 30 °C; MS, positive ion mode; full MS, resolution 70,000; mass range 400–1,200 *m/z*; dd-MS2 (data-dependent MS/MS), resolution 17,500; AGC target 1 x 10e^5^, loop count 5, isolation width 1.0 *m/z*, collision energy 25 eV and dynamic exclusion 0.5 s. Sample numbers were *n* = 3 (biological replicates) for each source plant extraction and each transient gene expression experiment in *N. benthamiana*. The negative control of the transient gene expression experiment in *N. benthamiana* was an empty pEAQ-HT vector agroinfiltration experiment. For *Z. jujuba* chemotyping, young stem and leaf tissue samples were prepared and analyzed for LC-MS chemotyping like *N. benthamiana* leaf samples except that stem samples were ground in metal homogenizer tubes with a 0.25 in. steel ball (2 mL, MP Biomedicals, Cat#: MP116991006).

### Peptide optimization and diversification in transgenic *Nicotiana benthamiana*

SkrBURP constructs for peptide optimization (*ZjuPrec-5xFPIY, ZjuBURP, ZjuPrec-5xFPIY-ZjuBURP-fusion, SkrBURP-1xFLLY, SkrBURP-1xFLLY*..*Y, SkrBURP-5xFLLY, SkrBURP-5xFLLY*..*Y, SkrBURP-5xLLIY*..*F*) were expressed transiently in *Nicotiana benthamiana* for 7 days with 4-5 replicates. Infiltrated leaves were frozen at -80 °C and dried by freeze-drying. 50 mg of dried leaves were extracted as described above with 1 mL 80% methanol. Methanol extracts were analyzed as above for peptide chemotyping except for using FullMS mode. Target peptide peak AUC values were characterized in QualBrowser and plotted and analyzed in GraphPad Prism (v10.4.2).

SkrBURP constructs for cyFLxY and cyLLxY diversification (*SkrBURP-5xLLxY*..*F-ACDEF, SkrBURP-5xLLxY*..*F-GHIKL, SkrBURP-5xLLxY*..*F-MNPQR, SkrBURP-5xLLxY*..*F-STVWY, SkrBURP-5xFLxY*..*F-ACDEF, SkrBURP-5xFLxY*..*F-GHIKL, SkrBURP-5xFLxY*..*F-MNPQR, SkrBURP-5xFLxY*..*F-STVWY*) were expressed transiently in *N. benthamiana* for 7 days with 3 replicates. Samples were prepared from fresh weight tissue as described for peptide chemotyping above. Target cyFLxY and cyLLxY peptides were characterized in QualBrowser based on MS1 and MS2 data. For CPA diversification, each SkrBURP-5xLLxY or SkrBURP-5xFLxY construct was co-expressed with ZjuDC and ZjuNMT transiently in *N. benthamiana* for 7 days with 3 replicates. Samples were prepared from fresh weight tissue as described for peptide chemotyping above. Target cyFLxY CPA and cyLLxY CPA peptides were characterized in QualBrowser based on MS1 and MS2 data.

### Peptide isolation

For scaled production of cyclopeptide alkaloid intermediates cyFPIY, cyFLLY, or cyLLIY, *Agrobacterium tumefaciens* LBA4404 was transformed with expression plasmids pEAQ-HT-*ZjuPrec-5xFPIY-ZjuBURP-fusion*, pEAQ-HT-*SkrBURP-5xFLLY*, or pEAQ-HT-*SkrBURP-5xLLIY* as described above for pathway expression^80^. A 25 mL YM medium liquid starter culture with 50 µg/mL rifampicin, 50 µg/mL kanamycin, and 100 µg/mL streptomycin was inoculated with a colony of a target gene construct transformant and incubated for 24-36 h at 30 °C and 220 rpm in a 50 mL Bioreactor tube. The culture was then used to inoculate a 1 L YM liquid medium culture with 50 ug/mL rifampicin, 50 ug/mL kanamycin, and 100 ug/mL streptomycin and incubated for 24 h at 30 °C and 220 rpm. The culture was then centrifugated at 3000 x *g* at 25 °C for 30 min and resuspended in MMA medium to a final OD_600_ of 1.5. 4-8 week-old *Nicotiana benthamiana* plants were infiltrated with a transformant of each construct in the bottom leaves by syringe infiltration at scale (total fresh weight: cyFPIY - 1 kg, cyFLLY - 1.1 kg, cyLLIY - 1.2 kg). Transgenic *N. benthamiana* leaves were frozen and ground in a food processor. Ground plant material was extracted with methanol (8 L for 1 kg fresh weight plant material) by incubation for 16 h at 37 °C and 160 rpm. Methanol extracts were filtered with a silica filter and dried *in vacuo*. Dried methanol extract of each construct was resuspended in 2 L deionized water and partitioned twice with 2 L hexanes, twice with 2 L of ethyl acetate and finally extracted with 2 L of n-butanol. n-butanol extract of each construct was dried *in vacuo* and separated by Sephadex LH20 (Cytiva, Cat#: 17009001, 300 g) liquid chromatography in a Chemglass 8 x 60 cm column, 45/50 (Cat#: CG118931) glass column with a starting mobile phase of 10% methanol. During Sephadex LH20 LC, methanol content of the mobile phase increased by 10% every 500 mL. Sephadex LH20 LC fractions were analyzed for the target cyclic peptide by LC-MS with the following method: Injection volume 2 µL, LC - Phenomenex Kinetex® 2.6 μm C18 reverse phase 100 Å 50 x 3 mm LC column; LC gradient, solvent A, 0.1% formic acid; solvent B, acetonitrile (0.1% formic acid); 0 min: 10% B; 2.5 min: 95% B; 3 min: 95% B; 3.1 min: 10% B; 5 min: 10% B; 0.5 mL/min, 30 °C; MS, positive ion mode; full MS, resolution 70,000; mass range 400–1,200 *m/z*, dd-MS2 (data-dependent MS/MS), resolution 17,500; AGC target 1e5, loop count 5, isolation width 1 *m/z*, collision energy 25 eV and dynamic exclusion 0.5 s. LC fractions with the target peptide were combined, dried *in vacuo*, and subjected to two steps of preparative HPLC with the following parameters: Phenomenex Kinetex® 5 µm C18 100 Å LC Column 150 x 21.2 mm, LC gradients: solvent A – 0.1% trifluoroacetic acid (TFA), solvent B – acetonitrile (0.1% TFA), 1. step: 0 min: 10% B, 1 min: 10% B, 36 min: 50% B, 39 min: 95% B, 42 min: 95% B, 42.5 min: 10% B, 60.1 min: 10% B. 2. step (cyFPIY): 0 min: 20% B, 1 min: 20% B, 36 min: 40% B, 39 min: 95% B, 42 min: 95% B, 42.5 min: 20% B, 60.1 min: 20% B. 2. step (cyFLLY): 0 min: 20% B, 1 min: 20% B, 36 min: 40% B, 39 min: 95% B, 42 min: 95% B, 42.5 min: 20% B, 60.1 min: 20% B. 2. step (cyLLIY): 0 min: 20% B, 1 min: 20% B, 36 min: 40% B, 39 min: 95% B, 42 min: 95% B, 42.5 min: 20% B, 60.1 min: 20% B. Preparative HPLC fractions were analyzed for target cyclic peptides as described above. Target peptide fractions were combined and dried *in vacuo*. Each target peptide fraction was subjected to two steps of semipreparative HPLC with the following parameters: Kinetex® 5 µm C18 100 Å LC Column 250 x 10 mm, 1. step: solvent A – 0.1% TFA, solvent B – acetonitrile (0.1% TFA), LC gradient (cyFPIY): 0 min: 25% B, 1 min: 25% B, 21 min: 35% B, 22 min: 95% B, 24.5 min: 95%, 25 min: 25% B, 45 min: 25% B. LC gradient (cyFLLY): 0 min: 30% B, 1 min: 30% B, 21 min: 40% B, 22 min: 95% B, 24.5 min: 95%, 25 min: 30% B, 45 min: 30% B, LC gradient (cyLLIY): 0 min: 29% B, 1 min: 29% B, 21 min: 34% B, 22 min: 95% B, 24.5 min: 95%, 25 min: 29% B, 45 min: 29% B. 2. step: solvent A – 0.1% TFA, solvent B – methanol (0.1% TFA), LC gradient (cyFPIY): 0 min: 40% B, 1 min: 40% B, 21 min: 60% B, 22 min: 95% B, 24.5 min: 95%, 25 min: 40% B, 45 min: 40% B. LC gradient (cyFLLY): 0 min: 58% B, 1 min: 58% B, 21 min: 58% B, 22 min: 95% B, 24.5 min: 95%, 25 min: 58% B, 45 min: 58% B. LC gradient (cyLLIY): 0 min: 57% B, 1 min: 57% B, 21 min: 57% B, 22 min: 95% B, 24.5 min: 95%, 25 min: 57% B, 45 min: 57% B. Semipreparative HPLC fractions were analyzed for target cyclic peptides as described above. Target peptide fractions were combined and dried *in vacuo*. cyFPIY (5 mg) was a white powder, cyFLLY (4.5 mg) was a white powder, cyLLIY (6.7 mg) was a white powder.

### Stereochemical analysis

About 400 μg of each purified cyclopeptide alkaloid was hydrolyzed in 600 μL of 6 M HCl in a sealed thick-walled reaction vessel^81^. The sample was heated at 110 °C and stirred overnight for 12 h. Subsequently, the hydrolysate was concentrated to dryness under nitrogen gas and redissolved in 100 μL water. Then, 100 μL of 1 M NaHCO_3_ and 100 μL of a 1% acetone solution of 1-fluoro-2,4-dinitrophenyl-5-L-alanine amide (L-FDAA) were added to the solution. The reaction mixture was incubated at 40 °C for 1 h and then quenched by adding 100 μL of 1 M HCl. For the preparation of L-FDAA-amino acid standard derivatives, 50 mM of each amino acid (D/L-Phe, D/L-Pro, D/L-Ile, D/L-*allo*-Ile, D/L-Leu and D/L-Tyr) dissolved in water (50 μL) was treated with 1 M NaHCO_3_ (20 μL) and 1% l-FDAA (100 μL) at 40 °C for 1 h, respectively. After the reaction, the solution was quenched with 1 M HCl (20 μL) and diluted with acetonitrile (810 μL) for LC-MS analysis. LC-MS/MS analysis parameters were as follows: injection volume 2 µL; LC, Phenomenex Kinetex 2.6 μm C18 reverse phase 100 Å 150 x 3 mm LC column; LC gradient: solvent A, 0.1% formic acid; solvent B, acetonitrile (0.1% formic acid); 0 min, 10% B; 40 min, 60% B; 41 min, 95% B; 44 min, 95% B; 45 min, 10% B, 50 min, 10% B, 0.5 mL/min, 30 °C; MS, positive ion mode; full MS, resolution 70,000, mass range 100–1,000 *m/z*.

To determine the configuration of Ile, the C3-Marfey’s method was employed^82^. Hydrolysis and derivatization were conducted as described above, with the analytical conditions modified as follows: injection volume 5 µL; LC, Agilent Zorbax StableBond C3 5 μm 150 x 4.6 mm LC column; LC gradient: solvent A, Water with 5% acetonitrile and 0.05% formic acid; solvent B, methanol with 5% acetonitrile and 0.05% formic acid; 0 min, 15% B; 53 min, 60% B; 54 min, 95% B; 59 min, 95% B; 60 min, 15% B, 70 min, 15% B, 0.7 mL/min, 50 °C; MS, positive ion mode; full MS, resolution 70,000, mass range 100–1,000 *m/z*. The configurations of the amino acid residues were determined by comparing their retention times with those of FDAA derivatives of amino acid standards.

To determine the configuration of PheNMe_2_ and LeuNMe_2_, PGME derivatization was performed^41^. Hydrolysis was carried out as described for Marfey’s analysis. After hydrolysis, the resulting peptide was dissolved in anhydrous tetrahydrofuran (1 mL), to which (S)-(+)-2-phenylglycine methyl ester (PGME, 2 mg) was added. N-(3-Dimethylaminopropyl)-N′-ethylcarbodiimide (EDC, 1.2 µL) was then added, and the reaction mixture was stirred at 25 °C for 2 h. The reaction was quenched by the addition of methanol (500 µL), concentrated under a stream of nitrogen, and the residue was redissolved in methanol (1 mL). For the preparation of PGME–amino acid standard derivatives, D-/L-PheNMe2 and D-/L-LeuNMe2 (1 mg each) were dissolved in anhydrous tetrahydrofuran (1 mL). To each solution, (S)-(+)-2-phenylglycine methyl ester (PGME, 2 mg) and N-(3-dimethylaminopropyl)-N′-ethylcarbodiimide (EDC, 1.2 µL) were added. The reaction mixtures were stirred at 25 °C for 2 h, quenched by addition of methanol (500 µL), concentrated under a stream of nitrogen, and the residues were redissolved in methanol (1 mL) for LC-MS analysis. LC-MS/MS analysis parameters: injection volume 5 µL; LC, Phenomenex Kinetex 2.6 μm C18 reverse phase 100 Å 150 × 3 mm LC column; LC gradient: solvent A, 0.1% formic acid; solvent B, acetonitrile (0.1% formic acid); 0 min, 10% B; 40 min, 50% B; 41 min, 95% B; 44 min, 95% B; 45 min, 10% B, 50 min, 10% B, 0.5 mL/min, 30 °C; MS, positive ion mode; full MS, resolution 70,000, mass range 100–1,000 m/z.

### Jubanine K isolation

Dried stems of *Ziziphus jujuba* (2.5 kg) were ground into powder and extracted with methanol (8 L) at 37 °C with shaking at 160 rpm for 16 h. The crude extract was resuspended in 2 L of deionized water, and the pH was adjusted to 2.5 using 2 M HCl. The acidified aqueous phase was extracted with ethyl acetate (3 x 2 L), and the organic layers were discarded to remove non-target components. The aqueous phase was then basified to pH 9.5 using 2 M NaOH and subsequently extracted with chloroform (3 x 2 L). The combined chloroform extracts were concentrated under reduced pressure to afford a residue for further purification. The chloroform fraction was subjected to Sephadex LH20 chromatography (60 x 8 cm) using a 1:1 mixture of dichloromethane and methanol. Fractions were analyzed by LC-MS with the following parameters: injection volume, 2 µL; column, Phenomenex Kinetex® 2.6 µm C18 100 Å (50 x 3 mm); solvent A, 0.1% formic acid in water; solvent B, acetonitrile (0.1% formic acid); 0.5 mL/min, 30 °C; gradient: 0 min, 5% B; min, 95% B; 3.0 min, 95% B; 3.1 min, 5% B; 5.0 min, 5% B. MS settings: positive ion mode; full MS resolution, 35,000; mass range, 400–1,200 *m/z*; dd-MS^2^ resolution, 17,500; collision energy, 25 eV; dynamic exclusion, 0.5 s. Jubanine K-containing fractions were combined and then separated by one round of preparative HPLC and one round of semipreparative HPLC with the following LC condition: PrepHPLC: Phenomenex Kinetex® 5 µm C18 100 Å LC Column 150 x 21.2 mm, solvent A - 0.1% TFA in water, solvent B - acetonitrile (0.1% TFA), LC gradients: 7.5 mL/min, 0 min: 10% B, 1 min: 10% B, 36 min: 70% B, 39 min: 95% B, 42 min: 95% B, 42.5 min: 10% B, 60.1 min: 10% B. SemiprepHPLC: Kinetex® 5 μm C18 100 Å 250 × 10 mm column, solvent A - 0.1% TFA, solvent B - acetonitrile (0.1% TFA), 1.5 mL/min, 0 min - 43% B, 1 min - 43% B, 21 min - 48% B, 22 min - 95% B, 25 min - 95% B, 25.1 min - 20% B, 45 min - 20% B. Jubanine K (1.7 mg) was obtained as a white amorphous powder.

### ZjuDC Expression and Purification

*E. coli* BL21 (DE3) was transformed with codon-optimized pHis_8_-*ZjuDC* (pHis_8_ is pET28a with an N-terminal His_8_-tag, cleavable by tobacco etch virus (TEV) protease, **Data S2**) by heat shock and plated on Luria-Bertani (LB)-agar plates with 50 μg/mL kanamycin. A colony was used to inoculate a 10 mL starter culture of LB with 50 μg/mL kanamycin. Starter cultures grew aerobically overnight at 37°C, 200-250 rpm in a shaking incubator. The starter was then used to inoculate 1L of sterile Terrific Broth in a 2.8 L Fernbach flask. Large-scale cultures grew at 37 °C, 200-250 rpm, until absorbance (OD_600_) reached 0.4-0.7. The flasks continued shaking as the temperature was lowered to 18 °C. At 18-20 °C the OD_600_ was 0.6-1.0, and cultures were induced with 50 μM IPTG and grown overnight.

The contents were centrifuged at 4,000 x *g*, 4 °C for 30 min. The supernatant was discarded, and cell pellets were resuspended in 5 mL lysis buffer (50 mM Tris-HCl pH 8, 500 mM NaCl, 1% Tween-20, 10% glycerol, 10 mM imidazole, and 10 mM β-mercaptoethanol) per gram of wet cell pellet. Lysozyme was added in a ratio of 0.5 mg per mL of lysate, then, ∼10 U of DNaseI was added. Contents stirred at 4 °C for 1 h, then sonicated on ice for eight, 1 min pulses of 80% amplitude, 2 s on, 2 s off. The lysate was centrifuged at 19,500 x *g*, 4 °C for 45 min. The clarified lysate was purified by gravity flow immobilized metal affinity chromatography (IMAC) at 4°C. Ni-NTA resin, 2 mL, was added to a 20 mL disposable column (Bio-Rad) and equilibrated with 10 CV of lysis buffer. Clarified lysate was poured over the resin and the flow through was discarded. The resin was washed for 10 CV with wash buffer (50 mM Tris-HCl pH 8, 500 mM NaCl, 10% glycerol, 30 mM imidazole, and 10 mM β-mercaptoethanol) and eluted with 5 CV of elution buffer with 300 mM imidazole pH 8. The eluted protein was added to 3.5 kDa MWCO dialysis tubing for dialysis in 50 mM HEPES pH 7.5, 100 mM NaCl, 3 mM reduced glutathione (GSH), and 0.3 mM oxidized glutathione (GSSG) with stirring at 4 °C overnight. Following dialysis, the protein was concentrated in Amicon Ultra-15 mL 10 kDa MWCO centrifugal concentrators (Millipore Sigma) before size-exclusion chromatography (SEC) and subsequent use in assays.

ZjuDC was further purified by SEC on a Cytiva HiLoad 26/600 Superdex 200 prep grade column precalibrated with the manufacturer’s kit (Cytiva) on an ÄKTA Fast Protein Liquid Chromatography system (FPLC) at 4-8 °C. ZjuDC was eluted 50 mM HEPES pH 7.5, 100 mM NaCl in 2 mL fractions. ZjuDC eluted as a monomer with an apparent molecular weight of 35 kDa. Fractions eluting as a monomer were >95% pure by SDS-PAGE (**Figure S25**), then pooled and concentrated using an Amicon Ultra-15 mL 10 kDa MWCO centrifugal concentrator for enzyme assays. SDS-PAGE was imaged with Google Pixel 8 and annotated in PowerPoint (Microsoft, 2023). The protein concentration was estimated by OD_280_ using a Thermo Scientific NanoDrop One microvolume ultraviolet-visible light spectrophotometer based on the calculated ExPASy ProtParam extinction coefficient^83^. The yield for ZjuDC following purification and SEC ranged from 15-18 mg monomer per L of culture.

### Anaerobic *in vitro* reconstitution assay for ZjuDC

Assays were handled in a Vacuum Atmospheres, Inc. anaerobic chamber (Ragsdale laboratory, University of Michigan, Ann Arbor) containing <1 ppm O_2_. Buffers were 0.2 μm filtered and degassed with nitrogen prior to use in the anaerobic chamber. ZjuDC was buffer exchanged in the chamber with 50 mM HEPES pH 7.5, 100 mM NaCl (Amicon Ultra Centrifugal Filter, 10 kDa MWCO) before addition to assays. Assays were 50 μL total volume with final concentrations of 50 mM HEPES pH 7.5, 200 μM FeSO_4_-7H_2_O, 500 μM 2-oxoglutarate (Chem-Impex Int’l Inc.), 400 μM L-ascorbic acid sodium salt (Acros Organics), 4% dimethyl sulfoxide, and 90 μM cyFPIY substrate. Control assays were quenched with a final concentration of 555 mM formic acid for 0.5 h prior to ZjuDC enzyme addition (5 μM final concentration). The pH of the assay solution dropped from 7.40 to 2.56 in the presence of formic acid. Experimental assays had 5 μM ZjuDC added and were only quenched with the same concentration of formic acid prior to removal from the chamber. All assays were done in quadruplicate and incubated in the chamber for 26 h at 21-24 °C. The same stock solutions, final concentrations, controls, and experimental assays were replicated in triplicate in the presence of O_2_ in the Kersten lab. The O_2_ containing samples were incubated for 24 h at 21-24 °C prior to analysis.

The reaction mixtures were analyzed by LC-MS with the following settings: 5 μL injection volume on a Kinetex 2.6 μm C18 100Å 150 x 3 mm column, where solvent A is 0.1% formic acid in water and solvent B is 100 % acetonitrile with 0.1% formic acid, a 0.5 mL/min flow rate and 30 °C column chamber. The LC gradient was as follows: 0 min: 10 % B, 0.7 min: 10 % B, 5.7 min: 60% B, 5.8 min: 95% B, 6.7 min: 95% B, 6.8 min: 10% B, 10.6 min: 10% B. MS was run in positive ion mode with Full MS: resolution 35,000, mass range *m/z* 300-1200, AGC target 1e6, and maximum IT 50 ms. The dd-MS2 settings were as follows: resolution 17,500, AGC target 1e5, maximum IT 50 ms, loop count 5, isolation window *m/z* 1, and (N)CE/stepped NCE 20, 25, 30 eV. MS data of cyFPIY (*m/z* 537.2708, z=1) and cyFPIY-dc (*m/z* 491.2653, z=1) were analyzed manually with Qualbrowser. We observe less than 1% of hydroxylated cyFPIY (*m/z* 553.2662, z=1) in our replicates.

The positive controls for dioxygen-containing assays were also tested via LC-MS/MS for succinate production, along with a standard of 100 μM succinic acid. The assays and standard were analyzed by LC-MS with the following settings: 5 μL injection volume on a Kinetex 2.6 μm C18 100Å 150×3 mm column, where solvent A is 20 mM ammonium formate pH 3.2 in water and solvent B is 90 % acetonitrile with 10% ammonium formate, a 0.5 mL/min flow rate and 30 °C column chamber. The LC gradient was as follows: 0 min: 0 % B, 5.0 min: 40% B, 5.1 min: 95% B, 6.0 min: 95% B, 6.1 min: 0% B, 9.9 min: 0% B. MS was run in negative ion mode with Full MS: resolution 35,000, mass range *m/z* 50-500, AGC target 1e6, and maximum IT 50 ms. The dd-MS2 settings were as follows: resolution 17,500, AGC target 1e5, maximum IT 50 ms, loop count 5, isolation window *m/z* 1, and (N)CE/stepped NCE 20, 25, 30 eV.

### ZjuDC Steady State Kinetics

End point assays were 50 µL total volume containing final concentrations of 50 mM HEPES pH 7.5, 0.5 μM ZjuDC, 20 μM FeSO_4_-7H_2_O, 500 μM 2-oxoglutarate, 400 μM L-ascorbic acid sodium salt, and 4 % DMSO to maintain solubility of cyFPIY. The initial velocity of ZjuDC-catalyzed styrylamine formation was monitored by discontinuous LC-MS/MS-based assays. Nine concentrations of cyFPIY, 0.25, 0.50, 1.0, 2.0, 3.5, 5, 7.5, 10, and 15 μM, were incubated at room temperature and quenched with 100% methanol after 30 sec, 2 min, 4min, 6 min, 8 min, and 10 min to generate an initial rate. Each substrate concentration and time point was conducted in quadruplets. All assays were analyzed by LC-MS with the following settings: 5 μL injection volume on a Kinetex 2.6 μm C18 100Å 150×3 mm column, where solvent A is 0.1% formic acid in water and solvent B is 100 % acetonitrile with 0.1% formic acid, a 0.5 mL/min flow rate and 30 °C column chamber. The LC gradient was as follows: 0 min: 10 % B, 5.7 min: 60% B, 5.8 min: 95% B, 6.7 min: 95% B, 6.8 min: 10% B, 10.6 min: 10% B. MS was run in positive ion mode with Full MS: resolution 35,000, mass range *m/z* 300-1,200, AGC target 1e6, and maximum IT 50 ms. The dd-MS2 settings were as follows: resolution 17,500, AGC target 1e5, maximum IT 50 ms, loop count 5, isolation window *m/z* 1, and (N)CE/stepped NCE 20, 25, 30 eV. MS data of cyFPIY-dc (*m/z* 491.2653, z=1) was analyzed manually with Qualbrowser.

The substrate cyFPIY and product cyFPIY-dc were quantified using corresponding MS1 extracted ion chromatograms (EIC) peak areas normalized to total detected EIC peak areas. EIC peak areas were determined by peak integration in QualBrowser and were further analyzed and plotted in GraphPad Prism. Product formation was converted to μM by calculating percent product formation through area under the curve and then using that percentage to calculate product concentration from the known substrate concentration added to the assays. The k_cat_ and K_M_ values were determined by fitting the initial velocity data using the equation V_0_ = V_max_[S]/(K_M_ + [S]) where V_0_ is the initial velocity, V_max_ is the maximal velocity, [S] is the concentration of substrate, and K_M_ is the Michaelis constant. Substrate inhibition occured when ascorbate and 2-oxoglutarate concentrations were 4 mM and 5 mM, respectively.

### ZjuDC enzyme assays

*In vitro* enzyme assays were 50 μL total volume containing 50 mM HEPES pH 7.5, 5 μM ZjuDC, 100 μM cyFPIY, 200 μM FeSO_4_-7H_2_O, 500 μM 2-oxoglutarate, 400 μM L-ascorbic acid sodium salt, and 4% dimethyl sulfoxide (DMSO) maintain solubility of cyFPIY. cyFPIY was dissolved in 100% DMSO and added to enzyme assays for a total of 4% DMSO in the assay. All assays conducted for the cofactor/substrate screen contained 50 mM HEPES pH 7.5, 5 μM ZjuDC, 90 μM cyFPIY, and 400 μM L-ascorbic acid sodium salt where the 200 μM FeSO_4_-7H_2_O and 500 μM 2-oxoglutarate where added back in respectively. It was later determined that ascorbate was not required for ZjuDC activity. Negative control assays were performed with ZjuDC that had been boiled at 95 °C for 20 min while maintaining the same conditions described. All assays were done in triplicates and incubated at 30 °C for 24 h before quenching with an equal volume of 100 % methanol. Quenched assays were then centrifuged at 16,200 x *g* for 5 min to pellet precipitate. The resulting supernatant was collected for analysis.

The assays were analyzed by LC-MS with the following settings: 5 μL injection volume on a Kinetex μm C18 100Å 150×3 mm column, where solvent A is 0.1% formic acid in water and solvent B is 100 % acetonitrile with 0.1% formic acid, a 0.5 mL/min flow rate and 30 °C column chamber. The LC gradient was as follows: 0 min: 10 % B, 0.7 min: 10 % B, 5.7 min: 60% B, 5.8 min: 95% B, 6.7 min: 95% B, 6.8 min: 10% B, 10.6 min: 10% B. MS was run in positive ion mode with Full MS: resolution 35,000, mass range *m/z* 300-1,200, AGC target 1e6, and maximum IT 50 ms. The dd-MS2 settings were as follows: resolution 17,500, AGC target 1e5, maximum IT 50 ms, loop count 5, isolation window *m/z* 1, and (N)CE/stepped NCE 20, 25, 30 eV. MS data of cyFPIY (*m/z* 537.2708, z=1) and cyFPIY-dc (*m/z* 491.2653, z=1) were analyzed manually with Qualbrowser.

### Scaled *in vitro* formation of cyFPIY-dc

The assay was 10 mL in total volume containing final concentrations of 50 mM HEPES pH 7.5, 19.5 μM ZjuDC, 390 μM cyFPIY, 780 μM FeSO_4_-7H_2_O, 1.95 mM 2-oxoglutarate, 1.56 mM L-ascorbic acid sodium salt, and 4 % DMSO to maintain solubility of cyFPIY. cyFPIY (9832 µM, 3.2 mg total) was dissolved in 100% DMSO prior to use. The reaction was incubated at 30°C for 24 h and quenched with equal volume of 100% methanol. Contents were centrifuged at 4000 x *g* for 10 min to pellet precipitate. The resulting supernatant was dried *in vacuo* and extracted four times with 25 mL of ethyl acetate. The organic layer was kept, dried *in vacuo*, and resuspended in 30% acetonitrile for semi preparative HPLC purification with the following LC settings: Kinetex® 5 μm C18 100 Å 250 × 10 mm column, solvent A - 0.1% TFA, solvent B - acetonitrile (0.1% TFA), 1.5 mL/min, 0 min - 35% B, 1 min - 35% B, 21 min - 40% B, 22 min - 95% B, 25 min - 95% B, 25.1 min - 35% B, 45 min - 35% B. cyFPIY-dc (**2**) was obtained as a white powder (2.9 mg) and stored at - 80 °C for later use with enzyme assays.

### ZjuNMT expression and purification

The expression and purification of His_8_-ZjuNMT (**Data S2**) followed the same protocol described for ZjuDC with a slight deviation in the buffer used in dialysis prior to assays. After IMAC elution, ZjuNMT was added to 3.5 kDa MWCO dialysis tubing for dialysis in 50 mM Tris-HCl pH 8.0, 100 mM NaCl, 3 mM GSH, and 0.3 mM GSSG with stirring at 4 °C overnight. After dialysis, the protein was concentrated in Amicon Ultra-15 mL 10 kDa MWCO centrifugal concentrators and stored at -80°C for later use with enzyme assays.

### ZjuNMT enzyme Assays

Both cyFPIY and cyFPIY-dc were used in enzyme assays to test for promiscuity of ZjuNMT. *In vitro* enzyme assays were 50 μL total volume containing 50 mM Tris-HCl pH 8.0, 10 μM ZjuNMT, 100 μM cyFPIY or 100 μM cyFPIY-dc, and 1 mM *S*-adenosylmethionine (SAM). Stocks of cyFPIY-dc and cyFPIY were dissolved in 100% DMSO and added to enzyme assays for a total of 4% DMSO in the assay. The cofactor screen contained 50 mM Tris-HCl pH 8, 10 μM ZjuNMT, and 100 μM cyFPIY without SAM. Negative control assays were performed with ZjuNMT that boiled at 95 °C for 20 min while maintaining the same conditions with cyFPIY-dc as described. All assays were done in triplicates and incubated at 30 °C for 24 h before quenching with an equal volume of 100 % methanol. Time course assays were conducted under the same conditions described and incubated at 30 °C for 10 min, 30 min, 1 h, 2 h, and 4 h being quenched with an equal volume of 100% methanol. Quenched assays were then centrifuged using a at 16,200 x *g* for 5 min to pellet precipitate. The resulting supernatant was collected for analysis.

The assays were analyzed by LC-MS with the following settings: 5 μL injection volume on a Kinetex 2.6 μm C18 100Å 150 x 3 mm column, where solvent A is 0.1% formic acid in water and solvent B is 100 % acetonitrile with 0.1% formic acid, a 0.5 mL/min flow rate and 30 °C column chamber. The LC gradient was as follows: 0 min: 10 % B, 0.7 min: 10 % B, 5.7 min: 60% B, 5.8 min: 95% B, 6.7 min: 95% B, 6.8 min: 10% B, 10.6 min: 10% B. MS was run in positive ion mode with Full MS: resolution 35,000, mass range *m/z* 300-1,200, AGC target 1e6, and maximum IT 50 ms. The dd-MS2 settings were as follows: resolution 17,500, AGC target 1e5, maximum IT 50 ms, loop count 5, isolation window *m/z* 1, and (N)CE/stepped NCE 20, 25, 30 eV. MS data of cyFPIY-dc (*m/z* 491.2653, z=1), Lotusine A (*m/z* 519.2966, z=1), and *N,N*-Me_2_-cyFPIY (*m/z* 565.3021, z=1) were analyzed manually with Qualbrowser.

### Scaled in vitro formation of Lotusine A

*In vitro* lotusine A reaction was 13 mL total volume, aliquoted in 250 μL reactions. Each 250 uL volume contained a final concentration of 50 mM Tris-HCl pH 8.0, 10 μM ZjuNMT, 100 μM cyFPIY-dc (total 2.6 mg), 4 % DMSO to maintain solubility of cyFPIY-dc, and 1 mM SAM. The reactions were incubated at 30°C for 30 min and quenched with equal volume of 100% methanol. After 30 min of incubation, an increase in trimethylation occurred. Reaction contents were centrifuged at 16,200 x *g* for 5 min to pellet precipitate. The supernatants were pooled, dried *in vacuo*, and resuspended in 30% acetonitrile for semipreparative HPLC purification with the following LC settings: Kinetex® 5 μm C18 100 Å 250 x 10 mm column, solvent A - 0.1% TFA, solvent B - acetonitrile (0.1% TFA), 1.5 mL/min, 0 min - 35% B, 1 min - 35% B, 21 min - 40% B, 22 min - 95% B, 25 min - 95% B, 25.1 min - 35% B, 45 min - 35% B. Lotusine A was obtained as a white amorphous powder (2.2 mg).

### One pot formation of sanjoinine A and adouetine X

One pot enzyme assays were 50 μL in total volume containing final concentrations of 50 mM HEPES pH 7.5, 10 μM ZjuDC, 10 μM ZjuNMT, 25 μM substrate, 200 μM FeSO_4_-7H_2_O, 500 μM 2-oxoglutarate, 400 μM L-ascorbic acid sodium salt, 1 mM SAM, and 4% dimethyl sulfoxide (DMSO) to maintain solubility of substrates. The substrates were cyFLLY and cyLLIY, each was dissolved in 100% DMSO and added to enzyme assays for a total of 4% DMSO. Assays were incubated for 1, 2, 4, 8, 16, and 24 h at 30 °C and quenched with equal volume of 100% methanol. Quenched assays were centrifuged at 16,200 x *g* for 5 min to pellet precipitate. The resulting supernatant was collected for product formation analysis.

The assays were analyzed by LC-MS with the following settings: 5 μL injection volume on a Kinetex 2.6 μm C18 100Å 150 x 3 mm column, where solvent A is 0.1% formic acid in water and solvent B is 100 % acetonitrile with 0.1% formic acid, a 0.5 mL/min flow rate and 30 °C column chamber. The LC gradient was as follows: 0 min: 10 % B, 0.7 min: 10 % B, 5.7 min: 60% B, 5.8 min: 95% B, 6.7 min: 95% B, 6.8 min: 10% B, 10.6 min: 10% B. MS was run in positive ion mode with Full MS: resolution 35,000, mass range *m/z* 300-1,200, AGC target 1e6, and maximum IT 50 ms. The dd-MS2 settings were as follows: resolution 17,500, AGC target 1e5, maximum IT 50 ms, loop count 5, isolation window *m/z* 1, and (N)CE/stepped NCE 20, 25, 30 eV. MS data of sanjoinine A (*m/z* 535.3279, z=1) and adouetine X (*m/z* 501.3435, z=1) were analyzed manually with Qualbrowser, and compared to authentic standards.

### Molecular Networking analysis for Cyclopeptide Alkaloids

For cyclopeptide alkaloid chemotyping, metabolomic datasets from Matthaei botanical garden (MassIVE-GNPS accession MSV000087872) and from *Ziziphus jujuba* stem and leaf samples of this study were filtered for analytes with iminium ion masses of unmethylated, monomethylated and dimethylated proteinogenic amino acids^11,80^ with MassQL^84^ (v31.4) with the following command: QUERY scaninfo(MS2DATA) WHERE MS2PROD=(58.06513 OR 60.04439 OR 70.06513 OR 72.08078 OR 74.06004 OR 84.04439 OR 84.08078 OR 86.09643 OR 87.05529 OR 88.0393 OR 88.07569 OR 100.11208 OR 101.07094 OR 101.10732 OR 102.05495 OR 102.09134 OR 104.05285 OR 110.07127 OR 114.12773 OR 115.08659 OR 115.12297 OR 116.0706 OR 118.0685 OR 120.08078 OR 124.08692 OR 129.10224 OR 129.11347 OR 129.13862 OR 130.08625 OR 132.08415 OR 134.09643 OR 136.07569 OR 138.10257 OR 143.12912 OR 148.11208 OR 150.09134 OR 157.14477 OR 159.09167 OR 164.10699 OR 173.10732 OR 187.12297):TOLERANCEPPM=10:INTENSITYPERCENT= 5. From the MassQL-filtered mass spectra and tandem mass spectra of CPA standards (sanjoinine A, adouetine X, selanine A, jubanine K), a molecular network was created using the online workflow (https://ccms-ucsd.github.io/GNPSDocumentation/) on the GNPS website (http://gnps.ucsd.edu)^72^. The data was filtered by removing all MS/MS fragment ions within +/-17 Da of the precursor *m/z*. MS/MS spectra were window filtered by choosing only the top 6 fragment ions in the +/-50 Da window throughout the spectrum. The precursor ion mass tolerance was set to 0.05 Da and a MS/MS fragment ion tolerance of 0.05 Da. A network was then created where edges were filtered to have a cosine score above 0.7 and more than 6 matched peaks. Further, edges between two nodes were kept in the network if and only if each of the nodes appeared in each other’s respective top 10 most similar nodes. Finally, the maximum size of a molecular family was set to 500, and the lowest scoring edges were removed from molecular families until the molecular family size was below this threshold. The spectra in the network were then searched against GNPS’ spectral libraries. The library spectra were filtered in the same manner as the input data. All matches kept between network spectra and library spectra were required to have a score above 0.7 and at least 6 matched peaks. The molecular network was analyzed for the main CPA cluster and visualized by Cytoscape (v3.10.3)^85^.

## Acknowledgements

This study was supported by NIGMS (grant R35GM146934 to R.D.K., grant F32GM146395 to L.S.M., grant F31GM155959 to K.S., Pharmacological Sciences Training Program predoctoral fellowship T32GM140223 to D.O.), the Hermann Frasch Foundation (R.D.K.), a Rackham Merit Fellowship (D.O.), the PhRMA foundation (predoctoral fellowship, D.N.C.), and the USDA Specialty Crop Block Grant Program through the New Mexico Department of Agriculture (S.Y.). We thank Dr. George Lomonosoff (John Innes Centre, UK) for sharing the pEAQ-HT vector. This research was supported in part through computational resources and services provided by Advanced Research Computing at the University of Michigan, Ann Arbor. We thank Dr. Stephen Ragsdale for anaerobic chamber access and Kareem Aboulhosn for assistance, Dr. Emily Scott for Fast Protein Liquid Chromatography use, and Dr. Janet Smith and Dr. Graham Moran for helpful discussions.

## Author Contributions

R.D.K. conceived the idea. J.H., R.D.K. performed bioinformatic analysis for biosynthetic gene identification and *in planta* characterization of biosynthetic genes. J.H., X.W. and R.D.K. purified and characterized peptide natural products from source and transgenic plants. J.H. and L.S.M. performed enzyme assays. J.H, D.O., X.W., K.S., D.N.C., G.M., W.L., L.S.M, K.M.M. performed scaled tobacco infiltration and peptide purification. L.M.P. performed SkrBURP metabolic engineering. S.Y. cultivated source plants. J.H., L.S.M., X.W. and R.D.K. wrote the manuscript. All authors have read and approved the manuscript.

