## Supporting Information for "Biosynthesis of 14-membered cyclopeptide alkaloids via non-heme-iron- and 2-oxoglutarate-dependent oxidative decarboxylation"

### Table of contents


| Table/Figure | Description | Page |
|---|---|---|
| Table S1 | $^{1}$H NMR (800 MHz) and $^{13}$C NMR (200 MHz) data of cyFPIY (1) in DMSO-$d_6$ | 4 |
| Table S2 | $^{1}$H-NMR (800 MHz) and $^{13}$C-NMR (200 MHz) data of jubanine K in DMSO-$d_6$ | 5 |
| Table S3 | Differential gene expression analysis towards reconstitution of 14-membered CPA biosynthesis pathway in *Ziziphus jujuba* | 6 |
| Table S4 | $^{1}$H-NMR (800 MHz) and $^{13}$C-NMR (200 MHz) data of cyFPIY-dc (2) in DMSO-$d_6$ | 7 |
| Table S5 | $^{1}$H-NMR (800 MHz) and $^{13}$C-NMR (200 MHz) data of lotusine A (3) in DMSO-$d_6$ | 8 |
| Table S6 | $^{1}$H NMR (800 MHz) and $^{13}$C NMR (200 MHz) data of cyFLLY (5) in DMSO-$d_6$ | 9 |
| Table S7 | $^{1}$H NMR (800 MHz) and $^{13}$C NMR (200 MHz) data of cyLLIY (6) in DMSO-$d_6$ | 10 |
| Table S8 | $^{1}$H-NMR (800 MHz) and $^{13}$C-NMR (200 MHz) data of sanjoinine A in DMSO-$d_6$ | 11 |
| Figure S1 | Chemical structures of representative RiPPs with α-N-methylations and/or decarboxylated C-termini | 12 |
| Figure S2 | Transient gene expression analysis of fused and split ZjuBURP constructs in *Nicotiana benthamiana* | 13 |
| Figure S3 | $^{1}$H NMR spectrum of cyFPIY in DMSO-$d_6$ (800 MHz) | 14 |
| Figure S4 | $^{1}$H-$^{1}$H COSY NMR spectrum of cyFPIY in DMSO-$d_6$ | 14 |
| Figure S5 | $^{1}$H-$^{1}$H TOCSY NMR spectrum of cyFPIY in DMSO-$d_6$ | 15 |
| Figure S6 | ROESY NMR spectrum of cyFPIY in DMSO-$d_6$ | 15 |
| Figure S7 | HSQC NMR spectrum of cyFPIY in DMSO-$d_6$ | 16 |
| Figure S8 | HMBC NMR spectrum of cyFPIY in DMSO-$d_6$ | 16 |
| Figure S9 | $^{13}$C NMR spectrum of cyFPIY in DMSO-$d_6$ (200 MHz) | 17 |
| Figure S10 | Key 2D NMR correlations of cyFPIY in DMSO-$d_6$ | 17 |
| Figure S11 | Marfey's analysis of cyFPIY | 18 |
| Figure S12 | Cyclopeptide alkaloid-containing spectral cluster of peptide-filtered LC-MS/MS datasets from specific plant sources | 19 |
| Figure S13 | $^{1}$H NMR spectrum of jubanine K in DMSO-$d_6$ (800 MHz). | 20 |
| Figure S14 | $^{1}$H-$^{1}$H COSY NMR spectrum of jubanine K in DMSO-$d_6$ | 20 |
| Figure S15 | $^{1}$H-$^{1}$H TOCSY NMR spectrum of jubanine K in DMSO-$d_6$ | 21 |
| Figure S16 | NOESY NMR spectrum of jubanine K in DMSO-$d_6$ | 21 |
| Figure S17 | HSQC NMR spectrum of jubanine K in DMSO-$d_6$ | 22 |
| Figure S18 | HMBC NMR spectrum of jubanine K in DMSO-$d_6$ | 22 |
| Figure S19 | $^{13}$C NMR spectrum of jubanine K in DMSO-$d_6$ (200MHz) | 23 |
| Figure S20 | Key 2D NMR correlations of jubanine K in DMSO-$d_6$ | 23 |
| Figure S21 | Stereochemical analysis of jubanine K | 24 |
| Figure S22 | MS characterization of jubanine K and related metabolites in *Ziziphus jujuba* samples. | 25-27 |
| Figure S23 | Characterization of candidate precursor peptide of jubanine K from *Ziziphus jujuba* var. Dongzao. | 28 |
| Figure S24 | Protein model prediction of ZjuDC and alignments with experimentally determined plant non-heme Fe(II)- and 2-oxoglutarate dependent enzymes. | 29 |
| Figure S25 | SDS-PAGE of heterologously expressed and purified proteins | 30 |
| Figure S26 | Succinate detection in ZjuDC in vitro enzyme assay compared to a succinate authentic standard | 31 |
| Figure S27 | $^{1}$H NMR spectrum of cyFPIY-dc in DMSO-$d_6$ (800 MHz) | 32 |
| Figure S28 | $^{1}$H-$^{1}$H COSY NMR spectrum of cyFPIY-dc in DMSO-$d_6$ | 32 |
| Figure S29 | $^{1}$H-$^{1}$H TOCSY NMR spectrum of cyFPIY-dc in DMSO-$d_6$ | 33 |
| Figure S30 | NOESY NMR spectrum of cyFPIY-dc in DMSO-$d_6$ | 33 |
| Figure S31 | HSQC NMR spectrum of cyFPIY-dc in DMSO-$d_6$ | 34 |
| Figure S32 | HMBC NMR spectrum of cyFPIY-dc in DMSO-$d_6$ | 34 |
| Figure S33 | $^{13}$C NMR spectrum of cyFPIY-dc in DMSO-$d_6$ (200MHz) | 35 |
| Figure S34 | Key 2D NMR correlations of cyFPIY-dc in DMSO-$d_6$ | 35 |
| Figure S35 | Stereochemical analysis of cyFPIY-dc | 36 |
| Figure S36 | ZjuNMT product methylation *in vitro* and *in planta* | 37-39 |
| Figure S37 | $^{1}$H NMR spectrum of Lotusine A in DMSO-$d_6$ (800 MHz) | 40 |
| Figure S38 | $^{1}$H-$^{1}$H COSY NMR spectrum of Lotusine A in DMSO-$d_6$ | 40 |
| Figure S39 | $^{1}$H-$^{1}$H TOCSY NMR spectrum of Lotusine A in DMSO-$d_6$ | 41 |
| Figure S40 | NOESY NMR spectrum of Lotusine A in DMSO-$d_6$ | 41 |
| Figure S41 | HSQC NMR spectrum of Lotusine A in DMSO-$d_6$ | 42 |

| Table/Figure | Description | Page |
|---|---|---|
| Figure S42 | HMBC NMR spectrum of Lotusine A in DMSO-$d_6$ | 42 |
| Figure S43 | $^{13}$C NMR spectrum of Lotusine A in DMSO-$d_6$ (200MHz) | 43 |
| Figure S44 | Key 2D NMR correlations of Lotusine A in DMSO-$d_6$ | 43 |
| Figure S45 | Stereochemical analysis of lotusine A | 44 |
| Figure S46 | Burpitide cyclase SkrBURP constructs for optimization of cyFLLY production | 45 |
| Figure S47 | SkrBURP optimization for cyclic peptide yields from transgenic *Nicotiana benthamiana* | 46 |
| Figure S48 | $^{1}$H NMR spectrum of cyLLIY in DMSO-$d_6$ (800 MHz) | 47 |
| Figure S49 | $^{1}$H-$^{1}$H COSY-NMR spectrum of cyLLIY in DMSO-$d_6$ | 47 |
| Figure S50 | $^{1}$H-$^{1}$H TOCSY NMR spectrum of cyLLIY in DMSO-$d_6$ | 48 |
| Figure S51 | ROESY NMR spectrum of cyLLIY in DMSO-$d_6$ | 48 |
| Figure S52 | HSQC NMR spectrum of cyLLIY in DMSO-$d_6$ | 49 |
| Figure S53 | HMBC NMR spectrum of cyLLIY in DMSO-$d_6$ | 49 |
| Figure S54 | $^{13}$C NMR spectrum of cyLLIY in DMSO-$d_6$ (200 MHz) | 50 |
| Figure S55 | Key 2D NMR correlations of cyLLIY in DMSO-$d_6$ | 50 |
| Figure S56 | Stereochemical analysis of cyLLIY | 51 |
| Figure S57 | $^{1}$H NMR spectrum of cyFLLY in DMSO-$d_6$ (800MHz) | 52 |
| Figure S58 | $^{1}$H-$^{1}$H COSY NMR spectrum of cyFLLY in DMSO-$d_6$ | 52 |
| Figure S59 | $^{1}$H-$^{1}$H TOCSY NMR spectrum of cyFLLY in DMSO-$d_6$ | 53 |
| Figure S60 | ROESY NMR spectrum of cyFLLY in DMSO-$d_6$ | 53 |
| Figure S61 | HSQC NMR spectrum of cyFLLY in DMSO-$d_6$ | 54 |
| Figure S62 | HMBC NMR spectrum of cyFLLY in DMSO-$d_6$ | 54 |
| Figure S63 | $^{13}$C NMR spectrum of cyFLLY in DMSO-$d_6$ (200MHz) | 55 |
| Figure S64 | Key 2D NMR correlations of cyFLLY in DMSO-$d_6$ | 55 |
| Figure S65 | Stereochemical analysis of cyFLLY. | 56 |
| Figure S66 | $^{1}$H NMR spectrum of sanjoinine A authentic standard in DMSO-$d_6$ (800MHz) | 57 |
| Figure S67 | $^{1}$H-$^{1}$H COSY NMR spectrum of sanjoinine A authentic standard in DMSO-$d_6$ | 57 |
| Figure S68 | $^{1}$H-$^{1}$H TOCSY NMR spectrum of sanjoinine A authentic standard in DMSO-$d_6$ | 58 |
| Figure S69 | ROESY NMR spectrum of sanjoinine A authentic standard in DMSO-$d_6$ | 58 |
| Figure S70 | HSQC NMR spectrum of sanjoinine A authentic standard in DMSO-$d_6$ | 59 |
| Figure S71 | HMBC NMR spectrum of sanjoinine A authentic standard in DMSO-$d_6$ | 59 |
| Figure S72 | $^{13}$C NMR spectrum of sanjoinine A authentic standard in DMSO-$d_6$ (200 MHz) | 60 |
| Figure S73 | Key 2D NMR correlations of sanjoinine A authentic standard in DMSO-$d_6$ | 60 |
| Figure S74 | $^{1}$H NMR spectrum of adouetineX in $CD_3OD$ (800 MHz) | 61 |
| Figure S75 | $^{1}$H-$^{1}$H COSY NMR spectrum of adouetineX in $CD_3OD$ | 61 |
| Figure S76 | $^{1}$H-$^{1}$H TOCSY-NMR spectrum of adouetine X in $CD_3OD$ | 62 |
| Figure S77 | ROESY NMR spectrum of adouetine X in $CD_3OD$ | 62 |
| Figure S78 | HSQC NMR spectrum of adouetineX in $CD_3OD$ | 63 |
| Figure S79 | HMBC NMR spectrum of adouetineX in $CD_3OD$ | 63 |
| Figure S80 | $^{13}$C NMR spectrum of adouetineX in $CD_3OD$ (200MHz) | 64 |
| Figure S81 | Selective 1D-TOCSY NMR spectrum of adouetineX in $CD_3OD$ | 64 |
| Figure S82 | Key 2D NMR correlations of adouetineX in $CD_3OD$ | 65 |
| Figure S83 | Stereochemical analysis of adouetine X | 66 |
| Figure S84 | cyFLxY diversification via transient expression of SkrBURP-FLxY in *N. benthamiana* | 67-75 |
| Figure S85 | cyLLxY diversification via transient expression of SkrBURP-LLxY in *N. benthamiana* | 76-84 |
| Figure S86 | cyFLxY CPA diversification via transient expression of SkrBURP-FLxY, ZjuDC and ZjuNMT in *N. benthamiana* | 85-88 |
| Figure S87 | cyLLxY CPA diversification via transient expression of SkrBURP-LLxY, ZjuDC and ZjuNMT in *N. benthamiana* | 89-90 |
| Figure S88 | Biosynthetic pathway of lotusine A in *Ziziphus jujuba* | 91 |
| Figure S89 | Proposed mechanisms for ZjuDC based on intermediates | 92 |
| Figure S90 | References | 93 |

### Table S1 | $^{1}$H NMR (800 MHz) and $^{13}$C NMR (200 MHz) data of cyFPIY (1) in DMSO-$d_6$.

| Residue | C | δ($^{13}$C) [ppm] | H | δ($^{1}$H) (int, m, *J* [Hz]) [ppm] | COSY | ROESY | HMBC |
|---|---|---|---|---|---|---|---|
| Phe$^{1}$ | α | 51.2 | α | 4.38 (1H, t, 5.3) | Phe$^{1}$: Hβ1/2, NH | Phe$^{1}$: Hβ1/2, H2/6, NH; Pro$^{2}$: Hδ2 | n/a |
| | β | 36.2 | β1 | 2.96 (1H, dd, 5.3, 14.0) | Phe$^{1}$: Hα, Hβ2 | Phe$^{1}$: Hα, Hβ2, H2/6 | Phe$^{1}$: Cα, C1, C2/6, C=O |
| | C1 | 134.0 | β2 | 2.90 (1H, dd, 7.6, 14.0) | Phe$^{1}$: Hα, Hβ1 | Phe$^{1}$: Hα, Hβ1, H2/6 | Phe$^{1}$: Cα, C1, C2/6, C=O |
| | C2 | 130.0 | H2 | 7.23 (1H, dd, 2.9, 6.6) | Phe$^{1}$: H3/5, H4 | Phe$^{1}$: Hα, Hβ1/2, H3/5, H4 | Phe$^{1}$: Cβ, C1, C3/5, C4, C6 |
| | C3 | 128.4 | H3 | 7.27 (1H, m) | Phe$^{1}$: H2/6, H4 | Phe$^{1}$: H2/6, H4 | Phe$^{1}$: C1, C2/6, C4 |
| | C4 | 127.2 | H4 | 7.26 (1H, m) | Phe$^{1}$: H2/6, H3/5 | Phe$^{1}$: H2/6, H3/5 | Phe$^{1}$: C2/6, C3/5 |
| | C5 | 128.4 | H5 | 7.27 (1H, m) | Phe$^{1}$: H2/6, H4 | Phe$^{1}$: H2/6, H4 | Phe$^{1}$: C1, C2/6, C4 |
| | C6 | 130.0 | H6 | 7.23 (1H, dd, 2.9, 6.6) | Phe$^{1}$: H3/5, H4 | Phe$^{1}$: Hα, Hβ1/2, H3/5, H4 | Phe$^{1}$: Cβ, C1, C2, C3/5, C4 |
| | C=O | 166.5 | NH | 8.14 (1H, s) | Phe$^{1}$: Hα | Phe$^{1}$: Hα | n/a |
| Pro$^{2}$ | α | 65.1 | α | 4.03 (1H, d, 7.2) | Pro$^{2}$: Hβ | Pro$^{2}$: Hβ, Hγ1/2, Hδ1; Ile$^{3}$: NH, Tyr$^{4}$: H5 | Phe$^{1}$: C=O, Pro$^{2}$: Cβ, C=O |
| | β | 80.6 | | | | | |
| | γ | 30.7 | β | 5.14 (1H, dt, 10.6, 7.2) | Pro$^{2}$: Hα, Hγ1/2 | Pro$^{2}$: Hα, Hγ1/2, Hδ1; Tyr$^{4}$: H3 | Pro$^{2}$: C=O; Tyr$^{4}$: C4 |
| | δ | 44.7 | γ1 | 2.34 (1H, dt, 11.3, 5.7) | Pro$^{2}$: Hβ, Hγ2, Hδ1 | Pro$^{2}$: Hβ, Hγ2, Hδ1/2 | n/a |
| | | | γ2 | 1.94 (1H, m) | Pro$^{2}$: Hβ, Hγ1, Hδ1/2 | Pro$^{2}$: Hα, Hβ, Hγ1, Hδ1/2 | Pro$^{2}$: Cβ, Cδ |
| | | | δ1 | 2.87 (1H, m) | Pro$^{2}$: Hγ1/2, Hδ2 | Pro$^{2}$: Hα, Hβ, Hγ1/2, Hδ2 | n/a |
| | C=O | 168.5 | δ2 | 3.87 (1H, t, 9.4) | Pro$^{2}$: Hγ2, Hδ1 | Phe$^{1}$: Hα; Pro$^{2}$: Hγ1/2, Hδ1 | Phe$^{1}$: C=O; Pro$^{2}$: Cα, Cβ, Cγ |
| Ile$^{3}$ | α | 57.3 | α | 3.72 (1H, t, 8.3) | Ile$^{3}$: Hβ, NH | Ile$^{3}$: Hβ, Hγ1A/B, Hγ2, NH; Tyr$^{4}$: NH | Pro$^{2}$: C=O, Ile$^{3}$: Cβ, Cγ1, Cδ, C=O |
| | β | 37.7 | β | 1.41 (1H, m) | Ile$^{3}$: Hα, Hγ1A/B | Ile$^{3}$: Hα, Hγ1A/B, Hγ2, Hδ, NH | Ile$^{3}$: Cα |
| | γ1 | 24.3 | γ1A ($CH_2$) | 1.46 (1H, m) | Ile$^{3}$: Hβ, Hγ1B, Hδ | Ile$^{3}$: Hα, Hβ, Hγ1B, Hγ2, Hδ | Ile$^{3}$: Cδ |
| | γ2 | 14.7 | γ1B ($CH_2$) | 1.00 (1H, m) | Ile$^{3}$: Hβ, Hγ1A, Hδ | Ile$^{3}$: Hα, Hβ, Hγ1A, Hγ2, Hδ | Ile$^{3}$: Cβ, Cδ |
| | δ | 11.1 | γ2 | 0.73 (3H, d, 6.8) | Ile$^{3}$: Hβ | Ile$^{3}$: Hα, Hβ, Hγ1A/B, Hδ | Ile$^{3}$: Cα, Cβ, Cγ1 |
| | | | δ | 0.76 (3H, t, 7.4) | Ile$^{3}$: Hγ1A/B, Hβ | Ile$^{3}$: Hα, Hβ, Hγ1A/B, Hγ2 | Ile$^{3}$: Cα, Cβ, Cγ1 |
| | C=O | 169.5 | NH | 6.91 (1H, d, 8.7) | Ile$^{3}$: Hα | Pro$^{2}$: Hα; Ile$^{3}$: Hα, Hβ; Tyr$^{4}$: NH | Pro$^{2}$: C=O; Ile$^{3}$: Cα |
| Tyr$^{4}$ | α | 52.5 | α | 4.82 (1H, m) | Tyr$^{4}$: Hβ1/2, NH | Tyr$^{4}$: Hβ1, H2, NH | Ile$^{3}$: C=O; Tyr$^{4}$: Cβ, C=O |
| | β | 36.7 | β1 | 3.23 (1H, dd, 5.8, 12.9) | Tyr$^{4}$: Hα, Hβ2 | Tyr$^{4}$: Hα, Hβ2, H2 | Tyr$^{4}$: C1 |
| | C1 | 131.8 | β2 | 2.53 (1H, d, 12.9) | Tyr$^{4}$: Hα, Hβ1 | Tyr$^{4}$: Hβ1, H6, NH | Tyr$^{4}$: Cα, C1, C2, C6, C=O |
| | C2 | 132.6 | H2 | 7.01 (1H, dd, 2.1, 8.4) | Tyr$^{4}$: H3 | Tyr$^{4}$: Hβ2, H5, NH | Tyr$^{4}$: Cβ, C1, C3, C4, C5, C6 |
| | C3 | 115.9 | H3 | 6.94 (1H, dd, 2.1, 8.4) | Tyr$^{4}$: H2 | Pro$^{2}$: Hβ; Tyr$^{4}$: H2 | Tyr$^{4}$: C1, C2, C4, C5 |
| | C4 | 156.1 | H5 | 6.79 (1H, dd, 2.1, 8.3) | Tyr$^{4}$: H6 | Pro$^{2}$: Hα; Tyr$^{4}$: H6 | Tyr$^{4}$: C1, C3, C4 |
| | C5 | 119.8 | H6 | 7.03 (1H, dd, 2.1, 8.3) | Tyr$^{4}$: H5 | Tyr$^{4}$: Hα, Hβ1, H3 | Tyr$^{4}$: Cβ, C1, C2, C3, C4, C5 |
| | C6 | 129.7 | NH | 7.95 (1H, d, 10.2) | Tyr$^{4}$: Hα | Ile$^{3}$: Hα, Tyr$^{4}$: Hα, Hβ2, H2 | Ile$^{3}$: C=O; Tyr$^{4}$: Cα |

**Table S2 | $^{1}$H-NMR (800 MHz) and $^{13}$C-NMR (200 MHz) data of jubanine K in DMSO-$d_6$.**

| Residue | C | δ($^{13}$C) [ppm] | H | δ($^{1}$H) (int, m, *J* [Hz]) [ppm] | COSY | NOESY | HMBC |
|---|---|---|---|---|---|---|---|
| Phe$^{1}$ | α | 53.4 | α | 4.25a (1H, m) | Phe$^{1}$: Hβ1, Hβ2 | Phe$^{1}$: Hβ1, Hβ2, Ile$^{2}$: NH | Phe$^{1}$: Cβ1, Cβ2, C1, C=O |
| | β | 36.1 | β1 | 3.03a (1H, m) | Phe$^{1}$: Hα, Hβ2 | Phe$^{1}$: Hα, Hβ2, H2, H6 | Phe$^{1}$: Cα, C1, C2, C6 |
| | C1 | 137.3 | β2 | 2.77a (1H, m) | Phe$^{1}$: Hα, Hβ1 | Phe$^{1}$: Hα, Hβ1, H2, H6 | Phe$^{1}$: Cα, C1, C2, C6 |
| | C2 | 129.0 | H2 | 7.14 (1H, d, 7.3) | Phe$^{1}$: H3 | Phe$^{1}$: Hβ1, Hβ2, H3 | Phe$^{1}$: Cβ, C1, C3, C4, C6 |
| | C3 | 128.6 | H3 | 7.17 (1H, t, 7.3) | Phe$^{1}$: H2, H4 | Phe$^{1}$: H2, H4 | Phe$^{1}$: C1, C2, C4, C5 |
| | C4 | 126.3 | H4 | 7.20 (1H, t, 7.3) | Phe$^{1}$: H3, H5 | Phe$^{1}$: H3, H5 | Phe$^{1}$: C2, C3, C5, C6 |
| | C5 | 128.6 | H5 | 7.17 (1H, t, 7.3) | Phe$^{1}$: H4, H6 | Phe$^{1}$: H4, H6 | Phe$^{1}$: C1, C3, C4, C6 |
| | C6 | 129.0 | | | | | |
| | C=O | 168.5 | H6 | 7.14 (1H, d, 7.3) | Phe$^{1}$: H5 | Phe$^{1}$: Hβ1, Hβ2, H5 | Phe$^{1}$: Cβ, C1, C2, C4, C5 |
| | Me-1 | 41.4 | Me-1 | 2.50[a] (6H, s) | n/a | n/a | Phe$^{1}$: Cα |
| | Me-2 | 41.4 | Me-2 | 2.50[a] (6H, s) | n/a | n/a | Phe$^{1}$: Cα |
| Ile$^{2}$ | α | 53.4 | α | 4.26[a] (1H, m) | Ile$^{2}$: Hβ, NH | Ile$^{2}$: Hβ, Hγ1, Hγ2, NH; Pro$^{3}$: Hδ2 | Ile$^{2}$: Cβ, Cγ1, Cγ2, C=O |
| | β | 36.1 | β | 1.49 (1H, m) | Ile$^{2}$: Hα, Hγ1, Hγ2 | Ile$^{2}$: Hα, Hγ1, Hγ2, Hδ | Ile$^{2}$: Cα, Cγ1, Cγ2, Cδ |
| | γ1 | 24.1 | γ1A ($CH_2$) | 1.34 (1H, m) | Ile$^{2}$: Hβ, Hγ1B, Hδ | Ile$^{2}$: Hα, Hβ, Hγ1B, Hγ2, Hδ | Ile$^{2}$: Cα, Cβ, Cγ2, Cδ |
| | γ2 | 14.0 | γ1B ($CH_2$) | 0.96 (1H, m) | Ile$^{2}$: Hβ, Hγ1A, Hδ | Ile$^{2}$: Hα, Hβ, Hγ1A, Hγ2, Hδ | Ile$^{2}$: Cα, Cβ, Cγ2, Cδ |
| | δ | 10.2 | γ2 | 0.18 (3H, d, 6.7) | Ile$^{2}$: Hβ | Ile$^{2}$: Hα, Hβ, Hγ1A, Hγ1, Hδ | Ile$^{2}$: Cα, Cβ, Cγ1 |
| | | | δ | 0.74 (3H, t, 7.5) | Ile$^{2}$: Hγ1A, Hγ1B | Ile$^{2}$: Hβ, Hγ1, Hγ2 | Ile$^{2}$: Cβ, Cγ1 |
| | C=O | 168.4 | NH | 8.76 (1H, br s) | n/a | n/a | n/a |
| Pro$^{3}$ | α | 63.8 | α | 4.44 (1H, s) | Pro$^{3}$: Hβ | Pro$^{3}$: Hβ, Hγ1, Hγ2, Hδ1, Hδ2; Phe$^{4}$: NH; Tyr$^{5}$: H6 | Pro$^{3}$: Hβ, Hγ1, Hγ2; Phe$^{4}$: NH |
| | β | 77.1 | | | | | |
| | γ | 32.2 | β | 5.28 (1H, br t, 7.2) | Pro$^{3}$: Hα, Hγ1, Hγ2 | Pro$^{3}$: Hα, Hγ1, Hγ2, Hδ1, Hδ2; Tyr$^{5}$: H6 | Pro$^{3}$: Hα, Hγ1, Hγ2, Hδ2; Phe$^{4}$: H6 |
| | δ | 46.0 | γ1 | 2.57 (1H, m) | Pro$^{3}$: Hβ, Hγ1, Hδ1, Hδ2 | Pro$^{3}$: Hα, Hβ, Hγ1, Hδ1, Hδ2 | Pro$^{3}$: Hα, Hβ, Hγ1, Hδ1, Hδ2 |
| | | | γ2 | 2.11 (1H, m) | Pro$^{3}$: Hγ1, Hγ2, Hδ2 | Pro$^{3}$: Hα, Hβ, Hγ1, Hγ2, Hδ2 | Pro$^{3}$: Hγ2, Hδ2 |
| | | | δ1 | 4.14 (1H, m) | Pro$^{3}$: Hγ1, Hγ2, Hδ1 | Pro$^{3}$: Hα, Hβ, Hγ1, Hγ2, Hδ1 | Pro$^{3}$: Hδ1, Ile$^{2}$: Hα |
| | C=O | 169.4 | δ2 | 3.40 (1H, m) | Pro$^{3}$: Hβ | Pro$^{3}$: Hβ, , Hγ1, Hγ2, Hδ1, Hδ2 | Pro$^{3}$: Hβ, Hγ1, Hγ2; Phe$^{4}$: NH |
| Phe$^{4}$ | α | 57.0 | α | 4.25[a] (1H, m) | Phe$^{4}$: Hβ1, Hβ2 | Phe$^{4}$: Hβ1, Hβ2, NH, Tyr$^{5}$: NH | Phe$^{4}$: Cβ1, Cβ2, C1, C=O |
| | β | 36.1 | β1 | 3.03[a] (1H, m) | Phe$^{4}$: Hα, Hβ2 | Phe$^{4}$: Hα, Hβ2, H2, H6 | Phe$^{4}$: Cα, C1, C2, C6 |
| | C1 | 137.3 | β2 | 2.77[a] (1H, m) | Phe$^{4}$: Hα, Hβ1 | Phe$^{4}$: Hα, Hβ1, H2, H6 | Phe$^{4}$: Cα, C1, C2, C6 |
| | C2 | 126.3 | γ1A ($CH_2$) | 7.15 (1H, d, 7.3) | Phe$^{4}$: H3 | Phe$^{4}$: Hβ1, Hβ2, H3 | Phe$^{4}$: Cβ, C1, C3, C4, C6 |
| | C3 | 128.1 | γ1B ($CH_2$) | 7.20 (1H, t, 7.3) | Phe$^{4}$: H2, H4 | Phe$^{4}$: H2, H4 | Phe$^{4}$: C1, C2, C4, C5 |
| | C4 | 128.2 | γ2 | 7.29 (1H, br t, 7.3) | Phe$^{4}$: H3, H5 | Phe$^{4}$: H3, H5 | Phe$^{4}$: C2, C3, C5, C6 |
| | C5 | 128.1 | δ | 7.20 (1H, t, 7.3) | Phe$^{4}$: H4, H6 | Phe$^{4}$: H4, H6 | Phe$^{4}$: C1, C3, C4, C6 |
| | C6 | 126.3 | | | | | |
| | C=O | 168.4 | NH | 7.15 (1H, d, 7.3) | Phe$^{4}$: H5 | Phe$^{4}$: Hβ1, Hβ2, H5 | Phe$^{4}$: Cβ, C1, C2, C4, C5 |
| Tyr$^{5}$ | α | 121.8 | α | 6.79 (1H, dd, 10.0, 8.9) | Tyr$^{5}$: Hβ, NH | Tyr$^{5}$: Hβ, NH | Tyr$^{5}$: Cβ, C1 |
| | β | 108.2 | β | 5.93 (1H, d, 8.9) | Tyr$^{5}$: Hα, NH | Tyr$^{5}$: Hα, NH, H6 | Tyr$^{5}$: Cα, C1, C2, C6 |
| | C1 | 124.1 | H3 | 7.11 (1H, d, 9.1) | Tyr$^{5}$: H4 | Tyr$^{5}$: H4, OMe | Tyr$^{5}$: C1, C2, C4, C5 |
| | C2 | 151.0 | H4 | 6.93 (1H, d, 9.1) | Tyr$^{5}$: H3 | Tyr$^{5}$: H3 | Tyr$^{5}$: C2, C3, C5, C6 |
| | C3 | 114.0 | H6 | 6.78 (1H, s) | n/a | Tyr$^{5}$: Hβ; Pro$^{3}$: Hβ | Tyr$^{5}$: Cβ, C1, C2, C4, C5 |
| | C4 | 116.9 | NH | 9.27 (1H, d, 10.0) | Tyr$^{5}$: Hα | Tyr$^{5}$: Hα; Phe$^{4}$: Hα | Tyr$^{5}$: Cα, Cβ; Phe$^{4}$: C=O |
| | C5 | 150.6 | | | | | |
| | C6 | 111.8 | | | | | |
| | OMe | 56.0 | OMe | 3.79 (3H, s) | n/a | Tyr$^{5}$: H3 | Tyr$^{5}$: C2 |

[a] - overlapped signals

**Table S3 | Differential gene expression analysis towards reconstitution of 14-membered CPA biosynthesis pathway in *Ziziphus jujuba*.**

| | |
|---|---|
| ZjuBURP-ZjuPrec-FIPFY correlation | |
| Gene | Pearson coefficient |
| ZjuPrec_FIPFY | Bait (1) |
| **ZjuBURP** | **0.978** |
| Oxidative gene candidates | |
| Gene | Pearson coefficient |
| ZjuBURP | Bait (1) |
| ZjuCYP | 0.983 |
| **ZjuKOx (ZjuDC)** | **0.974** |
| ZjuPAO | 0.967 |
| Methyltransferase gene candidates | |
| Gene | Pearson coefficient |
| ZjuBURP | Bait (1) |
| **ZjuOMT1 (ZjuNMT)*** | **0.992** |
| ZjuNMT2 | 0.987 |
| ZjuOMT3 | 0.978 |
| ZjuOMT2 | 0.972 |
| ZjuNMT1 | 0.951 |
| ZjuOMT4 | 0.871 |
| ZjuNMT3 | 0.850 |
| ZjuNMT4 | 0.819 |
| ZjuNMT5 | 0.707 |

* Gene annotation: desmethylxanthohumol 6'-O-methyltransferase-like [*Ziziphus jujuba*] (GenBank: XP_015881373.2), Closest characterized homolog: O-methyltransferase [*Mangifera indica*] (65%/47%, similarity/identity).

### Table S4 | $^{1}$H-NMR (800 MHz) and $^{13}$C-NMR (200 MHz) data of cyFPIY-dc (2) in DMSO-$d_6$.

| Residue | C | δ($^{13}$C) [ppm] | H | δ($^{1}$H) (int, m, *J* [Hz]) [ppm] | COSY | ROESY | HMBC |
|---|---|---|---|---|---|---|---|
| Phe$^{1}$ | α | 50.9 | α | 4.42 (1H, m) | Phe$^{1}$: Hβ1/2, NH | Phe$^{1}$: Hβ1/2, H2/6, NH; Pro$^{2}$: δ2 | n/a |
| | β | 36.3 | β1 | 2.84 (1H, dd, 7.6, 13.2) | Phe$^{1}$: Hα, Hβ2 | Phe$^{1}$: Hα, Hβ2, H2/6 | Phe$^{1}$: Cα, C1, C2/6, C=O |
| | C1 | 134.0 | β2 | 2.94 (1H, dd, 5.1, 13.1) | Phe$^{1}$: Hα, Hβ1 | Phe$^{1}$: Hα, Hβ1, H2/6 | Phe$^{1}$: C1, C2/6 |
| | C2 | 129.8 | H2 | 7.16 (2H, m) | Phe$^{1}$: H3/5, H4 | Phe$^{1}$: Hα, Hβ1/2, H3/5 | Phe$^{1}$: C3/5, C4 |
| | C3 | 128.5 | H3 | 7.26 (2H, m) | Phe$^{1}$: H2/6, H4 | Phe$^{1}$: H2/6, H4 | Phe$^{1}$: C2/6 |
| | C4 | 127.1 | H4 | 7.26 (1H, m) | Phe$^{1}$: H2/6, H3/5 | Phe$^{1}$: H2/6, H3/5 | Phe$^{1}$: C2/6 |
| | C5 | 128.5 | H5 | 7.26 (1H, m) | Phe$^{1}$: H2/6, H4 | Phe$^{1}$: H2/6, H4 | Phe$^{1}$: C2/6 |
| | C6 | 129.8 | H6 | 7.16 (2H, m) | Phe$^{1}$: H3/5, H4 | Phe$^{1}$: Hα, Hβ1/2, H3/5 | Phe$^{1}$: C3/5, C4 |
| | C=O | 166.5 | NH | 8.15 (1H, m) | Phe$^{1}$: Hα | Phe$^{1}$: Hα | n/a |
| Pro$^{2}$ | α | 64.3 | α | 4.09 (1H, d, 6.8) | Pro$^{2}$: Hβ | Pro$^{2}$: Hβ, Hγ1, Hδ1; Ile$^{3}$: NH; Tyr$^{4}$: H5 | Pro$^{2}$: Cβ |
| | β | 82.2 | | | | | |
| | γ | 30.9 | β | 5.19 (1H, dt, 6.8, 10.7) | Pro$^{2}$: Hα, Hγ1/2 | Pro$^{2}$: Hα, Hγ1/2, Hδ1; Tyr$^{4}$: H3 | n/a |
| | δ | 45.0 | γ1 | 1.92 (1H, m) | Pro$^{2}$: Hβ, Hγ2, Hδ1/2 | Pro$^{2}$: Hα, Hβ, Hγ1/2, Hδ1/2 | Pro$^{2}$: Cβ, Cδ |
| | | | γ2 | 2.38 (1H, m) | Pro$^{2}$: Hβ, Hγ1, Hδ1 | Pro$^{2}$: Hβ, Hγ1, Hδ1/2 | n/a |
| | | | δ1 | 2.67 (1H, m) | Pro$^{2}$: Hγ1/2, Hδ2 | Pro$^{2}$: Hβ, Hγ1/2, Hδ2 | n/a |
| | C=O | 169.3 | δ2 | 3.87 (1H, t, 9.6) | Pro$^{2}$: Hγ1, Hδ1 | Phe$^{1}$: Hα, Pro$^{2}$: Hγ1/2, Hδ1 | Pro$^{2}$: Cα, Cβ, Cγ |
| Ile$^{3}$ | α | 57.1 | α | 3.97 (1H, dd, 6.1, 8.4) | Ile$^{3}$: Hβ, NH | Ile$^{3}$: Hβ, Hγ1A/B, Hγ2, NH; Tyr$^{4}$: NH | Ile$^{3}$: Cβ, Cγ1, C=O |
| | β | 36.5 | β | 1.60 (1H, m) | Ile$^{3}$: Hα, Hγ1A/B, Hγ2 | Ile$^{3}$: Hα, Hγ1A, Hγ2, Hδ | n/a |
| | γ1 | 24.1 | γ1A ($CH_2$) | 1.36 (1H, m) | Ile$^{3}$: Hγ1B, Hδ | Ile$^{3}$: Hα, Hβ, Hγ1B, Hγ2, Hδ | n/a |
| | γ2 | 11.1 | γ1B ($CH_2$) | 1.09 (1h, m) | Ile$^{3}$: Hβ, Hγ1A, Hδ | Ile$^{3}$: Hα, Hβ, Hγ1A, Hγ2, Hδ | n/a |
| | δ | 15.0 | γ2 | 0.76 (3H, d, 7.3) | Ile$^{3}$: Hβ | Ile$^{3}$: Hα, Hβ, Hγ1A, Hδ | Ile$^{3}$: Cα, Cβ, Cγ1/2 |
| | | | δ | 0.75 (3H, d, 6.3) | Ile$^{3}$: Hγ1A/B | Ile$^{3}$: Hα, Hβ, Hγ1A, Hγ2 | Ile$^{3}$: Cα, Cβ, Cγ1, Cδ |
| | C=O | 168.5 | NH | 7.36 (1H, d, 8.6) | Ile$^{3}$: Hα | Pro$^{2}$: Hα | Pro$^{2}$: C=O |
| Tyr$^{4}$ | α | 126.2 | α | 6.24 (1H, dd, 4.7, 7.1) | Tyr$^{4}$: Hβ, NH | Tyr$^{4}$: Hβ, NH | Tyr$^{4}$: C1 |
| | β | 125.2 | β | 6.59 (1H, d, 7.3) | Tyr$^{4}$: Hα | Tyr$^{4}$: Hα, H2 | Tyr$^{4}$: Cα |
| | C1 | 131.6 | H2 | 7.01 (1H, m) | Tyr$^{4}$: H3 | Tyr$^{4}$: H3 | Tyr$^{4}$: Cβ, C1, C4 |
| | C2 | 129.8 | H3 | 7.14 (1H, m) | Tyr$^{4}$: H2 | Pro$^{2}$: Hβ; Tyr$^{4}$: NH, H2 | Tyr$^{4}$: C1, C5 |
| | C3 | 118.3 | H5 | 7.01 (1H, m) | Tyr$^{4}$: H6 | Pro$^{2}$: Hα; Tyr$^{4}$: NH, H6 | Tyr$^{4}$: C3, C4 |
| | C4 | 156.5 | H6 | 6.99 (1H, d, 8.1) | Tyr$^{4}$: H5 | Tyr$^{4}$: H5 | Tyr$^{4}$: C2, C4 |
| | C5 | 121.0 | | | | | |
| | C6 | 130.6 | NH | 7.77 (1H, d, 4.6) | Tyr$^{4}$: Hα | Ile$^{3}$: Hα, NH | n/a |

### Table S5 | $^1$H-NMR (800 MHz) and $^{13}$C-NMR (200 MHz) data of lotusine A (3) in DMSO-d$_6$.

| Residue | C | δ($^{13}$C) [ppm] | H | δ($^1$H) (int, m, *J* [Hz]) [ppm] | COSY | ROESY | HMBC |
|---|---|---|---|---|---|---|---|
| Phe$^1$ | α | 64.9 | α | 4.48 (1H, m) | Phe$^1$: Hβ1/2 | Phe$^1$: Hβ1/2, N(CH$_3$)$_2$A/B | n/d |
| | β | 33.9 | β1 | 3.35 (1H, d, 12) | Phe$^1$: Hβ2 | Phe$^1$: Hβ2, H2/6 | Phe$^1$: C1 |
| | C1 | 133.2 | β2 | 2.78 (1H, m, 12) | Phe$^1$: Hα, Hβ1 | Phe$^1$: Hβ1, H2/6 | Phe$^1$: Cα, C1, C2/6, C=O |
| | C2 | 129.4 | H2 | 7.13 (1H, m) | Phe$^1$: H3/5, H4 | Phe$^1$: Hβ1/2, H3/5, H4 | Phe$^1$: C3/5, C4, C6 |
| | C3 | 128.4 | H3 | 7.27 (1H, m) | Phe$^1$: H2/6, H4 | Phe$^1$: H2/6, H4 | Phe$^1$: C1, C2/6 |
| | C4 | 127 | H4 | 7.27 (1H, m) | Phe$^1$: H2/6, H3/5 | Phe$^1$: H2/6, H4 | Phe$^1$: C1, C2/6 |
| | C5 | 128.4 | H5 | 7.27 (1H, m) | Phe$^1$: H2/6, H4 | Phe$^1$: H2/6, H4 | Phe$^1$: C1, C2/6 |
| | C6 | 129.4 | H6 | 7.13 (1H, m) | Phe$^1$: H3/5, H4 | Phe$^1$: Hβ1/2, H3/5, H4 | Phe$^1$: C3/5, C4, C6 |
| | N(CH$_3$)2A | 40.4 | N(CH$_3$)2A | 2.95 (3H, s) | n/d | Phe$^1$: Hα, Hβ1/2 | n/d |
| | N(CH$_3$)2B | 41.8 | | | | | |
| | C=O | 165.2 | N(CH$_3$)2B | 2.8 (3H, s) | n/d | Phe$^1$: Hα, Hβ1/2, Pro$^2$: Hα | n/d |
| Pro$^2$ | α | 64.5 | α | 4.1 (1H, d, 6) | Pro$^2$: Hβ | Phe$^1$: N(CH3)2B; Pro$^2$: Hβ, Hγ2, Hδ2; Ile$^3$: NH; Tyr$^4$: H5 | Pro$^2$: Cβ |
| | β | 82.3 | β | 5.1 (1H, ddd, 6.3, 9.6) | Pro$^2$: Hα, Hγ1/2 | Pro$^2$: Hα, Hγ1/2, Hδ2; Tyr$^4$: H3 | Pro$^2$: C=O |
| | γ | 30.5 | | 2.28 (1H, m, 5.9) | Pro$^2$: Hβ, Hγ2 | | |
| | δ | 45.0 | γ1 | 1.95 (1H, m) | Pro$^2$: Hβ, Hγ1, Hδ1 | Pro$^2$: Hβ, Hγ2, Hδ1/2 | n/d |
| | | | γ2 | 3.84 (1H, t, 8.8) | Pro$^2$: Hγ2, Hδ2 | Pro$^2$: Hα, Hβ, Hγ1, Hδ1/2 | Pro$^2$: Cβ, Cδ |
| | | | δ1 | 2.13 (1H, m) | Pro$^2$: Hδ1 | Phe$^1$: Hα, N(CH3)2A/B; Pro$^2$: Hγ1/2, Hδ2 | Pro$^2$: Cα, Cβ, Cγ |
| | C=O | 169.2 | δ2 | 4.1 (1H, d, 6) | Pro$^2$: Hβ | Pro$^2$: Hβ, Hγ2, Hδ1 | n/d |
| Ile$^3$ | α | 57 | α | 3.94 (1H, d, 8.1) | Ile$^3$: Hβ, NH | Ile$^3$: Hβ, Hγ1A/B, Hγ2, NH; Tyr$^4$: NH | Ile3: Cβ, Cγ1, Cγ2, C=O |
| | β | 36.6 | β | 1.63 (1H, m) | Ile$^3$: Hα, Hγ1B, Hγ2 | Ile$^3$: Hα, Hγ1A/B, Hγ2, Hδ, NH | n/d |
| | γ1 | 23.7 | γ1A (CH$_2$) | 1.42 (1H, m) | Ile$^3$: Hβ, Hγ1B, Hδ | Ile$^3$: Hα, Hβ, Hγ1B, Hγ2, Hδ | n/d |
| | γ2 | 15.0 | γ1B (CH$_2$) | 1.16 (1H, m) | Ile$^3$: Hβ, Hγ1A, Hδ | Ile$^3$: Hα, Hβ, Hγ1A, Hγ2, Hδ, NH | n/d |
| | δ | 10.9 | γ2 | 0.79 (3H, d, 7.6) | Ile$^3$: Hβ | Ile$^3$: Hα, Hβ, Hγ1A/B, Hδ | Ile$^3$: Cα, Cβ, Cγ1 |
| | | | δ | 0.8 (3H, d, 7.6) | Ile$^3$: Hγ1A/B | Ile$^3$: Hβ, Hγ1A/B, Hγ2 | Ile$^3$: Cα, Cβ, Cγ1 |
| | C=O | 168.9 | NH | 7.22 (1H, d, 9) | Ile$^3$: Hα | Pro$^2$: Hα; Tyr$^4$: Hα, Hβ, Hγ1B | Pro$^2$: C=O, Ile$^3$: Cα |
| Tyr$^4$ | α | 126.2 | α | 6.24 (1H, d, 4.5, 7.1) | Tyr$^4$: Hβ, NH | Tyr$^4$: Hβ, NH | Tyr$^4$: C1 |
| | β | 125.7 | β | 6.6 (1H, d, 7.1) | Tyr$^4$: Hα | Tyr$^4$: Hα, H2/6 | Tyr$^4$: Cα, C6 |
| | C1 | 131.9 | H2 | 7 (1H, m) | Tyr$^4$: H3 | Tyr$^4$: H3, H5, H6, NH | Tyr$^4$: C1, C4 |
| | C2 | 129.6 | H3 | 7.1 (1H, d, 8.3) | Tyr$^4$: H2 | Pro$^2$: Hβ; Tyr$^4$: H2, H5, H6 | Tyr$^4$: C1, C4, C5 |
| | C3 | 118.4 | H5 | 7.02 (1H, m) | Tyr$^4$: H6 | Pro$^2$: Hα; Tyr$^4$: H2, H3, H6 | Tyr$^4$: C1, C2, C4 |
| | C4 | 156.4 | H6 | 6.98 (1H, d, 8.3) | Tyr$^4$: H5 | Tyr$^4$: H2, H3, H5 | Tyr$^4$: C2, C3, C4 |
| | C5 | 120.9 | | | | | |
| | C6 | 130.4 | NH | 7.83 (1H, br s) | Tyr$^4$: NH | Ile$^3$: Hα, Tyr$^4$: Hα | n/d |

### Table S6 | $^{1}H$ NMR (800 MHz) and $^{13}C$ NMR (200 MHz) data of cyFLLY (5) in DMSO-$d_6$.

| Residue | C | δ($^{13}$C) [ppm] | H | δ($^{1}$H) (int, m, *J* [Hz]) [ppm] | COSY | ROESY | HMBC |
|---|---|---|---|---|---|---|---|
| Phe¹ | α | 53.0 | α | 4.03 (1H, m) | Phe¹: Hβ1/2 | Phe¹: Hβ1/2, H2/6, Leu²: NH | Phe¹: Cβ |
| | β | 36.8 | β1 | 2.96 (1H, dd, 4.1, 14.4) | Phe¹: Hα, Hβ2 | Phe¹: Hα, Hβ2, H2/6 | Phe¹: Cα, C1, C2/6, C=O |
| | C1 | 134.4 | β2 | 2.78 (1H, dd, 8.3, 14.4) | Phe¹: Hα, Hβ1 | Phe¹: Hα, Hβ1, H2/6 | Phe¹: Cα, C1, C2/6, C=O |
| | C2 | 129.2 | H2 | 7.21 (1H, d, 7.4) | Phe¹: H3 | Phe¹: Hα, Hβ1/2, H3 | Phe¹: Cβ, C3, C4, C6 |
| | C3 | 128.4 | H3 | 7.29 (1H, t, 7.4) | Phe¹: H2, H4 | Phe¹: H2, H4 | Phe¹: C1, C2, C5 |
| | C4 | 127.0 | H4 | 7.26 (1H, t, 7.4) | Phe¹: H3/H5 | Phe¹: H3/H5 | Phe¹: C2/6, C3/5 |
| | C5 | 128.4 | H5 | 7.29 (1H, t, 7.4) | Phe¹: H2, H4 | Phe¹: H4, H6 | Phe¹: C1, C3, C6 |
| | C6 | 129.2 | H6 | 7.21 (1H, d, 7.4) | Phe¹: H5 | Phe¹: Hα, Hβ1/2, H5 | Phe¹: Cβ, C2, C4, C5 |
| | C=O | 166.7 | NH | n/d | - | - | - |
| Leu² | α | 54.8 | α | 4.48 (1H, t, 9.1) | Leu²: Hβ, NH | Leu²: Hβ, Hγ, Hδ1, NH; Leu³: NH; Tyr⁴: H5 | Phe¹: C=O; Leu²: Cβ, Cγ, C=O |
| | β | 79.2 | β | 4.67 (1H, d, 9.1) | Leu²: Hα, Hγ | Leu²: Hγ, Hδ2, NH; Tyr⁴: H3 | Leu²: Cα, Cγ, Cδ1/2, C=O; Tyr⁴: C4 |
| | γ | 27.6 | γ | 2.12 (1H, m) | Leu²: Hβ, Hδ1/2 | Phe¹: Hα; Leu²: Hα, Hβ, Hδ1/2 | - |
| | δ1 | 14.7 | δ1 | 0.94 (3H, d, 6.7) | Leu²: Hγ | Leu²: Hα, Hγ, Hδ2 | Leu²: Cβ, Cγ, Cδ2 |
| | δ2 | 20.2 | δ2 | 1.05 (3H, d, 6.7) | Leu²: Hγ | Leu²: Hβ, Hγ, Hδ1 | Leu²: Cβ, Cγ, Cδ1 |
| | C=O | 168.7 | NH | 8.77 (1H, d, 9.1) | Leu²: Hα | Phe¹: Hα; Leu²: Hα, Hβ, Hγ | Phe¹: C=O |
| Leu³ | α | 50.2 | α | 4.07 (1H, dd, 8.1, 12.1) | Leu³: Hβ2, NH | Leu³: Hβ, Hγ, Hδ1/2, NH; Tyr⁴: NH | Leu³: Cβ, Cγ, C=O |
| | β | 42.1 | β | 1.13 (2H, t, 7.0) | Leu³: Hα, Hγ | Leu³: Hα, Hγ, Hδ1/2 | Leu³: Cα, Cγ, C=O |
| | γ | 23.7 | γ | 1.31 (1H, m) | Leu³: Hβ, Hδ1/2 | Leu³: Hβ, Hδ1/2 | Leu³: Cβ, Cδ1/2 |
| | δ1 | 22.6 | δ1 | 0.77 (3H, d, 6.6) | Leu³: Hγ | Leu³: Hα, Hβ, Hγ, Hδ2 | Leu³: Cβ, Cγ, Cδ2 |
| | δ2 | 22.1 | δ2 | 0.75 (3H, d, 6.6) | Leu³: Hγ | Leu³: Hα, Hβ, Hγ, Hδ1 | Leu³: Cβ, Cγ, Cδ1 |
| | C=O | 169.3 | NH | 7.48 (1H, d, 9.1) | Leu³: Hα | Leu²: Hα; Leu³: Hα, Hβ | Leu²: C=O; Leu³: Cα |
| Tyr⁴ | α | 52.7 | α | 4.74 (1H, dt, 10.9, 5.7) | Tyr⁴: Hβ1/2, NH | Tyr⁴: Hβ1, H2 | Tyr⁴: Cβ, C=O |
| | β | 36.5 | β1 | 3.22 (1H, dd, 5.7, 13.2) | Tyr⁴: Hα, Hβ2 | Tyr⁴: Hα, Hβ2, H2, H6 | Tyr⁴: C1, C6 |
| | C1 | 130.4 | β2 | 2.47 (1H, t, 12.6) | Tyr⁴: Hα, Hβ1 | Tyr⁴: Hβ1, H6 | Tyr⁴: Cα, C1, C2, C6, C=O |
| | C2 | 131.5 | H2 | 6.91 (1H, dd, 2.2, 8.5) | Tyr⁴: H3 | Tyr⁴: Hα, Hβ2, H3 | Tyr⁴: Cβ, C4, C6 |
| | C3 | 119.1 | H3 | 6.70 (1H, dd, 2.2, 8.5) | Tyr⁴: H2 | Leu²: Hβ; Tyr⁴: H2 | Tyr⁴: C2, C4, C5 |
| | C4 | 155.4 | H5 | 6.74 (1H, dd, 2.2, 8.5) | Tyr⁴: H6 | Leu²: Hα; Tyr⁴: H6 | Tyr⁴: C2, C3, C4 |
| | C5 | 113.5 | H6 | 6.98 (1H, dd, 2.2, 8.5) | Tyr⁴: H5 | Tyr⁴: Hβ1, H5 | Tyr⁴: Cβ, C2, C4 |
| | C6 | 129.3 | | | | | |
| | C=O | 172.5 | NH | 7.83 (1H, d, 10.1) | Tyr⁴: Hα | Leu³: Hα | Leu³: C=O |

### Table S7 | $^{1}$H NMR (800 MHz) and $^{13}$C NMR (200 MHz) data of cyLLIY (6) in DMSO-$d_6$.

| Residue | C | δ($^{13}$C) [ppm] | H | δ($^{1}$H) (int, m, *J* [Hz]) [ppm] | COSY | ROESY | HMBC |
|---|---|---|---|---|---|---|---|
| Leu$^{1}$ | α | 50.4 | α | 3.8 (1H, m) | Leu$^{1}$: Hβ1/2 | Leu$^{1}$: Hβ1/2, Hγ, Hδ1/2, Leu$^{2}$: NH, Hγ | Leu$^{1}$: Cβ1/2, Cγ, C=O |
| | β | 40.2 | β1 | 1.44 (1H, m) | Leu$^{1}$: Hα, Hβ2, Hγ | Leu$^{1}$: Hα, Hβ2, Hγ, Hδ1/2 | Leu$^{1}$: Cα, Cγ, Cδ1/2, C=O |
| | γ | 23.2 | β2 | 1.38 (1H, m) | Leu$^{1}$: Hα, Hβ1, Hγ | Leu$^{1}$: Hα, Hβ1, Hγ, Hδ1/2 | Leu$^{1}$: Cα, Cγ, Cδ1/2, C=O |
| | δ1 | 21.6 | γ | 1.55 (1H, m) | Leu$^{1}$: Hβ1/2, Hδ1/2 | Leu$^{1}$: Hα, Hβ1/2, Hδ1/2 | - |
| | δ2 | 22.4 | δ1 | 0.83 (3H, m) | Leu$^{1}$: Hγ | Leu$^{1}$: Hα, Hβ1/2, Hγ, Hδ2 | Leu$^{1}$: Cβ, Cγ, Cδ2 |
| | | | δ2 | 0.84 (3H, m) | Leu$^{1}$: Hγ | Leu$^{1}$: Hα, Hβ1/2, Hγ, Hδ1 | Leu$^{1}$: Cβ, Cγ, Cδ1 |
| | C=O | 167.6 | NH | 8.17 (2H, m) | - | Leu$^{1}$: Hα | - |
| Leu$^{2}$ | α | 54.7 | α | 4.54 (1H, m) | Leu$^{2}$: Hβ, NH | Leu$^{2}$: Hβ, Hγ, Hδ1; Ile$^{3}$: NH; Tyr$^{4}$: H3 | - |
| | β | 79.3 | β | 4.70 (1H, m) | Leu$^{2}$: Hα, Hγ | Leu$^{2}$: Hγ, Hδ2, NH; Tyr$^{4}$: H5 | Leu$^{2}$: Cγ, Cδ1/2, C=O; Tyr$^{4}$: C4 |
| | γ | 27.4 | γ | 2.16 (1H, m) | Leu$^{2}$: Hδ1/2 | Leu$^{1}$: Hα; Leu$^{2}$: Hα, Hβ, Hδ1/2, NH | - |
| | δ1 | 14.8 | δ1 | 0.94 (1H, m) | Leu$^{2}$: Hγ | Leu$^{2}$: Hα, Hγ, Hδ2, Tyr$^{4}$: H3 | Leu$^{2}$: Cβ, Cγ, Cδ2 |
| | δ2 | 20.3 | δ2 | 1.05 (1H, m) | Leu$^{2}$: Hγ | Leu$^{2}$: Hβ, Hγ, Hδ1 | Leu$^{2}$: Cβ, Cγ, Cδ1 |
| | C=O | 169.5 | NH | 8.78 (1H, m) | Leu$^{2}$: Hα | Leu$^{1}$: Hα, Leu$^{2}$: Hα, Hβ, Hγ | - |
| Ile$^{3}$ | α | 56.3 | α | 3.75 (1H, m) | Leu$^{3}$: Hβ, NH | Leu$^{3}$: Hβ, Hγ1/2, Hδ, NH; Tyr$^{4}$: NH, H2 | Leu$^{3}$: Cβ, Cγ1/2, C=O |
| | β | 36.8 | β | 1.31 (1H, m) | Leu$^{3}$: Hα, Hγ1/2 | Leu$^{3}$: Hα, Hγ1/2, Hδ, NH | Leu$^{3}$: Cα, Cγ2 |
| | γ1 | 14.7 | γ1 | 0.68 (3H, m) | Leu$^{3}$: Hβ | Leu$^{3}$: Hα, Hγ2, Hδ | Leu$^{3}$: Cα, Cβ, Cγ2 |
| | γ2 | 23.6 | γ2 | 1.21 (2H, m) | Leu$^{3}$: Hβ, Hδ | Leu$^{3}$: Hα, Hβ, Hγ1, Hδ | Leu$^{3}$: Cβ, Cγ1, Cδ |
| | δ | 21.8 | δ | 0.80 (3H, m) | Leu$^{3}$: Hγ2 | Leu$^{3}$: Hα, Hβ, Hγ1/2 | Leu$^{3}$: Cβ, Cγ2 |
| | C=O | 169.3 | NH | 7.37 (1H, m) | Leu$^{3}$: Hα | Leu$^{2}$: Hα; Leu$^{3}$: Hα, Hβ | - |
| Tyr$^{4}$ | α | 52.6 | α | 4.78 (1H, m) | Tyr$^{4}$: Hβ1/2, NH | Tyr$^{4}$: Hβ1, H6 | Tyr$^{4}$: Cβ, C=O |
| | β | 36.4 | β1 | 3.21 (1H, m) | Tyr$^{4}$: Hα, Hβ2 | Tyr$^{4}$: Hα, Hβ2, H2, H6 | Tyr$^{4}$: C1 |
| | C1 | 130.4 | β2 | 2.45 (1H, t, 12.0) | Tyr$^{4}$: Hα, Hβ1 | Tyr$^{4}$: Hβ1, H2 | Tyr$^{4}$: Cα, C1, C2, C6, C=O |
| | C2 | 129.1 | H2 | 6.98 (1H, d, 7.3) | Tyr$^{4}$: H3 | Ile$^{3}$: Hα, Tyr$^{4}$: Hβ2, H3 | Tyr$^{4}$: Cβ, C6, C4 |
| | C3 | 119.2 | H3 | 6.71 (1H, d, 7.3) | Tyr$^{4}$: H2 | Leu$^{2}$: Hα, Tyr$^{4}$: H2 | Tyr$^{4}$: C4, C5, C6 |
| | C4 | 155.4 | H5 | 6.74 (1H, d, 7.3) | Tyr$^{4}$: H6 | Leu$^{2}$: Hβ, Hδ1; Tyr$^{4}$: H6 | Tyr$^{4}$: C1, C3, C4, C6 |
| | C5 | 113.9 | H6 | 6.90 (1H, d, 7.3) | Tyr$^{4}$: H5 | Tyr$^{4}$: Hα, Hβ1, H5 | Tyr$^{4}$: Cβ, C2, C4 |
| | C6 | 131.5 | | | | | |
| | C=O | 172.5 | NH | 7.9 (1H, m) | Tyr$^{4}$: Hα | Ile$^{3}$: Hα | - |

### Table S8 | $^{1}$H NMR (800 MHz) and $^{13}$C NMR (200 MHz) data of sanjoinine A in DMSO-$d_6$.

| Residue | C | δ($^{13}$C) [ppm] | H | δ($^{1}$H) (int, m, *J* [Hz]) [ppm] | COSY | ROESY | HMBC |
|---|---|---|---|---|---|---|---|
| Phe$^{1}$ | α | 67.2 | α | 3.36 (1H, dt, 5.2, 9.0) | Phe$^{1}$: Hβ1/2 | Phe$^{1}$: Hβ1/2, N(Me), H2/6, Leu$^{2}$: NH | Phe$^{1}$: Cβ, N(Me), C=O |
| | β | 33.2 | β1 | 2.83 (1H, dd, 9.0, 14.3) | Phe$^{1}$: Hα, Hβ2 | Phe$^{1}$: Hα, Hβ2, H2/6 | Phe$^{1}$: Cα, C1, C2/6, C=O |
| | C1 | 139.4 | β2 | 2.63 (1H, dd, 5.2, 14.3) | Phe$^{1}$: Hα, Hβ1 | Phe$^{1}$: Hα, Hβ1, H2/6 | Phe$^{1}$: Cα, C1, C2/6 |
| | C2 | 128.6 | H2 | 7.18 (2H, d, 7.4) | Phe$^{1}$: H3 | Phe$^{1}$: Hα, Hβ1/2, N(Me) | Phe$^{1}$: Cβ, C3, C4 |
| | C3 | 127.3 | H3 | 7.24 (2H, t, 7.4) | Phe$^{1}$: H2, H4 | Phe$^{1}$: H2, H4 | Phe$^{1}$: C1, C2, C5 |
| | C4 | 125.6 | H4 | 7.14 (1H, t, 7.4) | Phe$^{1}$: H3/H5 | Phe$^{1}$: H3/H5 | Phe$^{1}$: C2/6, C3/5 |
| | C5 | 127.3 | H5 | 7.24 (2H, t, 7.4) | Phe$^{1}$: H2, H4 | Phe$^{1}$: H4, H6 | Phe$^{1}$: C1, C3, C6 |
| | C6 | 128.6 | H6 | 7.18 (2H, d, 7.4) | Phe$^{1}$: H5 | Phe$^{1}$: Hα, Hβ1/2, N(Me) | Phe$^{1}$: Cβ, C3, C4 |
| | C=O | 169.8 | | | | | |
| | N(Me) | 40.7 | N(Me) | 2.24 (6H, s) | - | | Phe$^{1}$: Cα |
| Leu$^{2}$ | α | 54.6 | α | 4.48 (1H, dd, 7.9, 9.5) | Leu$^{2}$: Hβ, NH | Leu$^{2}$: Hβ, Hγ, Hδ1, NH; Leu$^{3}$: NH; Tyr$^{4}$: H5 | Phe$^{1}$: C=O; Leu$^{2}$: Cβ, C=O |
| | β | 81.9 | β | 4.82 (1H, d, 7.7) | Leu$^{2}$: Hα, Hγ | Leu$^{2}$: Hγ, Hδ2, NH; Tyr$^{4}$: H3 | Leu$^{2}$: Cα, Cγ, Cδ1/2, C=O; Tyr$^{4}$: C4 |
| | γ | 27.8 | γ | 2.18 (1H, q, 6.7) | Leu$^{2}$: Hβ, Hδ1/2 | Leu$^{2}$: Hα, Hβ, Hδ1/2 | - |
| | δ1 | 14.8 | δ1 | 0.94 (1H, d, 6.7) | Leu$^{2}$: Hγ | Leu$^{2}$: Hα, Hγ, Hδ2 | Leu$^{2}$: Cβ, Cγ, Cδ2 |
| | δ2 | 20.3 | δ2 | 1.15 (1H, d, 6.7) | Leu$^{2}$: Hγ | Leu$^{2}$: Hβ, Hγ, Hδ1 | Leu$^{2}$: Cβ, Cγ, Cδ1 |
| | C=O | 170.9 | NH | 8.35 (1H, d, 9.5) | Leu$^{2}$: Hα | Phe$^{1}$: Hα; Leu$^{2}$: Hα, Hβ, Hγ | Phe$^{1}$: Cα, C=O |
| Leu$^{3}$ | α | 51.5 | α | 3.88 (1H, dt, 4.5, 8.8) | Leu$^{3}$: Hβ, NH | Leu$^{3}$: Hβ1/2, Hγ, Hδ1/2, NH; Tyr$^{4}$: NH | Leu2: C=O; Leu$^{3}$: Cβ, Cγ, C=O |
| | β | 39.2 | β1 | 1.26 (1H, m) | Leu$^{3}$: Hα, Hγ | Leu$^{3}$: Hα, Hβ2, Hγ, Hδ1/2, NH | Leu$^{3}$: Cα, Cδ1/2, C=O |
| | γ | 23.4 | β2 | 1.33 (1H, m) | Leu$^{3}$: Hα, Hγ | Leu$^{3}$: Hα, Hβ1, Hγ, Hδ1/2, NH | Leu$^{3}$: Cγ |
| | δ1 | 20.4 | γ | 1.35 (1H, m) | Leu$^{3}$: Hβ, Hδ1/2 | Leu$^{3}$: Hα, Hβ1/2, Hδ1/2, NH | Leu$^{3}$: Cβ |
| | δ2 | 22.7 | δ1 | 0.62 (3H, d, 6) | Leu$^{3}$: Hγ | Leu$^{3}$: Hα, Hβ1/2, Hγ, Hδ2, NH | Leu$^{3}$: Cβ, Cγ, Cδ2 |
| | | | δ2 | 0.69 (3H, d, 6) | Leu$^{3}$: Hγ | Leu$^{3}$: Hα, Hβ1/2, Hγ, Hδ1, NH | Leu$^{3}$: Cβ, Cγ, Cδ1 |
| | C=O | 168.2 | NH | 7.68 (1H, d, 7.1) | Leu$^{3}$: Hα | Leu$^{3}$: Hα, Hβ1/2, Hγ, Hδ1/2 | - |
| Tyr$^{4}$ | α | 125.8 | α | 6.42 (1H, t, 7.0) | Tyr$^{4}$: Hβ | Tyr$^{4}$: Hβ, H2, NH | Tyr$^{4}$: Cβ, C1 |
| | β | 119.4 | β | 6.48 (1H, d, 7.4) | Tyr$^{4}$: Hα | Tyr$^{4}$: Hα, H2, H6, NH | Tyr$^{4}$: Cα, C1 |
| | C1 | 130.4 | H2 | 6.94 (1H, dd, 2.2, 8.5) | Tyr$^{4}$: H3 | Tyr$^{4}$: Hα, Hβ, H3 | Tyr$^{4}$: Cβ, C4, C6 |
| | C2 | 130.4 | H3 | 7.07 (1H, dd, 2.2, 8.5) | Tyr$^{4}$: H2 | Leu$^{2}$: Hβ; Tyr$^{4}$: H2 | Tyr$^{4}$: C2, C4, C5 |
| | C3 | 119.8 | H5 | 7.05 (1H, dd, 2.2, 8.5) | Tyr$^{4}$: H6 | Leu$^{2}$: Hα; Tyr$^{4}$: H6 | Tyr$^{4}$: C2, C3, C4 |
| | C4 | 155.7 | H6 | 7.00 (1H, dd, 2.2, 8.5) | Tyr$^{4}$: H5 | Tyr$^{4}$: Hβ, H5 | Tyr$^{4}$: Cβ, C1, C2, C4 |
| | C5 | 121.9 | | | | | |
| | C6 | 129.8 | NH | 6.92 (1H, m) | Tyr$^{4}$: Hα | Leu$^{3}$: Hα | - |

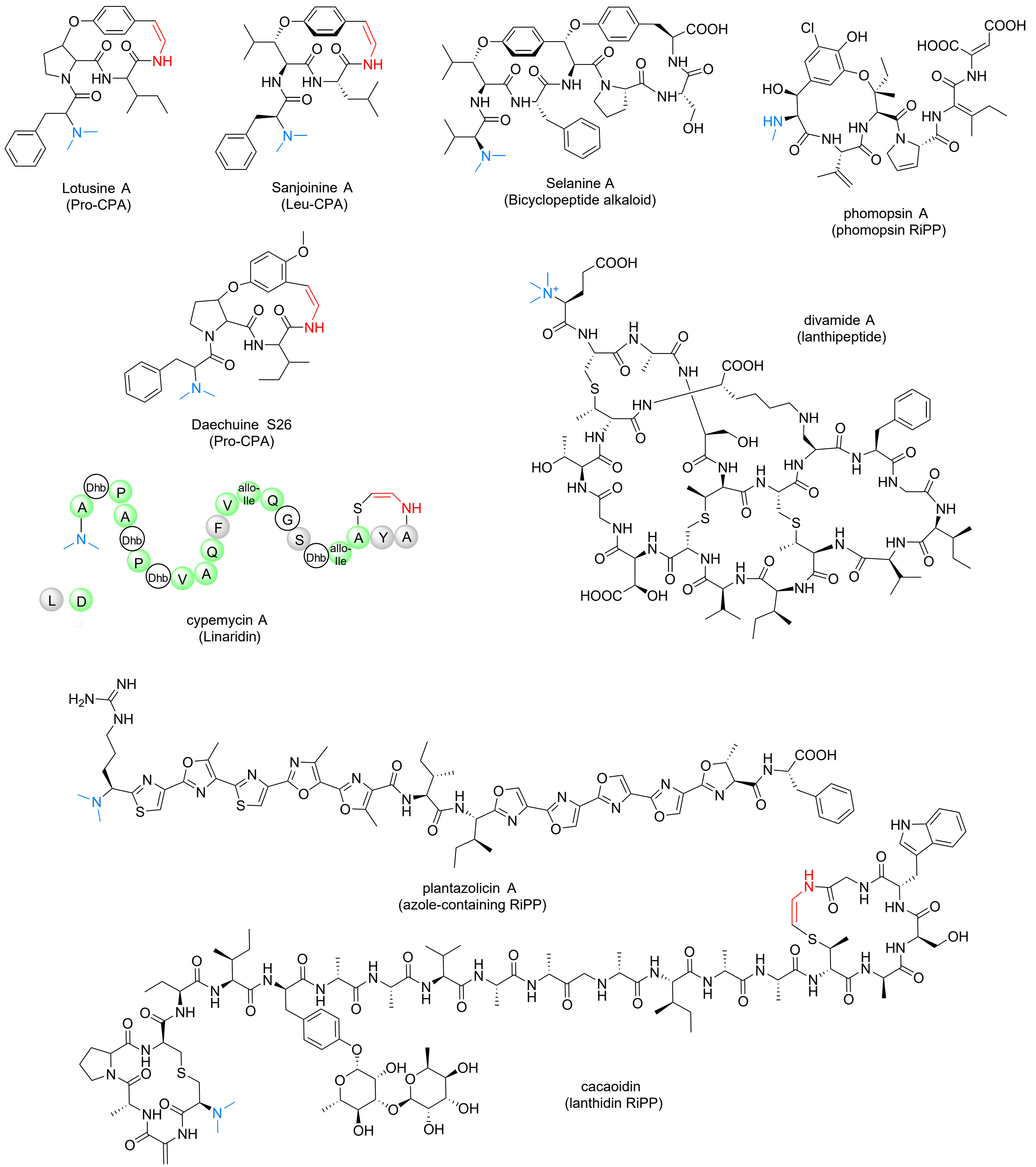


**Figure S1 | Chemical structures of representative RiPPs with α-*N*-methylations (highlighted in blue color) and/or decarboxylated C-termini (highlighted in red color).**

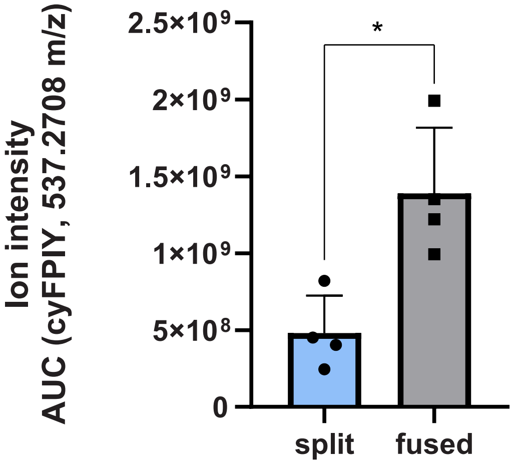


split = ZjuPrec-5xFPIY + ZjuBURP

>ZjuPrec-5xFPIY
**MKSFFALLAFSSLLLLSSIVIA**RKEPVEYVKTVVEQVAKDLSIDP
SSRFPIYHDNQNGQSVDTSSRFPIYHDNQNGQSIDPSSRFPIYHG
NQNGQSIDPSSRFPIYHGNQNGQSIDPSSRFPIYHGNQNGDQNSK
ANEEGAAKSASTDQ

>ZjuBURP
**MAKEIASCVLLIVYLSLLMLCSTGNG**HPQHHHQQQYGGDDPLDMA
RRVGFFTVDDLYVGKAVGIQFYTRDPSSLPPFLSKEVADRIPFSV
KELPLILQLFGMAPGSPEAQLAERTLRTCAEEPIKGEQKTCATSI
ESFIEFASSMLGGGQYGVDFRAIKTTHLGKPVSVYQNYTFLDIKE
VHSPIMVACHIMDYPYIVYMCHSQTSRVYQIKIVGQEVGDALNIA
VCHIDTSQWAPDHVSFKLLSVKPGTVSICHFFGPHNPVWVKN

fused = ZjuPrec-5xFPIY-ZjuBURP-fusion

>ZjuPrec-5xFPIY-ZjuBURP-fusion
**MKSFFALLAFSSLLLLSSIVIA**RKEPVEYVKTVVEQVAKDLSIDP
SSRFPIYHDNQNGQSVDTSSRFPIYHDNQNGQSIDPSSRFPIYHG
NQNGQSIDPSSRFPIYHGNQNGQSIDPSSRFPIYHGNQNHPQHHH
QQQYGGDDPLDMARRVGFFTVDDLYVGKAVGIQFYTRDPSSLPPF
LSKEVADRIPFSVKELPLILQLFGMAPGSPEAQLAERTLRTCAEE
PIKGEQKTCATSIESFIEFASSMLGGGQYGVDFRAIKTTHLGKPV
SVYQNYTFLDIKEVHSPIMVACHIMDYPYIVYMCHSQTSRVYQIK
IVGQEVGDALNAIAVCHIDTSQWAPDHVSFKLLSVKPGTVSICHF

**Figure S2 | Transient gene expression analysis of fused and split ZjuBURP constructs in *Nicotiana benthamiana.*** Comparison of EIC AUC (area under the curve) values of cyFPIY from methanolic extracts of transgenic *N. benthamiana* leaves 7 days after infiltration with either split ZjuPrec-5xFPIY and ZjuBURP (split) or ZjuPrec-5xFPIY-ZjuBURP-fusion (fused). Agroinfiltration occured from cultures with the same final $OD_{600}$ (0.8) in plants of the same age. The bar graphs represent means (n=4) of AUC values with the error bar representing one standard deviation. Split versus fused data was tested for significance by unpaired t-test (* $p < 0.05$). The protein sequences of split proteins and the fused protein are shown with the BURP-domain sequence underlined, the signal peptide shown in bold, the core peptides highlighted in red color and the fusion residues highlighted in blue color.

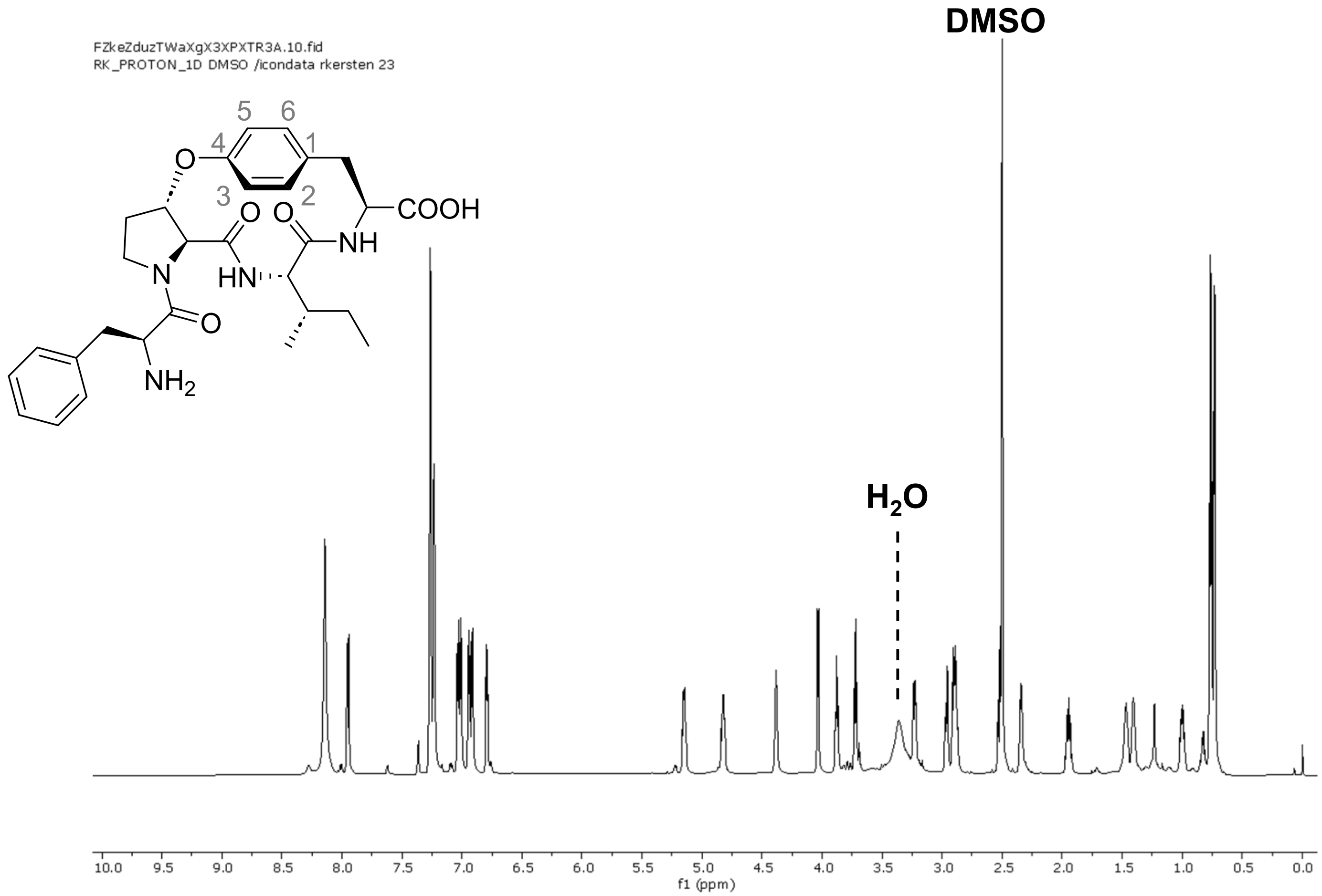


**Figure S3 | $^{1}H$ NMR spectrum of cyFPIY in DMSO-$d_6$ (800 MHz).**

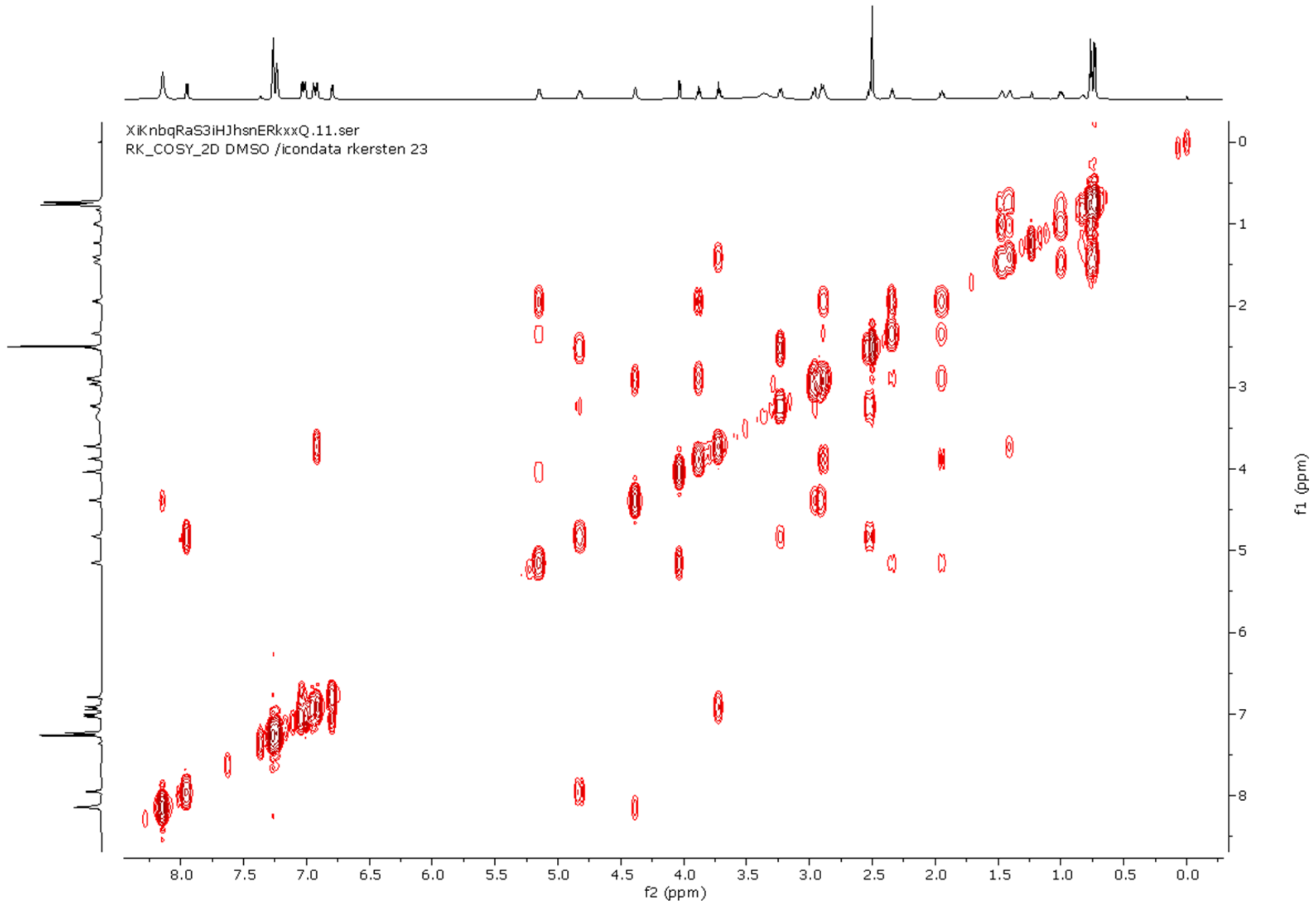


**Figure S4 | $^{1}H$-$^{1}H$ COSY NMR spectrum of cyFPIY in DMSO-$d_6$.**

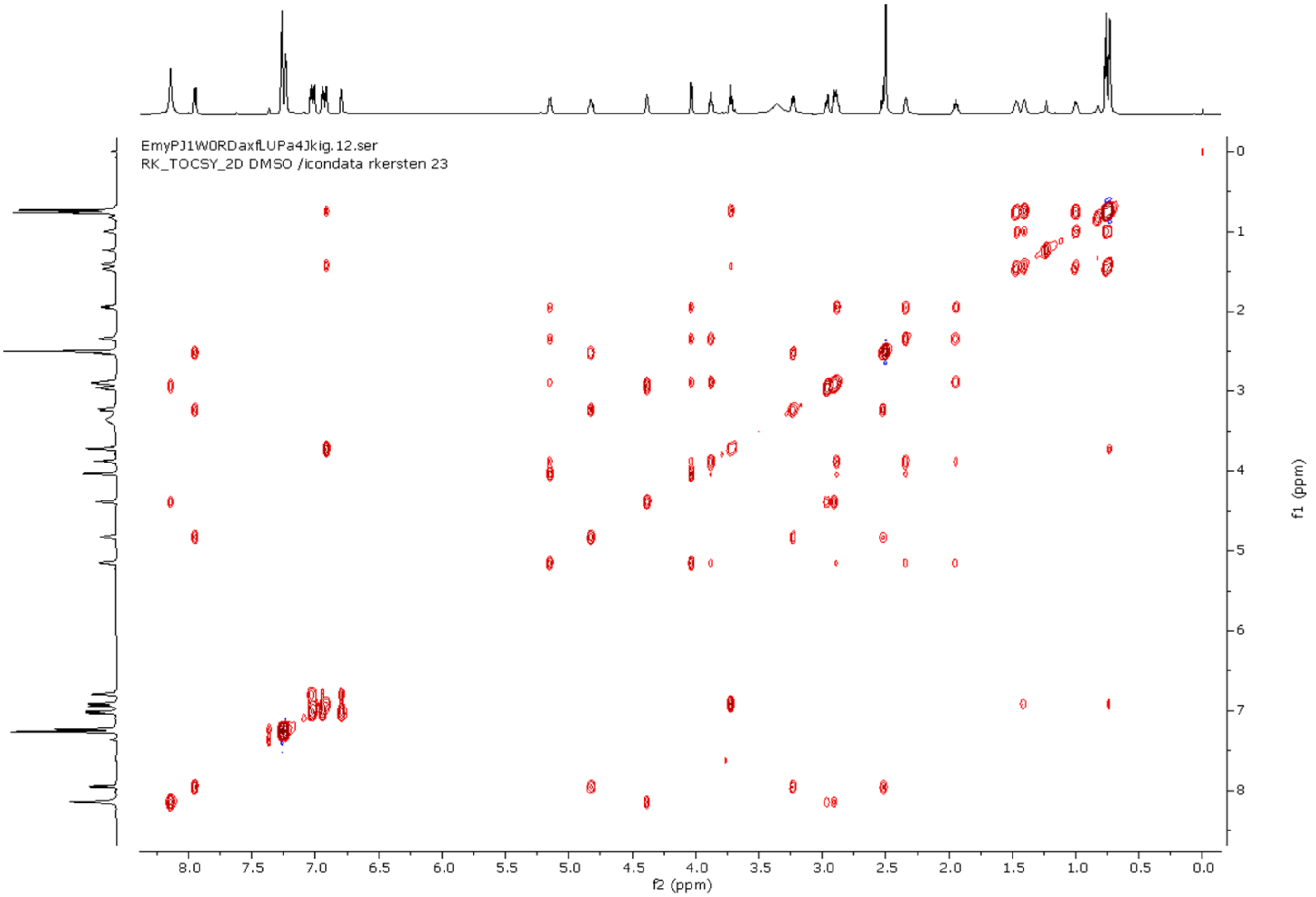


**Figure S5 | $^{1}H$-$^{1}H$ TOCSY NMR spectrum of cyFPIY in DMSO-d$_{6}$.**

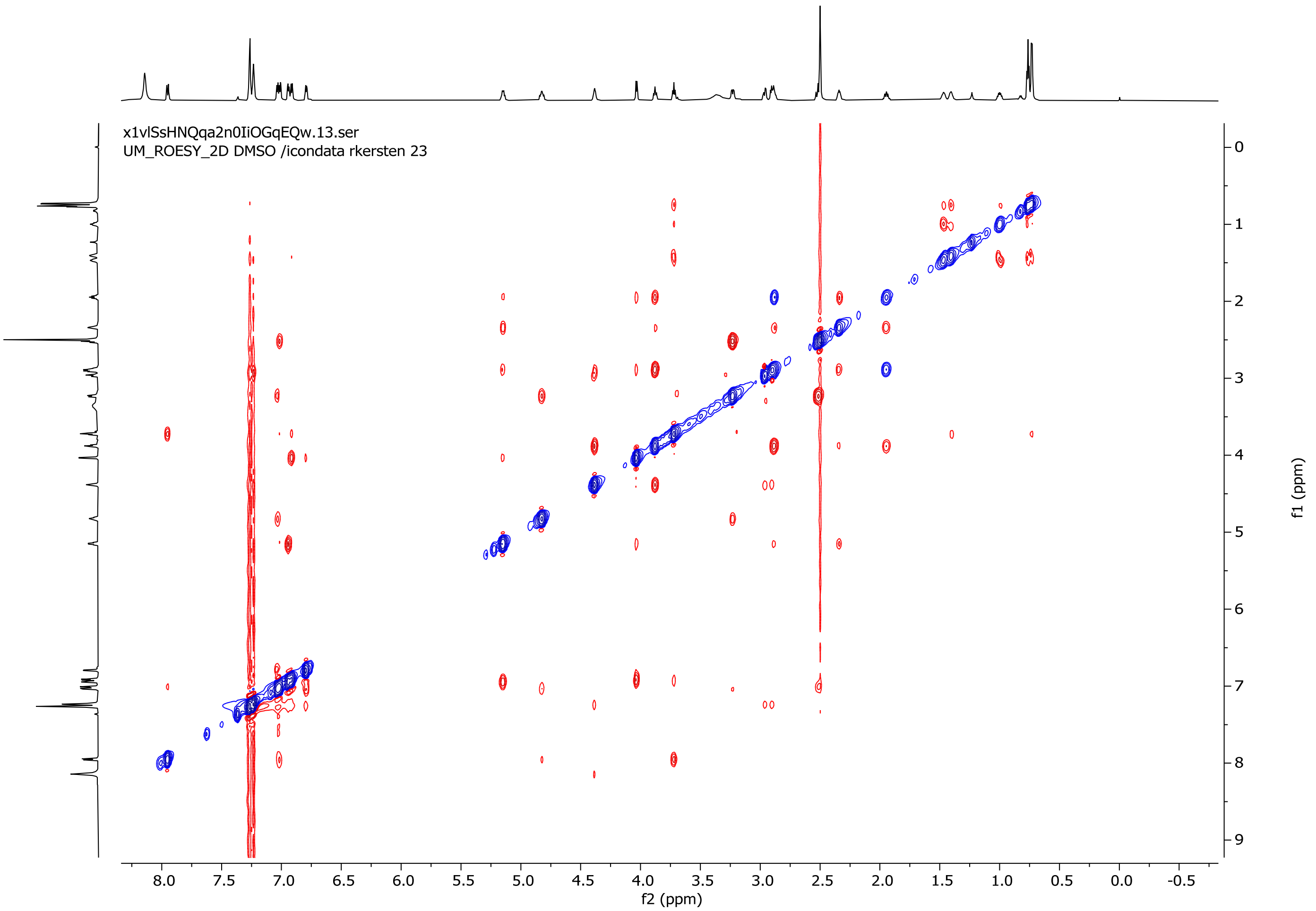


**Figure S6 | ROESY NMR spectrum of cyFPIY in DMSO-d$_{6}$.**

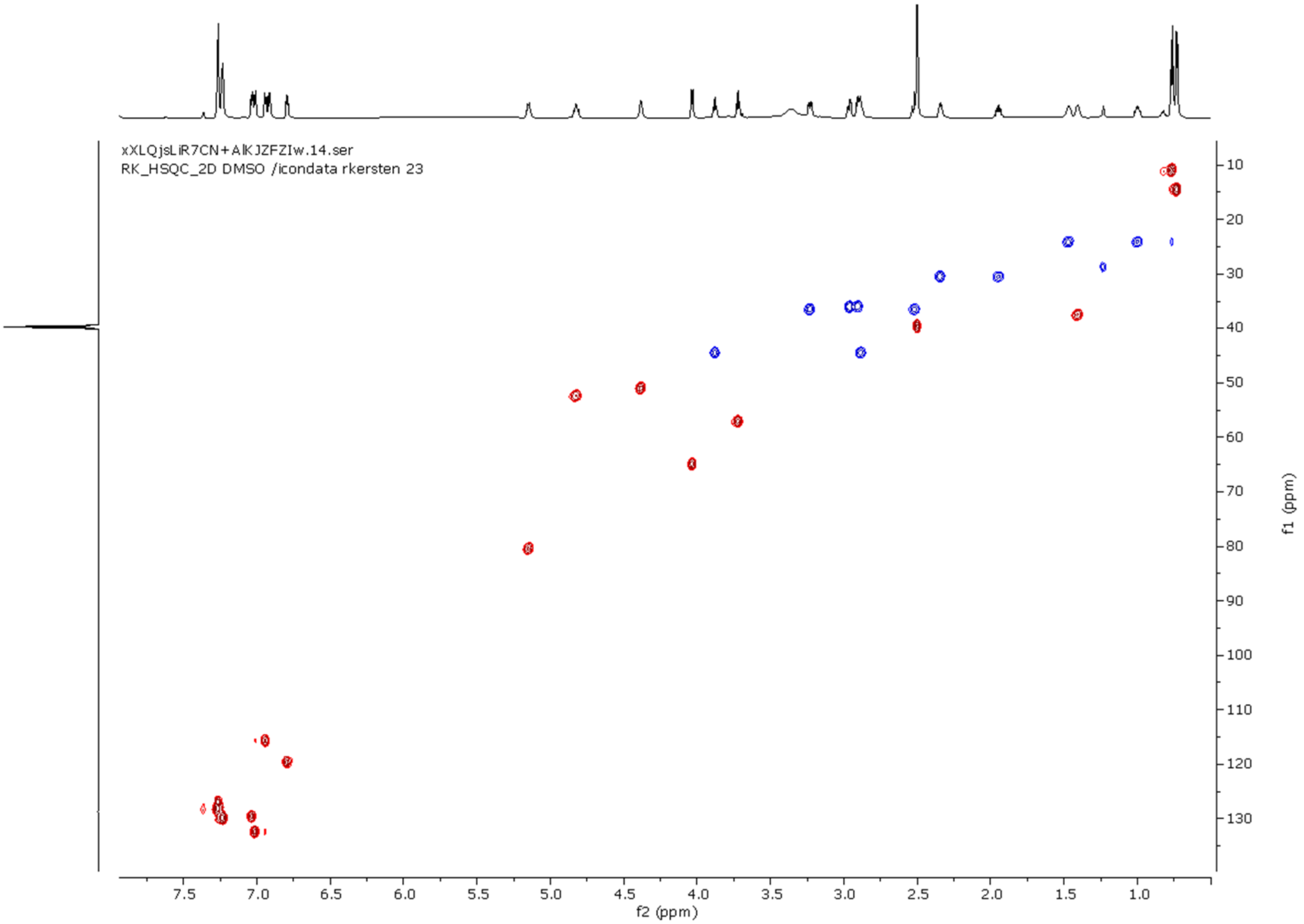


**Figure S7 | HSQC NMR spectrum of cyFPIY in DMSO-d$_6$.**

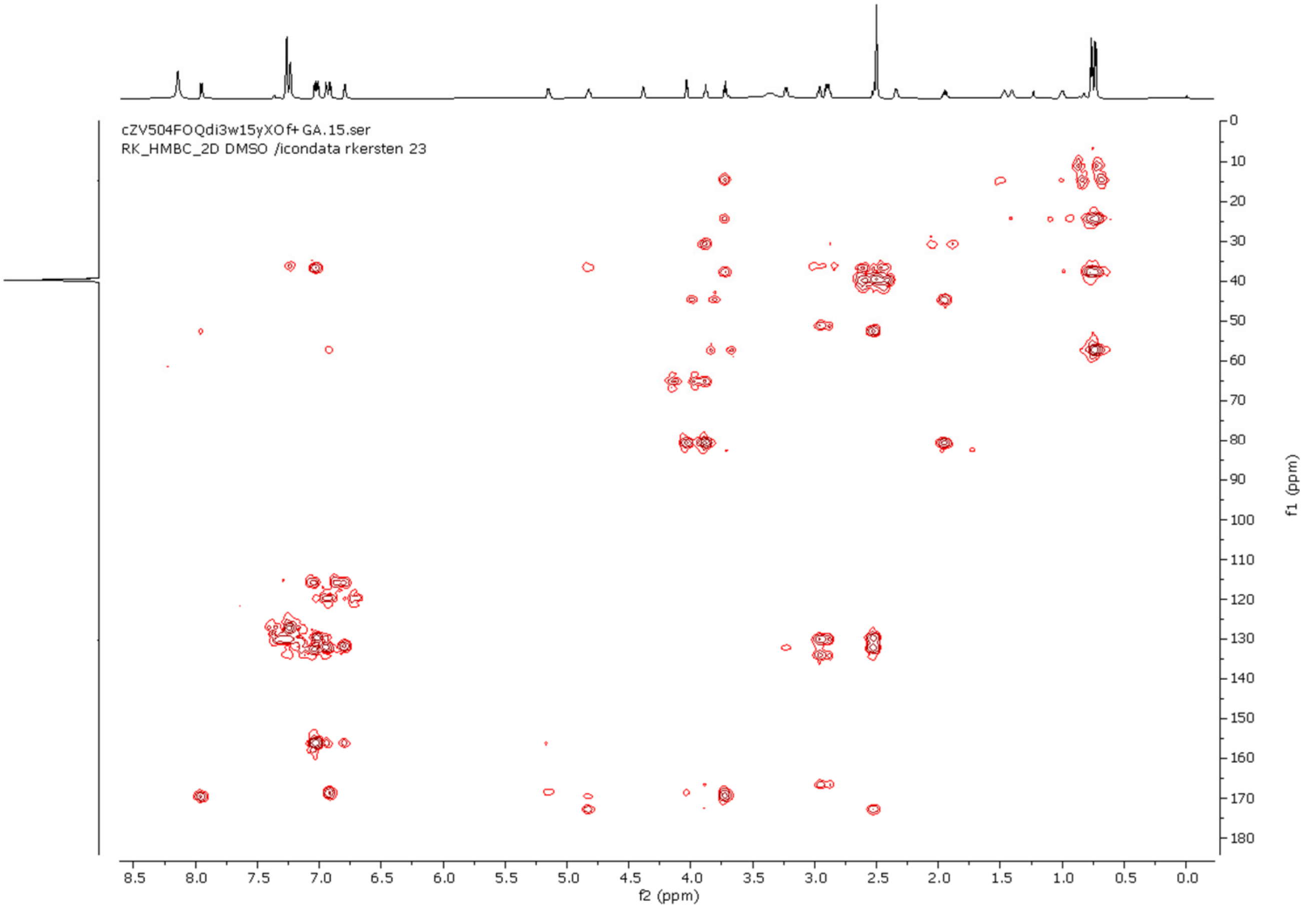


**Figure S8 | HMBC NMR spectrum of cyFPIY in DMSO-d$_6$.**

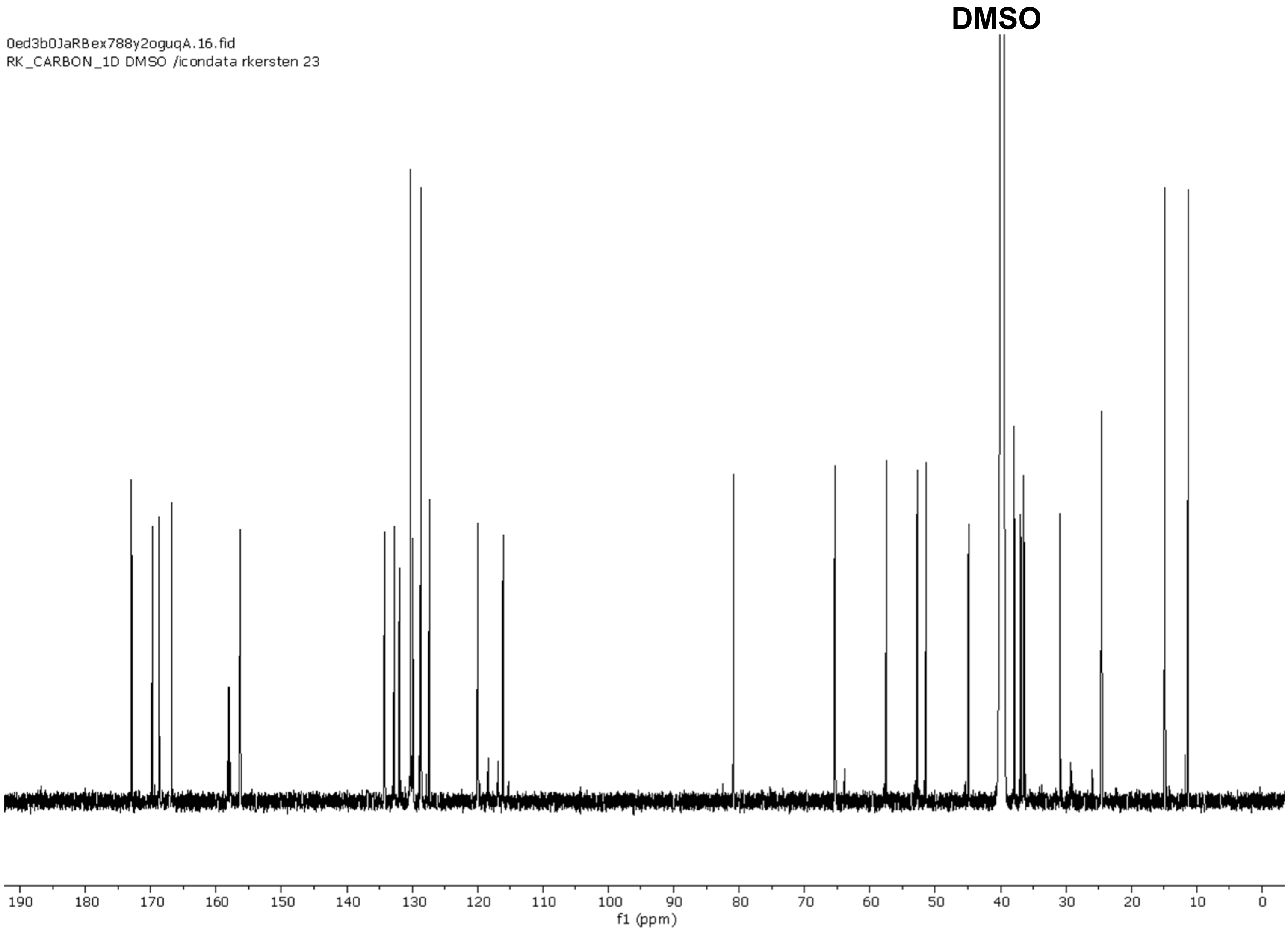


**Figure S9 | $^{13}$C NMR spectrum of cyFPIY in DMSO-d$_6$ (200 MHz).**

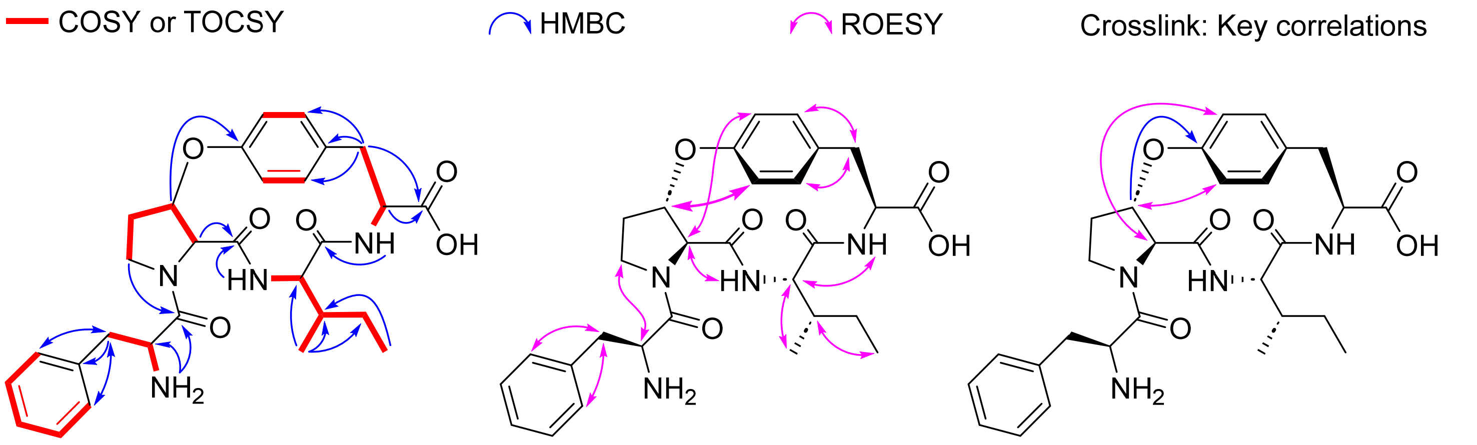


**Figure S10 | Key 2D NMR correlations of cyFPIY in DMSO-d$_6$.**

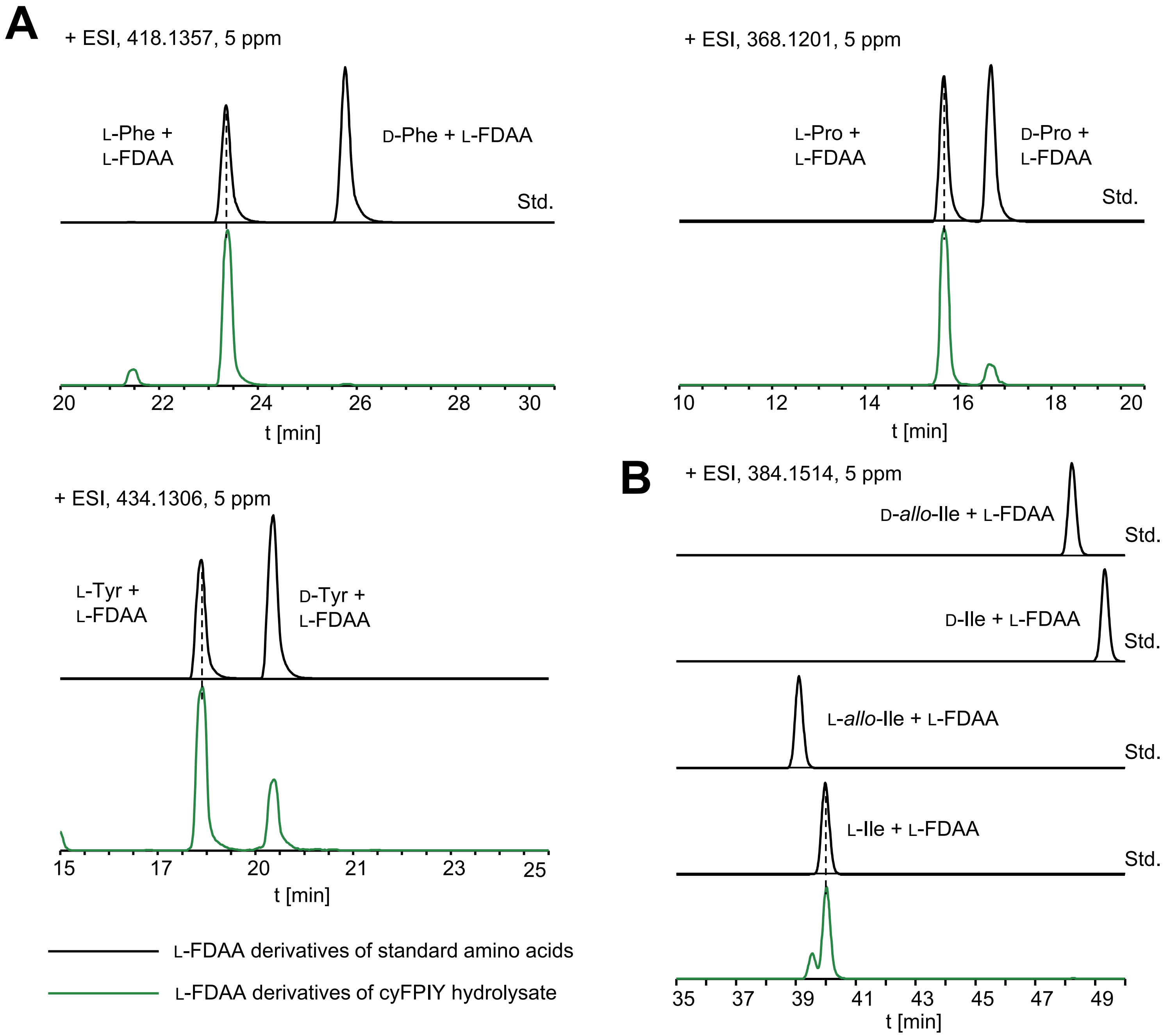


**Figure S11 | Marfey's analysis of cyFPIY.** (**A**) The Phe, Pro, and Tyr residues in cyFPIY were assigned as L by using Marfey's analysis. (**B**) The Ile residue in cyFPIY was assigned as L by using C3-Marfey's analysis.

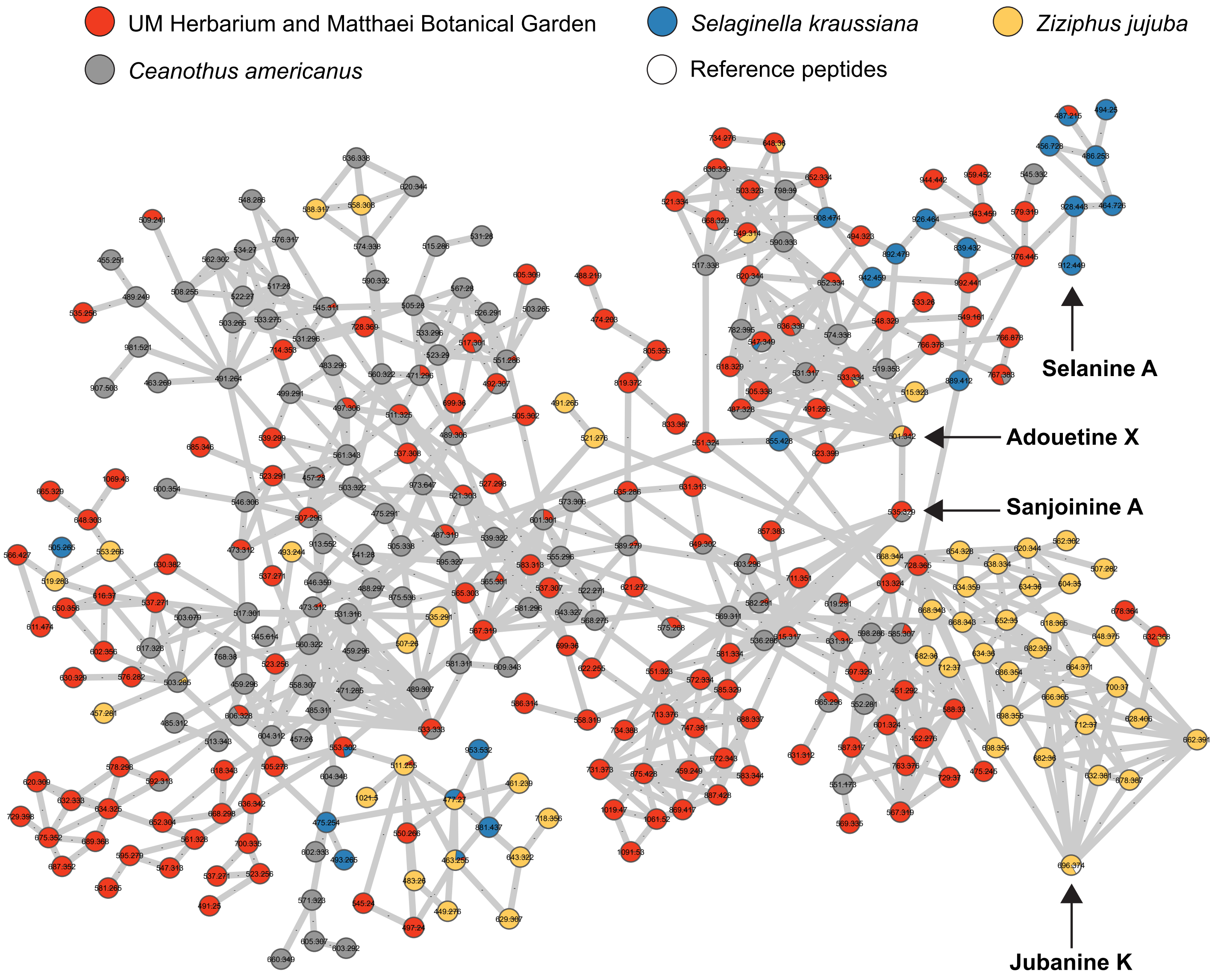


**Figure S12 | Cyclopeptide alkaloid-containing spectral cluster of peptide-filtered LC-MS/MS datasets from specific plant sources.** Nodes are colored by plant source listed above with precursor ion mass. Pie charts represent relative spectral counts of defined source materials. The relevant nodes for cyclic peptides are indicated with arrows, and reference peptides in spectral cluster are verified by NMR. Edges are default cosine values >0.7.

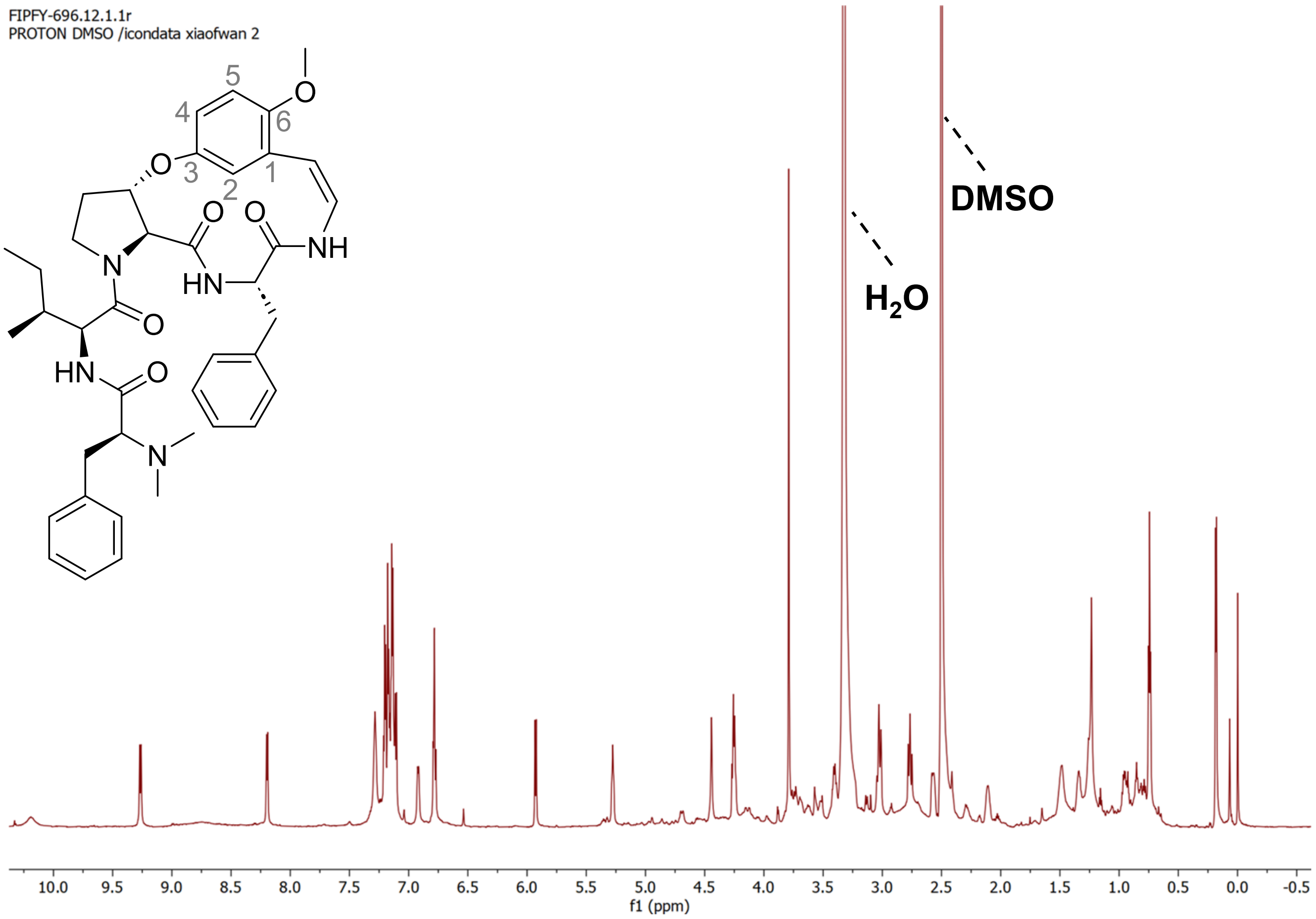


**Figure S13 | [1]H NMR spectrum of jubanine K in DMSO-$d_6$ (800 MHz).**

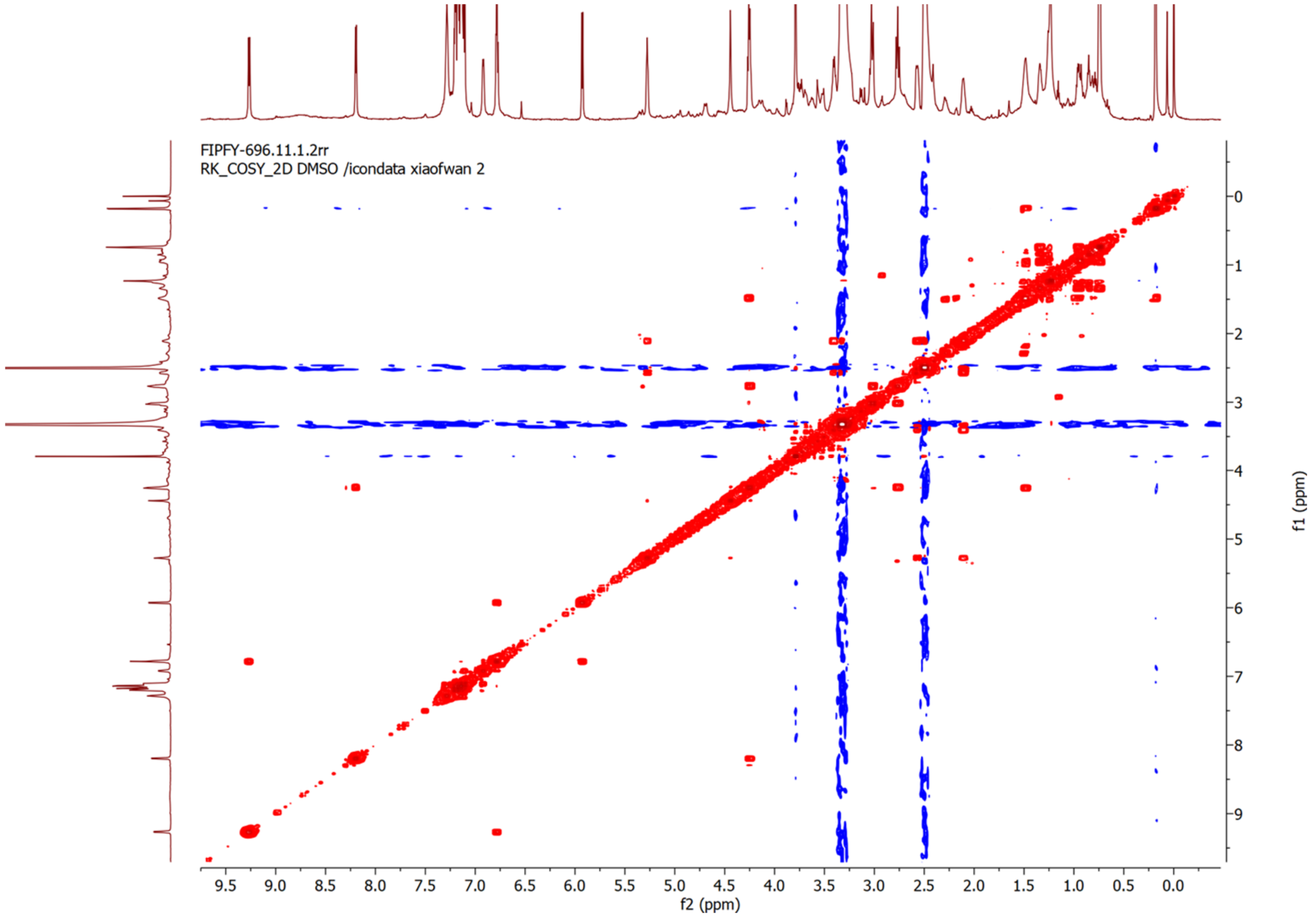


**Figure S14 | [1]H-[1]H COSY NMR spectrum of jubanine K in DMSO-$d_6$.**

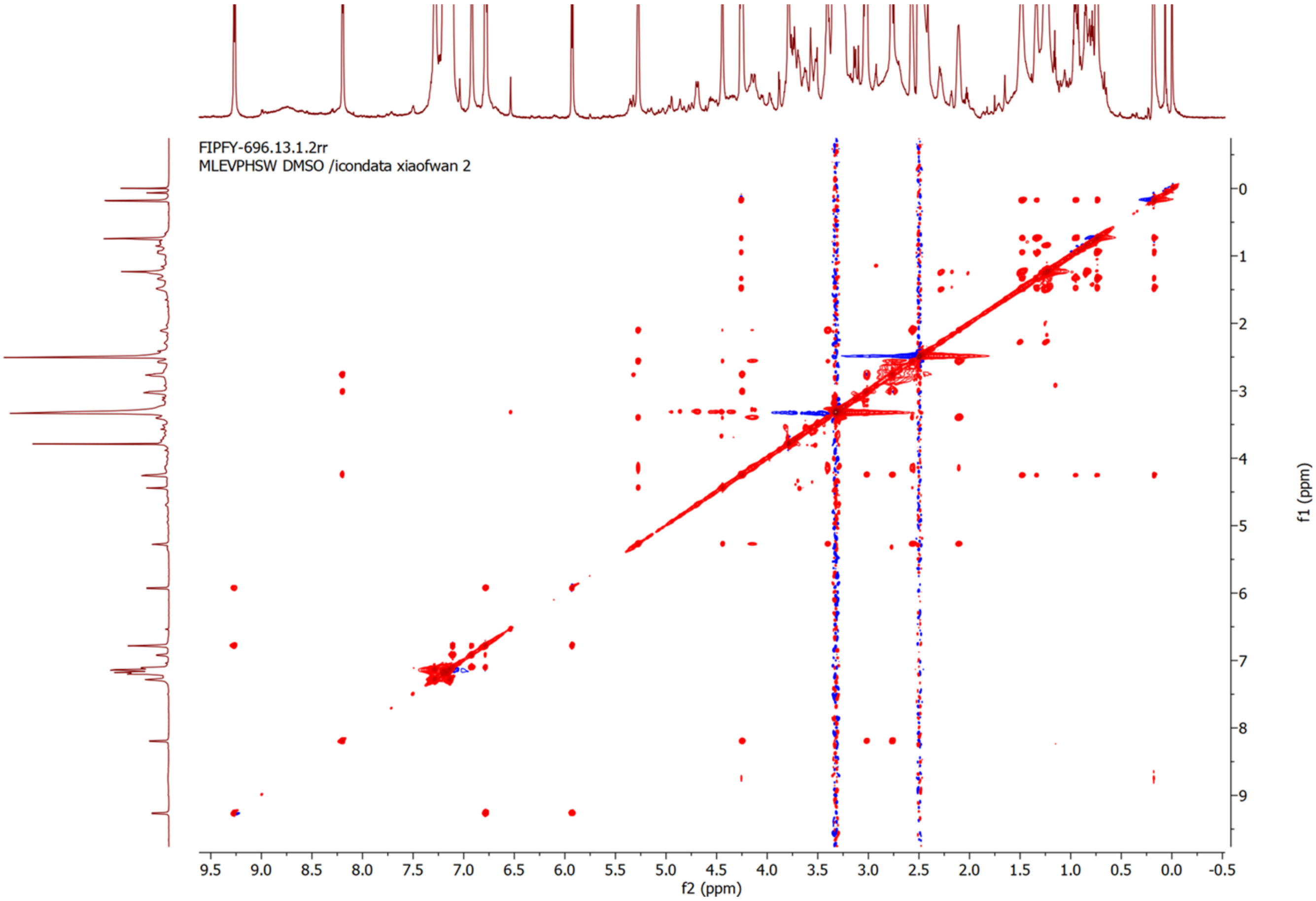


**Figure S15 | $^1$H-$^1$H TOCSY NMR spectrum of jubanine K in DMSO-$d_6$.**

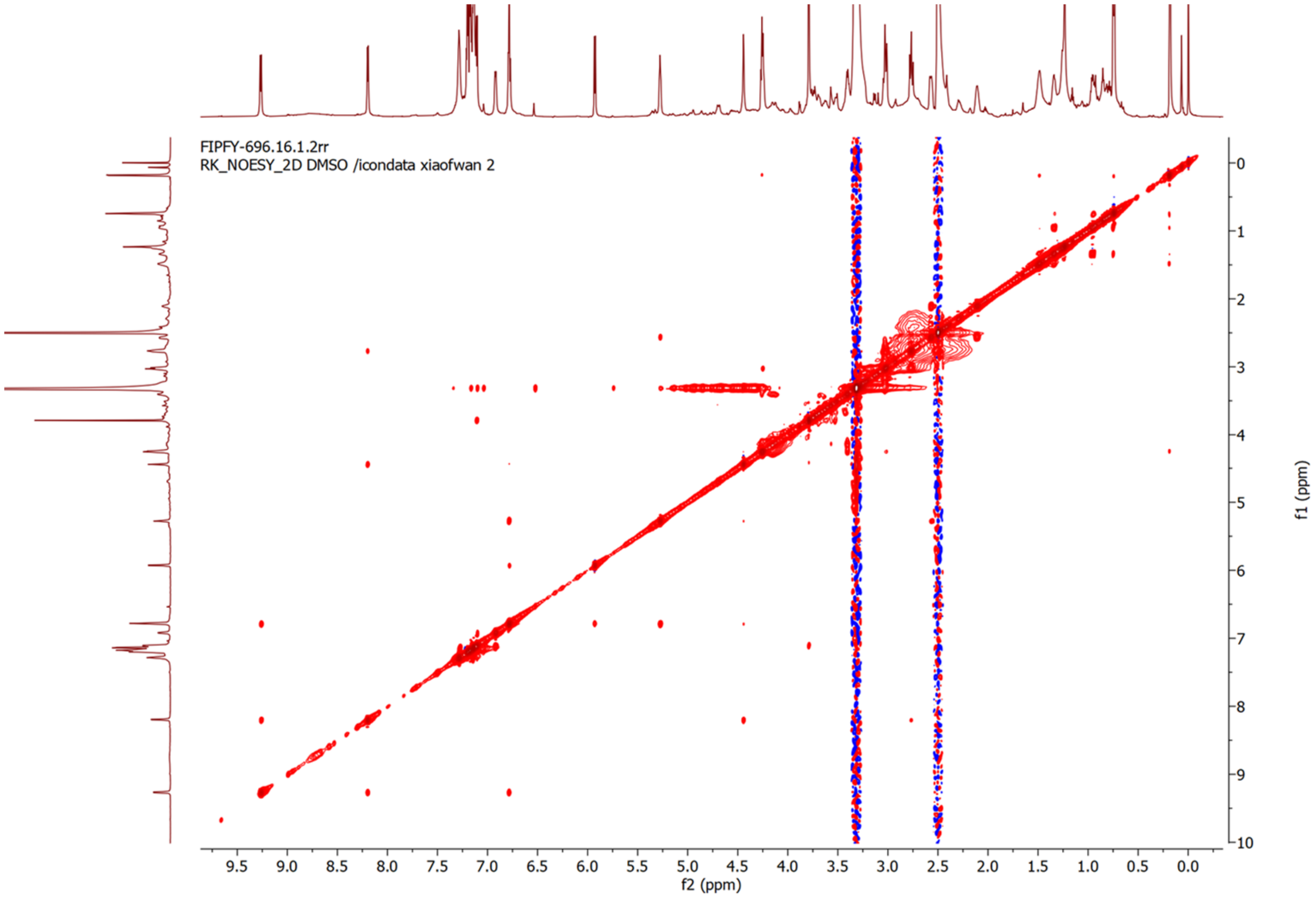


**Figure S16 | NOESY NMR spectrum of jubanine K in DMSO-$d_6$.**

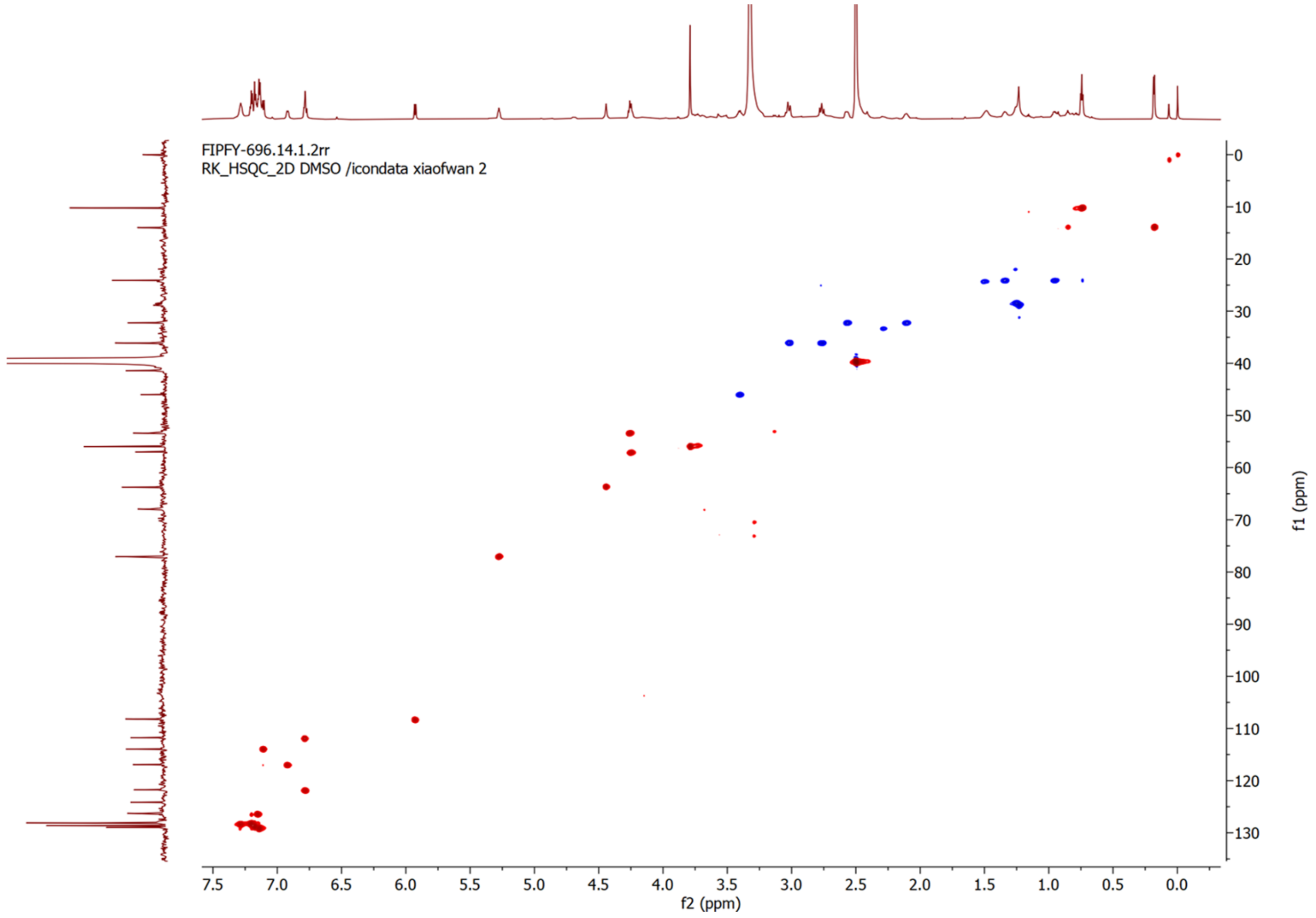


**Figure S17 | HSQC NMR spectrum of jubanine K in DMSO-$d_6$.**

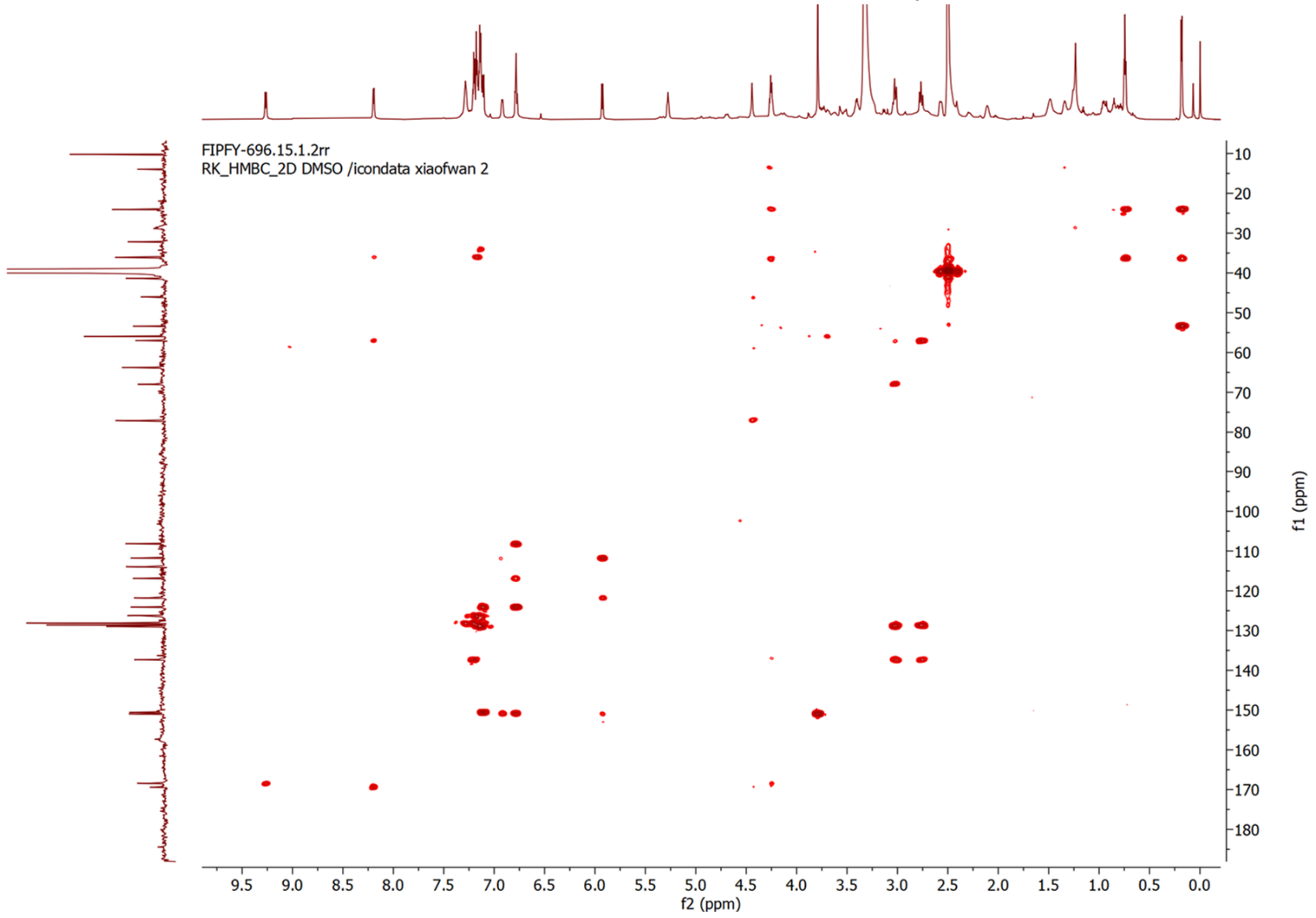


**Figure S18 | HMBC NMR spectrum of jubanine K in DMSO-$d_6$.**

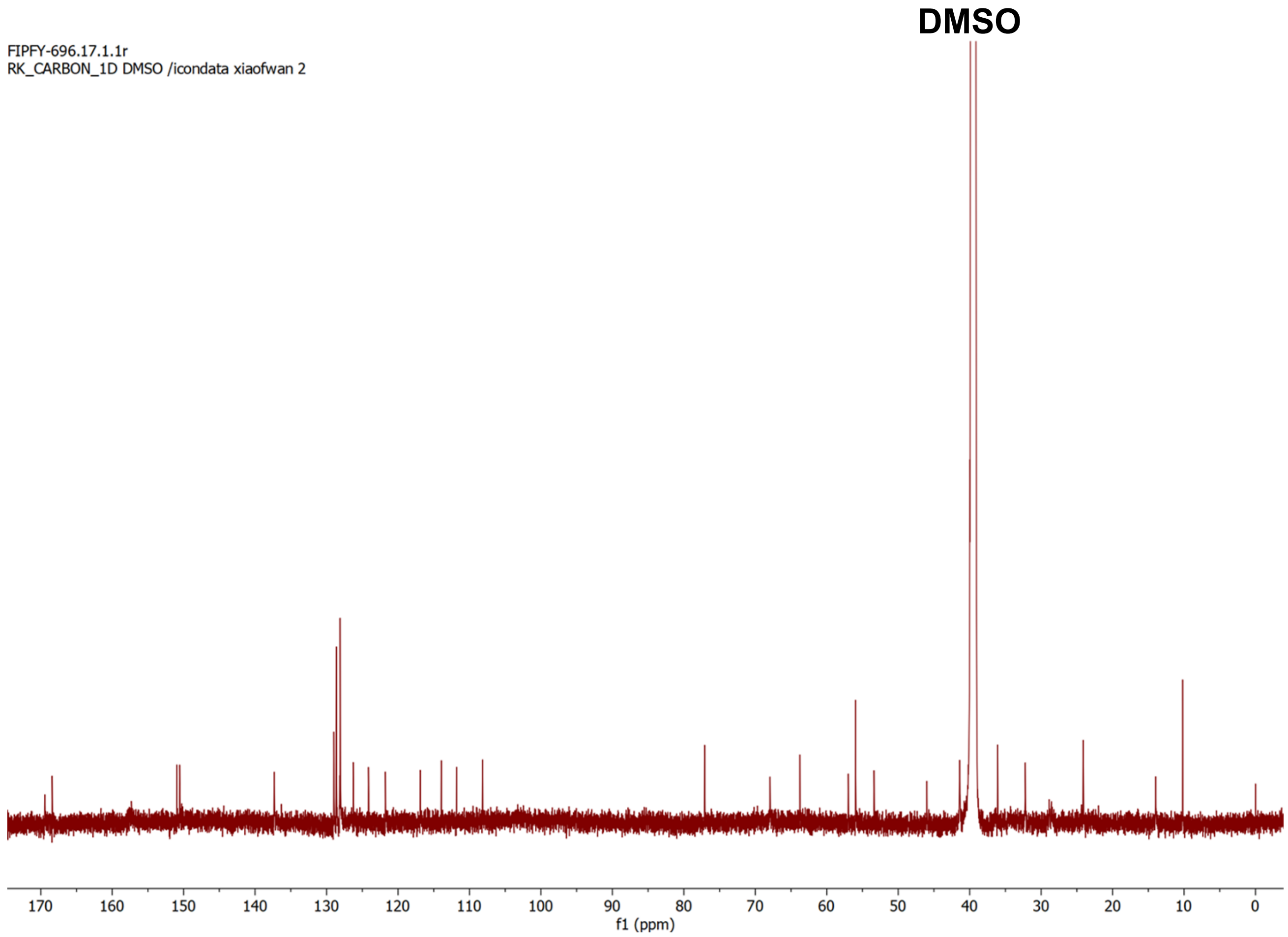


**Figure S19 | $^{13}$C NMR spectrum of jubanine K in DMSO-$d_6$ (200MHz).**

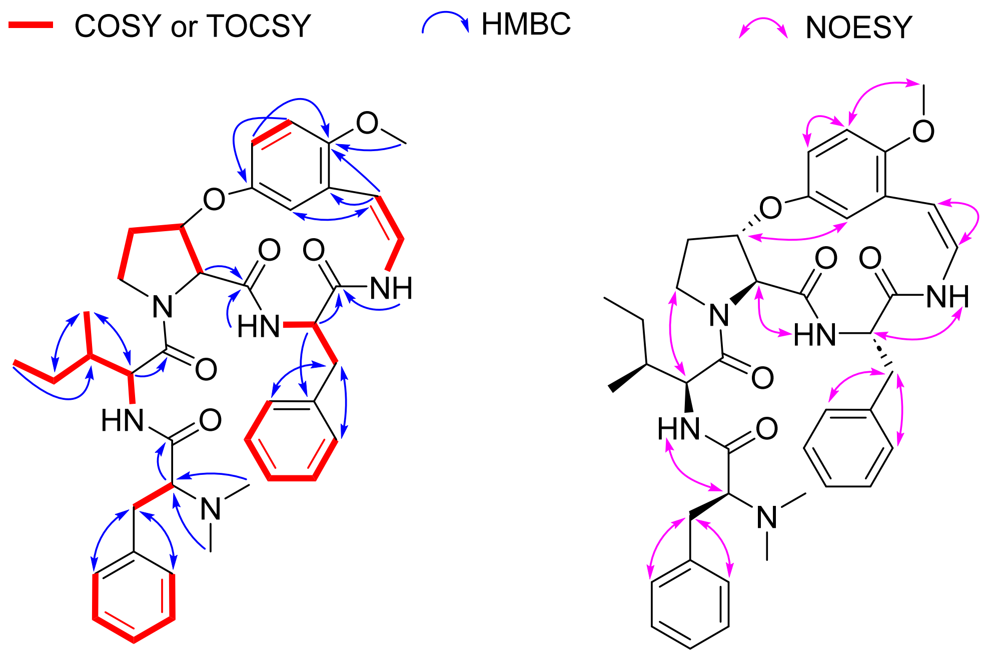


**Figure S20 | Key 2D NMR correlations of jubanine K in DMSO-$d_6$.**

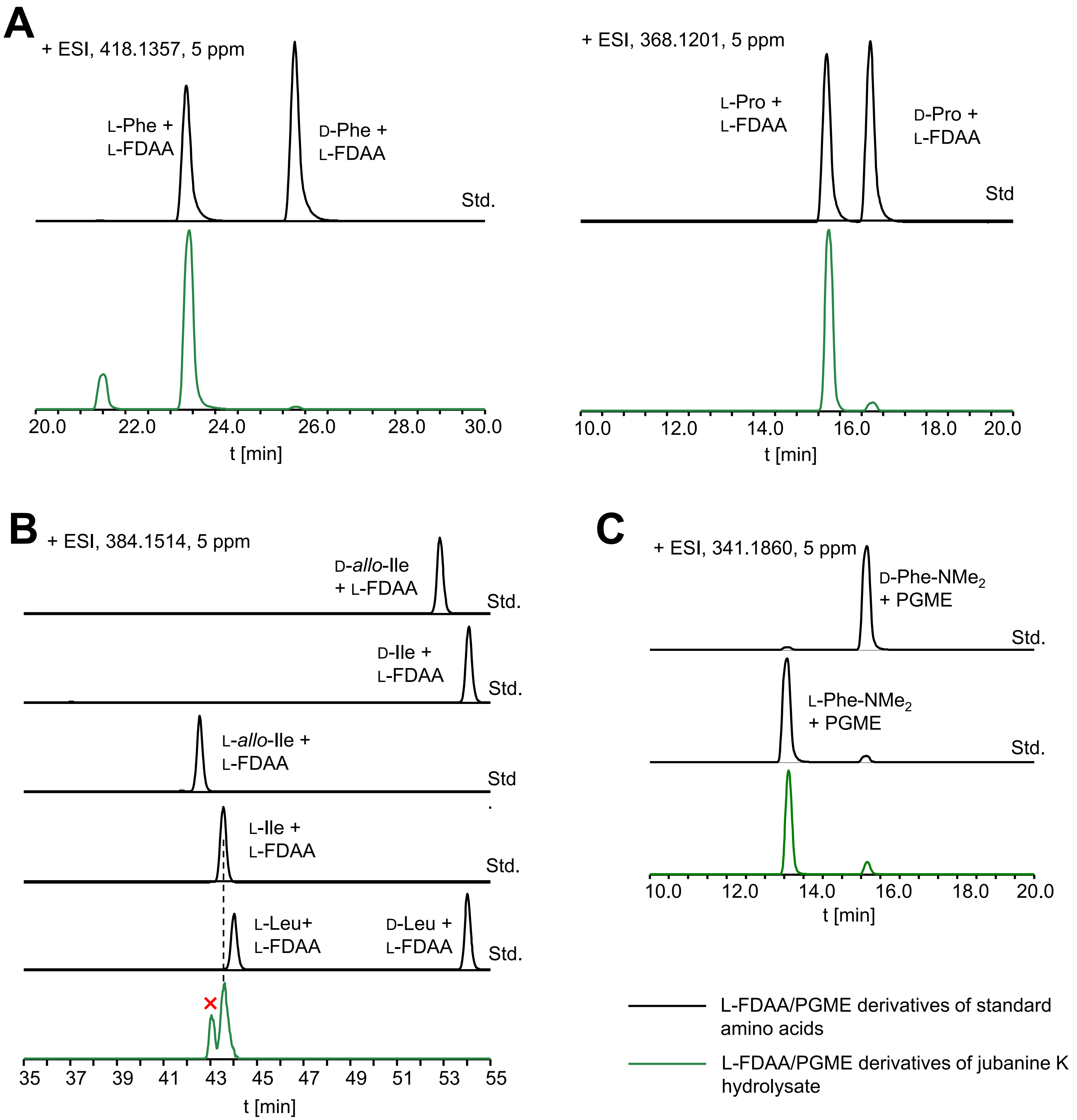


**Figure S21 | Stereochemical analysis of jubanine K.** (**A**) The Phe and Pro in jubanine K were assigned as L by Marfey's analysis. (**B**) The Ile in jubanine K was assigned as L by C3-Marfey's analysis. (**C**) The PheNMe$_2$ residue in jubanine K was assigned as L by PGME derivatization and LC-MS analysis.

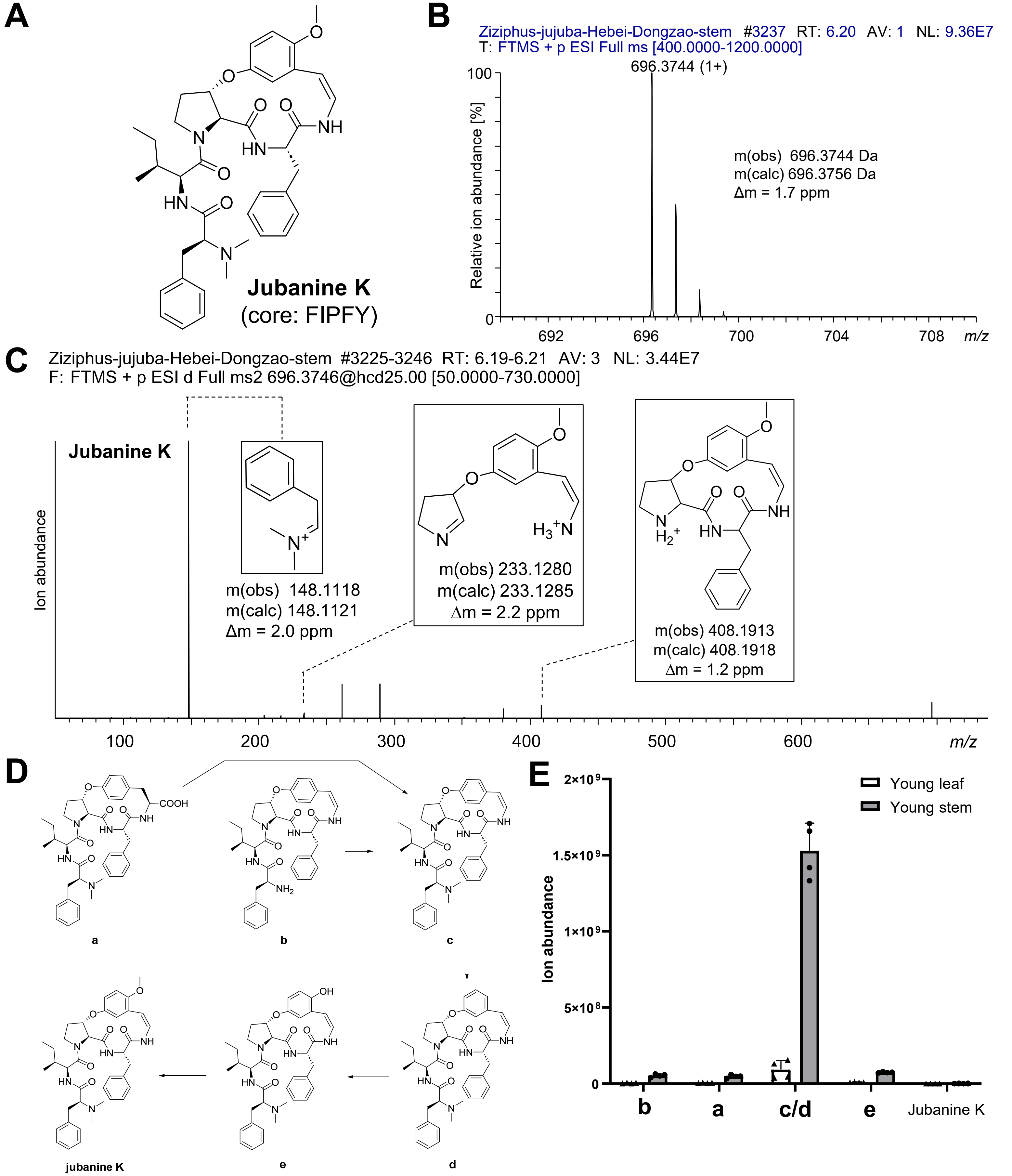


**Figure S22 | MS characterization of jubanine K and related metabolites in *Ziziphus jujuba* samples.** (**A**) Jubanine K structure. (**B**) MS spectrum of jubanine K from *Z. jujuba* var. Dongzao. (C) MS/MS spectrum of jubanine K and annotated predicted MS/MS fragments. (**D**) Predicted biosynthetic precursors of jubanine K. (**E**) Peak areas of predicted biosynthetic precursors of jubanine K. Bar graphs are means (*n*=4), error bars are one standard deviation.

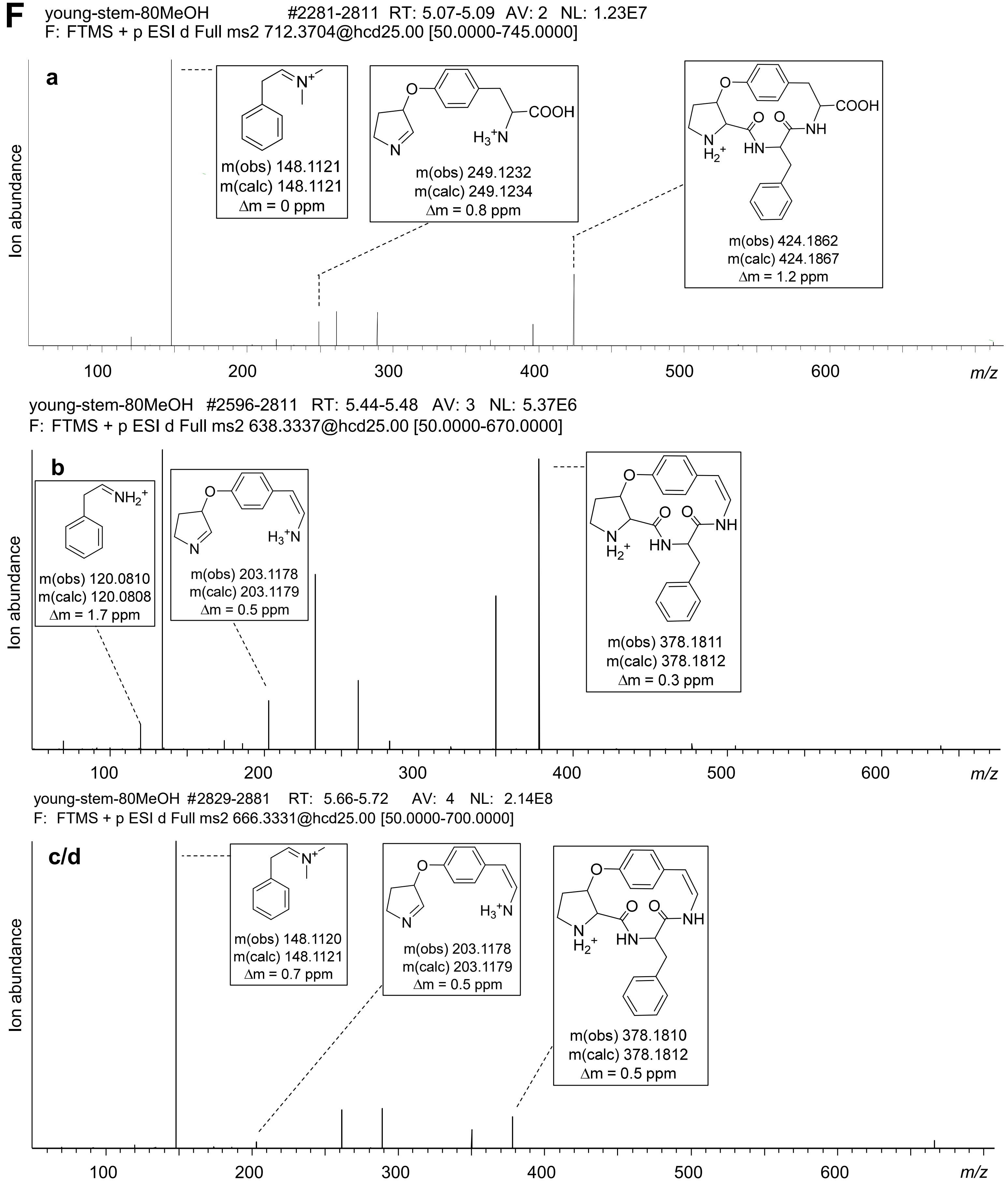


**Figure S22 | MS characterization of jubanine K and related metabolites in *Ziziphus jujuba* samples.** (**F**) MS/MS spectra of predicted biosynthetic precursors of jubanine K.

F

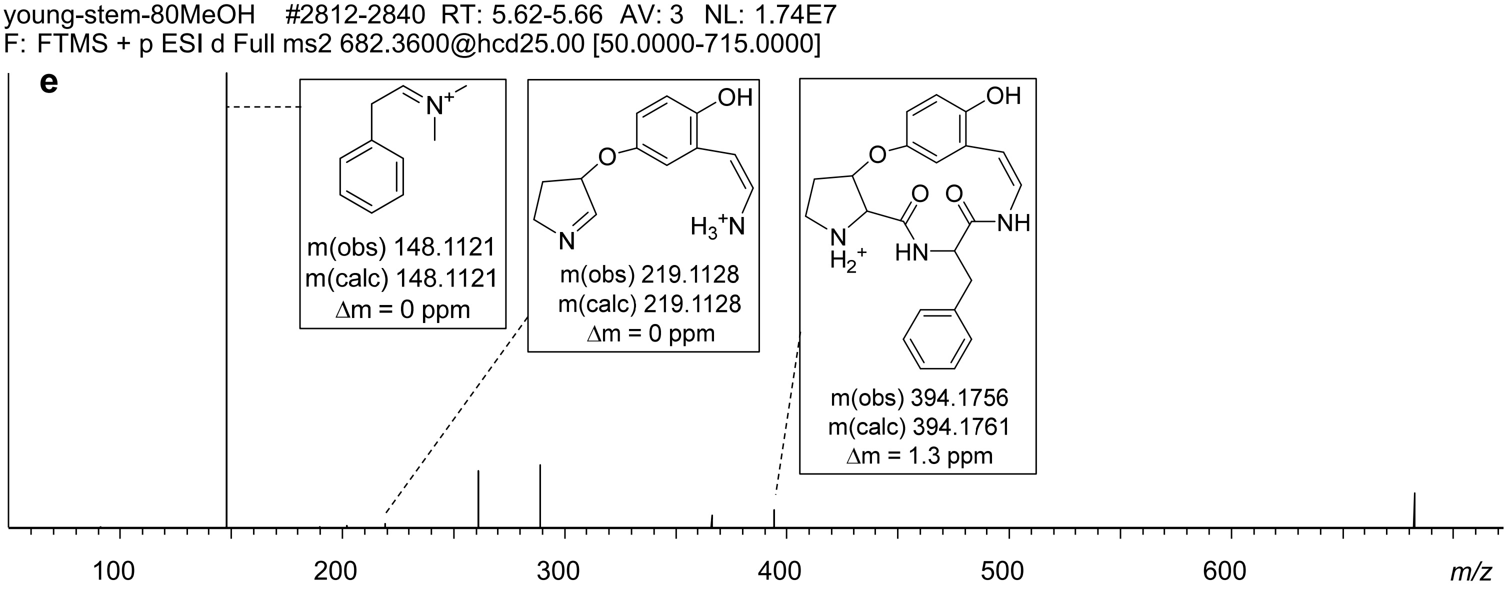


**Figure S22 | MS characterization of jubanine K and related metabolites in *Ziziphus jujuba* samples.** (**F**) MS/MS spectra of predicted biosynthetic precursors of jubanine K.

**A**

```
>ZjuPrec-FIPFY - XP_015880721.3 adhesin AWP1 [Ziziphus jujuba]
MREPLHHSSFSKFQFVIKMKSFFALLAFSSLLLLASTITARKEPAEYVKTTLEQVAKDLSIDPSSFVPFY
HRKQNDISVDPSSFIPFYHGKPNGLSVDPSSFIPFYHGKQNDLSVDPSSFVPFYHGKPNGLSVDPSSFVP
FYHGKPNDLSVDPSSFVPFYHGKPNDLSVDPSSFVPFYRGNKNGNSVDPSSFIPFYHGKANRLSVDPSSL
VPFYHGKPNGLSIDPSSFVPLYHGNRNGFSVDPSSFVPFYHGNRNDLSVDPSSFIPLYQGNQNGLSVDPS
SFIPFYHGNENSDQNTKINEEGVAKVVSPDHQ
```

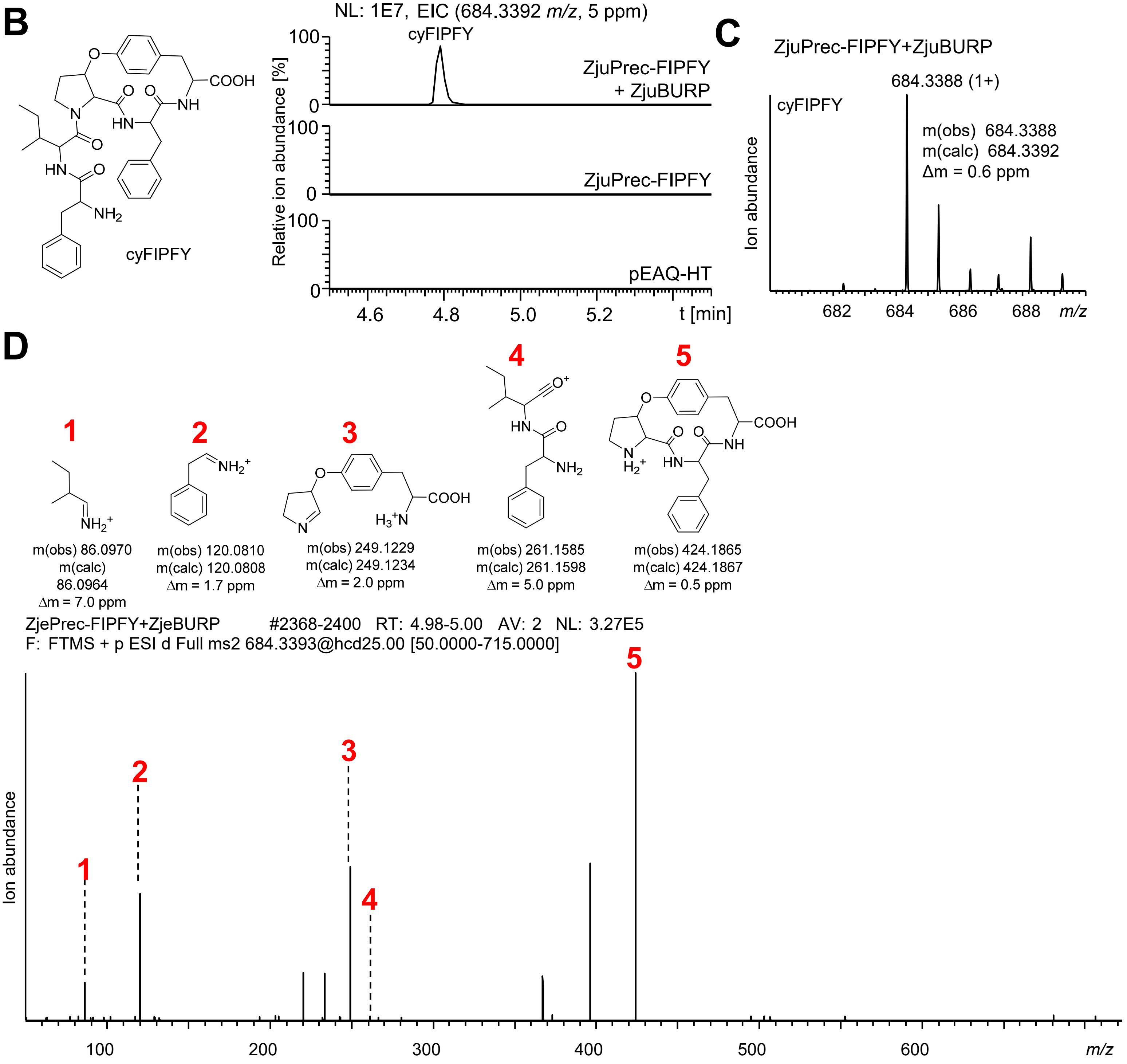


**Figure S23 | Characterization of candidate precursor peptide of jubanine K from *Ziziphus jujuba* var. Dongzao.** (**A**) Protein sequence of candidate precursor peptide ZjuPrec-FIPFY of jubanine K. Jubanine K matching core peptide FIPFY is highlighted in red. (**B**) Transient expression of ZjuPrec-FIPFY with or without ZjuBURP in *Nicotiana benthamiana* via pEAQ-HT. EIC traces for monocyclic FIPFY analyte mass $[M+H]^+$ are shown from methanolic extracts of *N. benthamiana* leaves 7 days after agroinfiltration. A predicted 2D cyclic FIPFY structure is shown. (**C**) Mass spectrum of monocyclic FIPFY analyte mass $[M+H]^+$. (**D**) Tandem mass spectrum of monocyclic FIPFY analyte mass $[M+H]^+$.

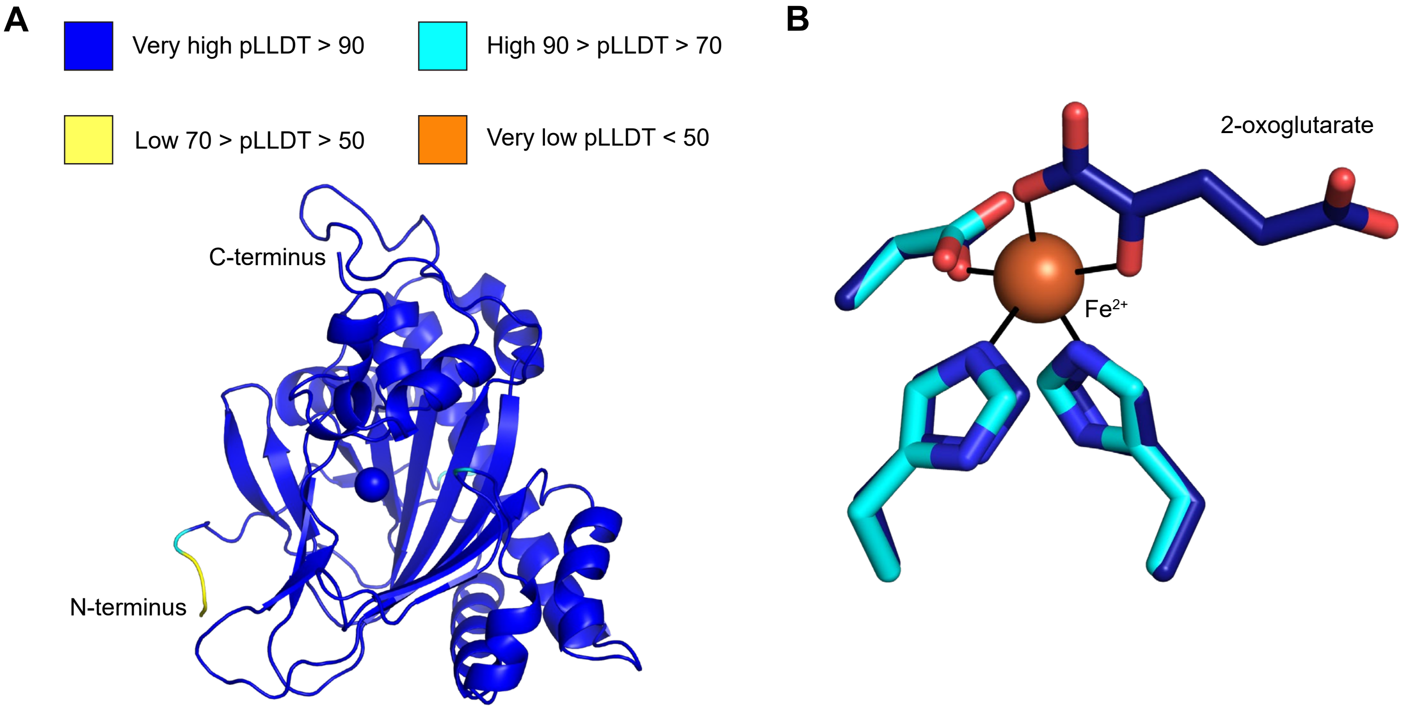


C

```
Zju-Decarboxylase                        LDKMVRRMVLESLGVEKY-FEEHMETTKCLLRVMKFKGPHIKDNKKLGLVPHTDKEFITI   197
8CI9-Deoxypodophyllotoxin-Synthase       VDHILKMMVFESFGMEKH-FDSFCESTNYLLHFMRYQQPGK-DGRSPALSLHKDKSILTI   192
6KUN-Dioxygenase-Auxin-Oxidation         LIVDVAGKVAASLGLHGA----SFQDWPCQFRMNRYNYTQD-SVGSPGVQVHTDSGFLTV   181
8CV8-Hyoscyamine-6-beta-Hydroxylase      LTMRILDYICEGLGLKLGYFDNELSQ--IQMMLTNYYPPCPDPSSTLGSGGHYDGNLITL   225
                                         :   :   :  .:*:.        .     : . .:         . .   * *  ::*:

Zju-Decarboxylase                        LYHNH-VSGLEVQIKDGTWISVERNSPDSFIVMFGDSFYAWTNGRCHSPRHQVMMTGDET   256
8CI9-Deoxypodophyllotoxin-Synthase       VNQND-VKGLEFETKDGEWILPT---ADNHIVLLGDCFMAWSNGRLHSPLHRVTLVANQA   248
6KUN-Dioxygenase-Auxin-Oxidation         LQEDECVGGLEVLDPAAGEFVPVDPLPGSFVVNVGDVGQAWSNGRLHNVKHRVQCVAAVP   241
8CV8-Hyoscyamine-6-beta-Hydroxylase      LQQN--LPGLQQLIEDAKWIAV-EPIPTAFVVNLGLTLKVITNEKFEGSIHRVVTNPTRD   282
                                         : .:  : **:     .  :         .:* .*    . :* : ..  *:*
```

**Figure S24 | Protein model prediction of ZjuDC and alignments with experimentally determined plant non-heme Fe(II)- and 2-oxoglutarate dependent enzymes.** (**A**) AlphaFold3 prediction[1] of ZjuDC (XP_048320867.1) colored based on pLLDT values, where Fe(III) is a sphere. (**B**) Conserved residues of non-heme-iron and 2-oxoglutarate-dependent enzyme sequence motif H-X-(D/E)-$X_n$-H[2]. ZjuDC model is cyan and hyoscyamine-6 β-hydroxylase [*Atropa belladonna*], PDB 8CV8[3] is dark blue. Sticks are colored according to atomic type, oxygen red, nitrogen blue, and iron sphere from PDB 8CV8 is bronze. Coordination bonds are thin black lines, and other ligands are omitted for clarity. (**C**) Excerpt of Clustal Omega 1.2.4[4] sequence alignment of top results and varying catalytic activity from ZjuDC model in FoldSeek[5]. ZjuDC sequence is compared to experimentally determined structures PDB 8CI9[6] deoxypodophyllotoxin synthase [*Sinopodophyllum hexandrum*]; PDB 6KUN[7] auxin dioxygenase [*Oryza sativa Indica Group*]; hyoscyamine-6 β-hydroxylase [*Atropa belladonna*], PDB 8CV8[3]. All experimental structures align to the ZjuDC model with Cα root-mean-squared deviation <1.8 Å across 192-253 atoms and <50 % sequence identity. The conserved sequence motif residues are highlighted in cyan rectangles.

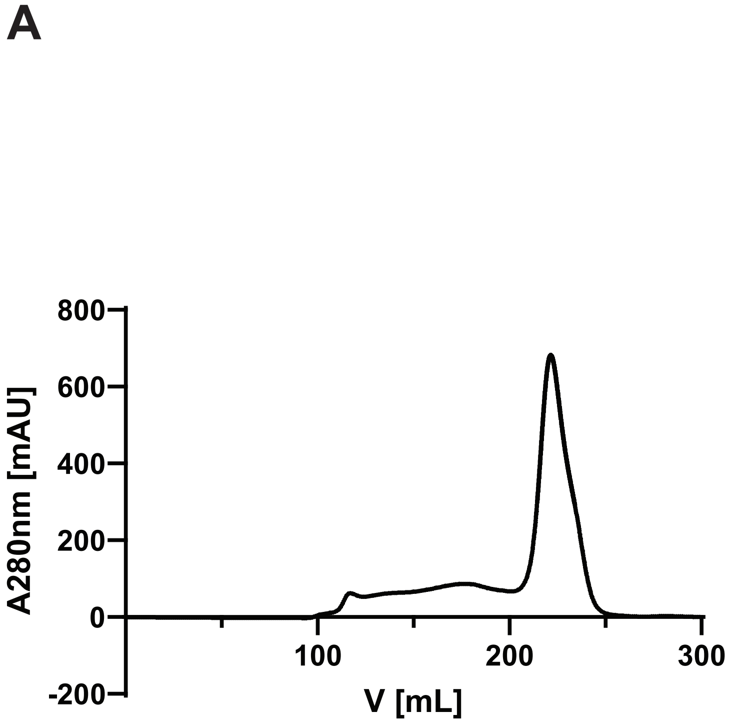


| Construct | Rentention [mL] | M.W. [Da] |
|---|---|---|
| ZjuDC | 221 | 35329 |

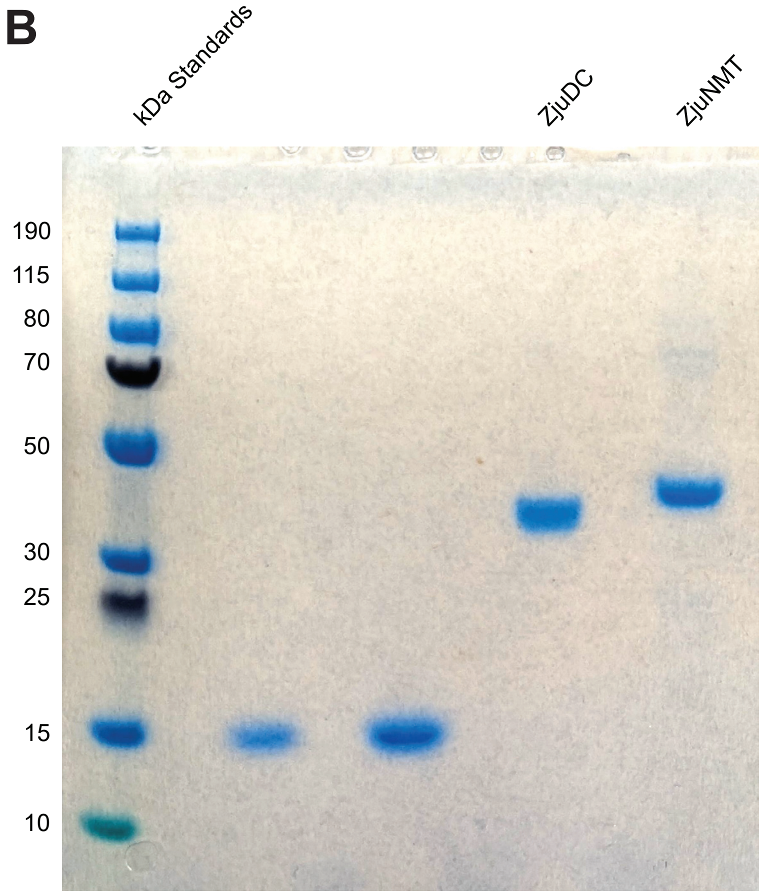


**Figure S25 | SDS-PAGE of heterologously expressed and purified proteins.** (**A**) Size exclusion chromatogram of ZjuDC with protein elution profile based on A280nm (mAu) and calculation of molecular weight estimate based on SEC elution volume. (**B**) SDS-PAGE of purified ZjuDC and ZjuNMT. All protein purifications were reproducible a minimum of three times.

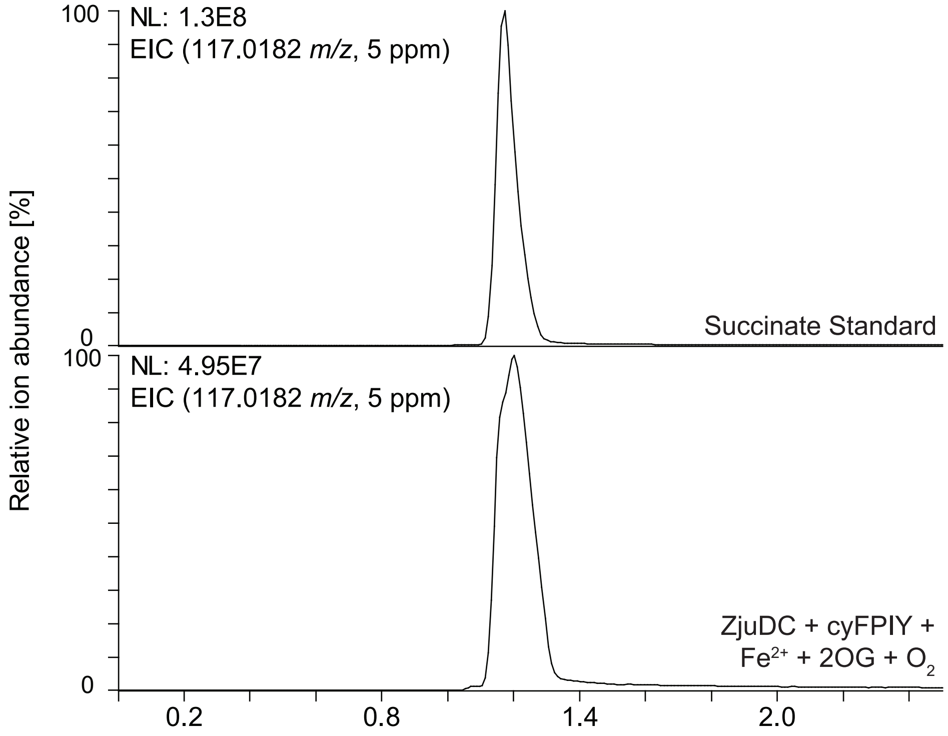


**Figure S26 |** Succinate detection in ZjuDC *in vitro* enzyme assay (bottom trace) compared to a succinate authentic standard (top trace).

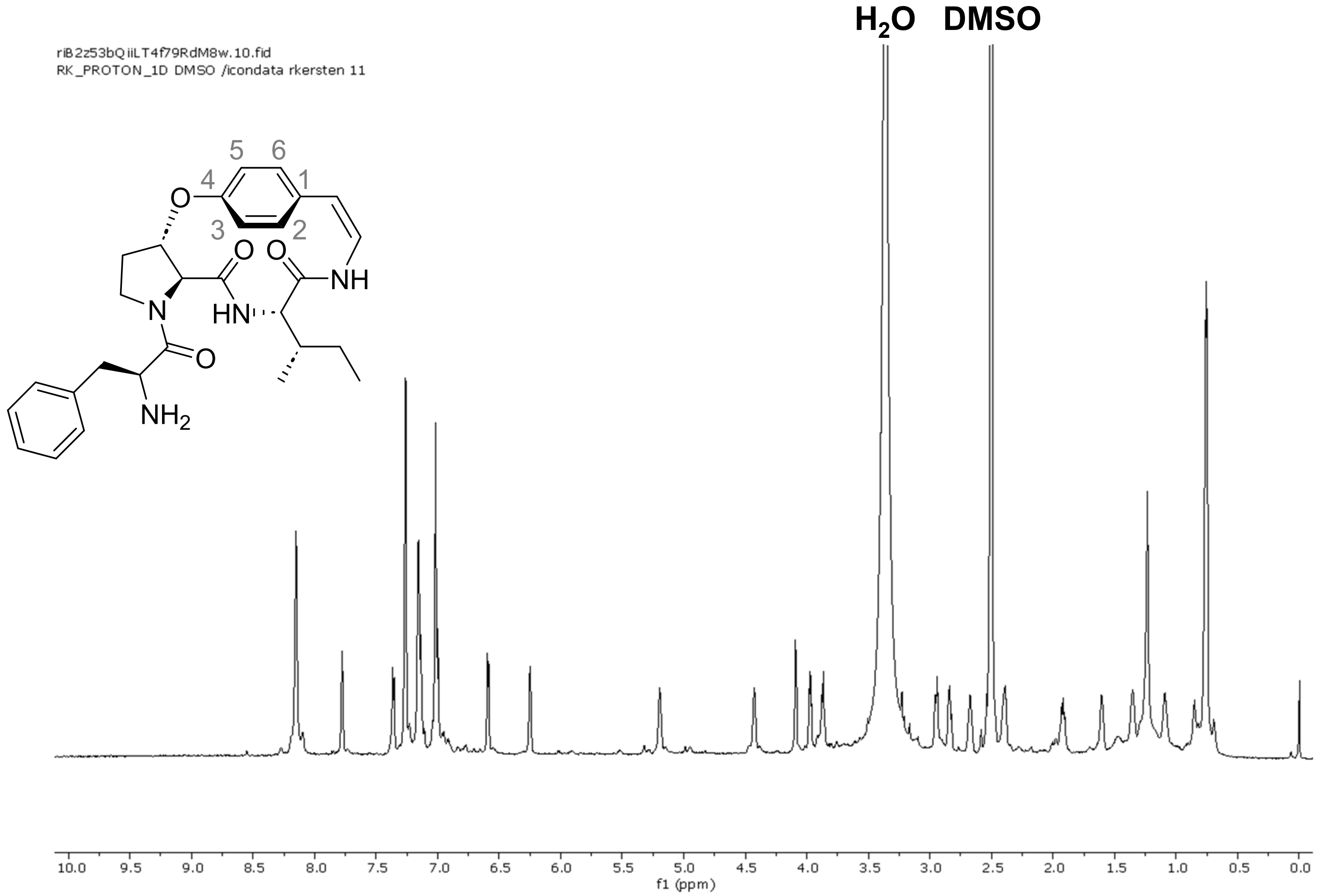


**Figure S27 | $^{1}$H NMR spectrum of cyFPIY-dc in DMSO-$d_6$ (800 MHz).**

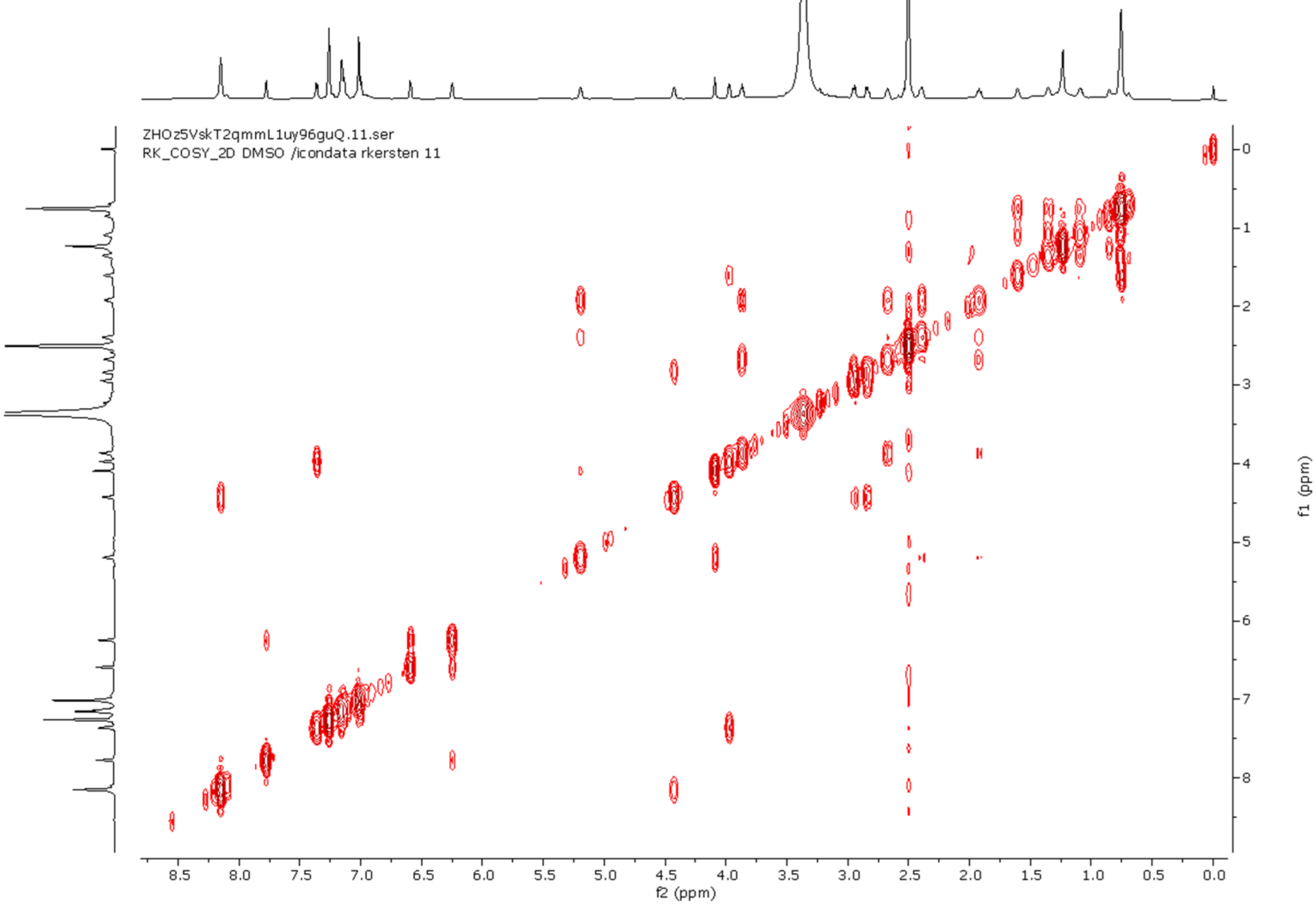


**Figure S28 | $^{1}$H-$^{1}$H COSY NMR spectrum of cyFPIY-dc in DMSO-$d_6$.**

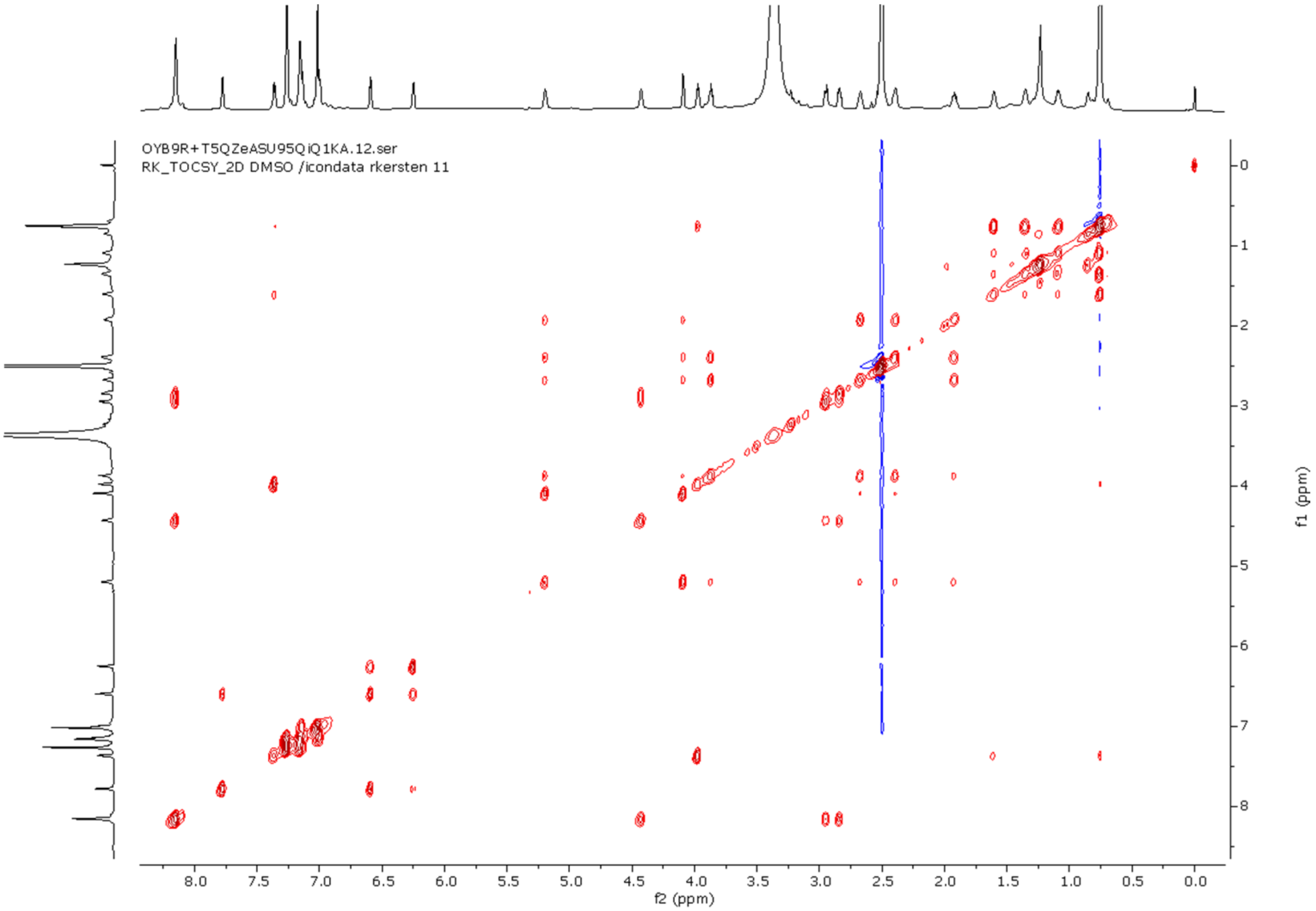


**Figure S29 | $^1H$-$^1H$ TOCSY NMR spectrum of cyFPIY-dc in DMSO-$d_6$.**

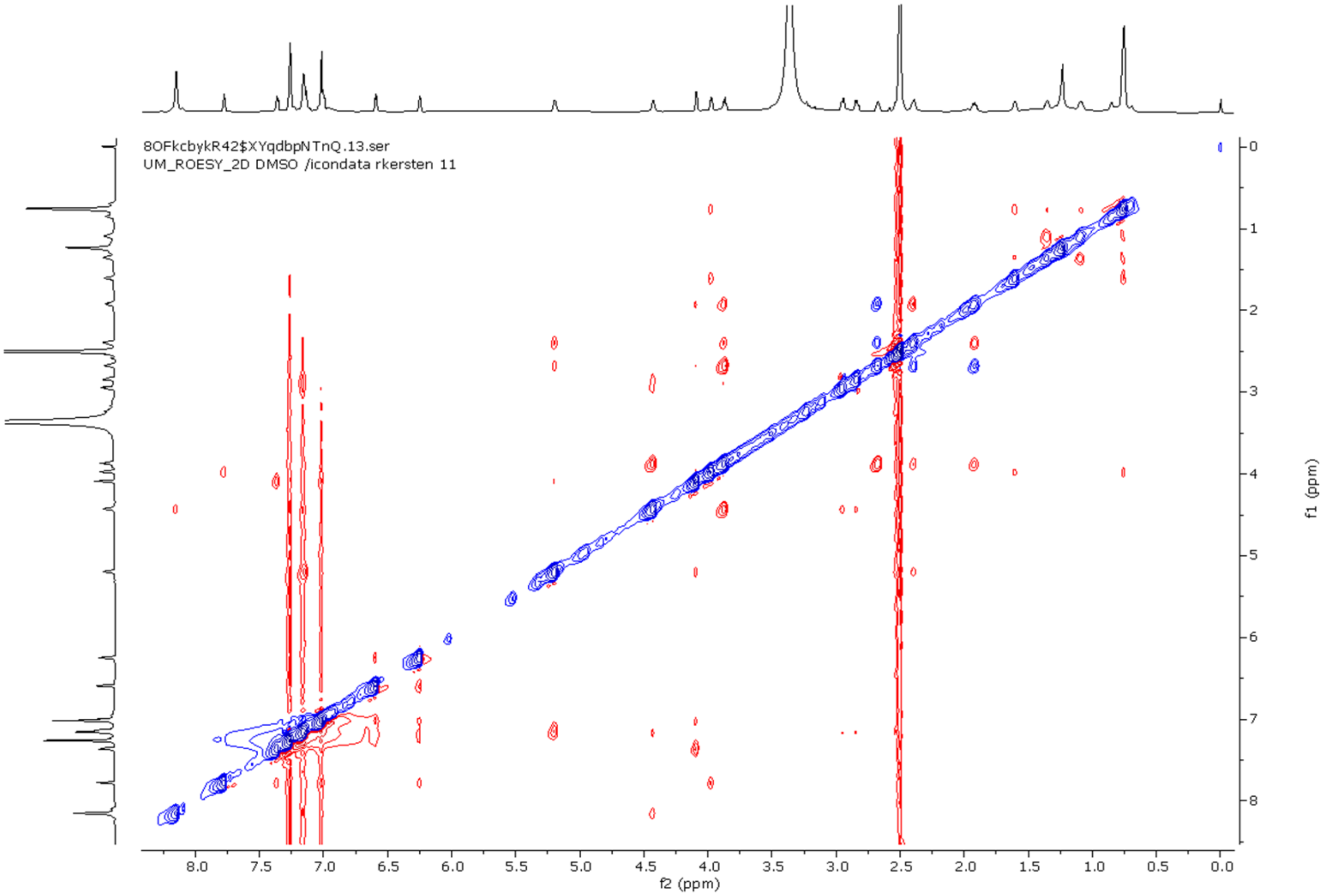


**Figure S30 | ROESY NMR spectrum of cyFPIY-dc in DMSO-$d_6$.**

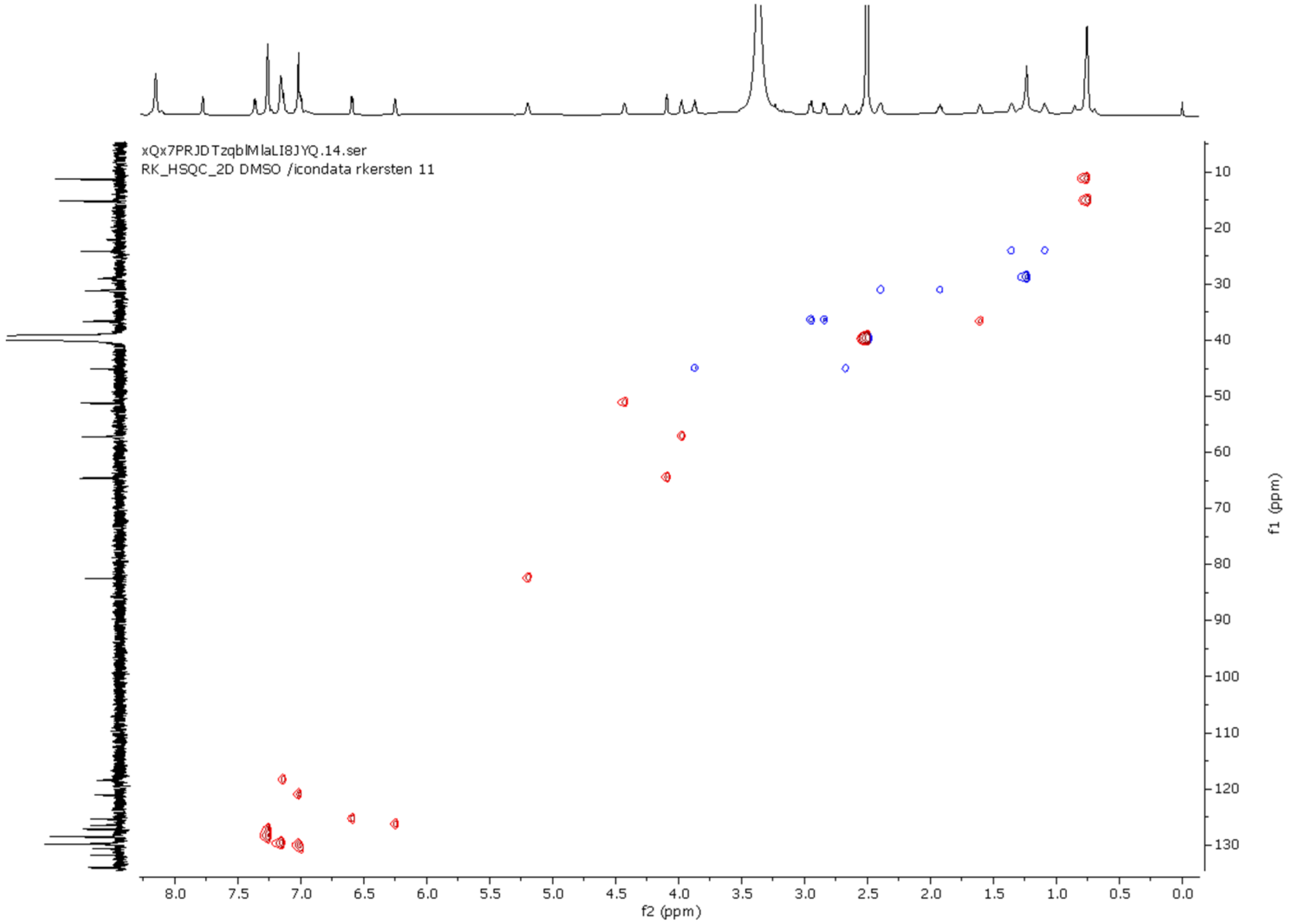


**Figure S31 | HSQC NMR spectrum of cyFPIY-dc in DMSO-d$_6$**

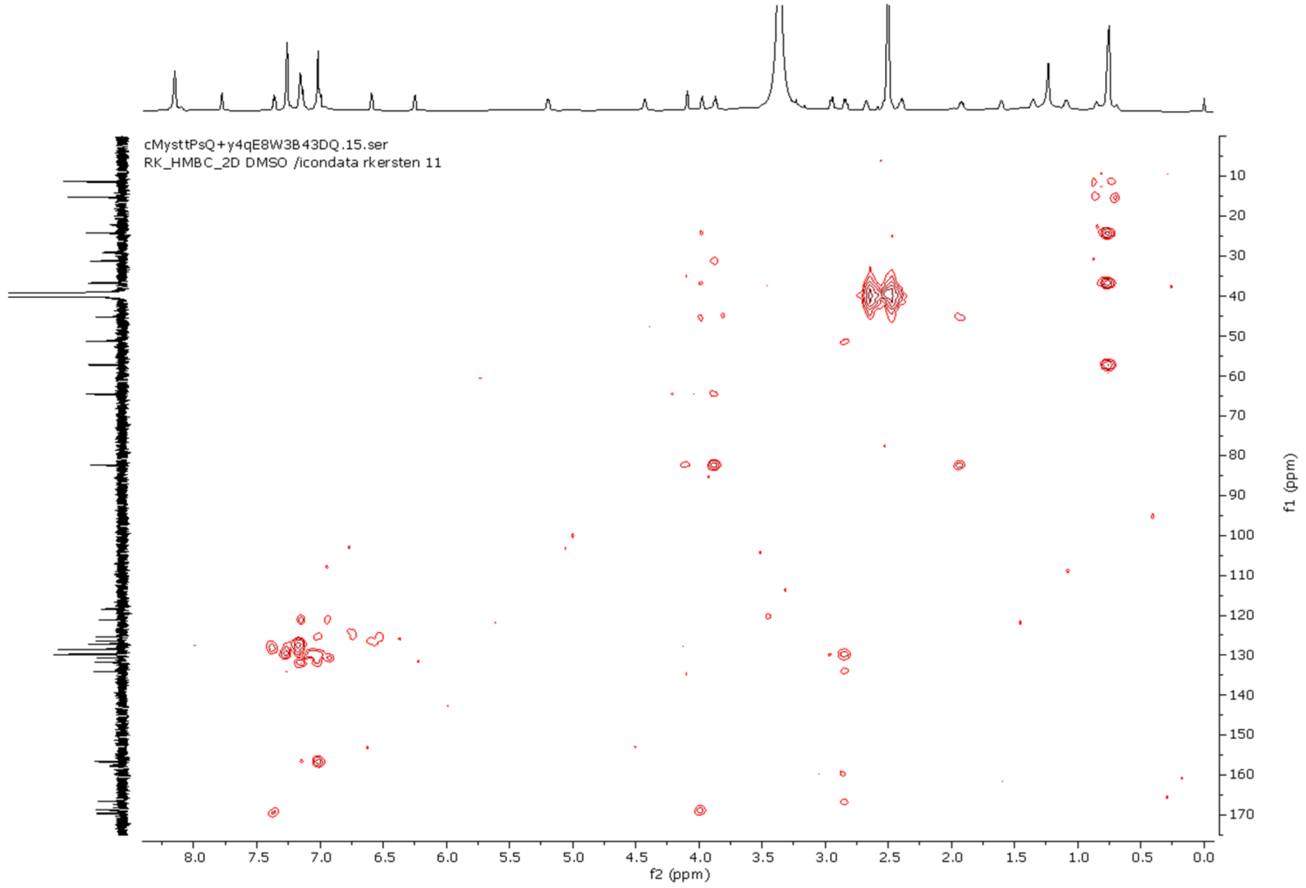


**Figure S32 | HMBC NMR spectrum of cyFPIY-DC in DMSO-d$_6$.**

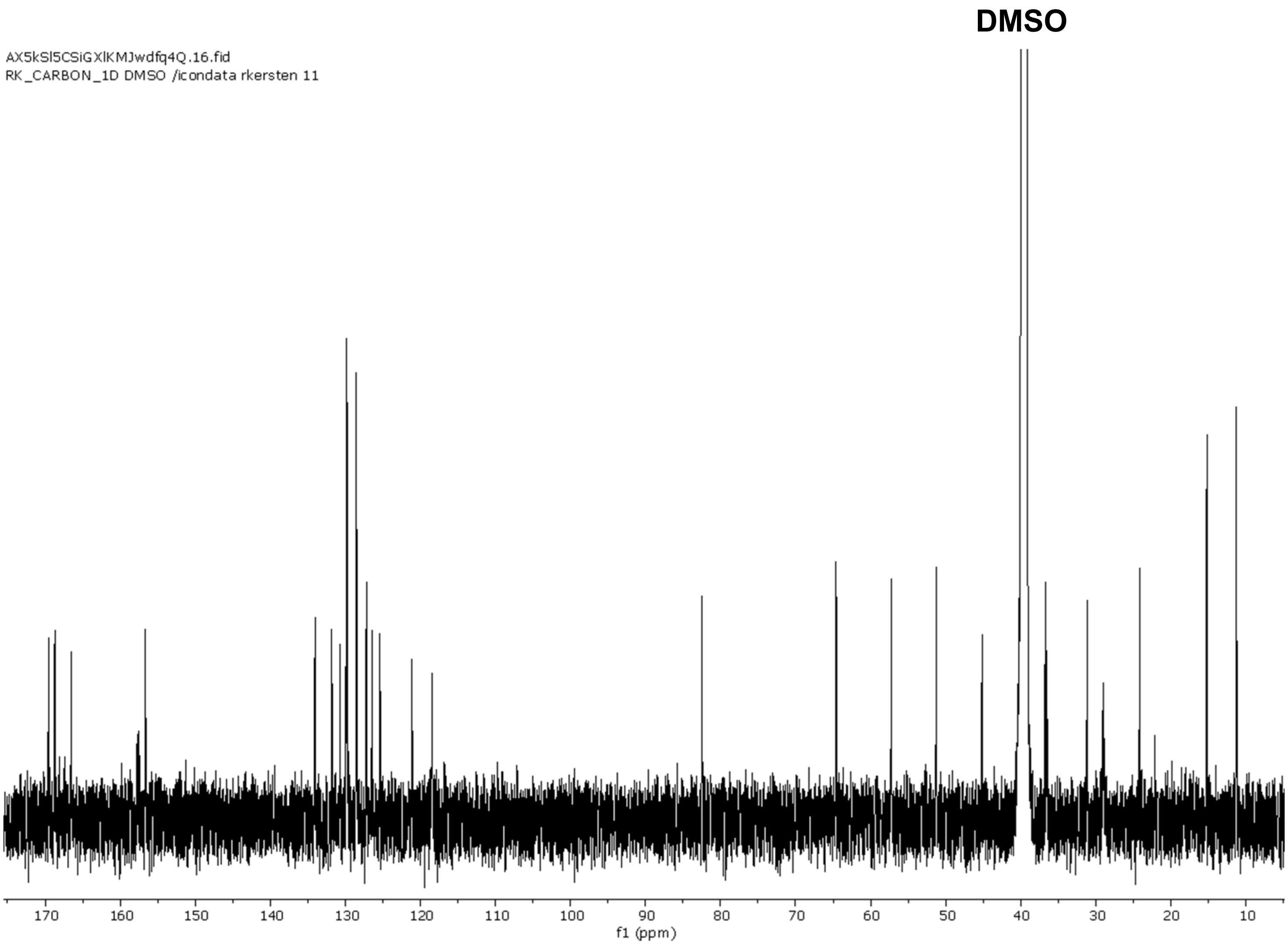


**Figure S33 | $^{13}$C NMR spectrum of cyFPIY-dc in DMSO-$d_6$ (200 MHz).**

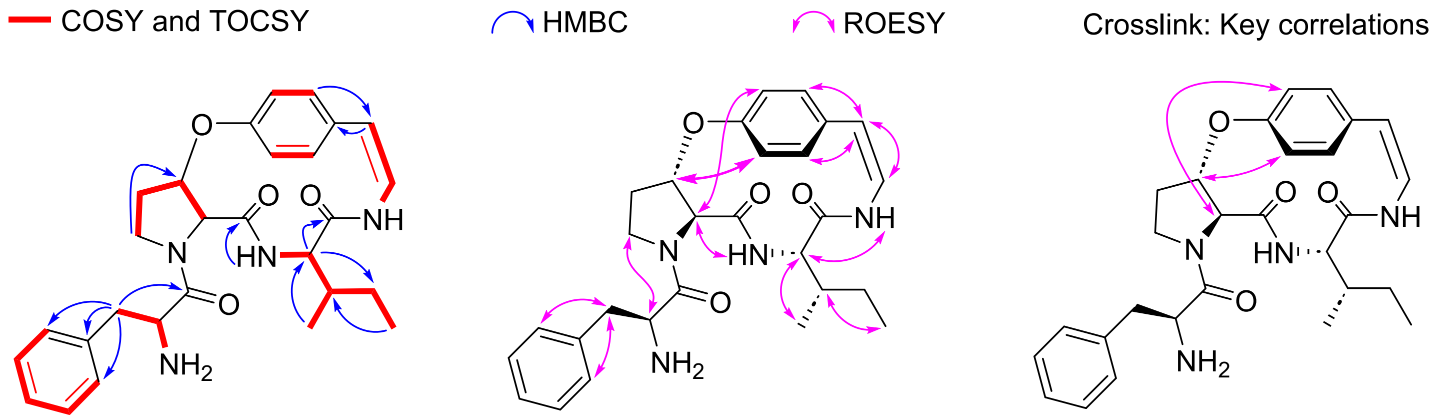


**Figure S34 | Key 2D NMR correlations of cyFPIY-dc in DMSO-$d_6$.**

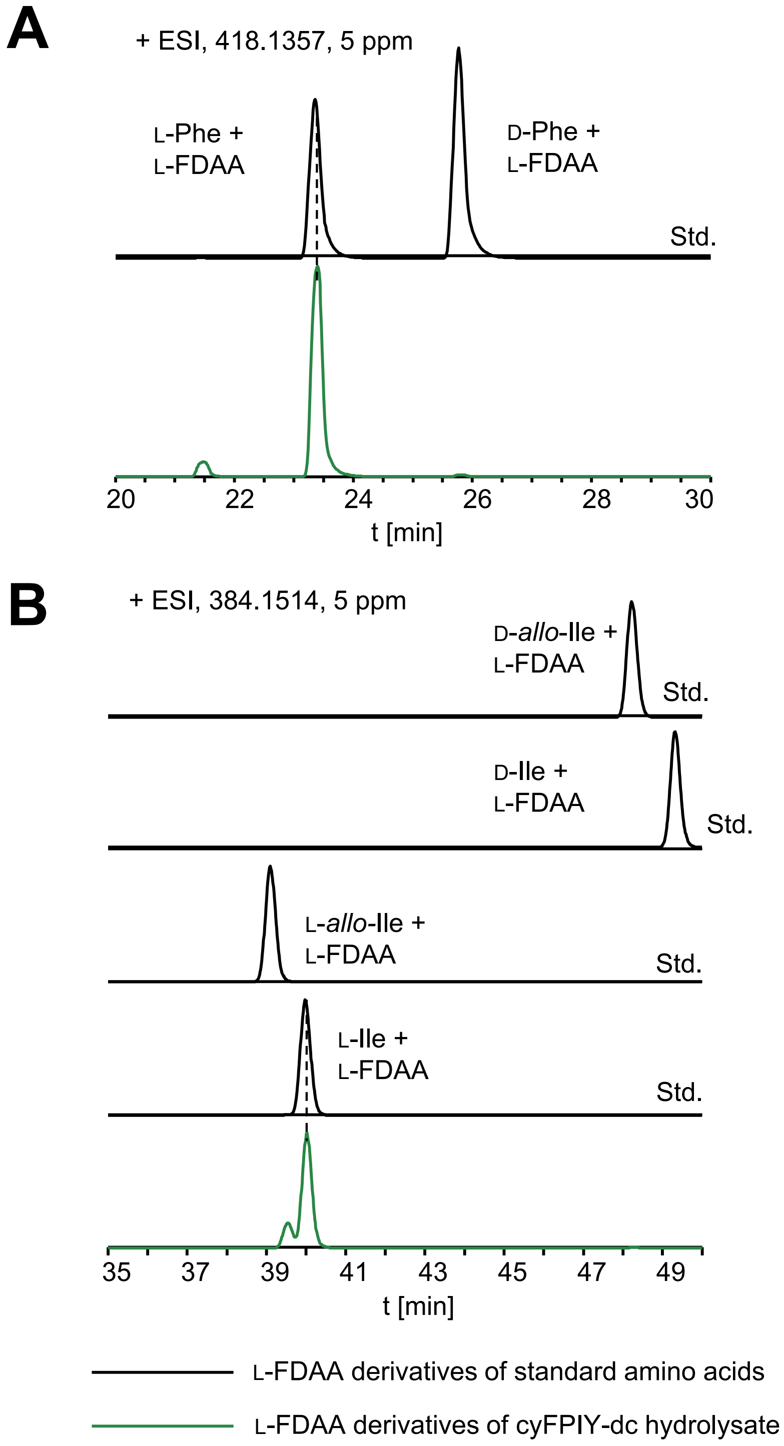


**Figure S35 | Stereochemical analysis of cyFPIY-dc.** (**A**) The Phe and Pro residues in cyFPIY were assigned as L by using Marfey's analysis. (**B**) The Ile residue in cyFPIY-dc was assigned as L by using C3-Marfey's analysis

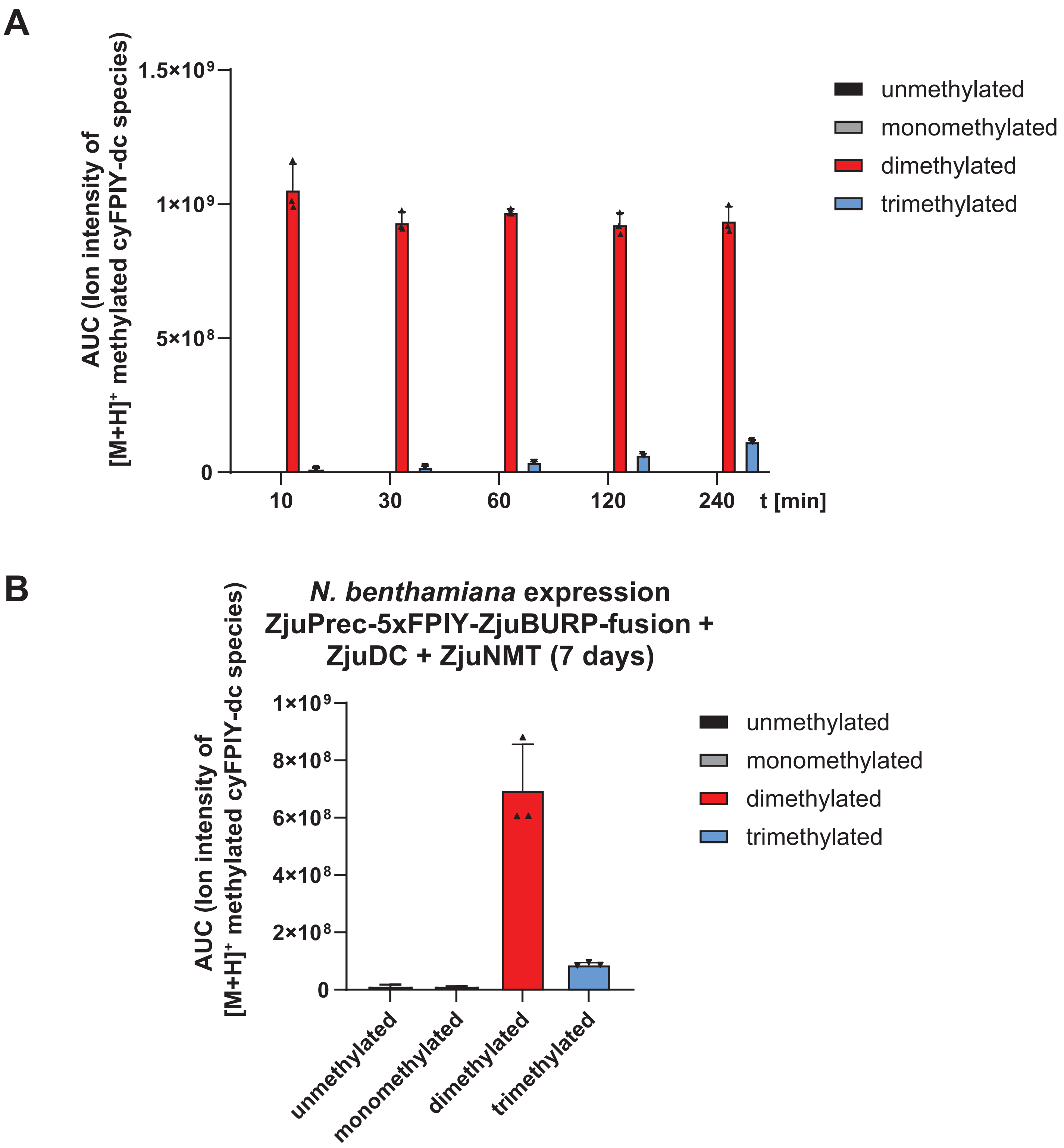


**Figure S36 | ZjuNMT product methylation *in vitro* and *in planta*.** (**A**) Time course of ZjuNMT *in vitro* reaction with cyFPIY substrate. Means (*n*=3) of methylated cyFPIY product mass signal AUC are displayed with error bars displaying one standard deviation. (**B**) ZjuNMT *in planta* reaction products in transgenic *N. benthamiana* 7 days after agro-infiltration with ZjuPrec-5xFPIY-ZjuBURP-fusion, ZjuDC and ZjuNMT. Means (*n*=3) of methylated cyFPIY-dc product mass signal AUC are displayed with error bar displaying one standard deviation.

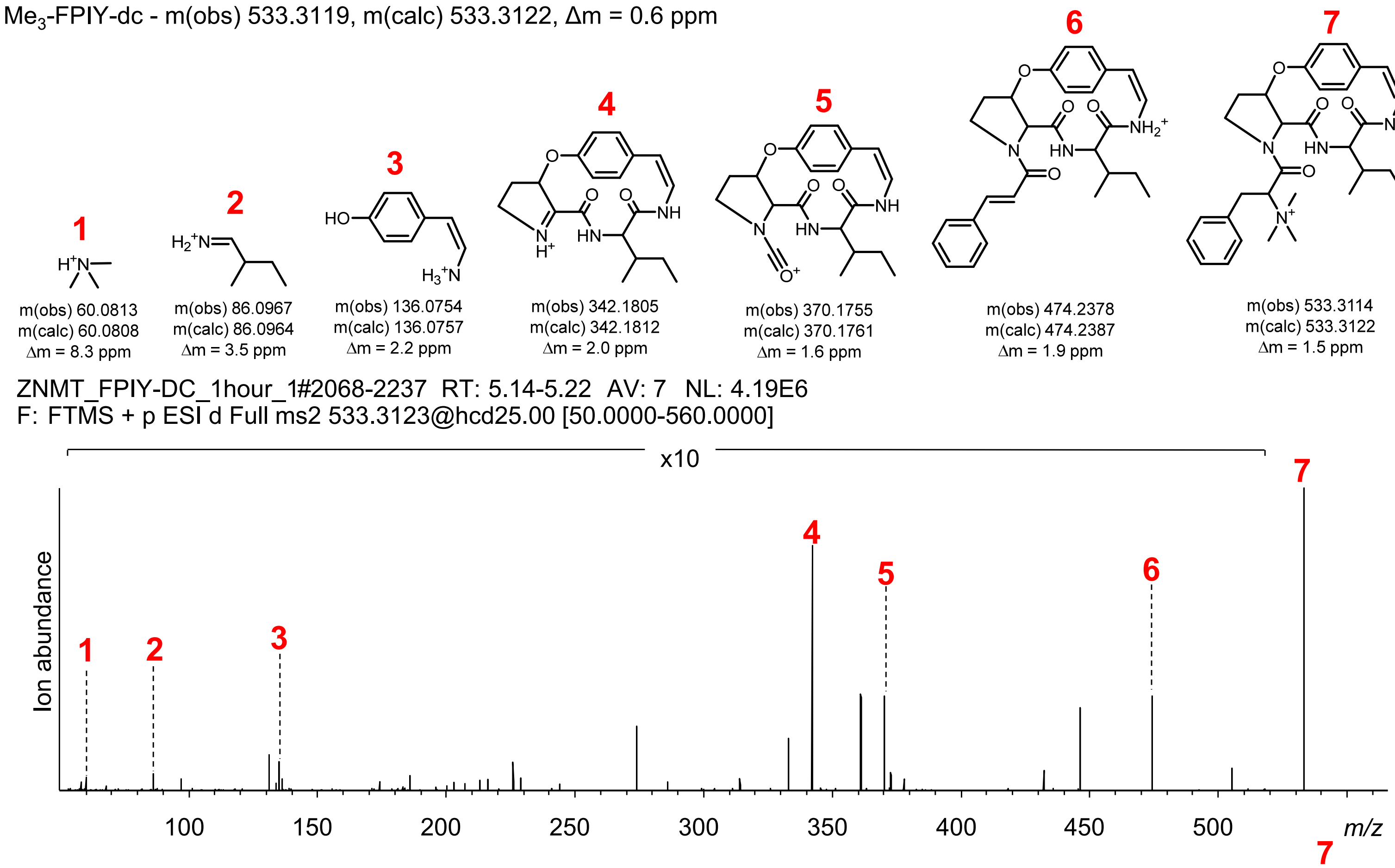


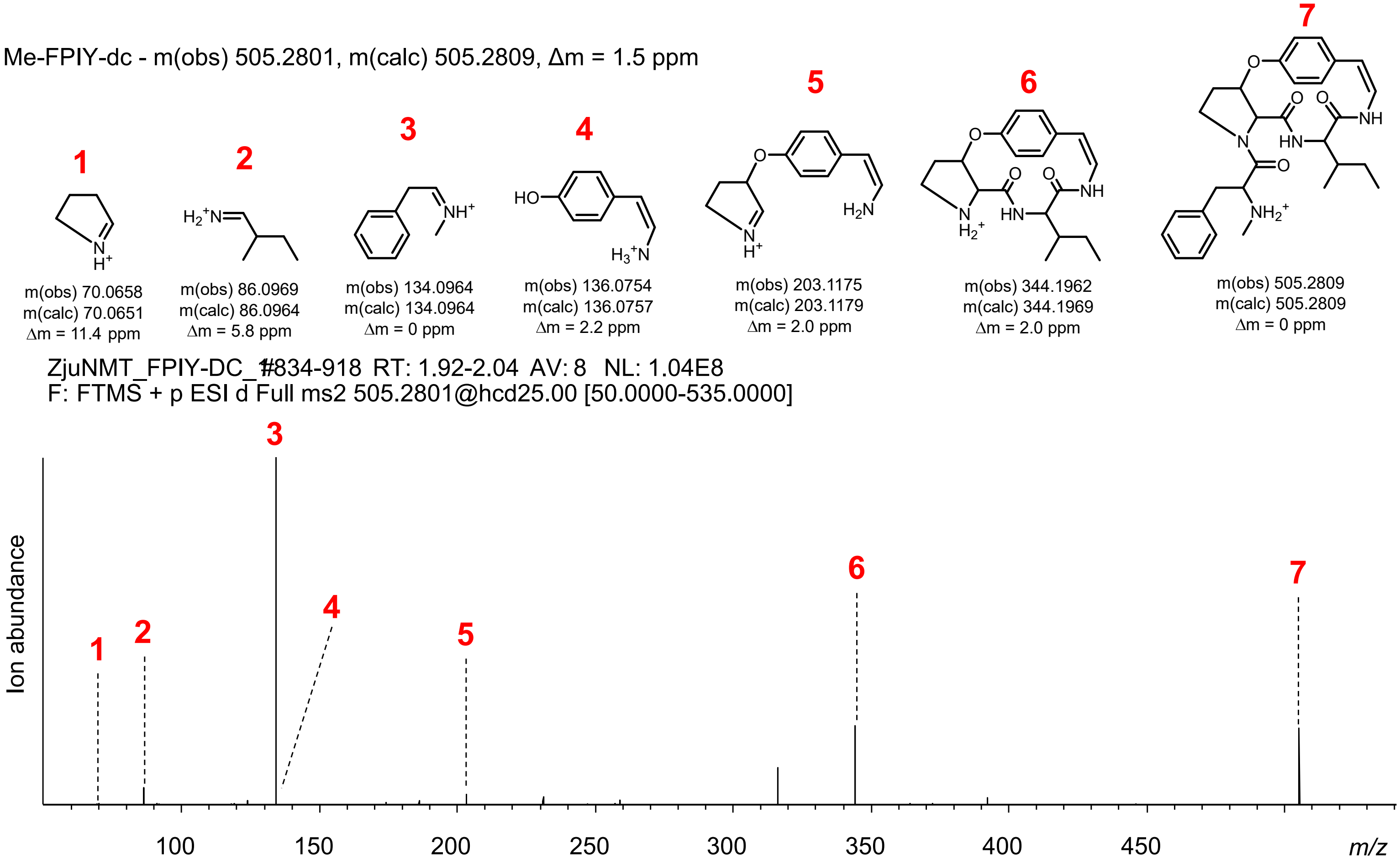


**Figure S36 | ZjuNMT product methylation *in vitro* and *in planta*.** (**C**) MS analysis of trimethylated cyFPIY-dc and monomethylated cyFPIY-dc from *in vitro* enzyme assay of ZjuNMT with cyFPIY-dc substrate.

Me$_3$-FPIY-dc - m(obs) 533.3132, m(calc) 533.3122, Δm = 1.9 ppm

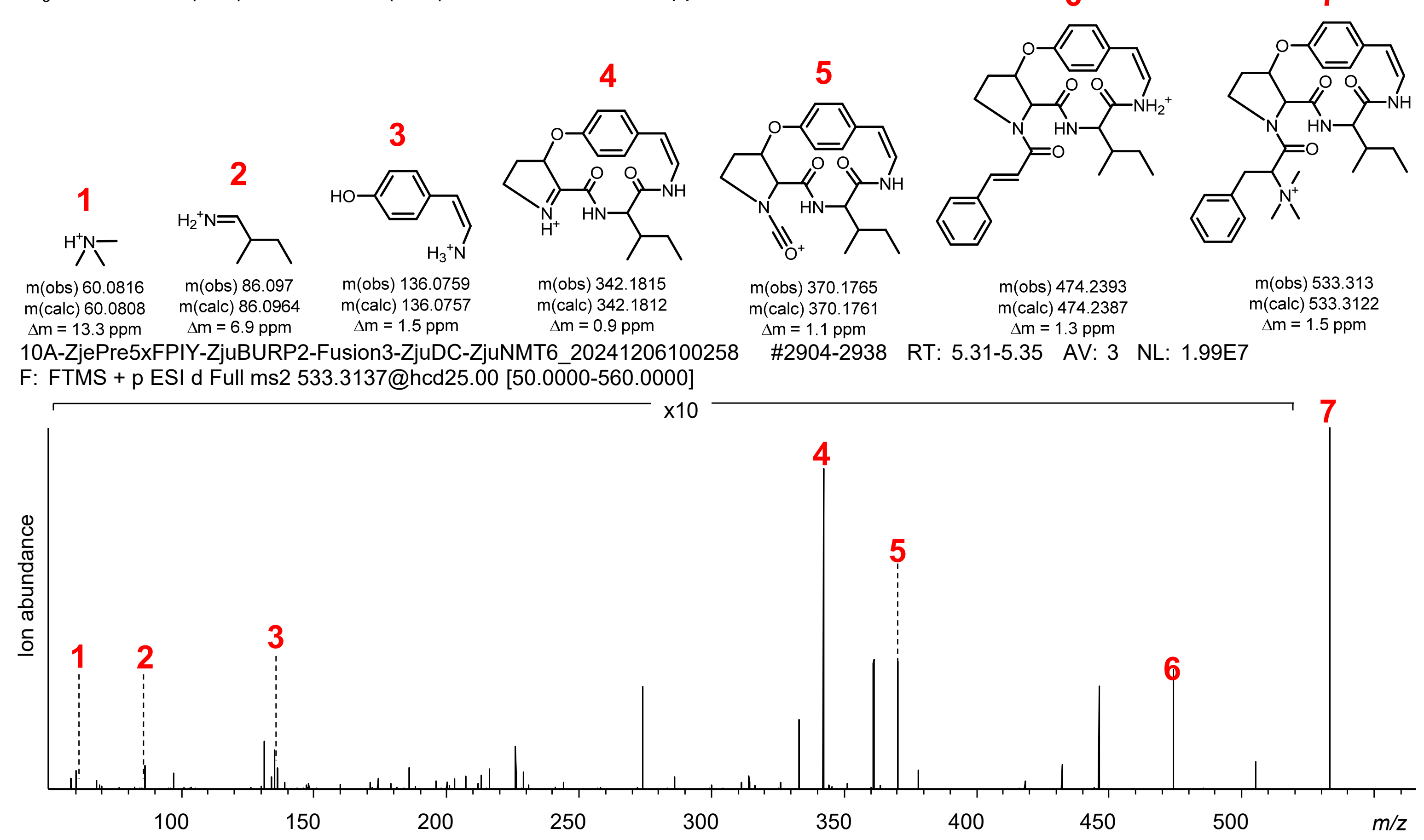


Me-FPIY-dc - m(obs) 505.2811, m(calc) 505.2809, Δm = 0.4 ppm

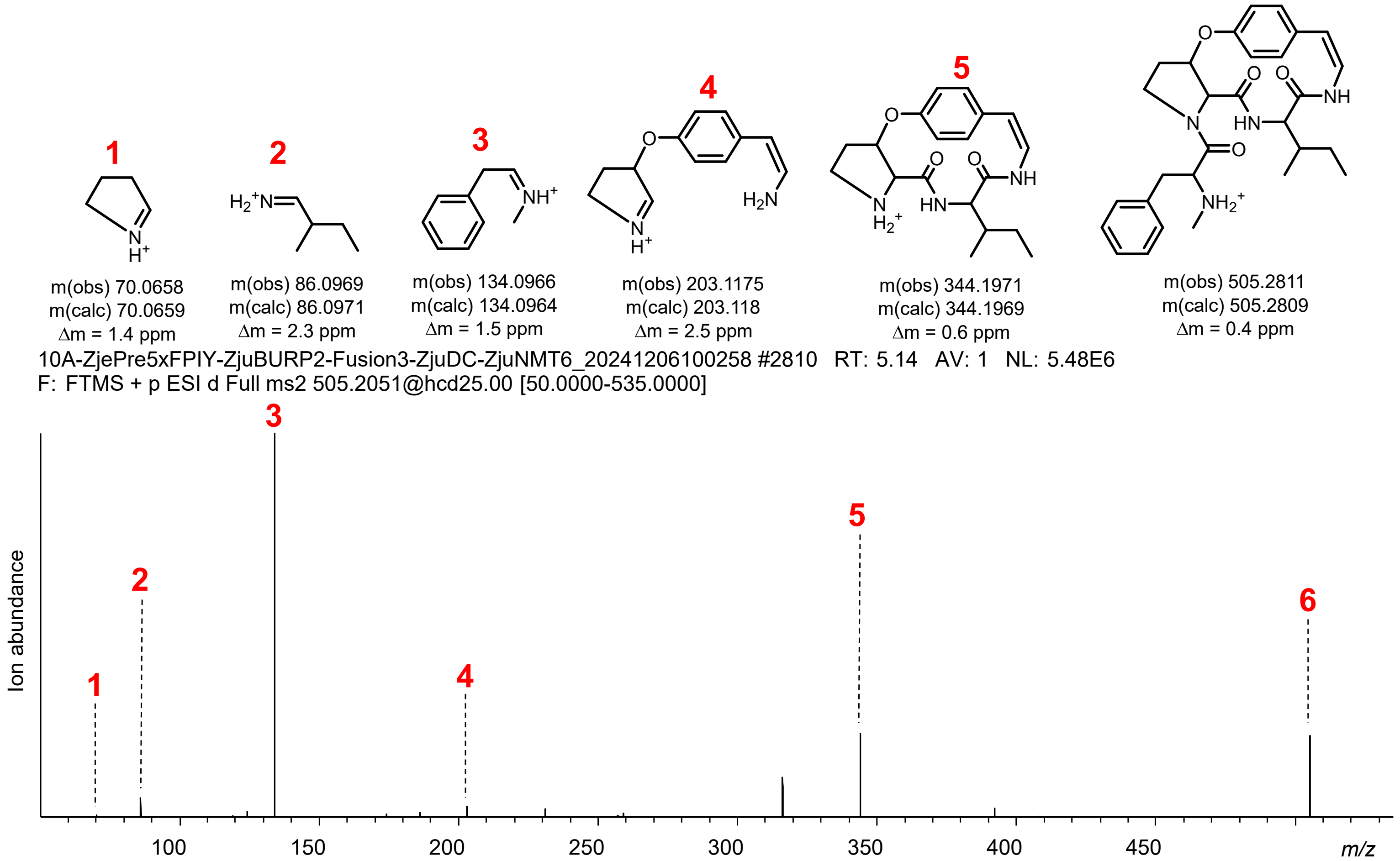


**Figure S36 | ZjuNMT product methylation *in vitro* and *in planta.*** (**D**) MS analysis of trimethylated cyFPIY-dc and monomethylated cyFPIY-dc from transgenic *N. benthamiana* after coexpression of ZjuPrec-5xFPIY-ZjuBURP-fusion with ZjuDC and ZjuNMT.

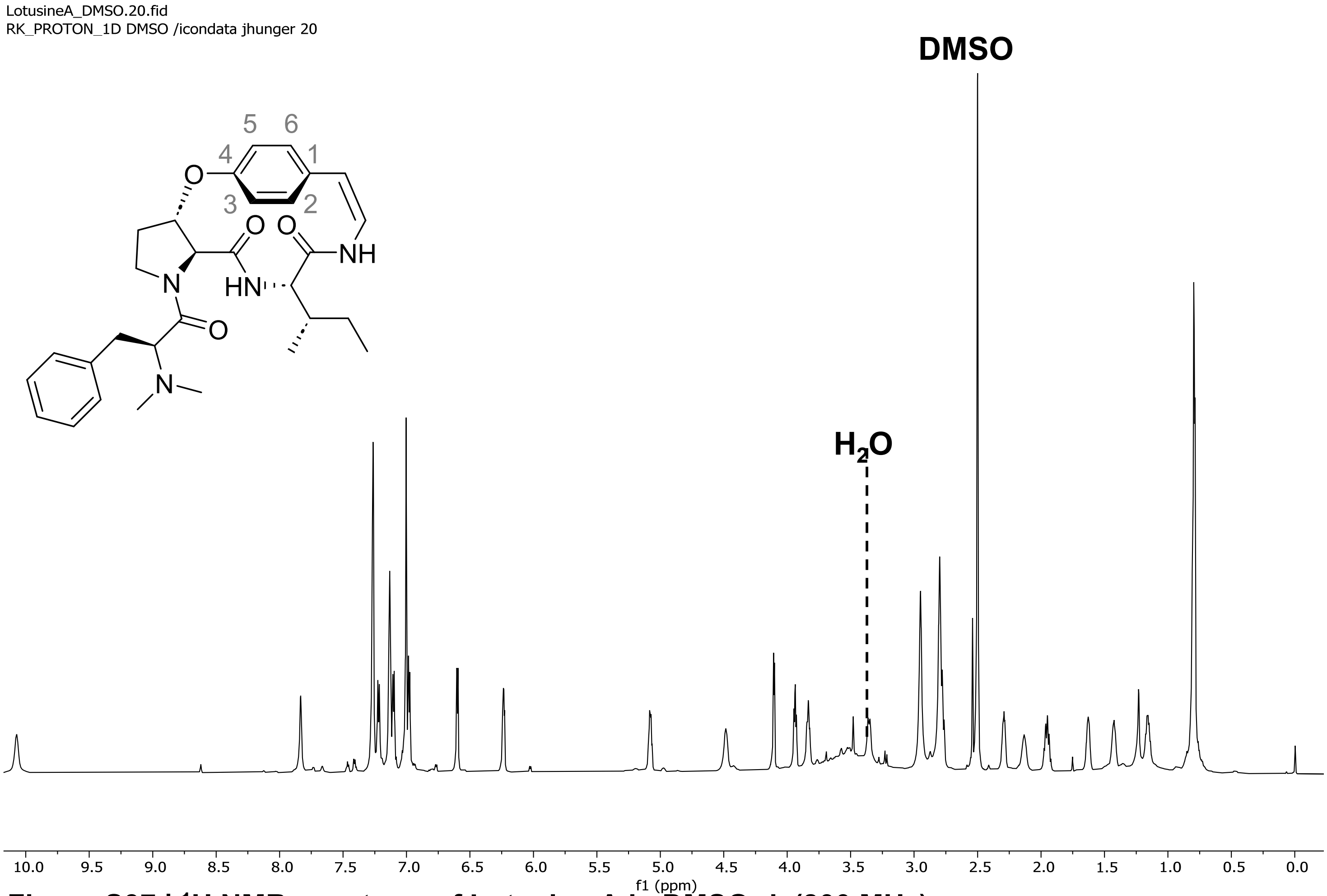


**Figure S37 | $^1$H NMR spectrum of Lotusine A in DMSO-$d_6$ (800 MHz).**

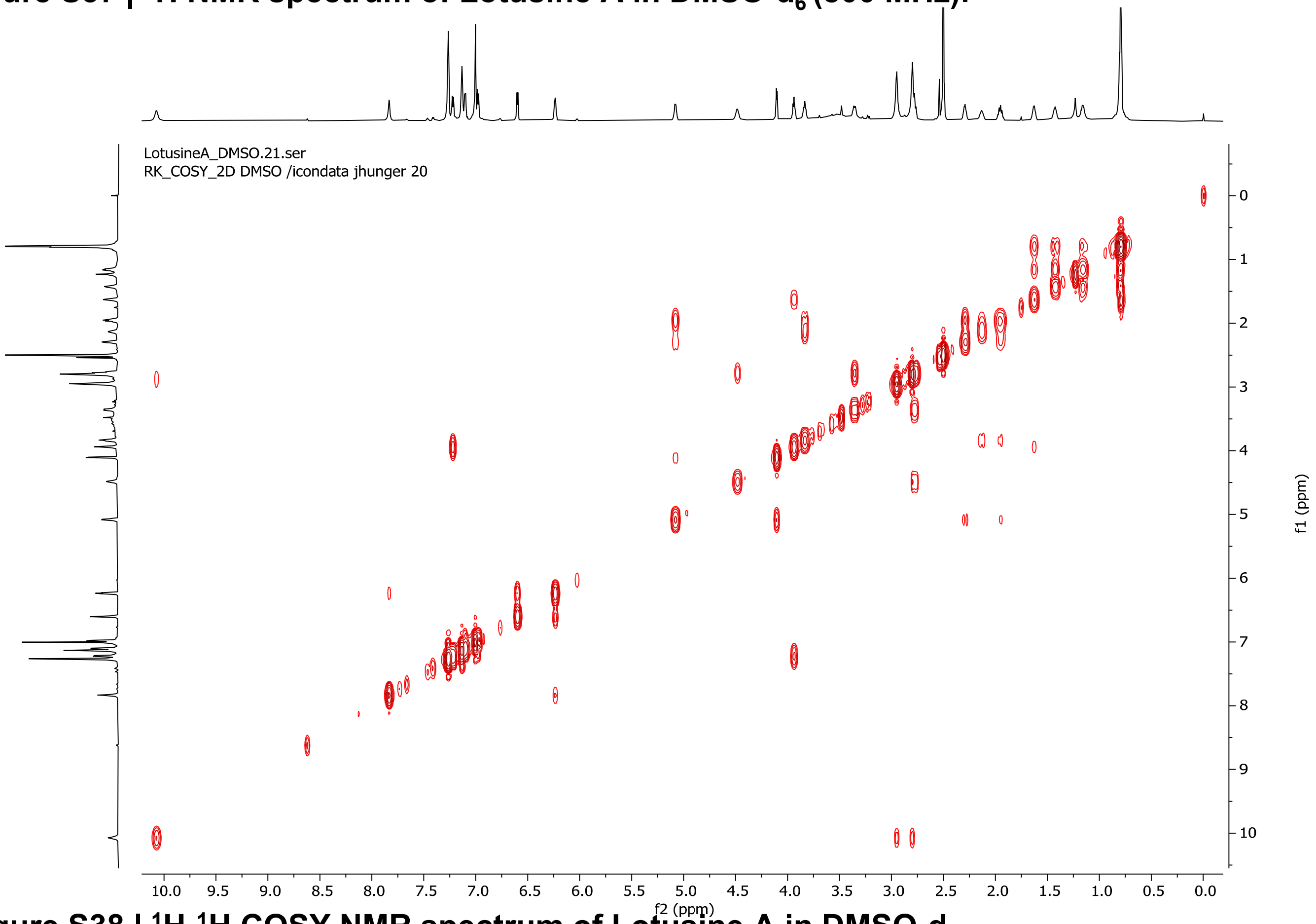


**Figure S38 | $^1$H-$^1$H COSY NMR spectrum of Lotusine A in DMSO-$d_6$.**

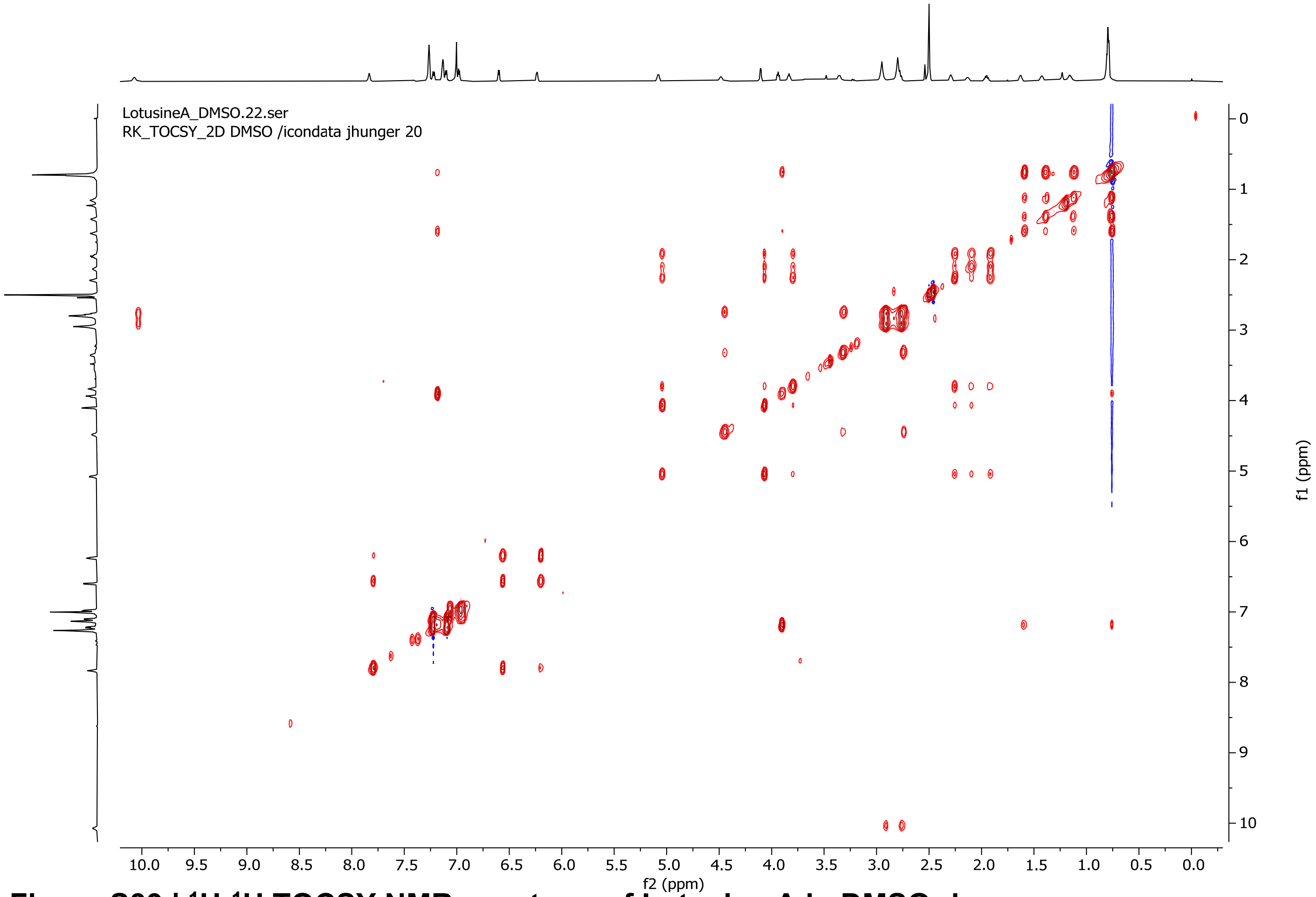


**Figure S39 | $^1$H-$^1$H TOCSY NMR spectrum of Lotusine A in DMSO-$d_6$.**

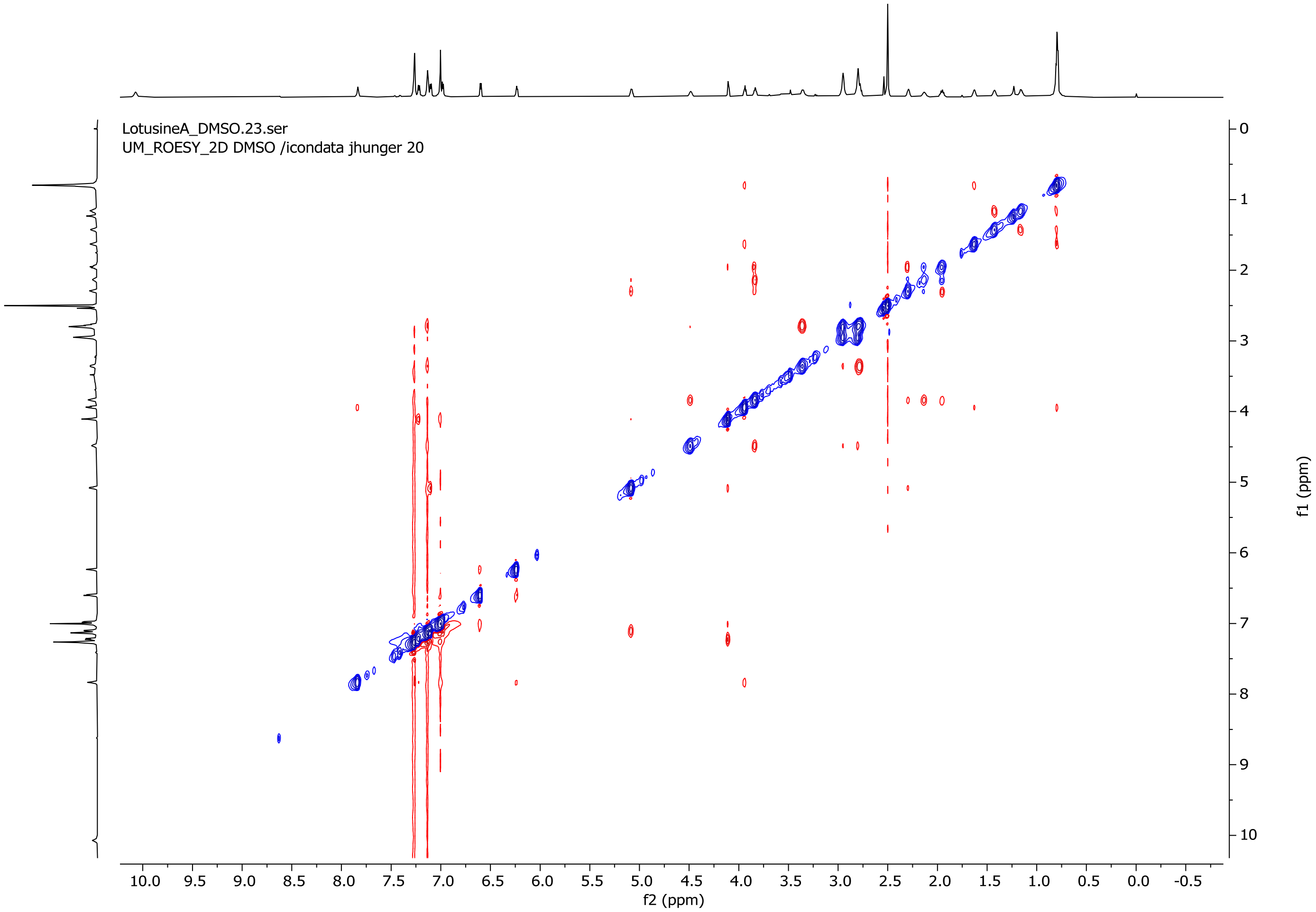


**Figure S40 | ROESY NMR spectrum of Lotusine A in DMSO-$d_6$.**

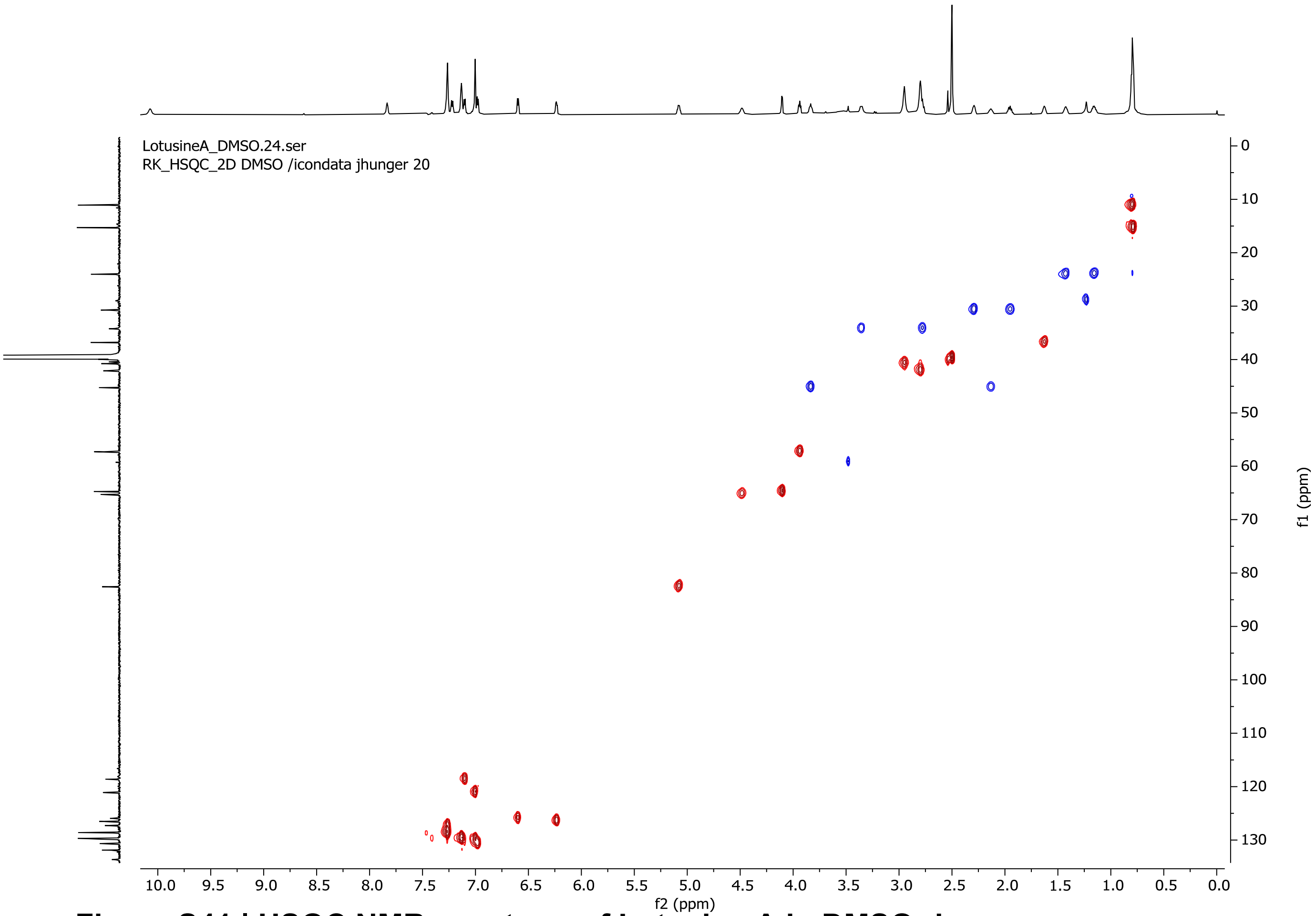


**Figure S41 | HSQC NMR spectrum of Lotusine A in DMSO-d$_6$.**

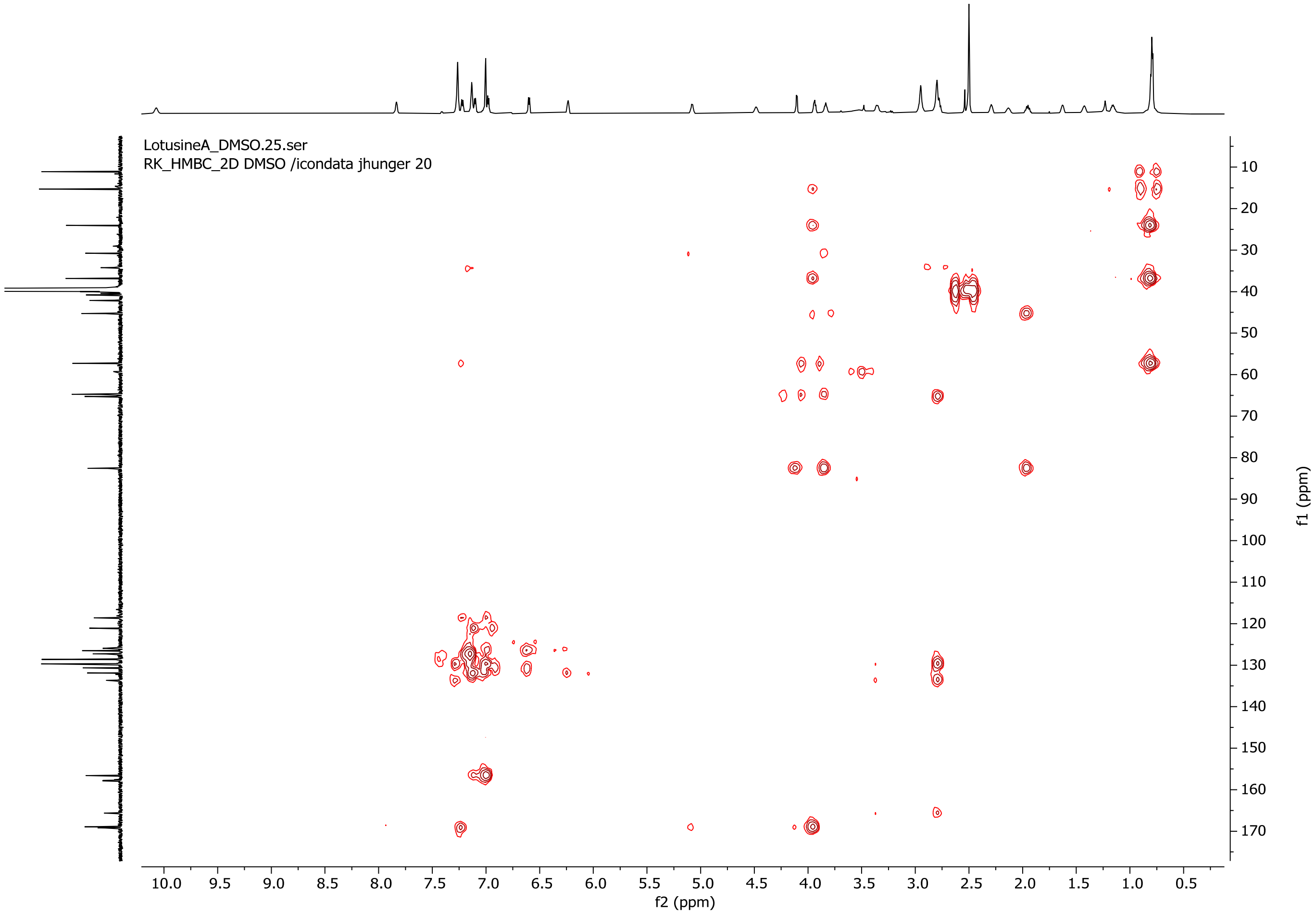


**Figure S42 | HMBC NMR spectrum of Lotusine A in DMSO-d$_6$.**

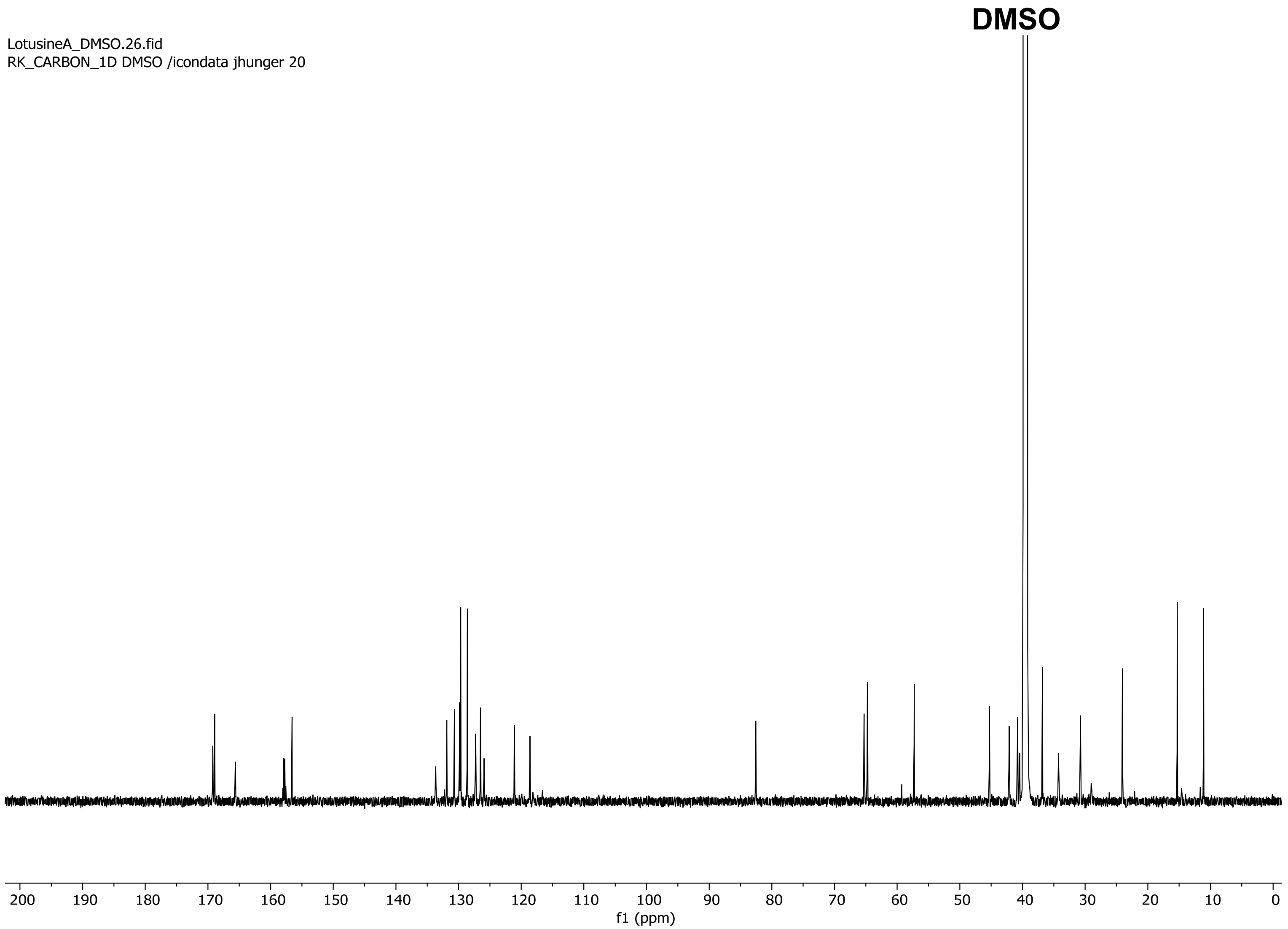


**Figure S43 | $^{13}$C NMR spectrum of Lotusine A in DMSO-$d_6$ (200 MHz).**

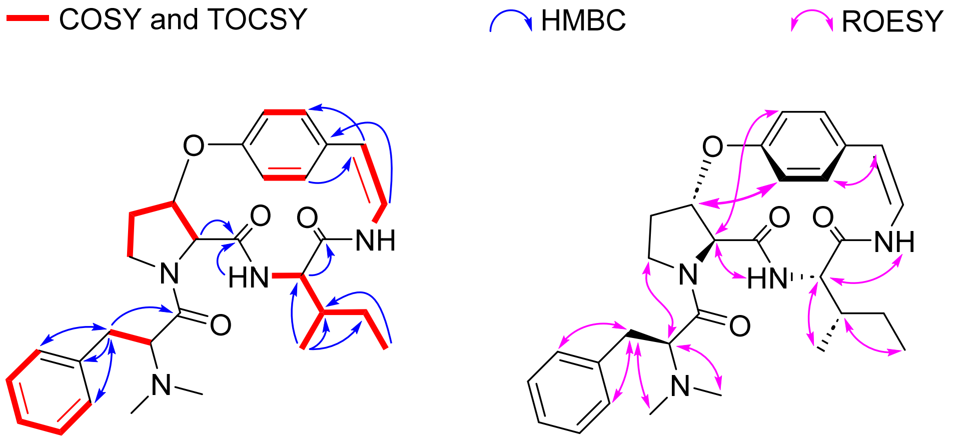


**Figure S44 | Key 2D NMR correlations of Lotusine A in DMSO-$d_6$.**

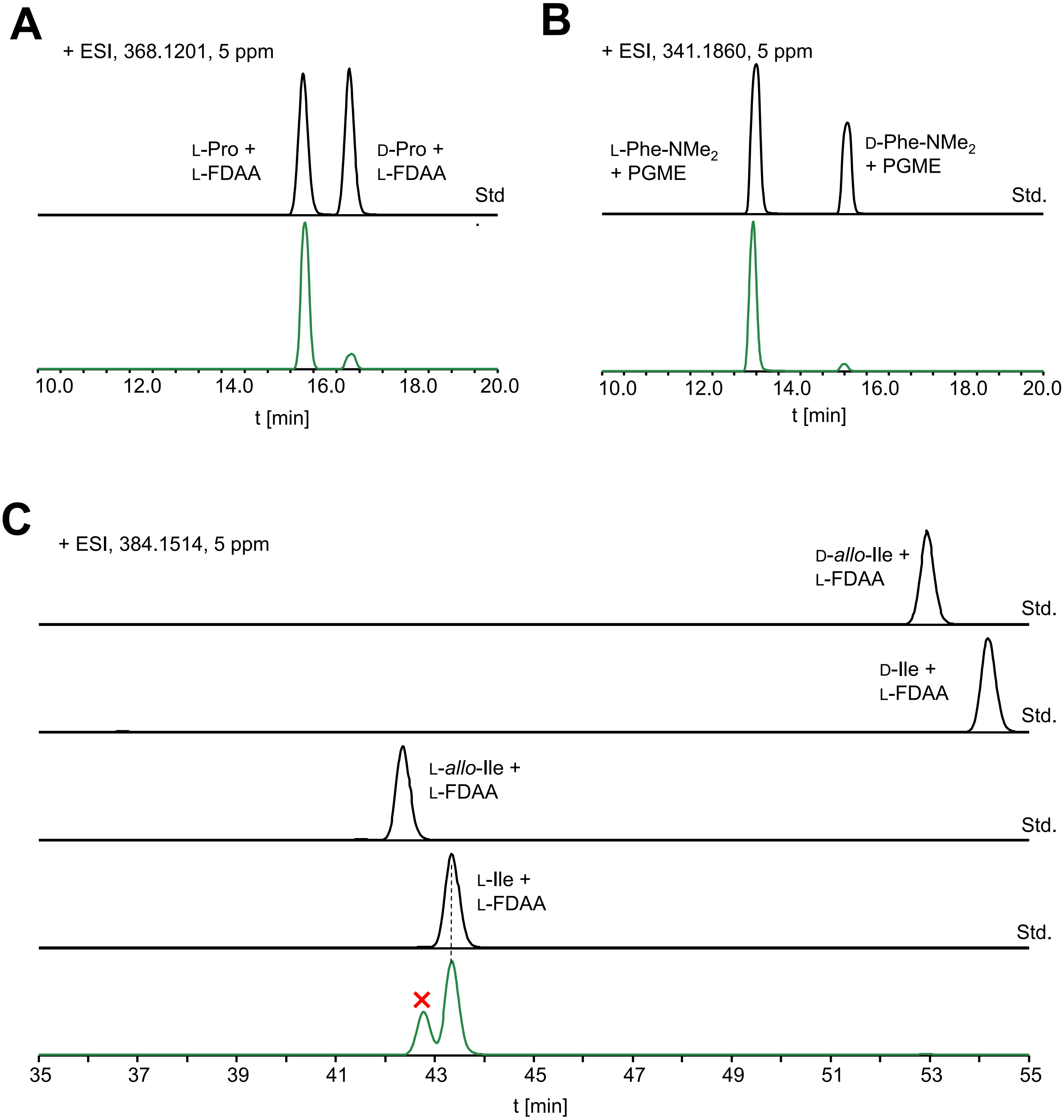


**Figure S45 | Stereochemical analysis of lotusine A.** (**A**) The Pro residue in lotusine A was assigned as L by Marfey's analysis. (**B**) The PheNMe$_2$ residue in lotusine A was assigned as L by PGME derivatization and LC-MS analysis. (**C**) The Ile residue in lotusine A was assigned as L by C3-Marfey's analysis

>SkrBURP-1xFLLYPSY
MAQSLILLFLIVMGCYDVGAIERVHSAKEGVSEAMNDQGHPTIATKAVPASVEDPSQADADNFLLYPSYAARPKAPSTMDHDATHN
AAQNMHHQKGALLFFRMNTLKVGGEVVIPSLASHALGGRKLMTPRLEAMLGSMELPRLLSVLKIPLDSTLARKSKTNLGNCRSPPL
QGEKKACVSSIRSMTNFARSVLEDKKPLEHLNPSRPVSSPEHFKIMDVKVVTENSVVTCHPMVFPYALYMCHFVPKSVPIKVTLQD
DHDKLVVVPVMCHMDTSEFDPSHLSFKILNTKPGEAEMCHWMPNSHIMWYTSDGKTRDVL

>SkrBURP-1xFLLYx(x)F
MAQSLILLFLIVMGCYDVGAIERVHSAKEGVSEAMNDQGHPTIATKAVPASVEDPSQADADNFLLYPSFAARPKAPSTMDHDATHN
AAQNMHHQKGALLFFRMNTLKVGGEVVIPSLASHALGGRKLMTPRLEAMLGSMELPRLLSVLKIPLDSTLARKSKTNLGNCRSPPL
QGEKKACVSSIRSMTNFARSVLEDKKPLEHLNPSRPVSSPEHFKIMDVKVVTENSVVTCHPMVFPYALYMCHFVPKSVPIKVTLQD
DHDKLVVVPVMCHMDTSEFDPSHLSFKILNTKPGEAEMCHWMPNSHIMWYTSDGKTRDVL

>SkrBURP-5xFLLYx(x)F
MAQSLILLFLIVMGCYDVGAIERVHSAKEGVSEAMNDQGHPTIATKAVPASVEDPSQADADNFLLYPSFAAKKAVPEDPSQADADN
FLLYRSFAAKKAVPASVEDPSHDADNFLLYPSFVAKKAVPASVEDPSHDADNFLLYPSFVAKKAVPASVEDPSQADADNFLLYPSF
AARPKAPSTMDHDATHNAAQNMHHQKGALLFFRMNTLKVGGEVVIPSLASHALGGRKLMTPRLEAMLGSMELPRLLSVLKIPLDST
LARKSKTNLGNCRSPPLQGEKKACVSSIRSMTNFARSVLEDKKPLEHLNPSRPVSSPEHFKIMDVKVVTENSVVTCHPMVFPYALY
MCHFVPKSVPIKVTLQDDHDKLVVVPVMCHMDTSEFDPSHLSFKILNTKPGEAEMCHWMPNSHIMWYTSDGKTRDVL

>SkrBURP-5xFLLY
MAQSLILLFLIVMGCYDVGAIERVHSAKEGVSEAMNDQGHPTIATKAVPASVEDPSQADADNFLLYPSYAAKKAVPEDPSQADADN
FLLYRSYAAKKAVPASVEDPSHDADNFLLYPSYVAKKAVPASVEDPSHDADNFLLYPSYVAKKAVPASVEDPSQADADNFLLYPSY
AARPKAPSTMDHDATHNAAQNMHHQKGALLFFRMNTLKVGGEVVIPSLASHALGGRKLMTPRLEAMLGSMELPRLLSVLKIPLDST
LARKSKTNLGNCRSPPLQGEKKACVSSIRSMTNFARSVLEDKKPLEHLNPSRPVSSPEHFKIMDVKVVTENSVVTCHPMVFPYALY
MCHFVPKSVPIKVTLQDDHDKLVVVPVMCHMDTSEFDPSHLSFKILNTKPGEAEMCHWMPNSHIMWYTSDGKTRDVL

>SkrBURP-5xLLIYx(x)F
MAQSLILLFLIVMGCYDVGAIERVHSAKEGVSEAMNDQGHPTIATKAVPASVEDPSQADADNLLIYPSFAAKKAVPEDPSQADADN
LLIYRSFAAKKAVPASVEDPSHDADNLLIYPSFVAKKAVPASVEDPSHDADNLLIYPSFVAKKAVPASVEDPSQADADNLLIYPSF
AARPKAPSTMDHDATHNAAQNMHHQKGALLFFRMNTLKVGGEVVIPSLASHALGGRKLMTPRLEAMLGSMELPRLLSVLKIPLDST
LARKSKTNLGNCRSPPLQGEKKACVSSIRSMTNFARSVLEDKKPLEHLNPSRPVSSPEHFKIMDVKVVTENSVVTCHPMVFPYALY
MCHFVPKSVPIKVTLQDDHDKLVVVPVMCHMDTSEFDPSHLSFKILNTKPGEAEMCHWMPNSHIMWYTSDGKTRDVL

**Figure S46 | Burpitide cyclase SkrBURP constructs for optimization of cyFLLY production.** Core peptides are highlighted in red, BURP-domain sequence is underlined.

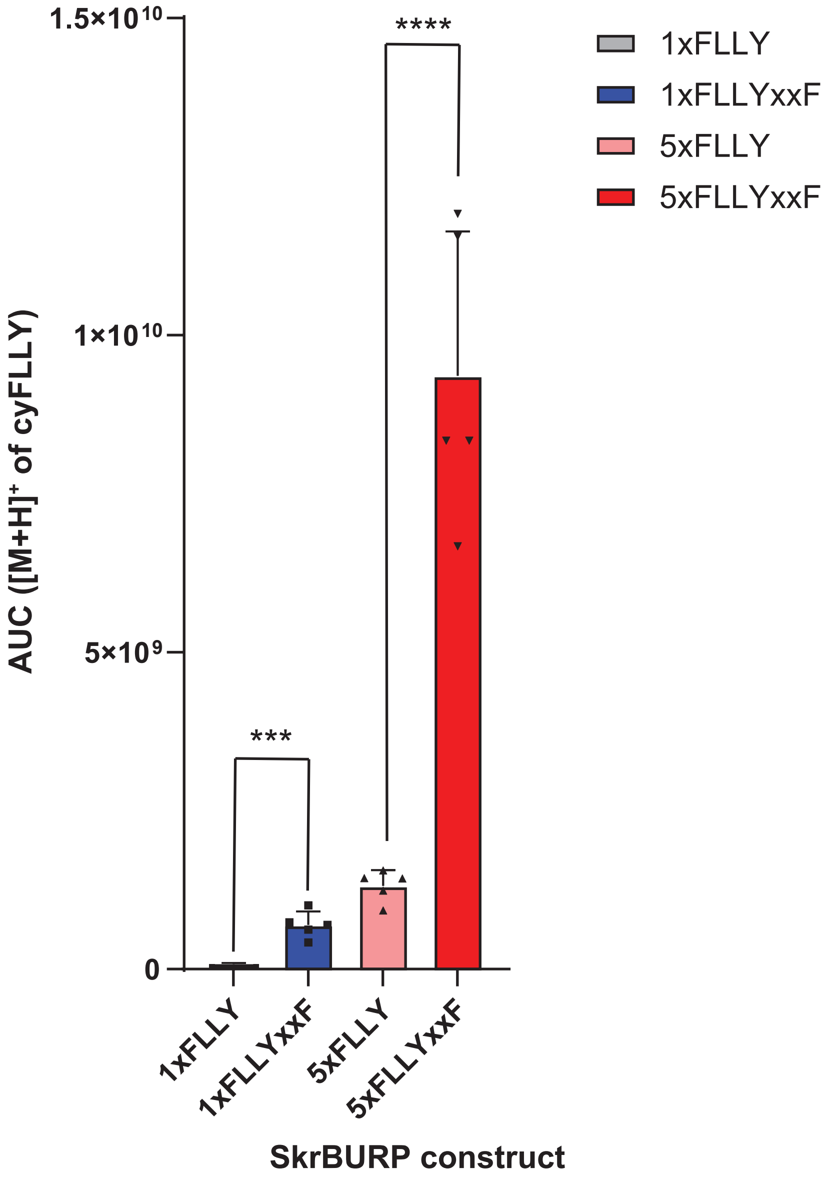


**Figure S47 | SkrBURP optimization for cyclic peptide yields from transgenic *Nicotiana benthamiana.*** AUC of cyFLLY analyte $(M+H]^+)$ for four SkrBURP constructs expressed for 7 days in *N. benthamiana* via pEAQ-HT constructs after agroinfiltration. Bar graphs represent means (*n*=5), error bars represent one standard deviation. FLLY and FLLYxxY constructs were compared by unpaired t-test (*** - p = 0.0002, **** p <0.0001).

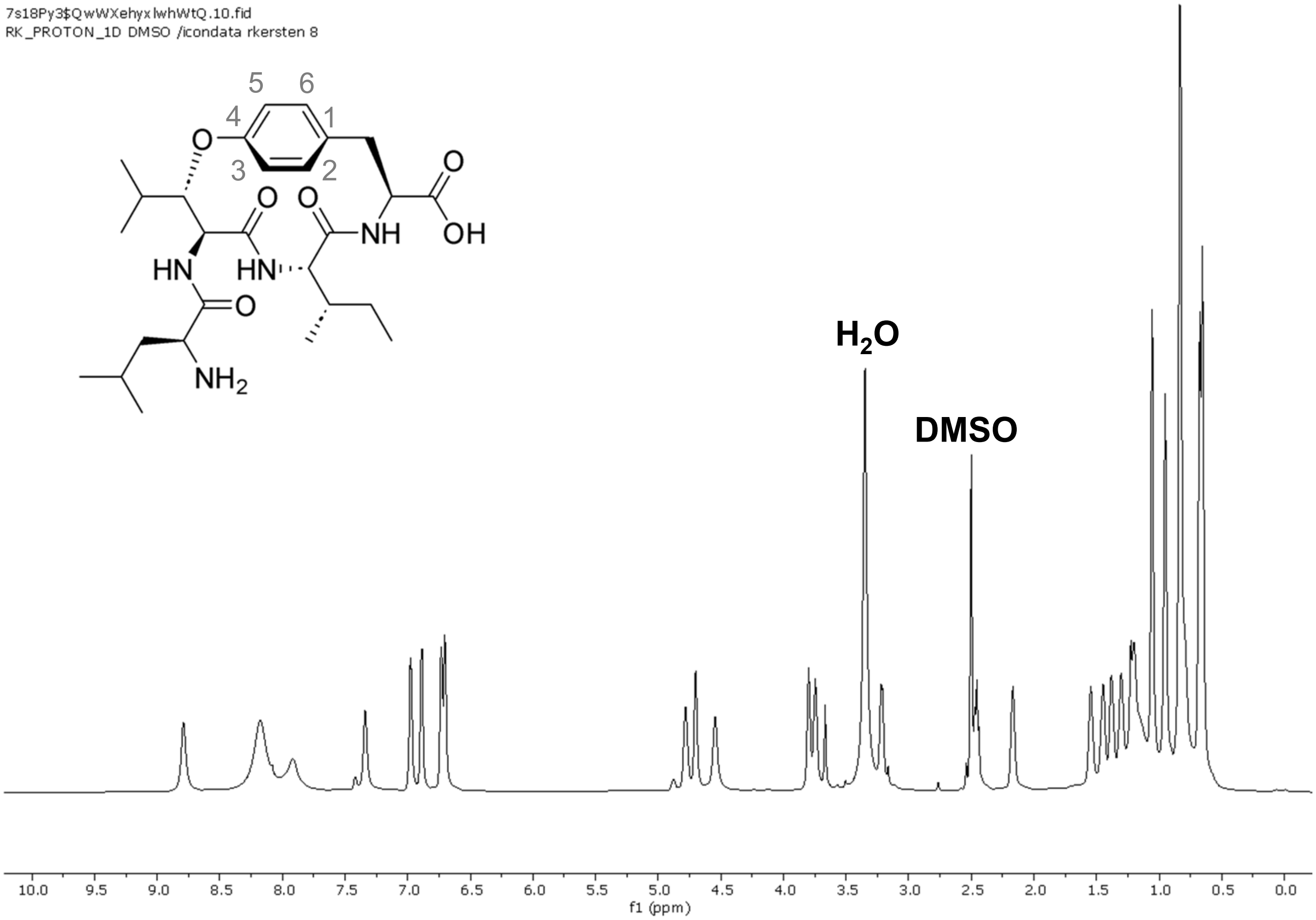


**Figure S48 | [1]H NMR spectrum of cyLLIY in DMSO-$d_6$ (800 MHz).**

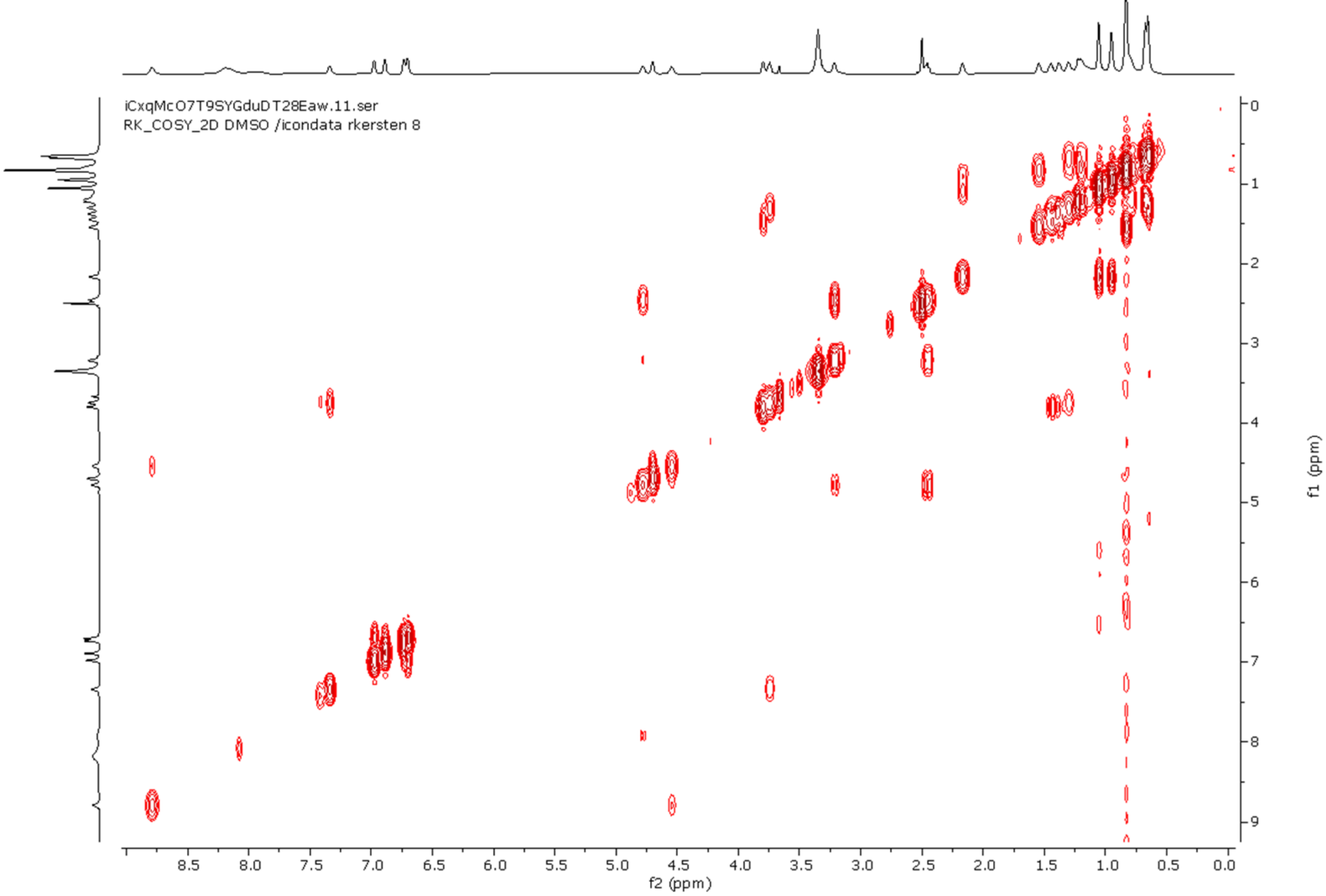


**Figure S49 | [1]H-[1]H COSY-NMR spectrum of cyLLIY in DMSO-$d_6$.**

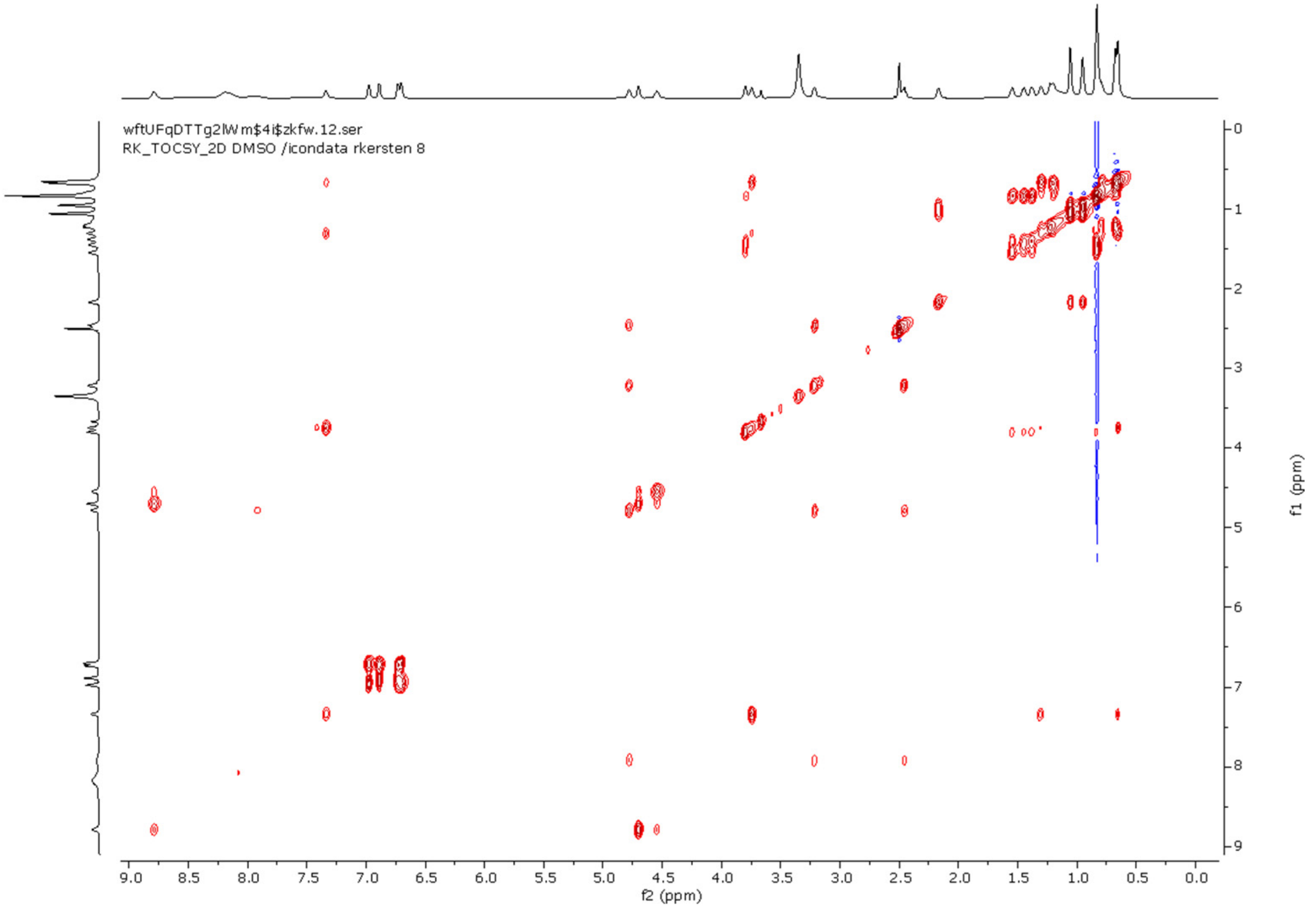


**Figure S50 | $^{1}H$-$^{1}H$ TOCSY NMR spectrum of cyLLIY in DMSO-d$_6$.**

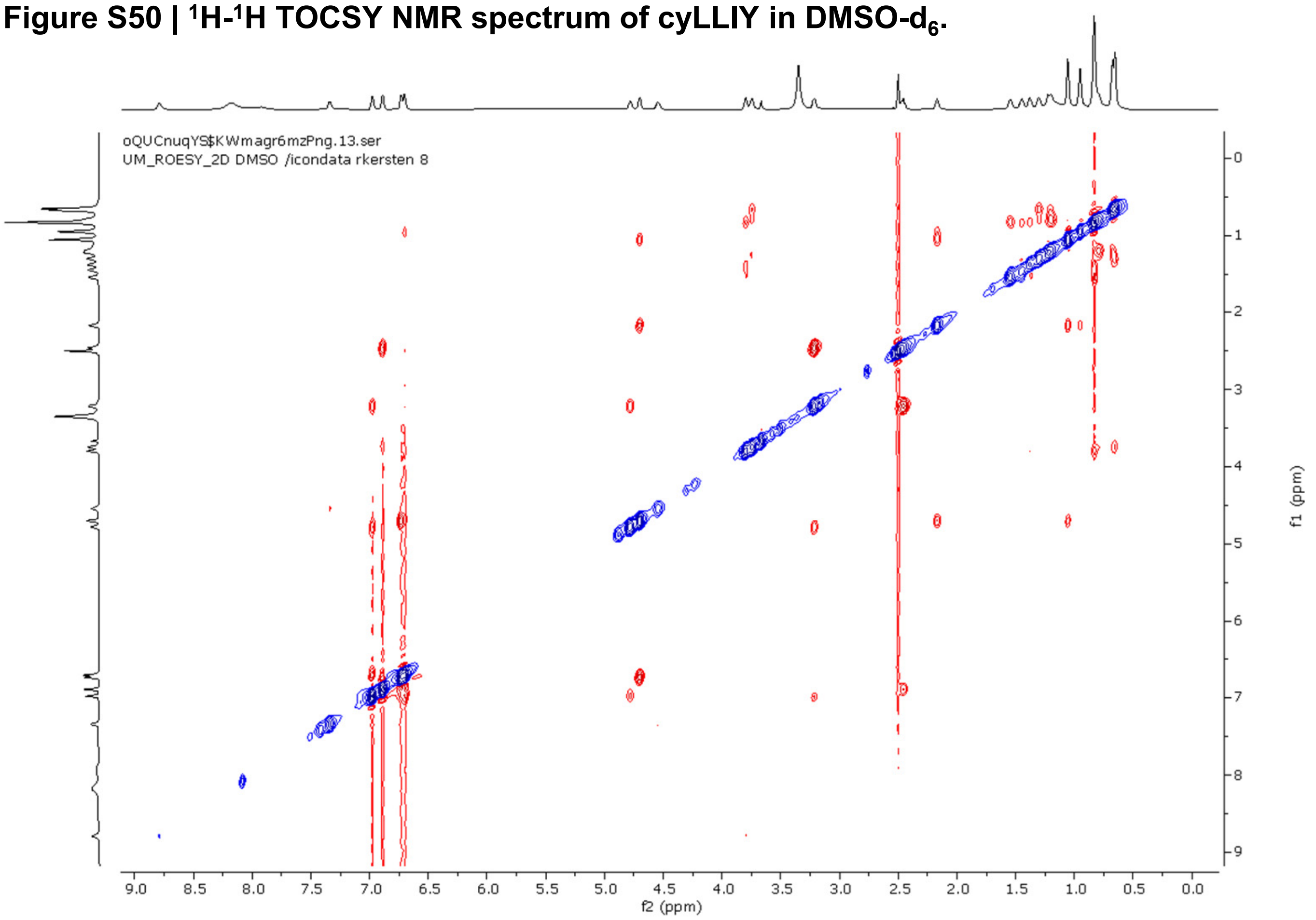


**Figure S51 | ROESY NMR spectrum of cyLLIY in DMSO-d$_6$.**

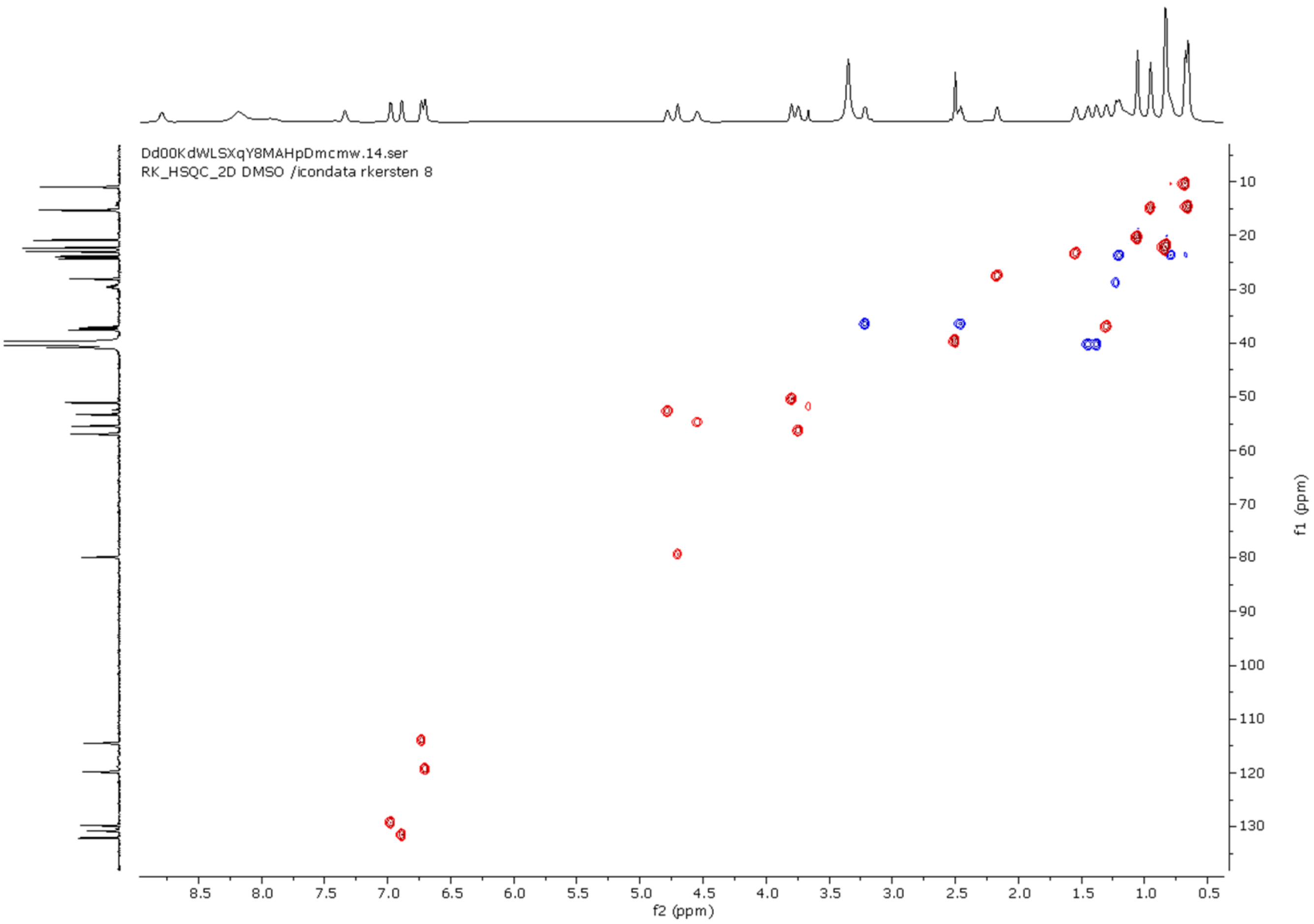


**Figure S52 | HSQC NMR spectrum of cyLLIY in DMSO-$d_6$.**

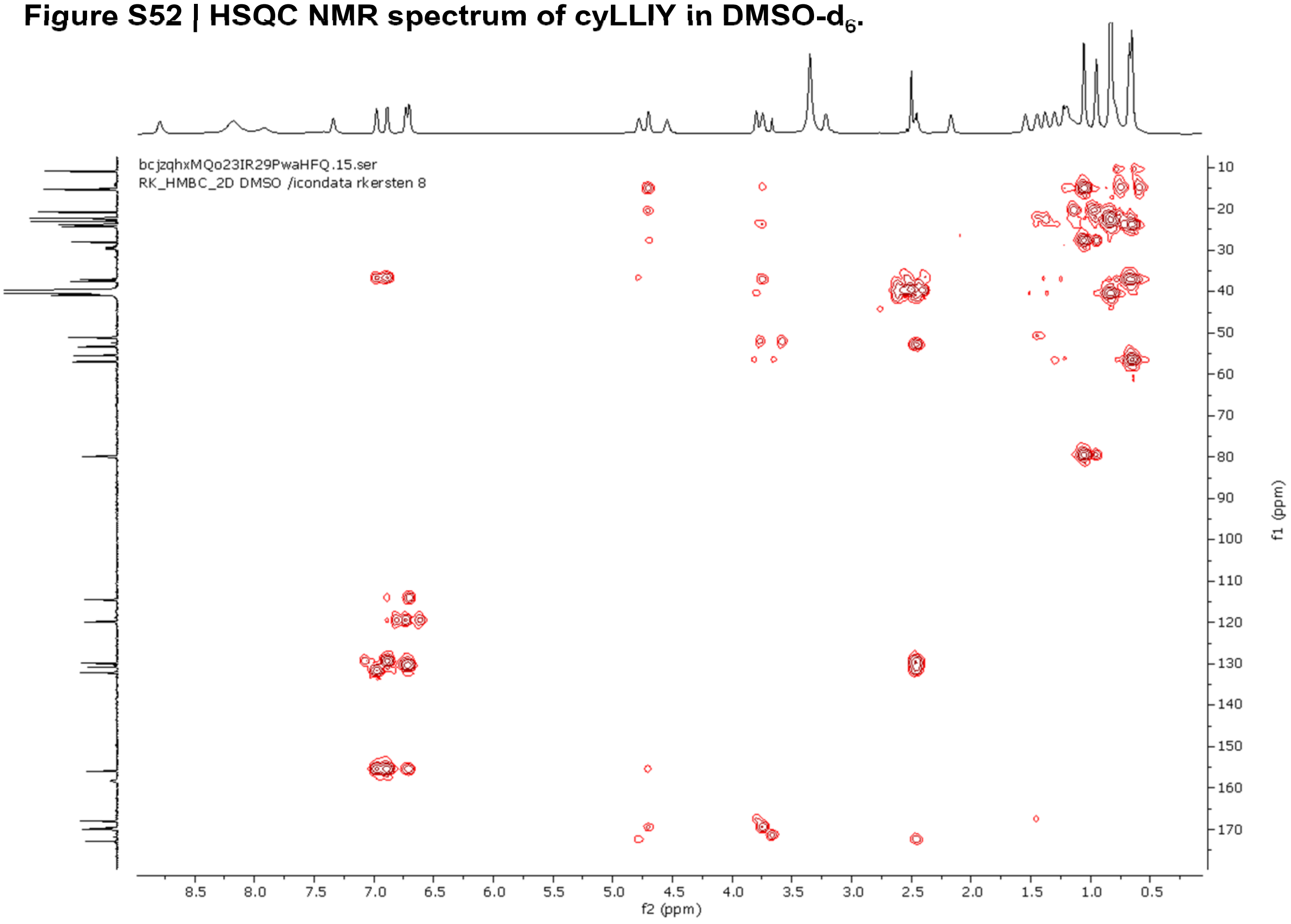


**Figure S53 | HMBC NMR spectrum of cyLLIY in DMSO-$d_6$.**

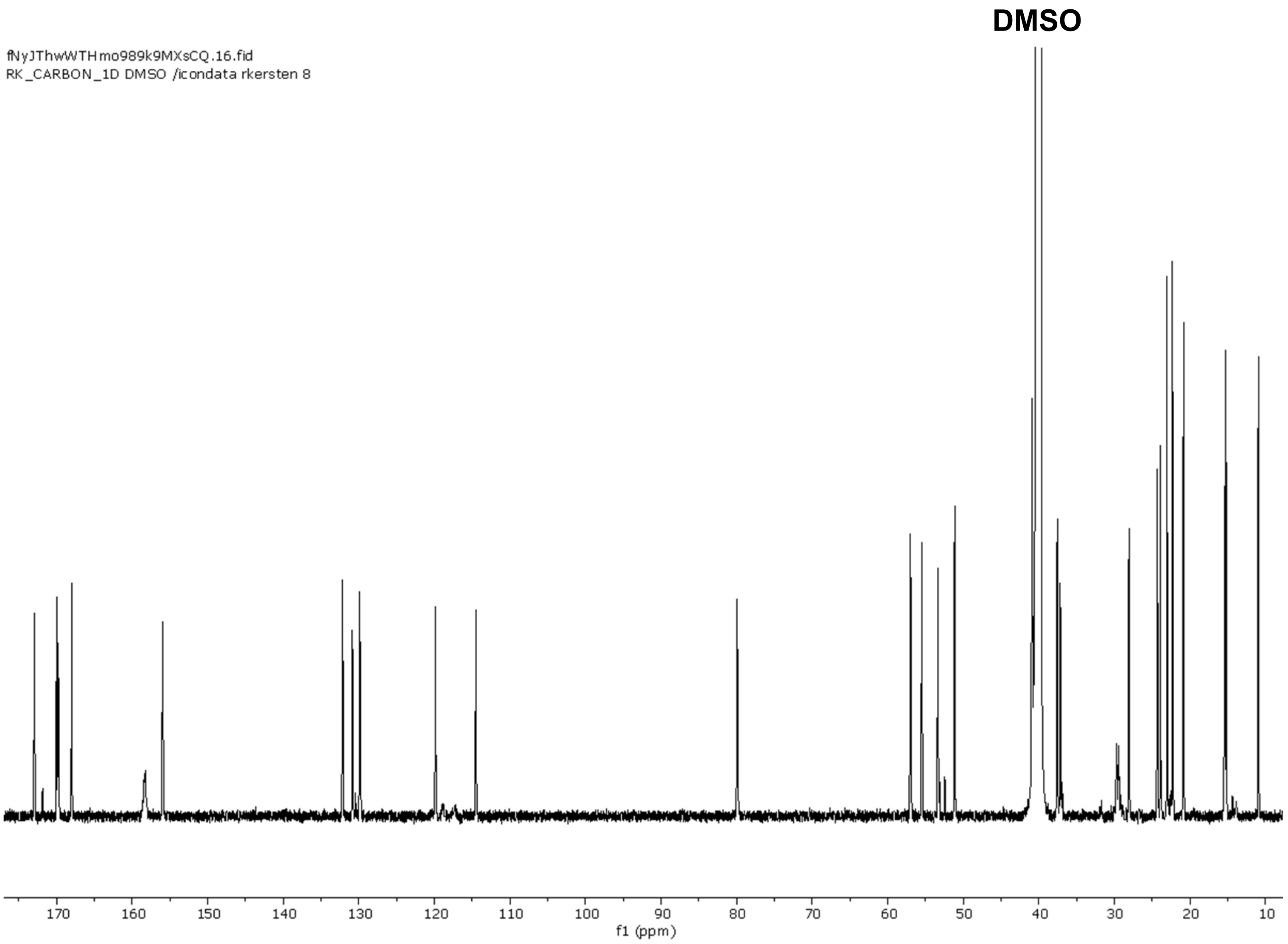


**Figure S54 | $^{13}$C NMR spectrum of cyLLIY in DMSO-$d_6$ (200 MHz).**

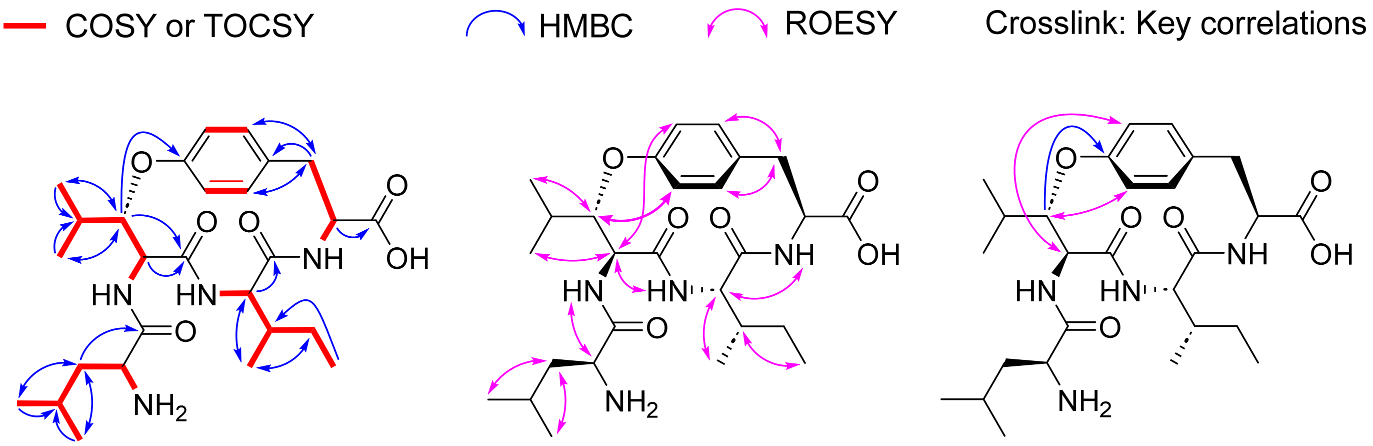


**Figure S55 | Key 2D NMR correlations of cyLLIY in DMSO-$d_6$.**

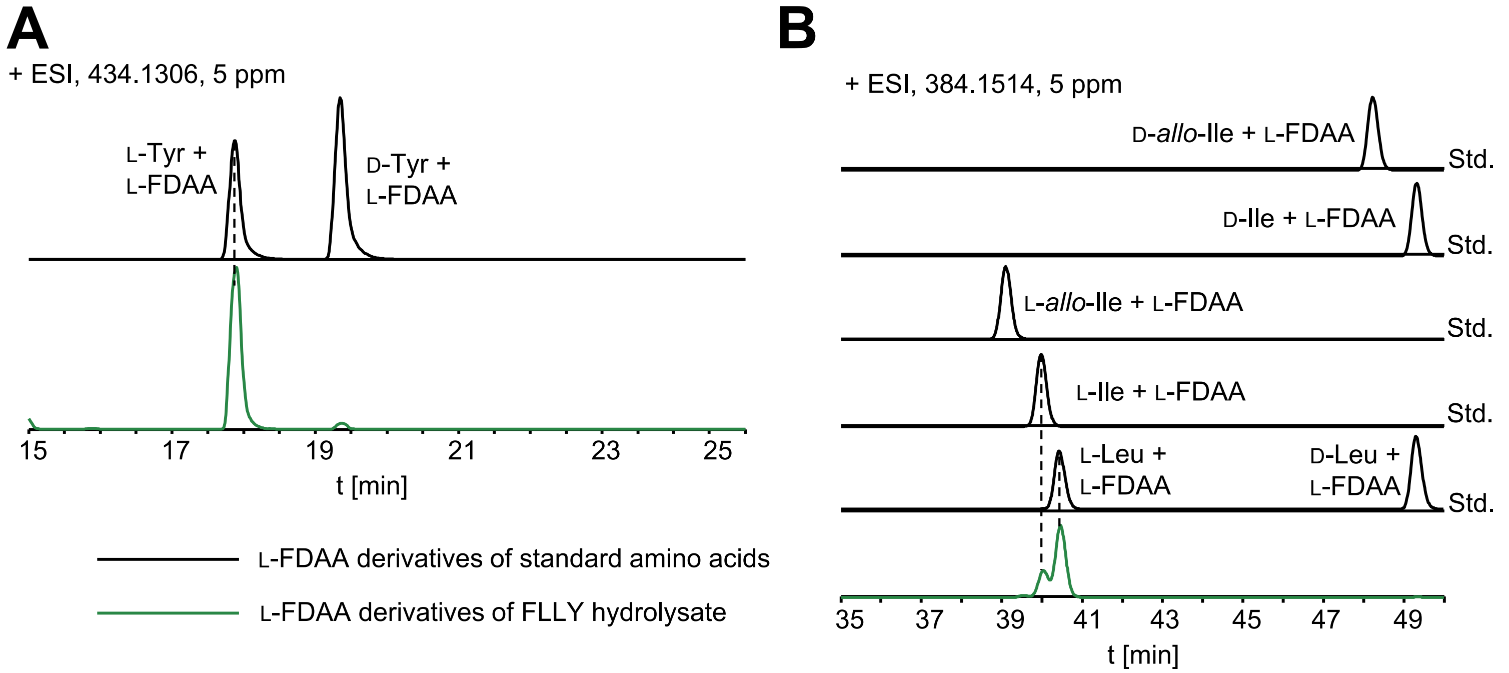


**Figure S56 | Stereochemical analysis of cyLLIY.** (**A**) The Tyr residue in cyLLIY was assigned as L by using Marfey's analysis. (**B**) The two Leu and Ile residues in cyLLIY were assigned as L by using C3-Marfey's analysis.

Vsil2OiLS7CVJF4h4Jq5SA.10.fid
RK_PROTON_1D DMSO /icondata rkersten 2

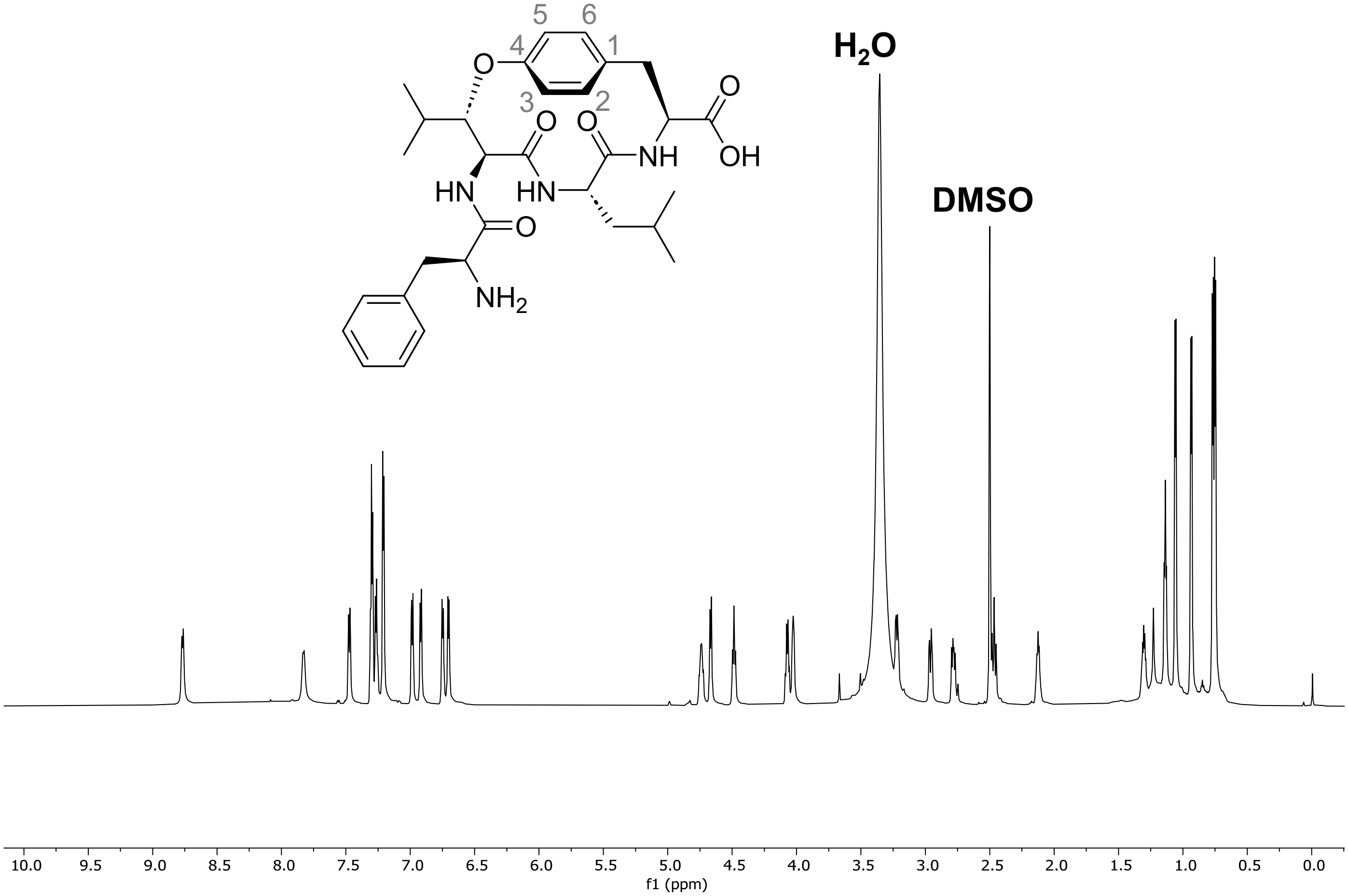


**Figure S57 | $^1$H NMR spectrum of cyFLLY in DMSO-d$_6$ (800MHz).**

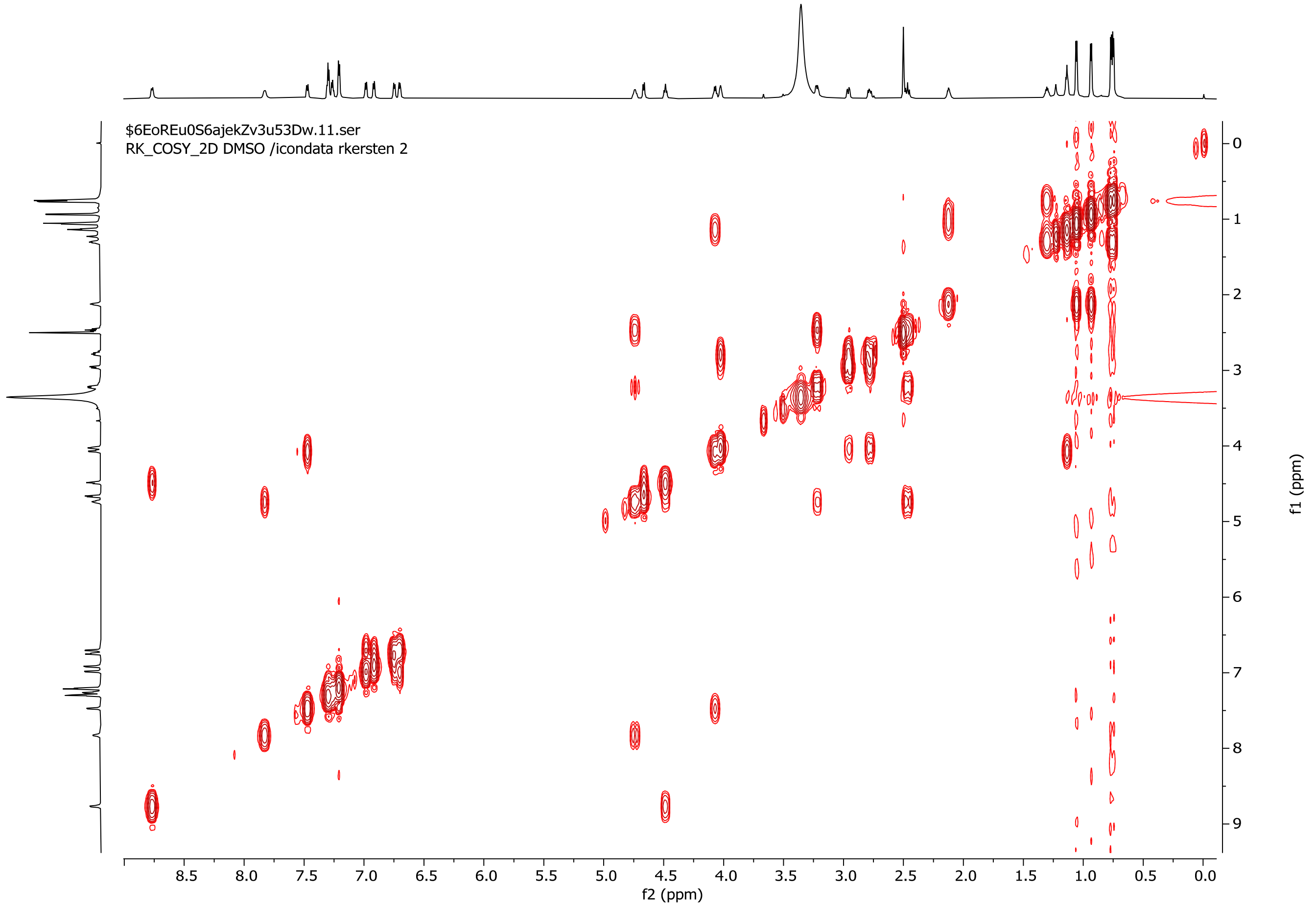


**Figure S58 | $^1$H-$^1$H COSY NMR spectrum of cyFLLY in DMSO-d$_6$.**

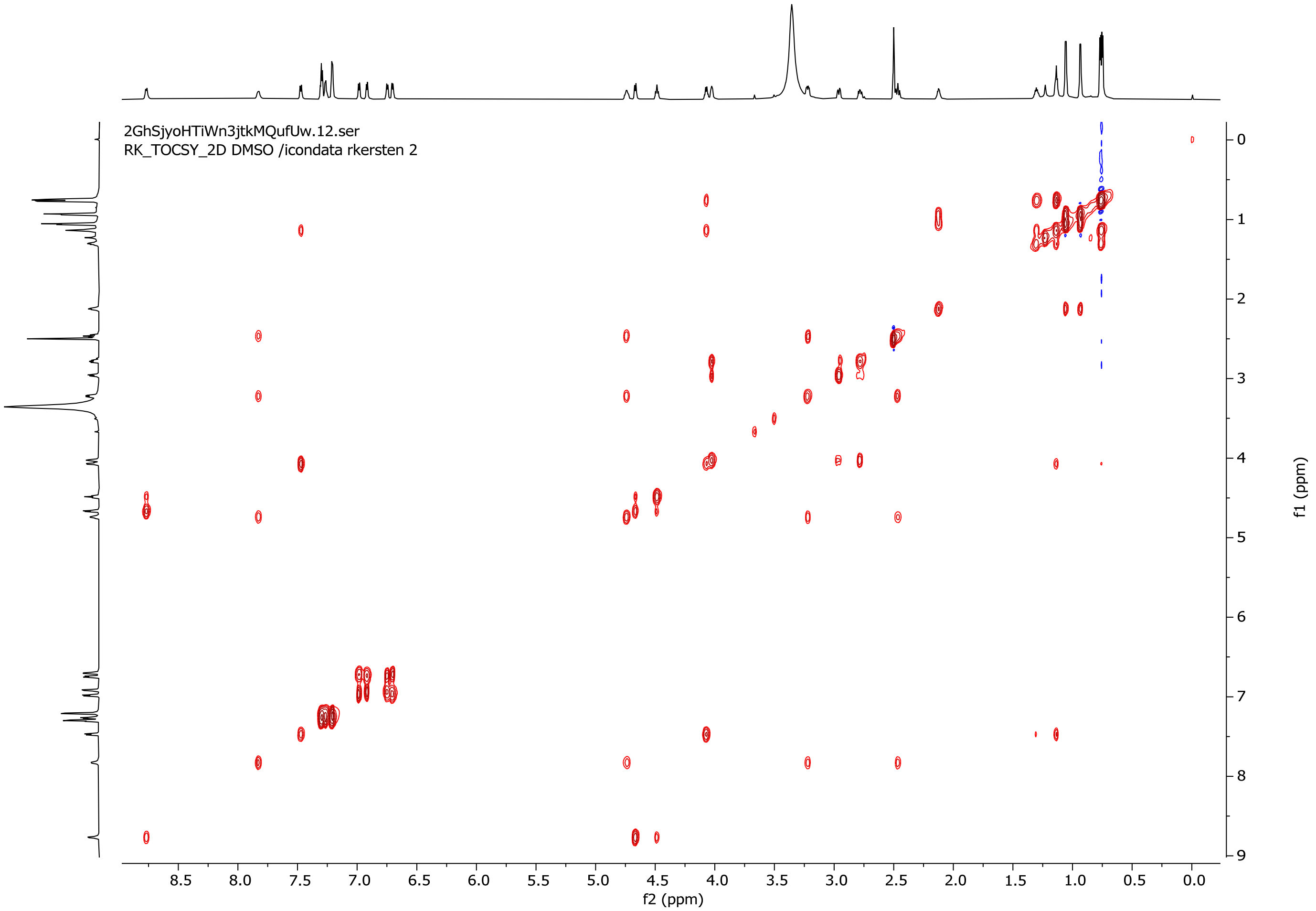


**Figure S59 | $^{1}H$-$^{1}H$ TOCSY NMR spectrum of cyFLLY in DMSO-$d_6$.**

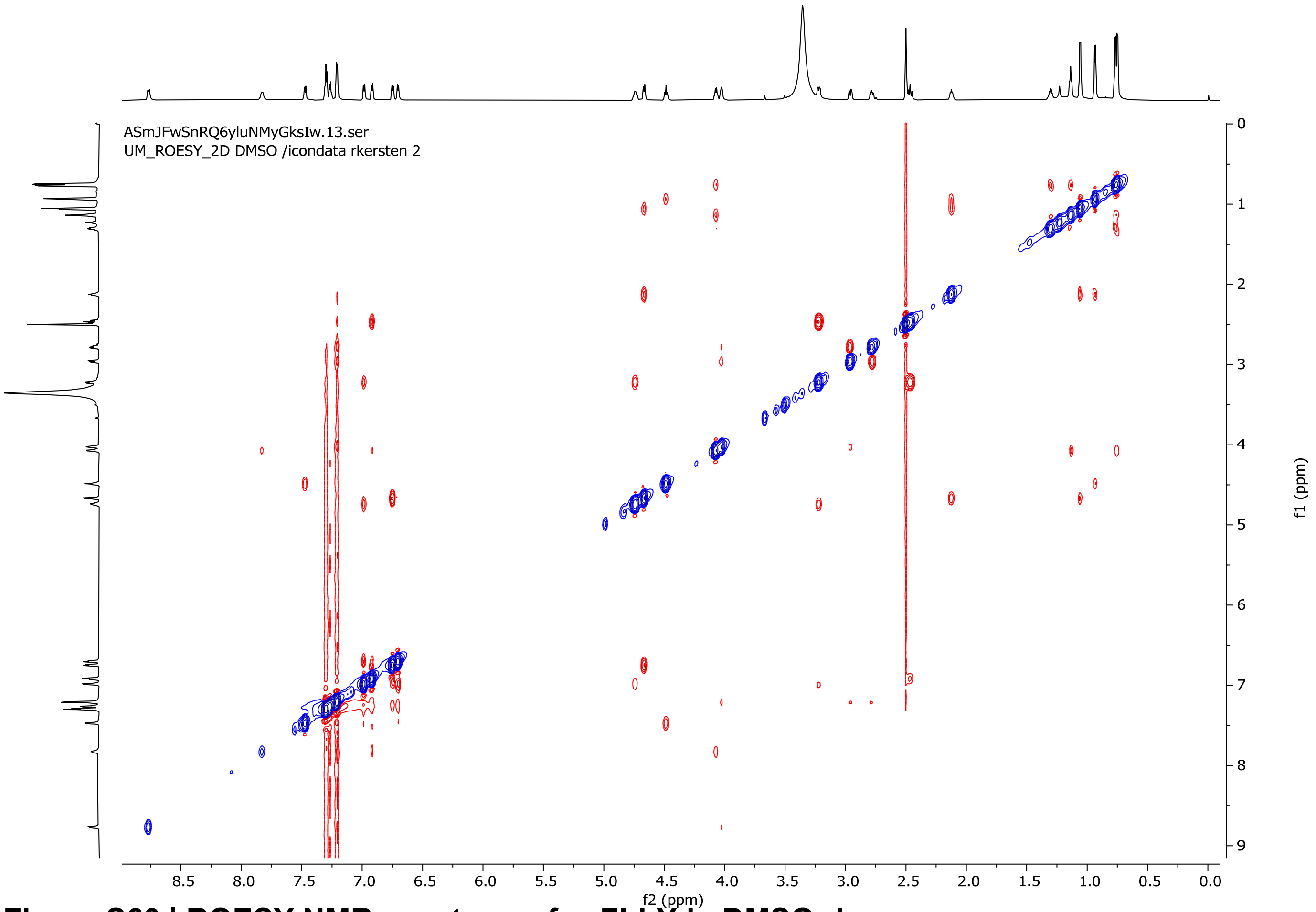


**Figure S60 | ROESY NMR spectrum of cyFLLY in DMSO-$d_6$.**

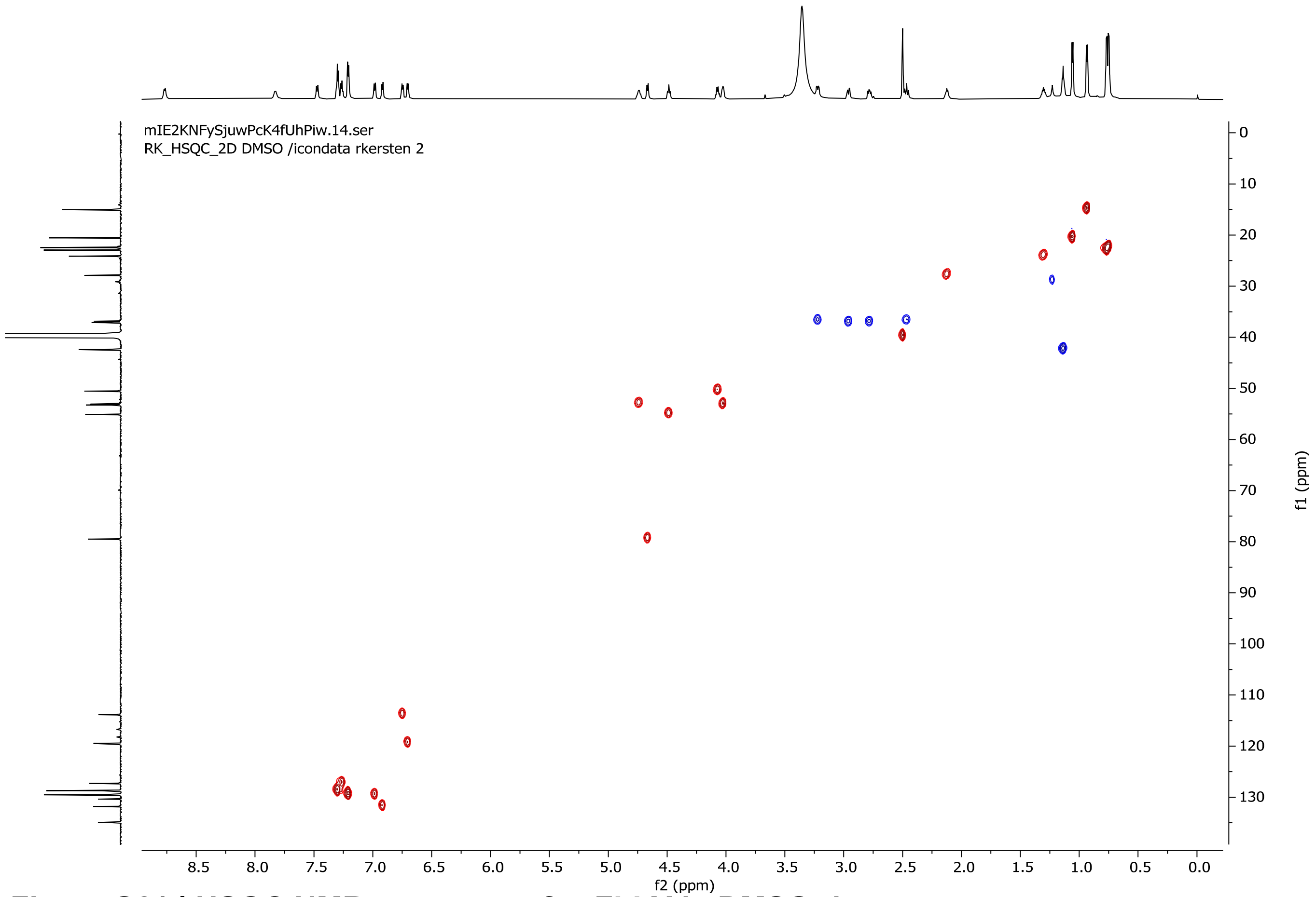


**Figure S61 | HSQC NMR spectrum of cyFLLY in DMSO-d$_6$.**

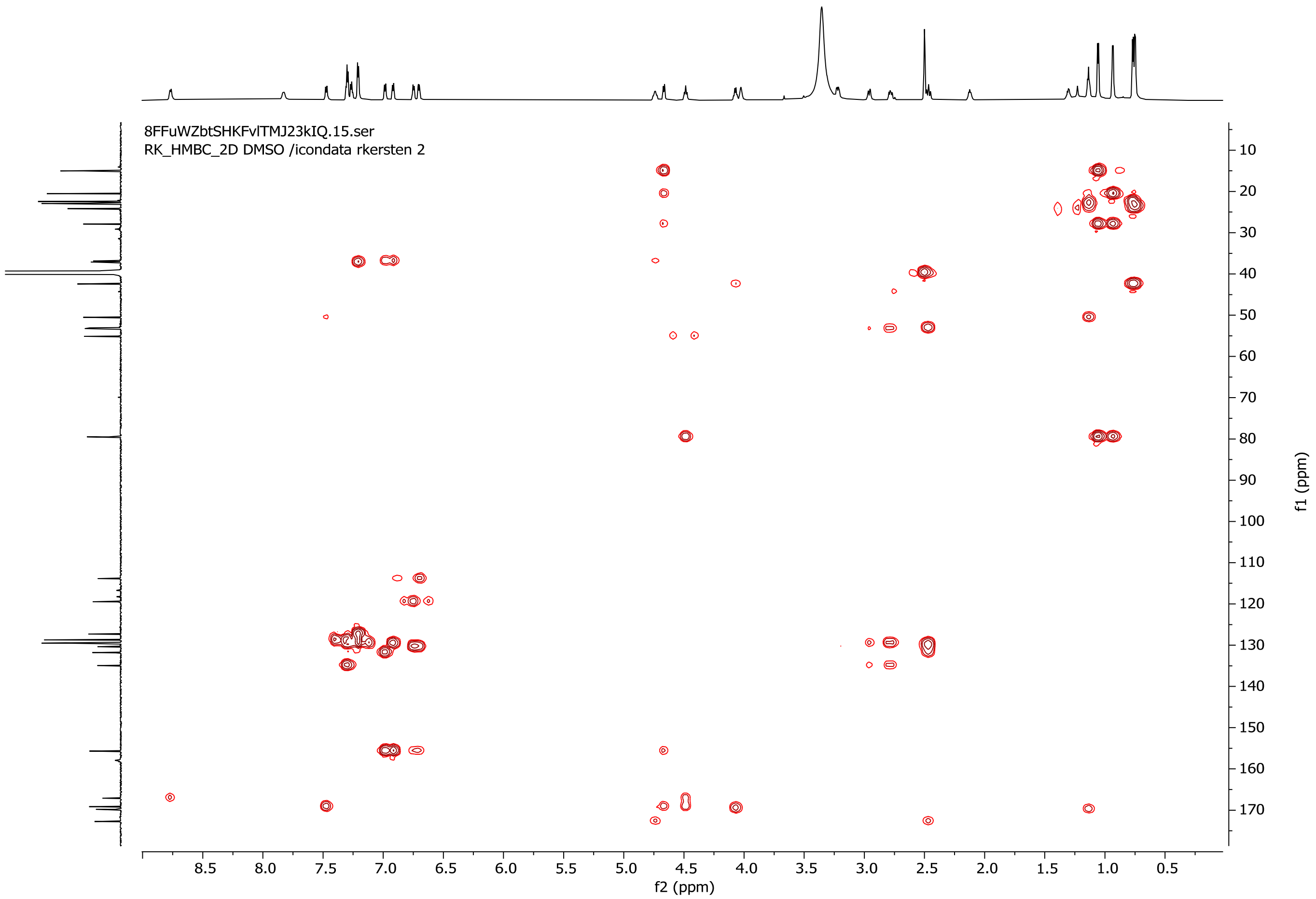


**Figure S62 | HMBC NMR spectrum of cyFLLY in DMSO-d$_6$.**

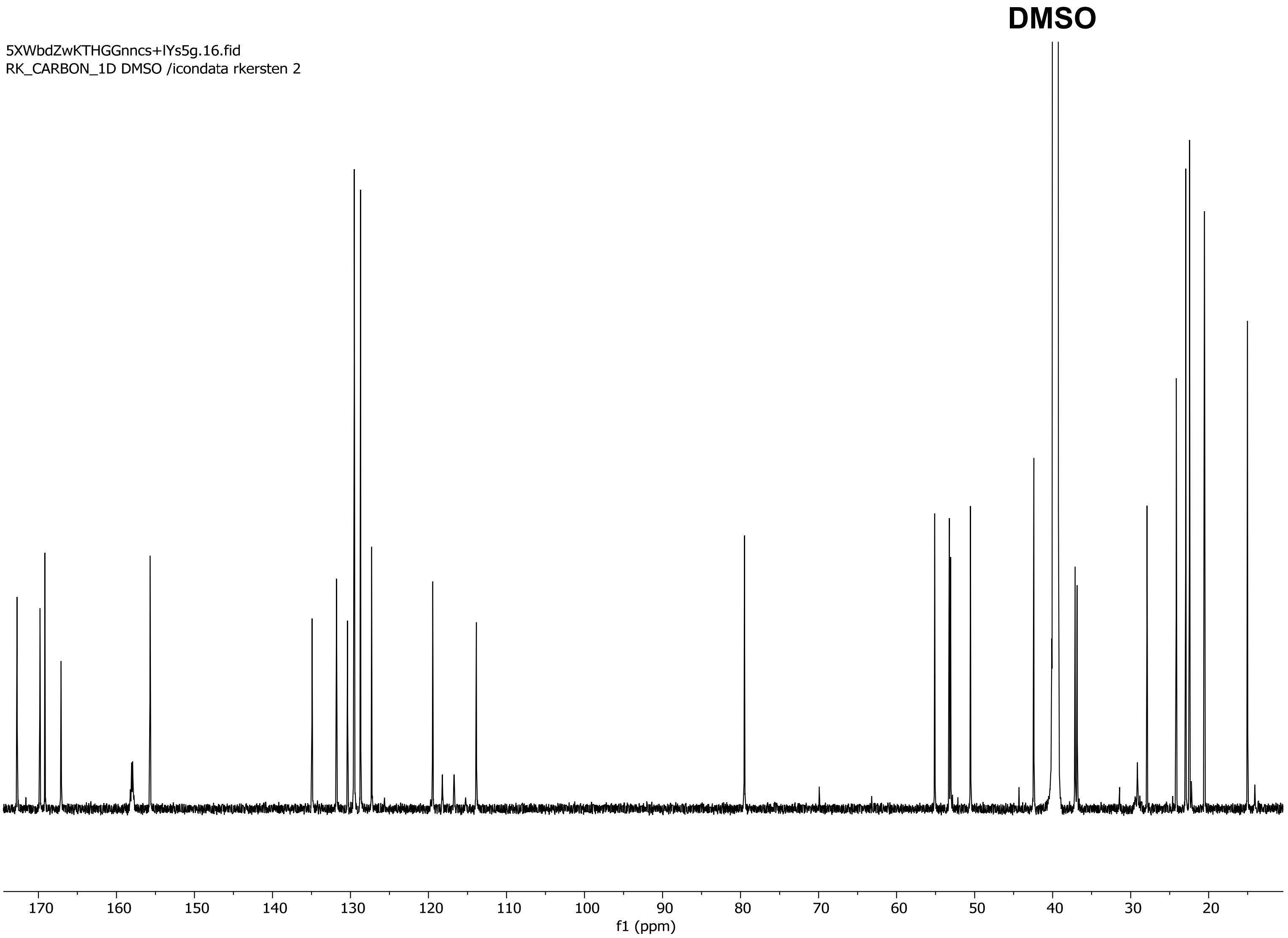


**Figure S63 | $^{13}$C NMR spectrum of cyFLLY in DMSO-$d_6$ (200MHz).**

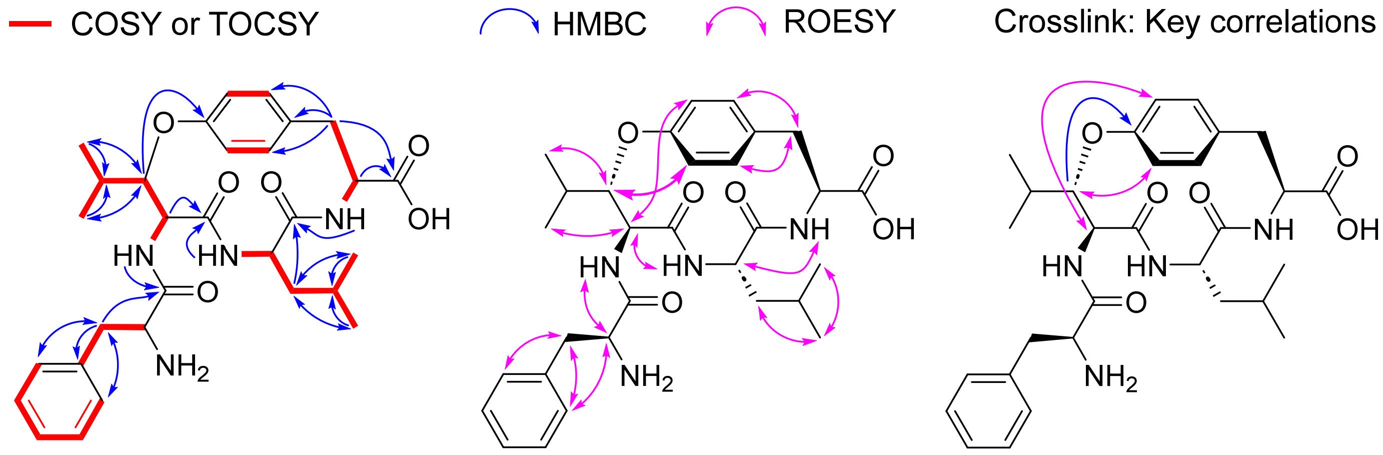


**Figure S64 | Key 2D NMR correlations of cyFLLY in DMSO-$d_6$.**

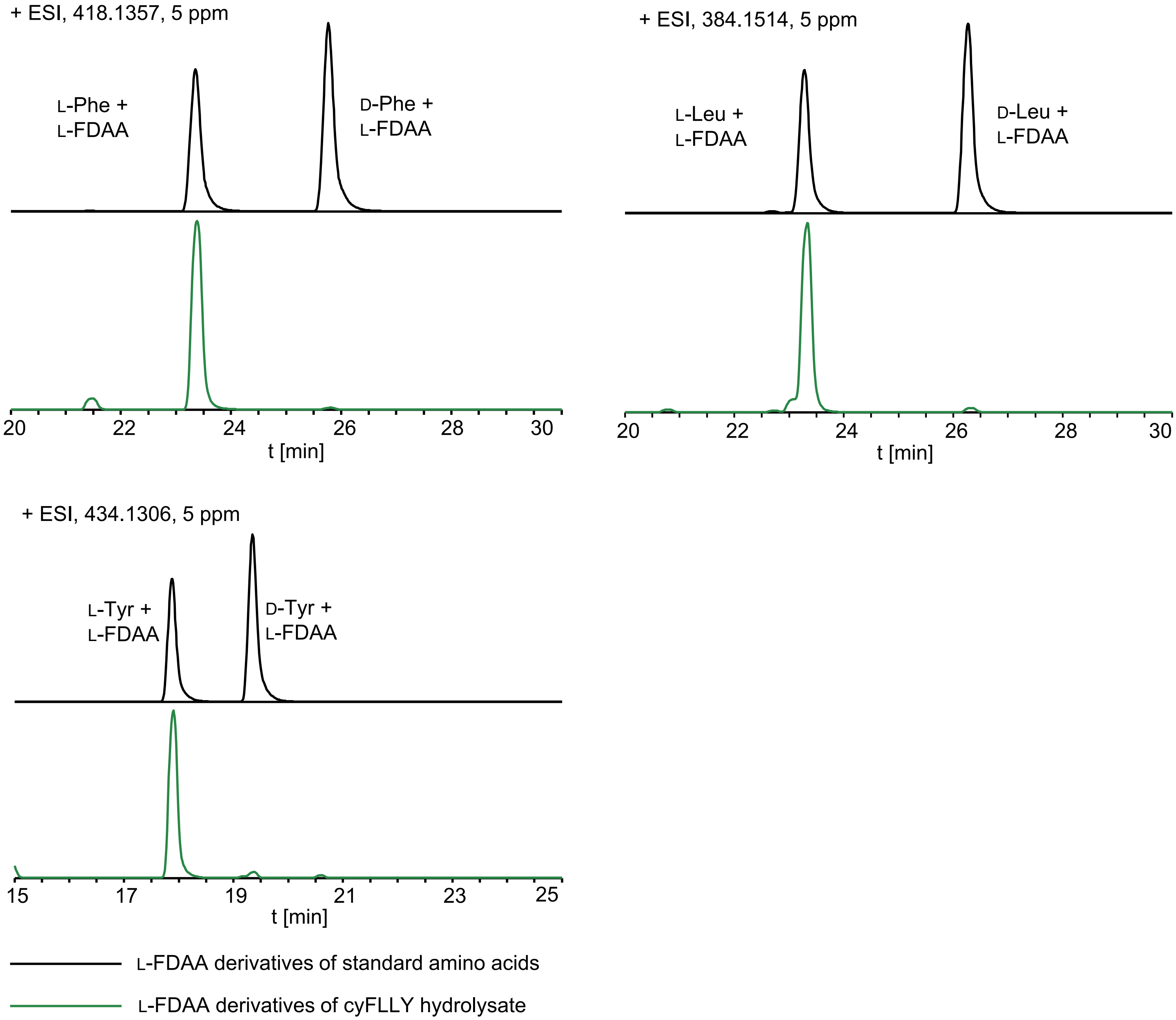


**Figure S65 | Stereochemical analysis of cyFLLY.** (**A**) The Phe, Leu and Tyr residues in cyFLLY were assigned as L by using Marfey’s analysis.

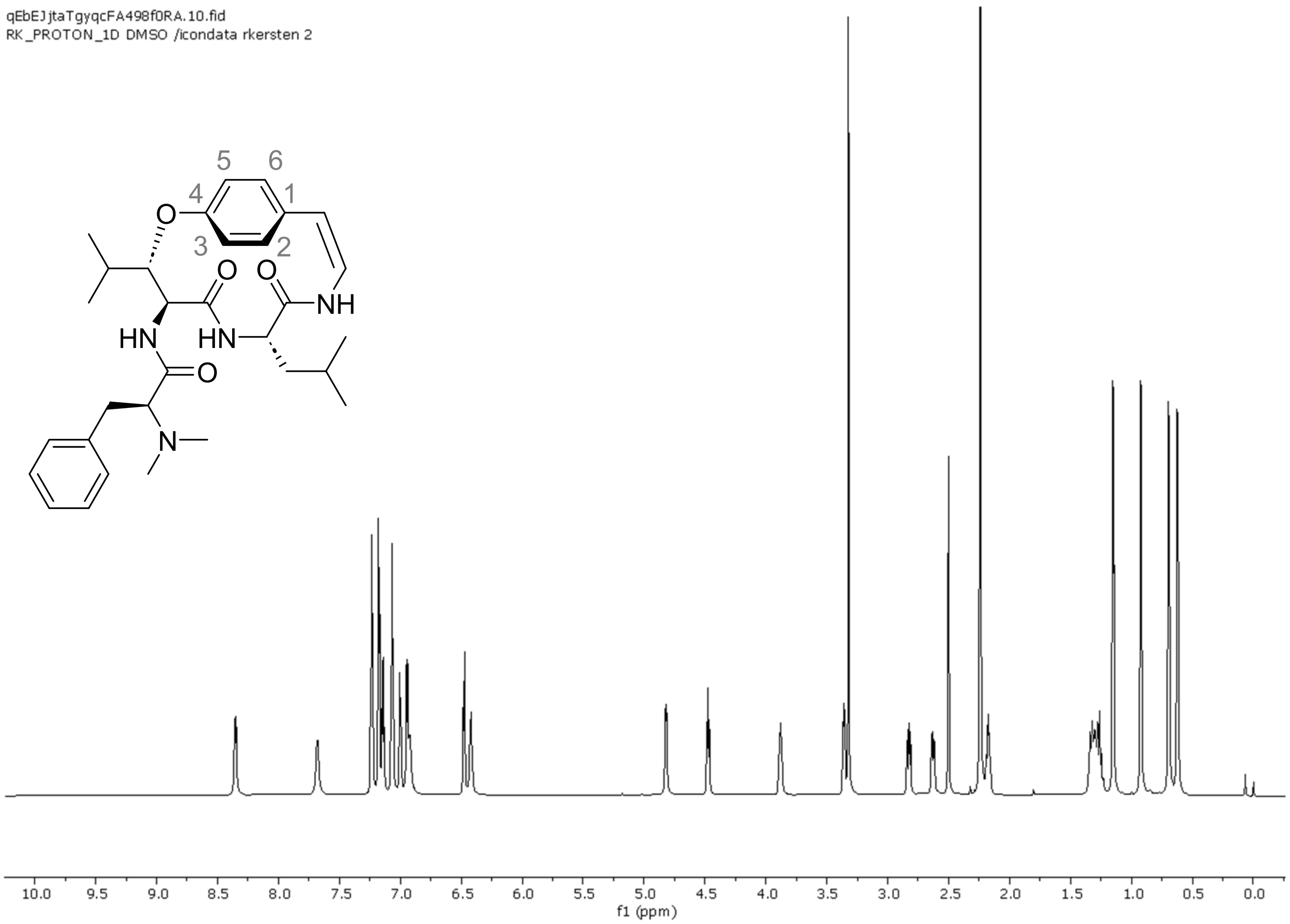


**Figure S66 | $^{1}$H NMR spectrum of sanjoinine A authentic standard in DMSO-$d_6$ (800MHz).**

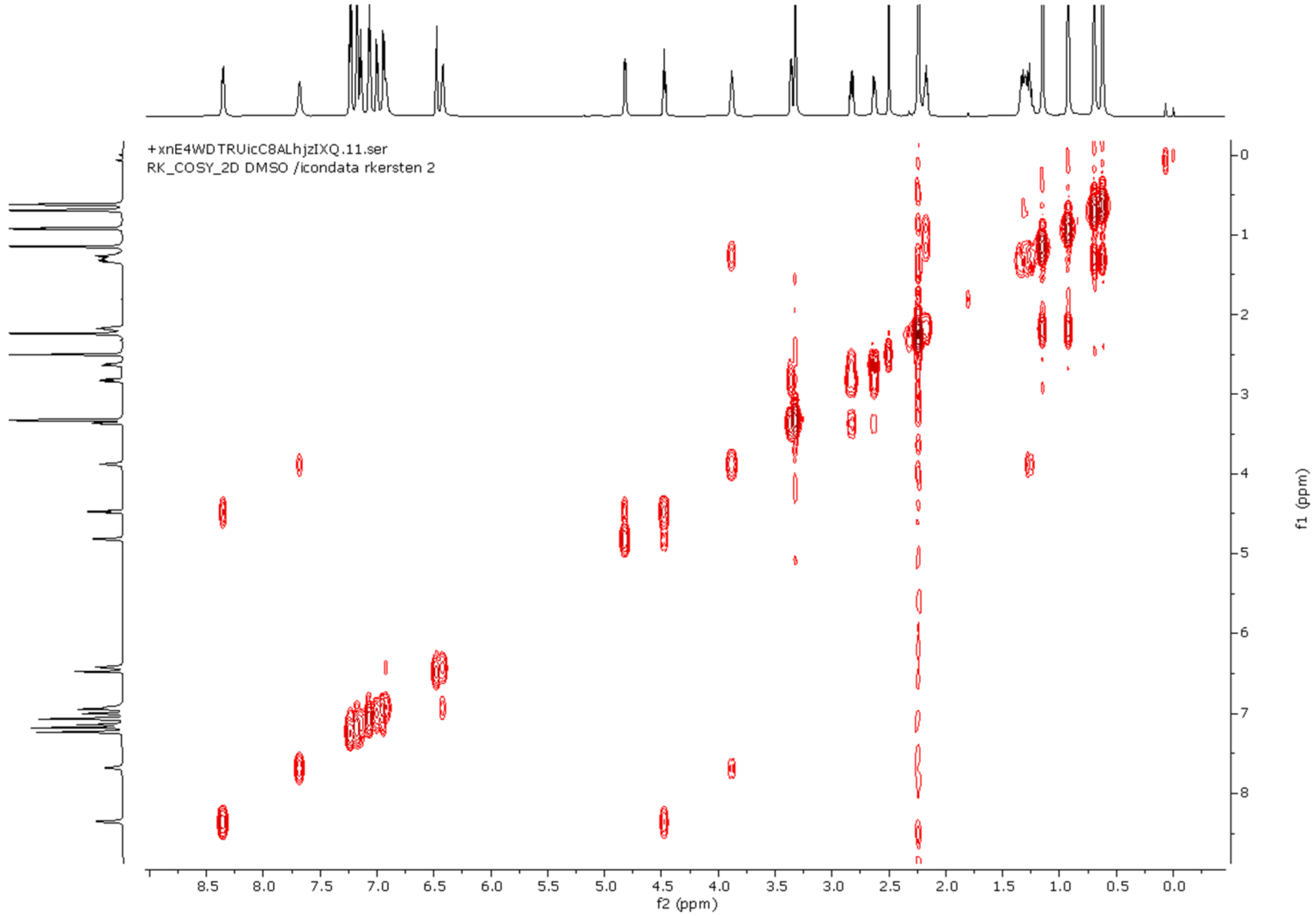


**Figure S67 | $^{1}$H-$^{1}$H COSY NMR spectrum of sanjoinine A authentic standard in DMSO-$d_6$.**

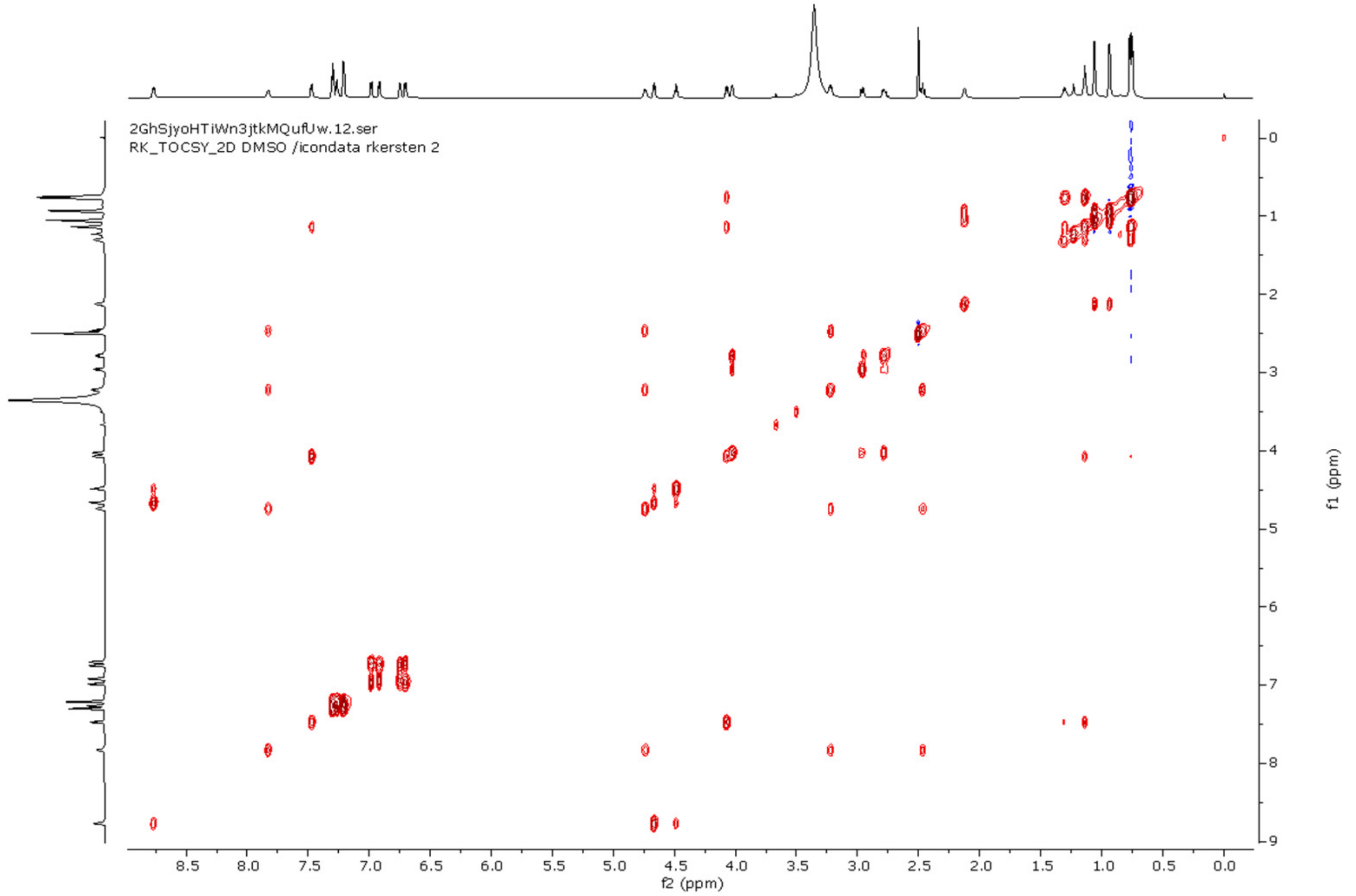


**Figure S68 | $^{1}H$-$^{1}H$ TOCSY NMR spectrum of sanjoinine A authentic standard in DMSO-$d_6$.**

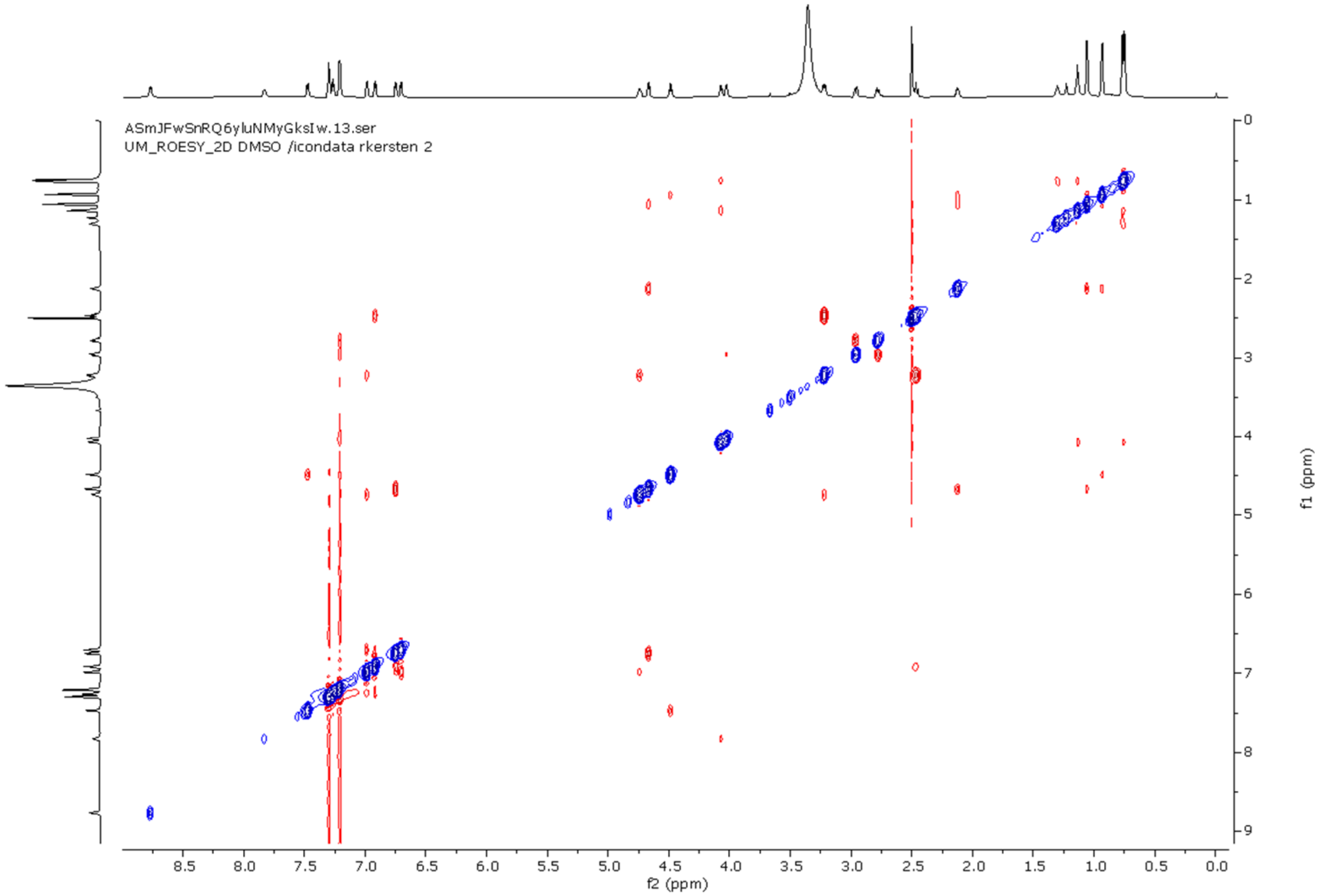


**Figure S69 | ROESY NMR spectrum of sanjoinine A authentic standard in DMSO-$d_6$.**

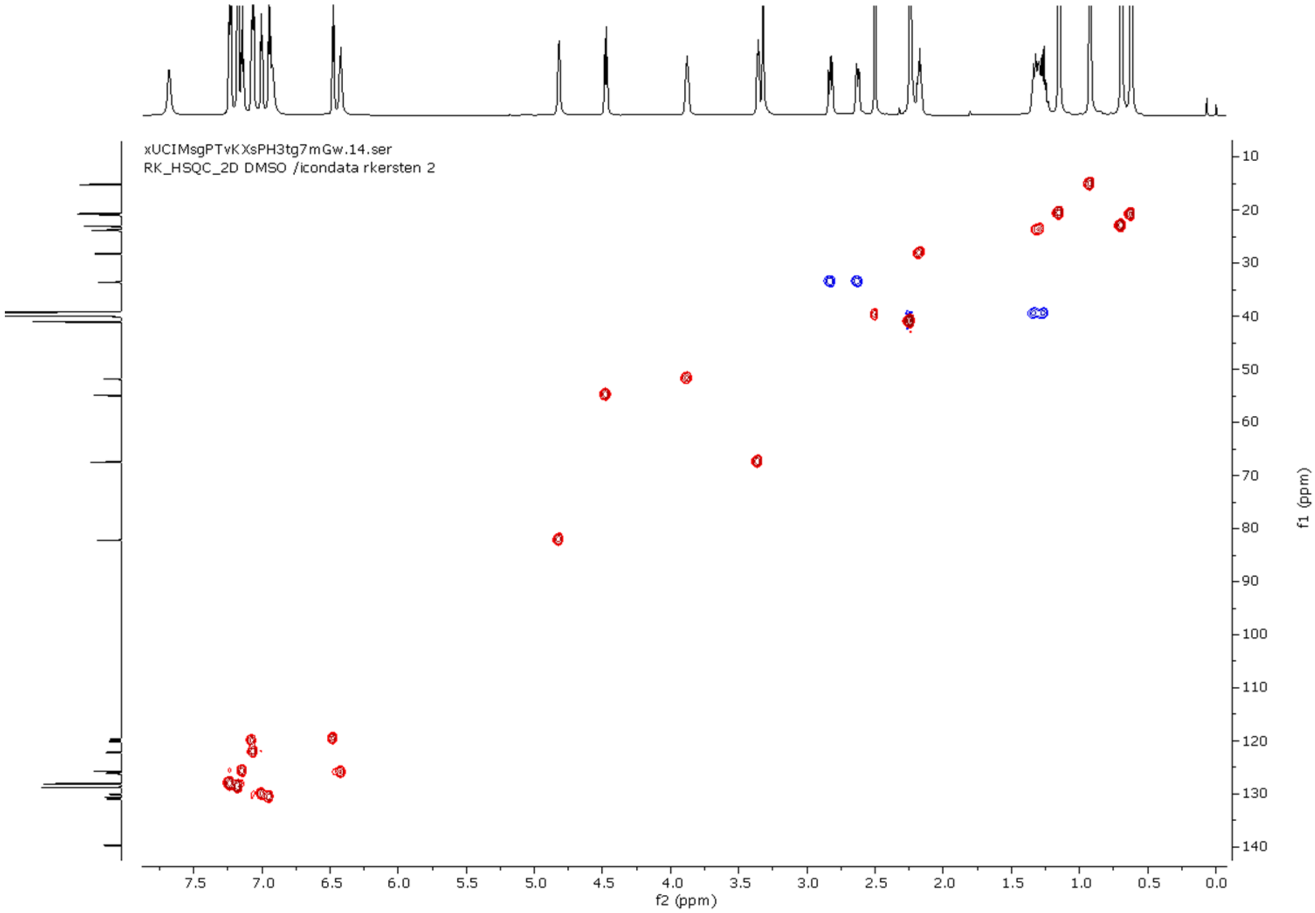


**Figure S70 | HSQC NMR spectrum of sanjoinine A authentic standard in DMSO-d$_6$.**

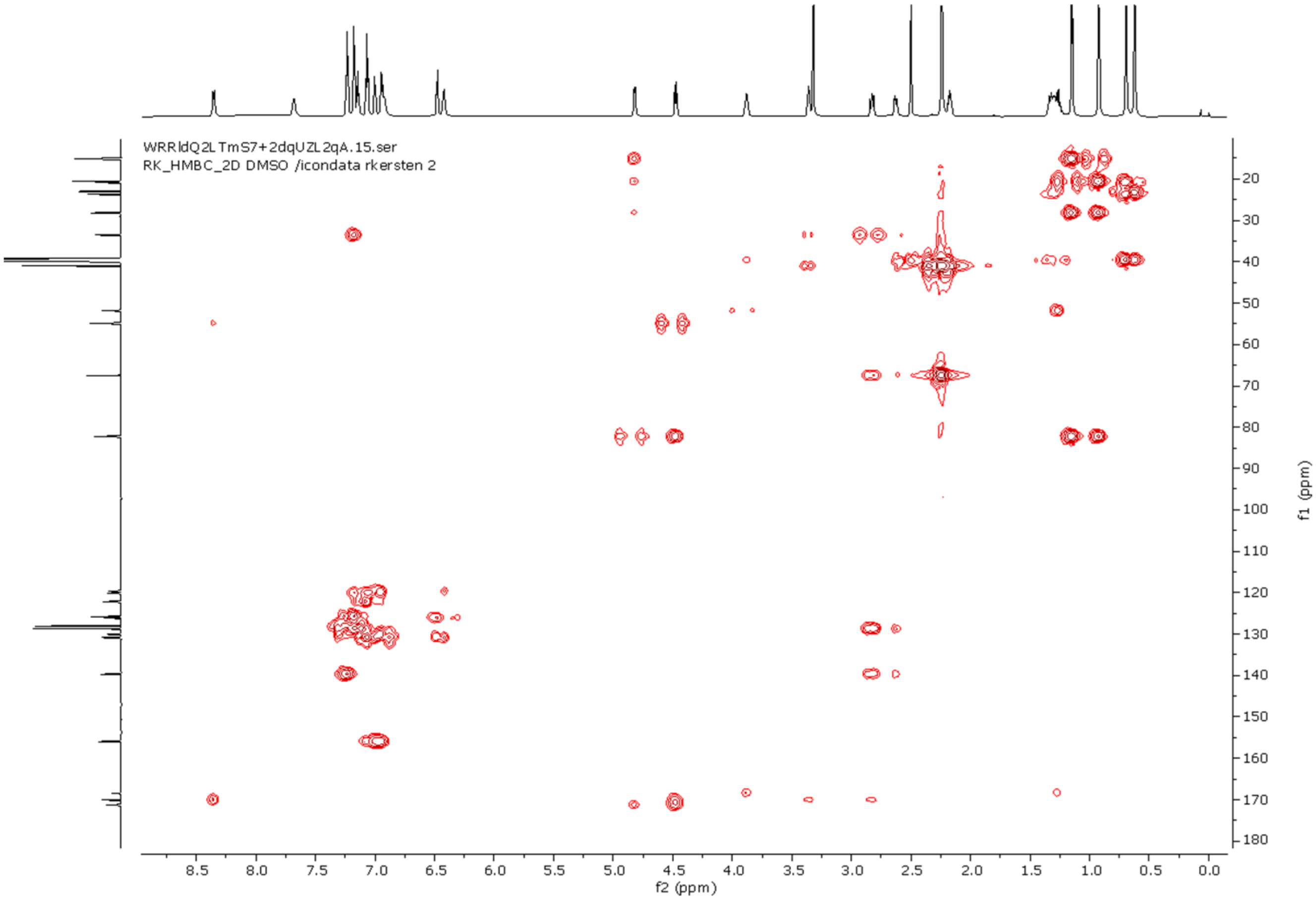


**Figure S71 | HMBC NMR spectrum of sanjoinine A authentic standard in DMSO-d$_6$.**

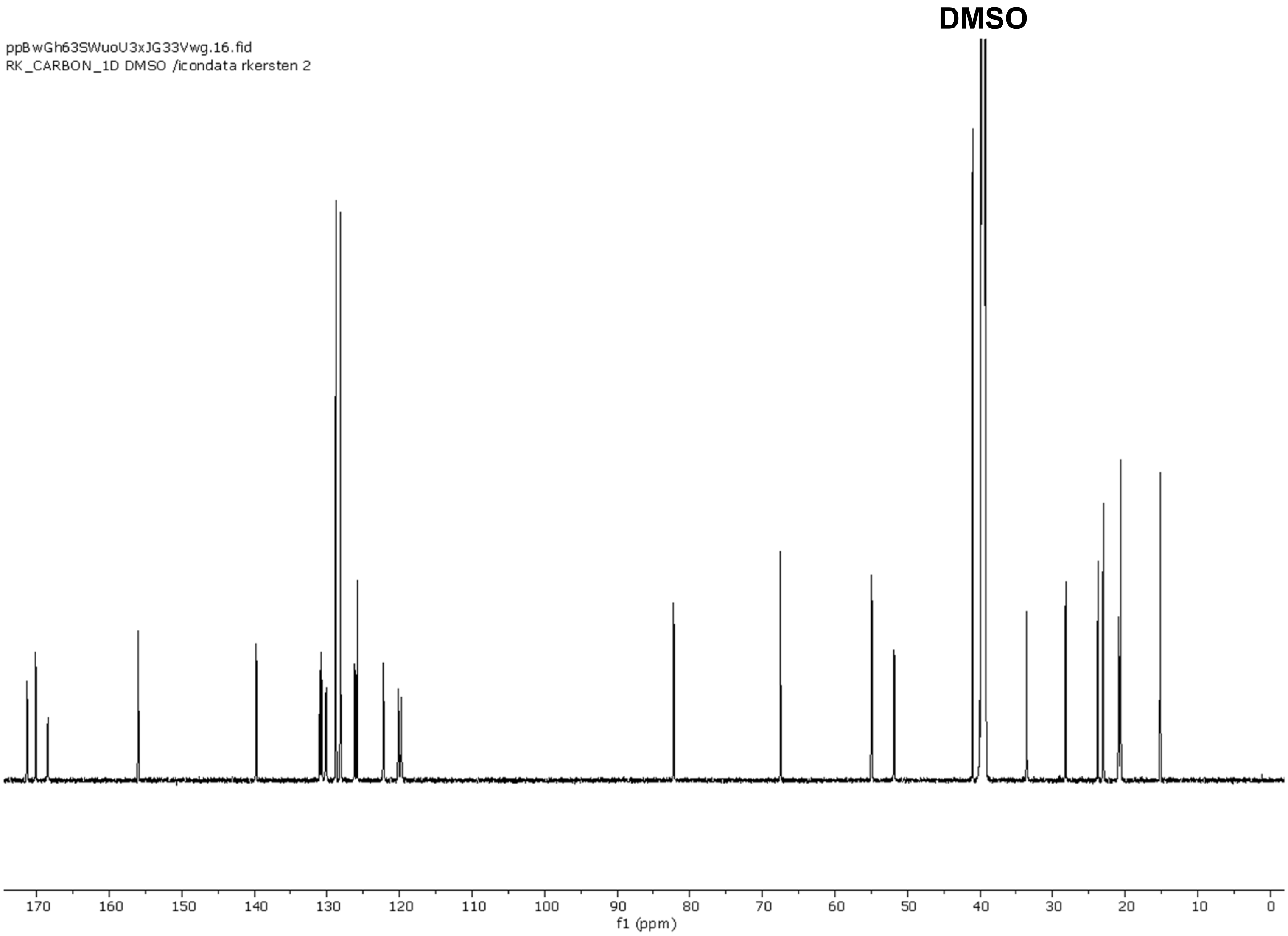


**Figure S72 | $^{13}$C NMR spectrum of sanjoinine A authentic standard in DMSO-$d_6$ (200 MHz).**

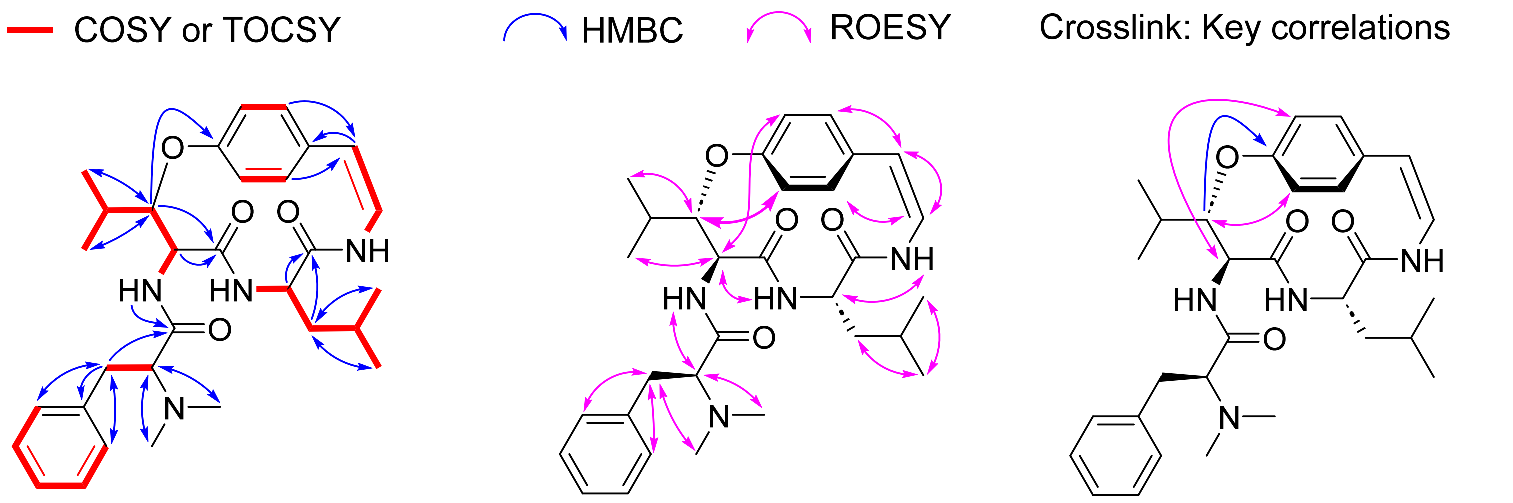


**Figure S73 | Key 2D NMR correlations of sanjoinine A authentic standard in DMSO-$d_6$.**

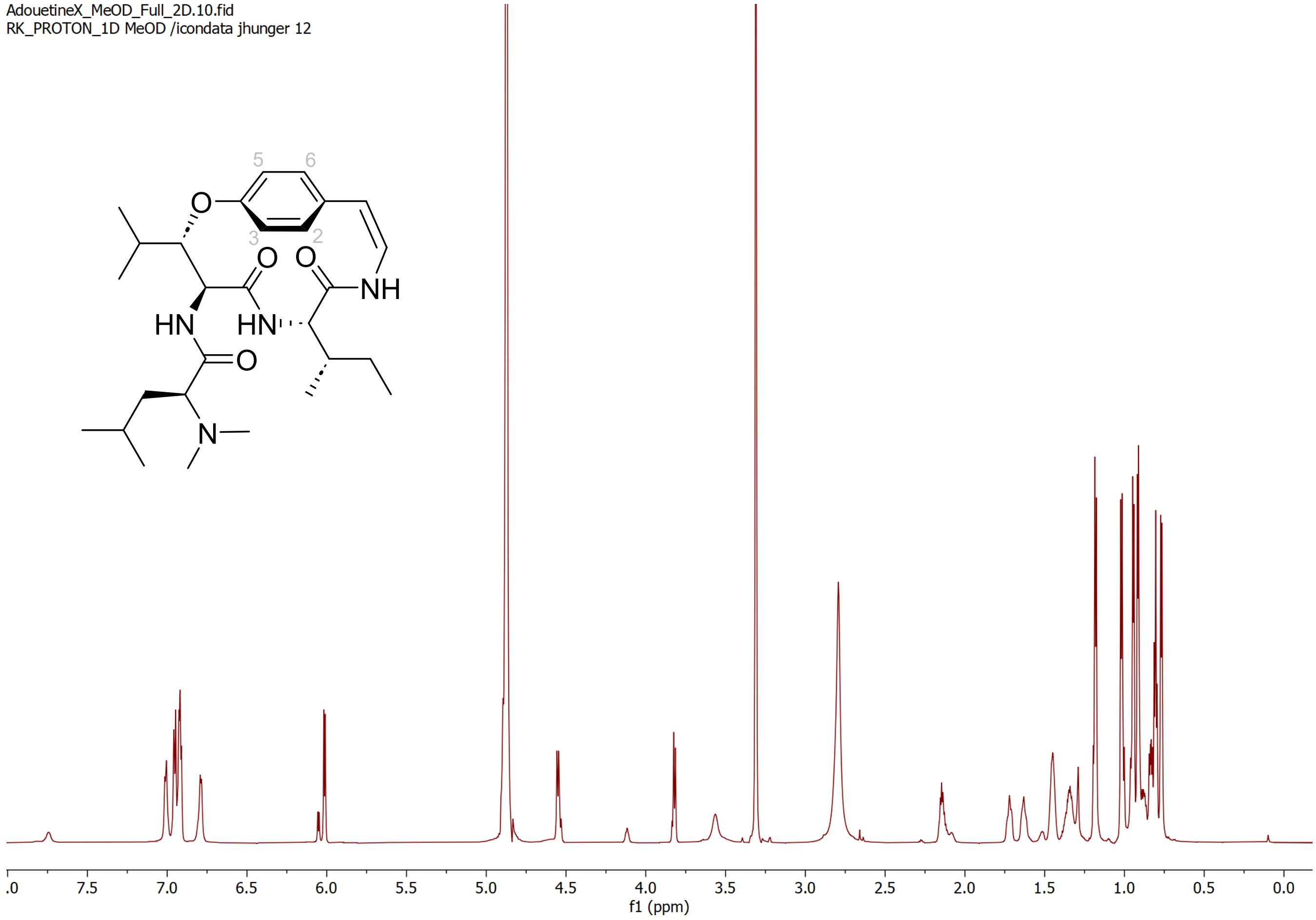


**Figure S74 | $^1$H NMR spectrum of adouetine X in $CD_3OD$ (800 MHz).**

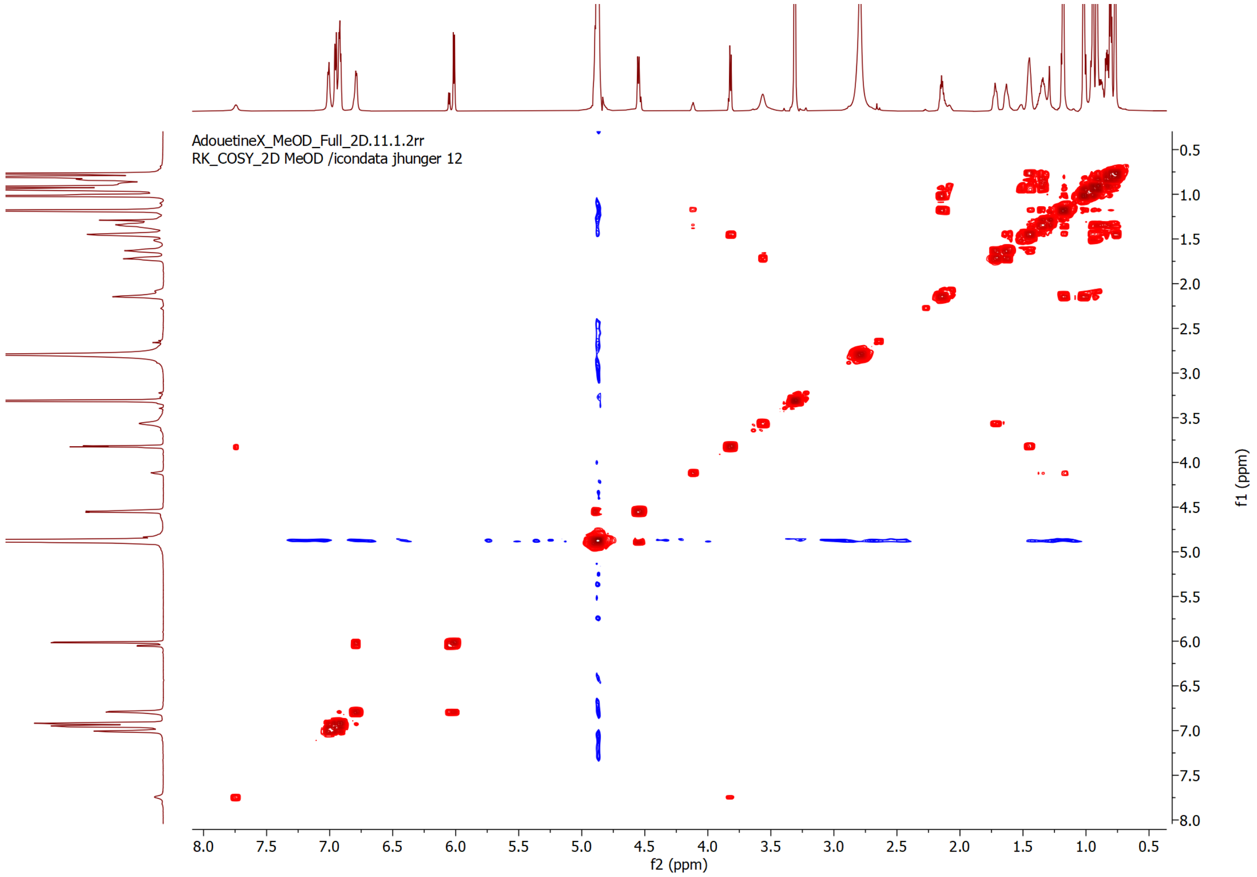


**Figure S75 | $^1$H-$^1$H COSY NMR spectrum of adouetine X in $CD_3OD$.**

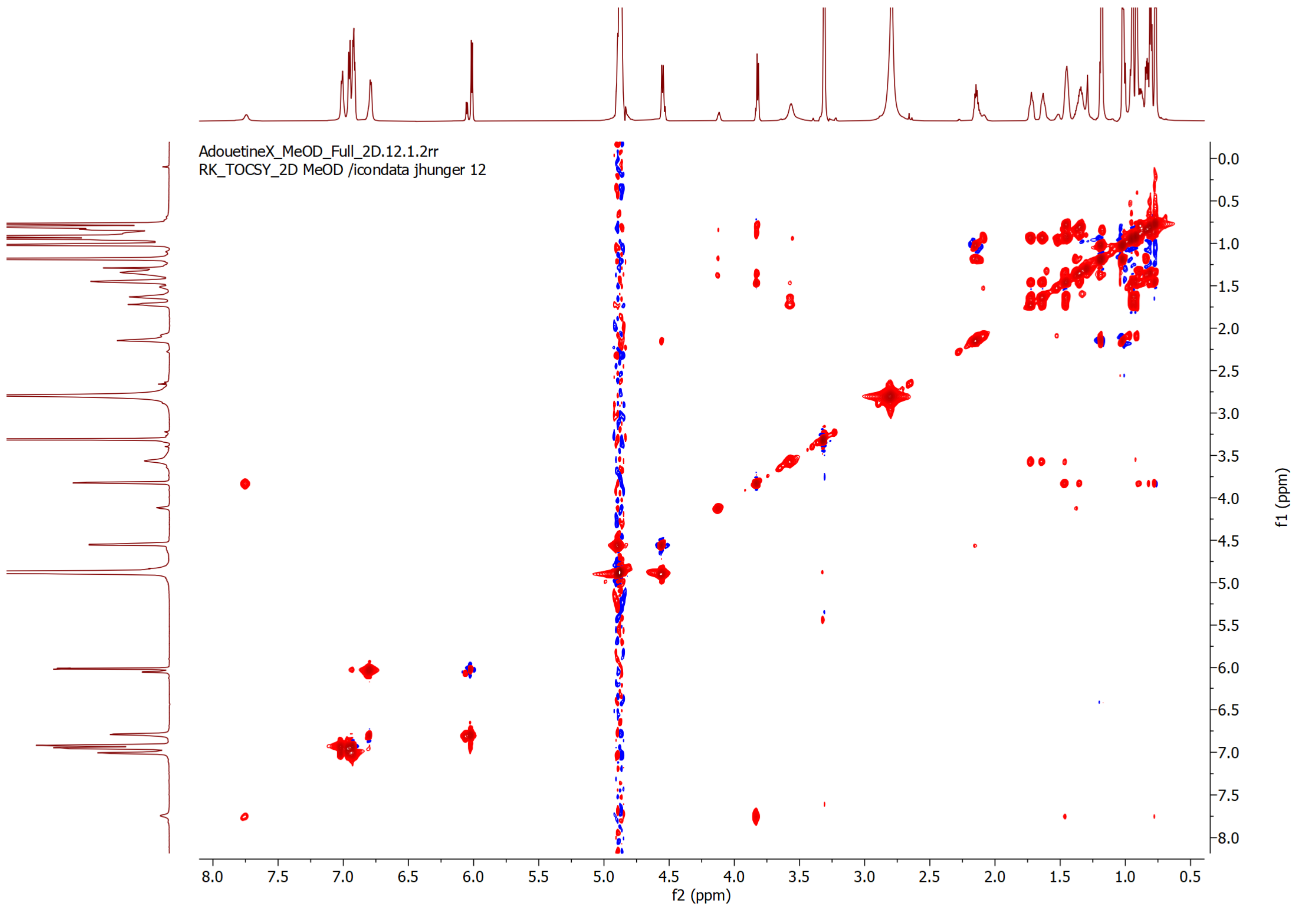


**Figure S76 | $^{1}H$-$^{1}H$ TOCSY NMR spectrum of adouetine X in $CD_3OD$.**

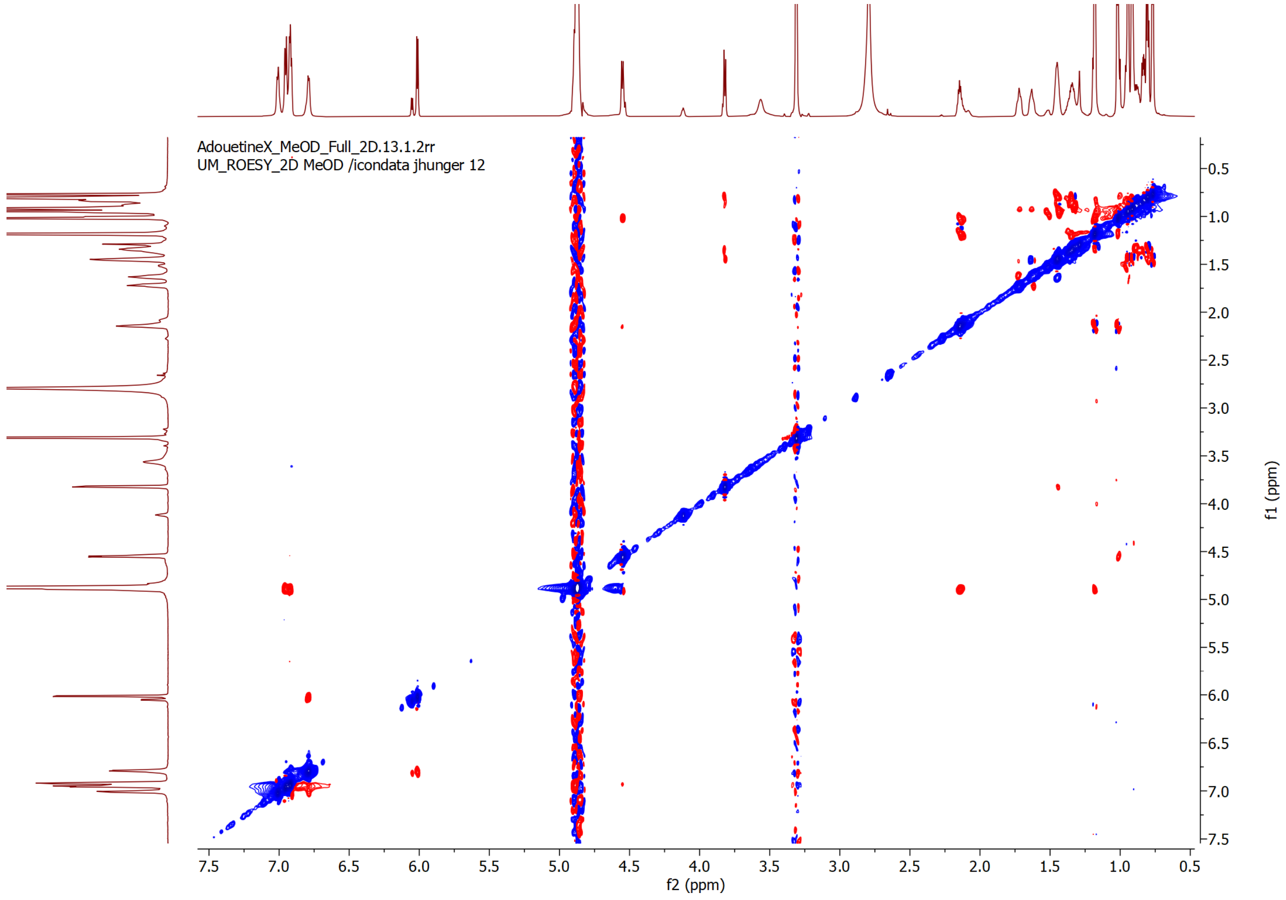


**Figure S77 | ROESY NMR spectrum of adouetine X in $CD_3OD$.**

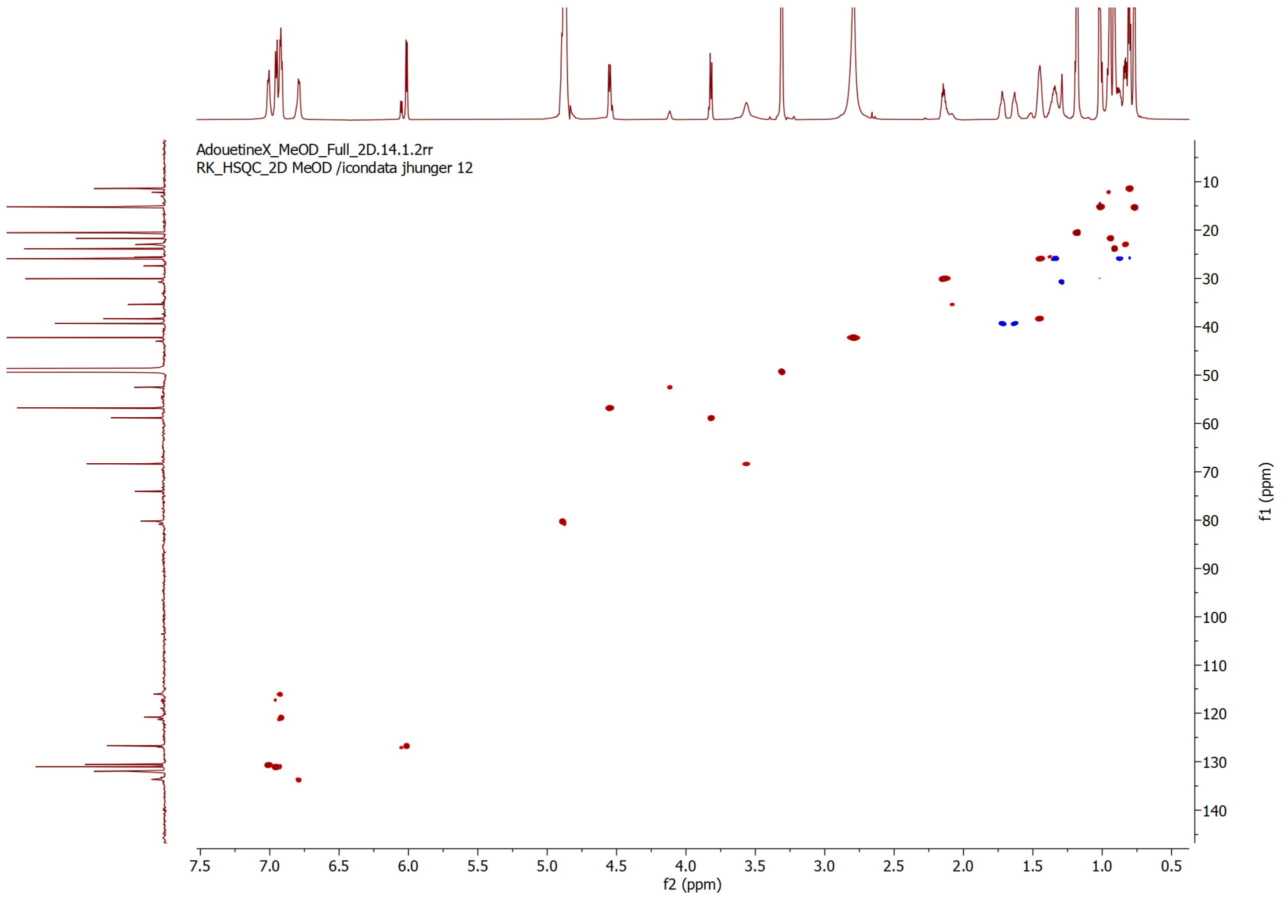


**Figure S78 | HSQC NMR spectrum of adouetine X in $CD_3OD$.**

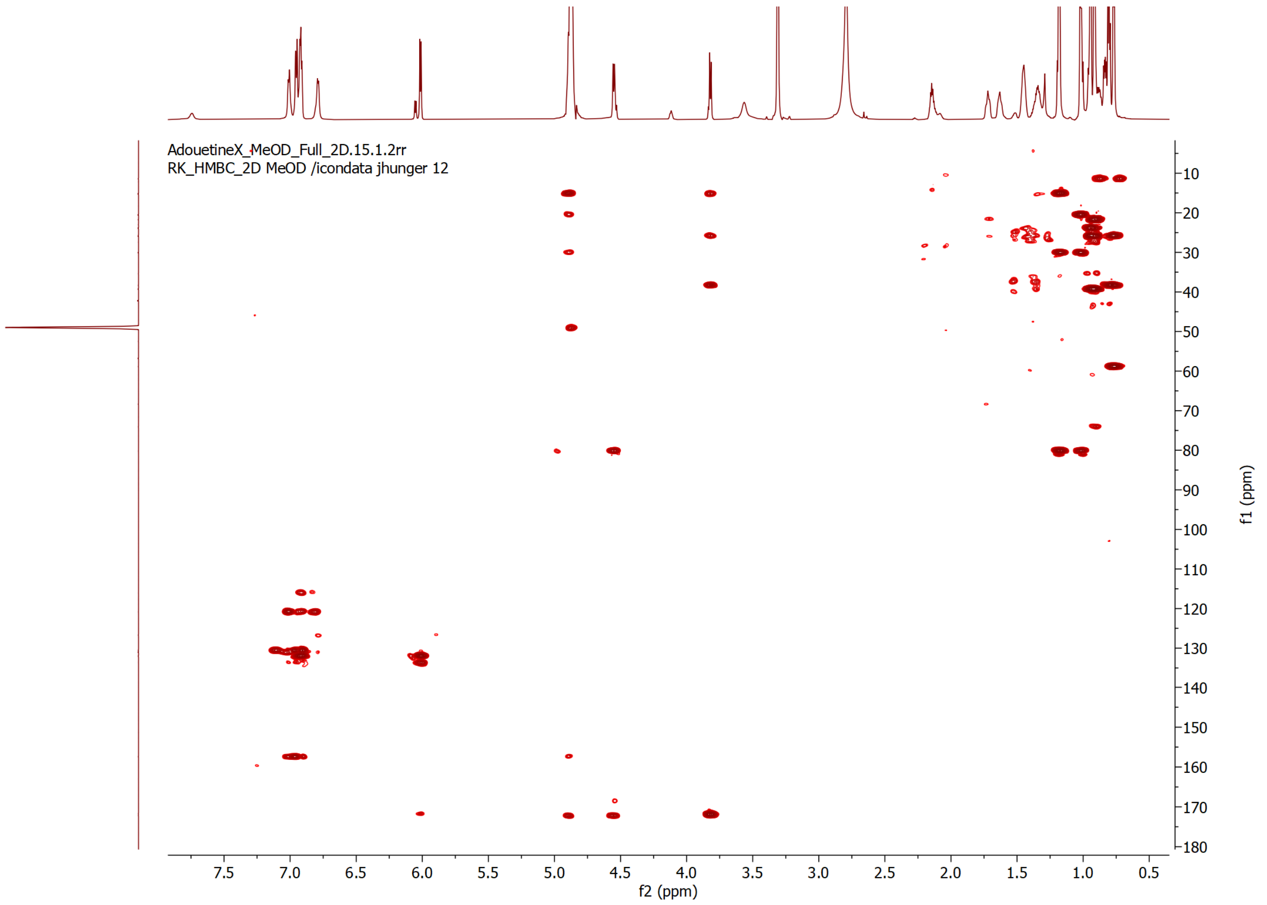


**Figure S79 | HMBC NMR spectrum of adouetine X in $CD_3OD$.**

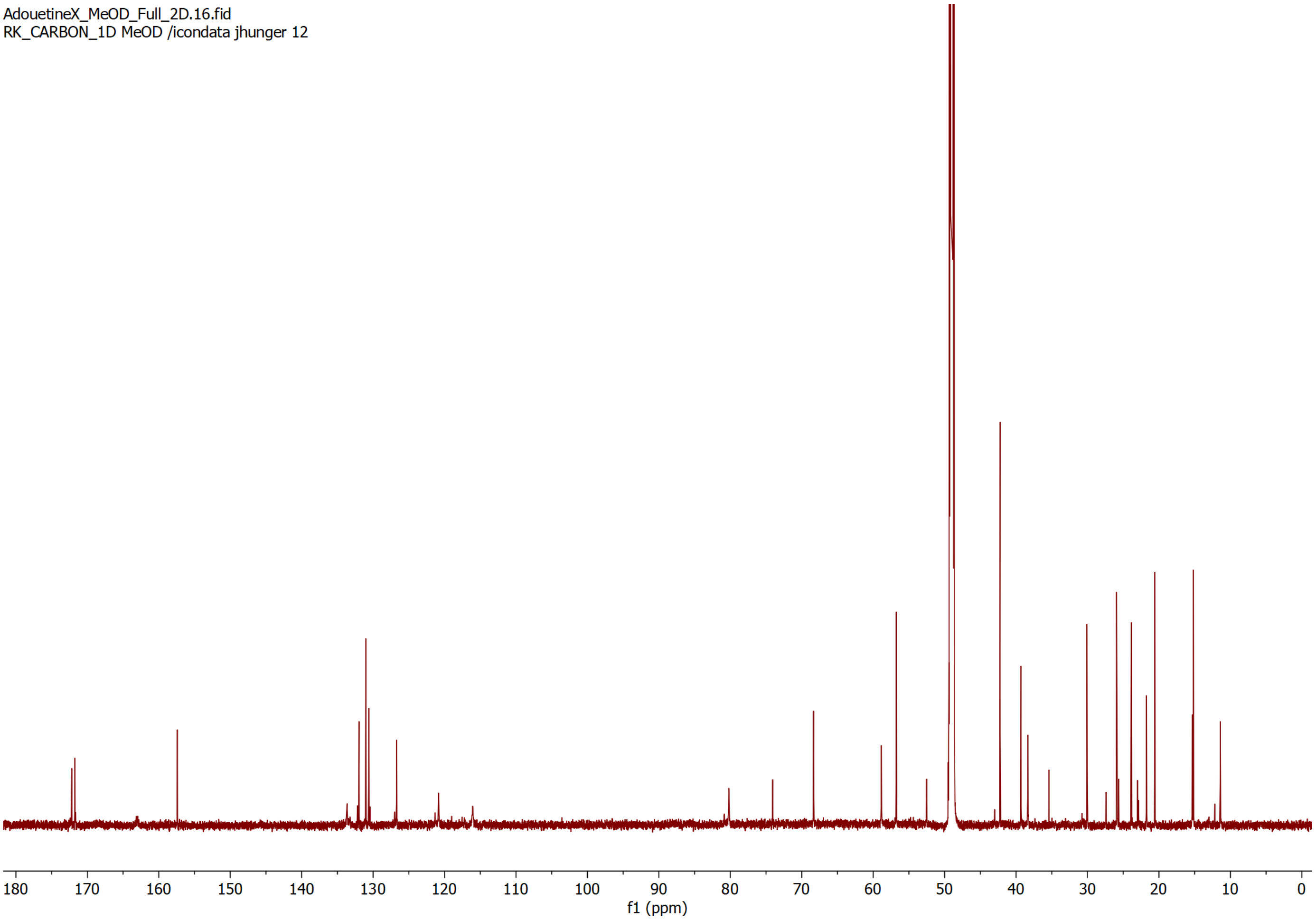


**Figure S80 | $^{13}C$ NMR spectrum of adouetine X in $CD_3OD$ (200MHz).**

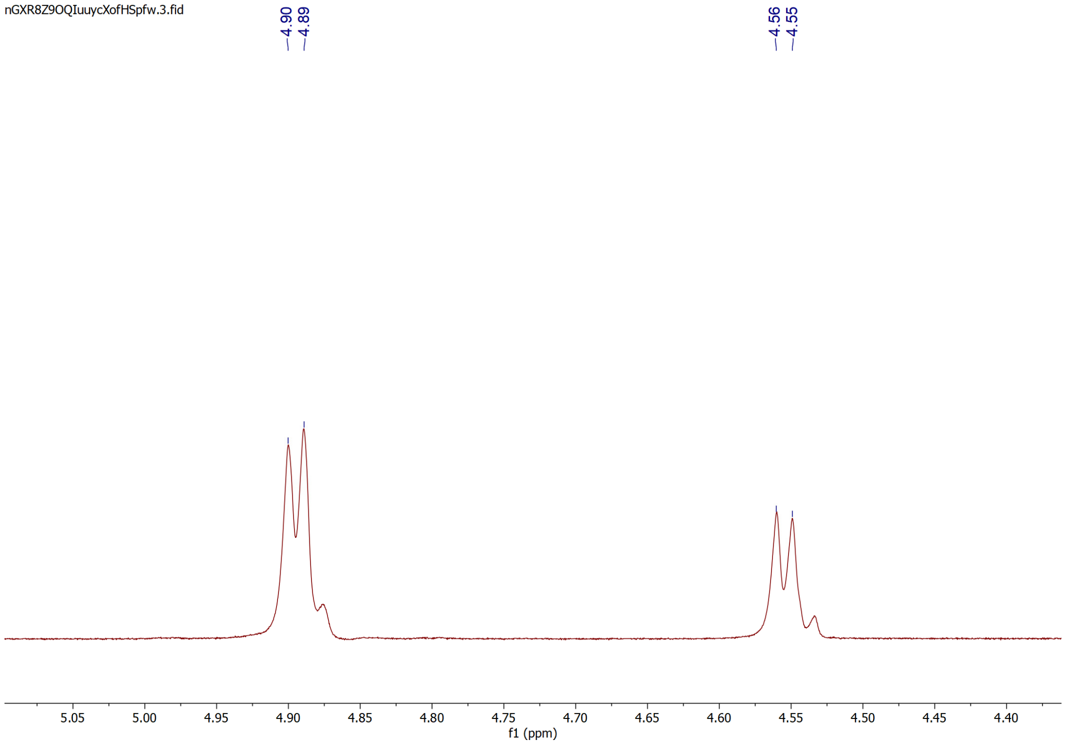


**Figure S81 | Selective 1D-TOCSY NMR spectrum of adouetine X in $CD_3OD$.**

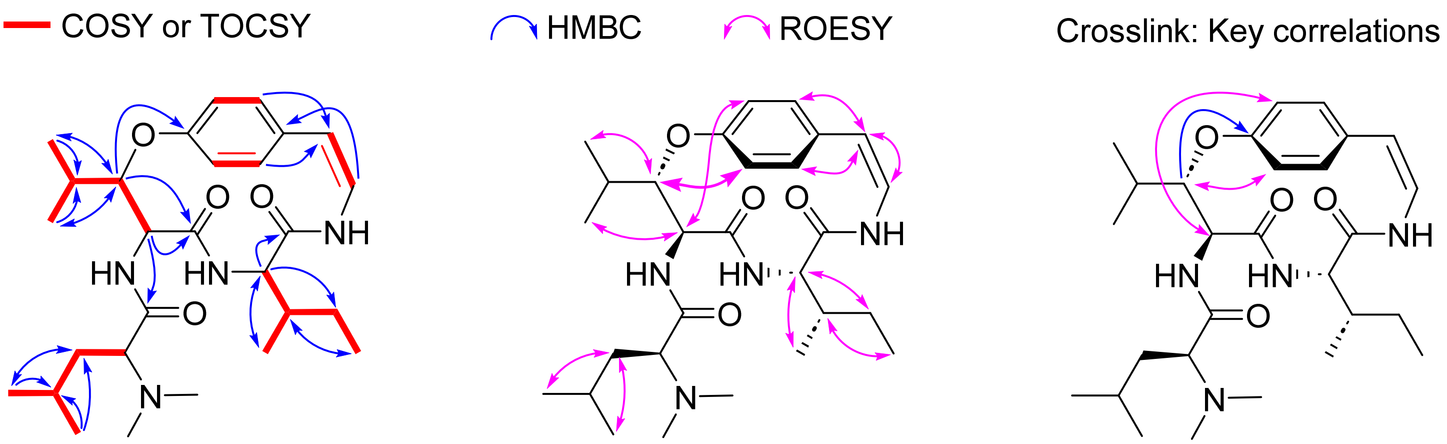


**Figure S82 | Key 2D NMR correlations of adouetine X in $CD_3OD$.**

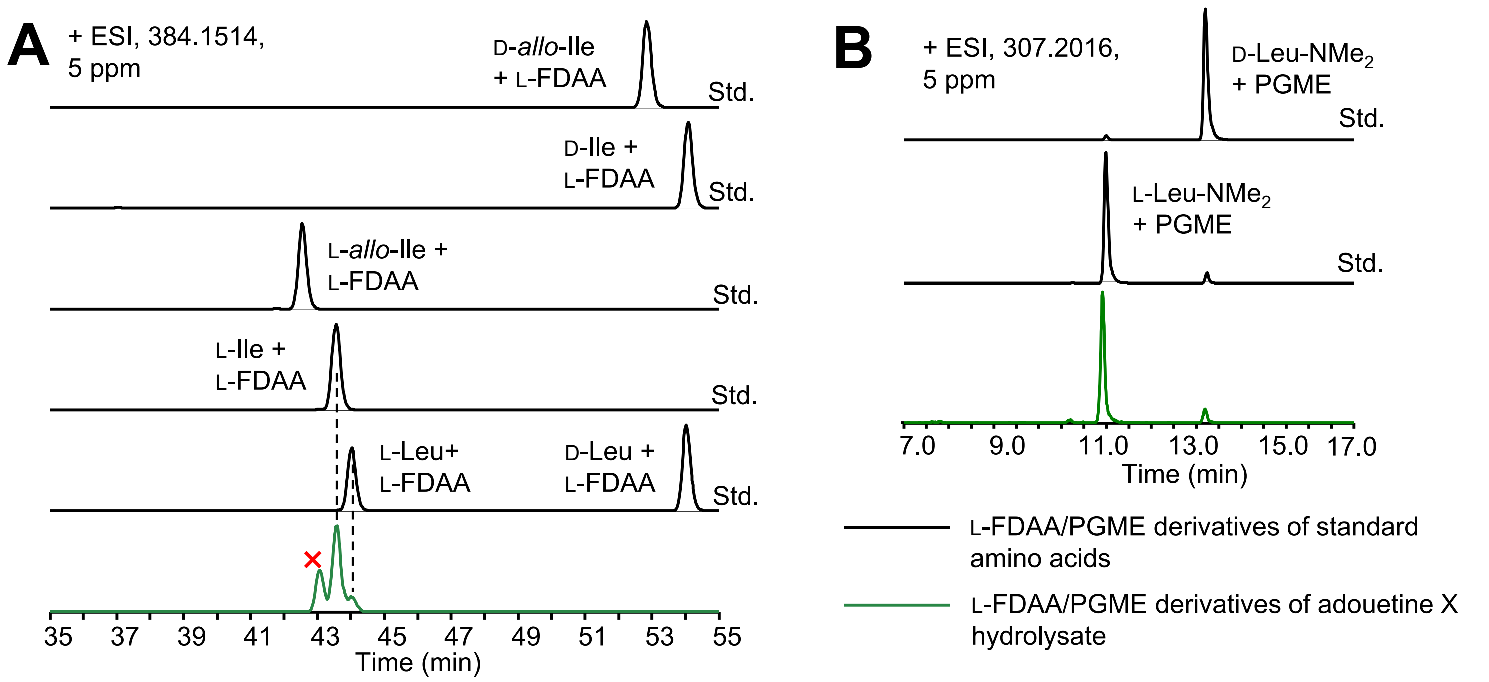


**Figure S83 | Stereochemical analysis of adouetine X** (**A**) The Leu and Ile residues in adouetine X were assigned as L by C3-Marfey's analysis. (**B**) The LeuNMe$_2$ residue in adouetine X was assigned as L by PGME derivatization and LC-MS analysis cyFLAY – m(obs) 511.2552, m(calc) 511.2551, Δm = 0.2 ppm

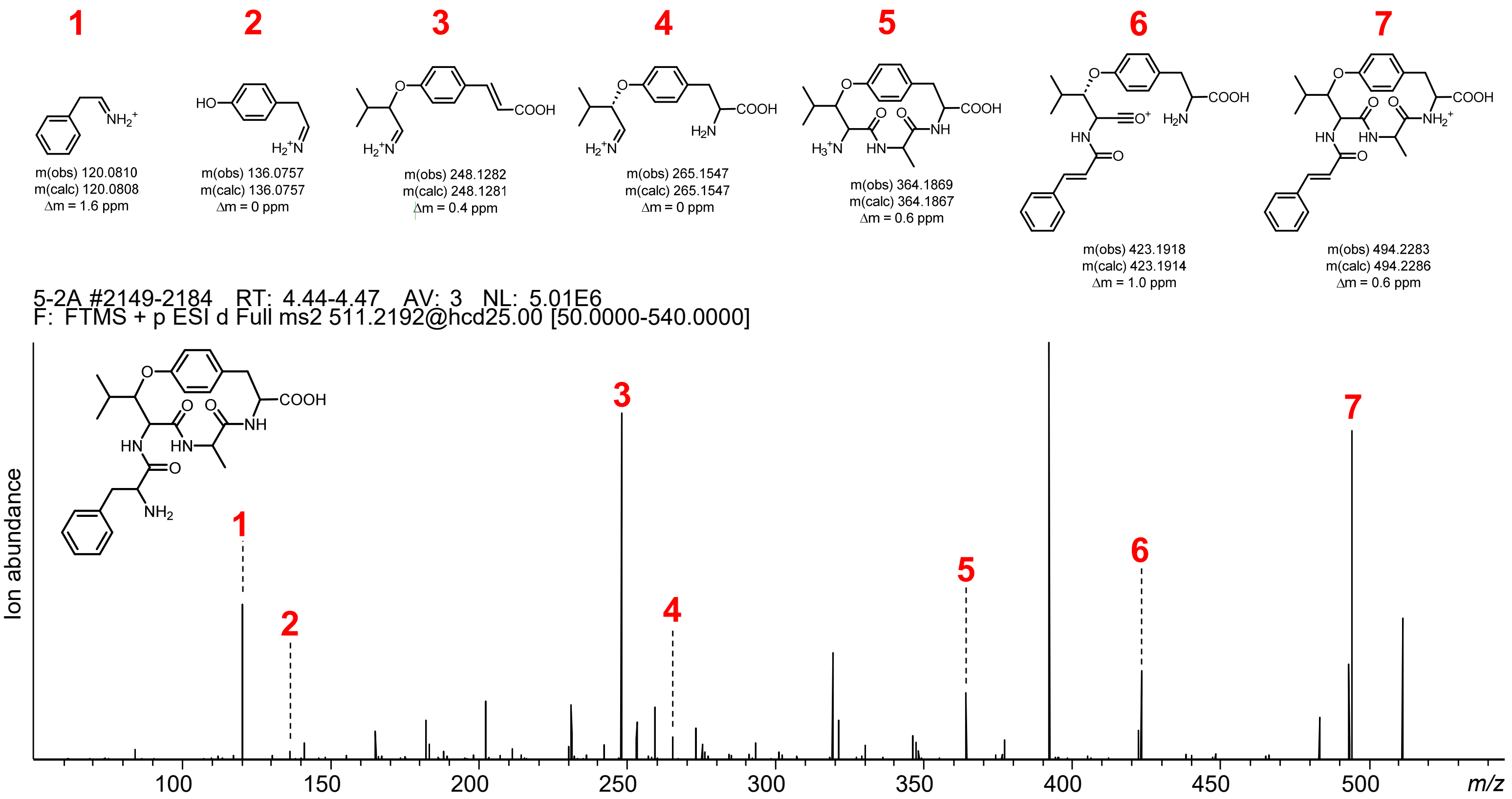


cyFLDY – m(obs) 555.2451, m(calc) 555.2449, Δm = 0.4 ppm

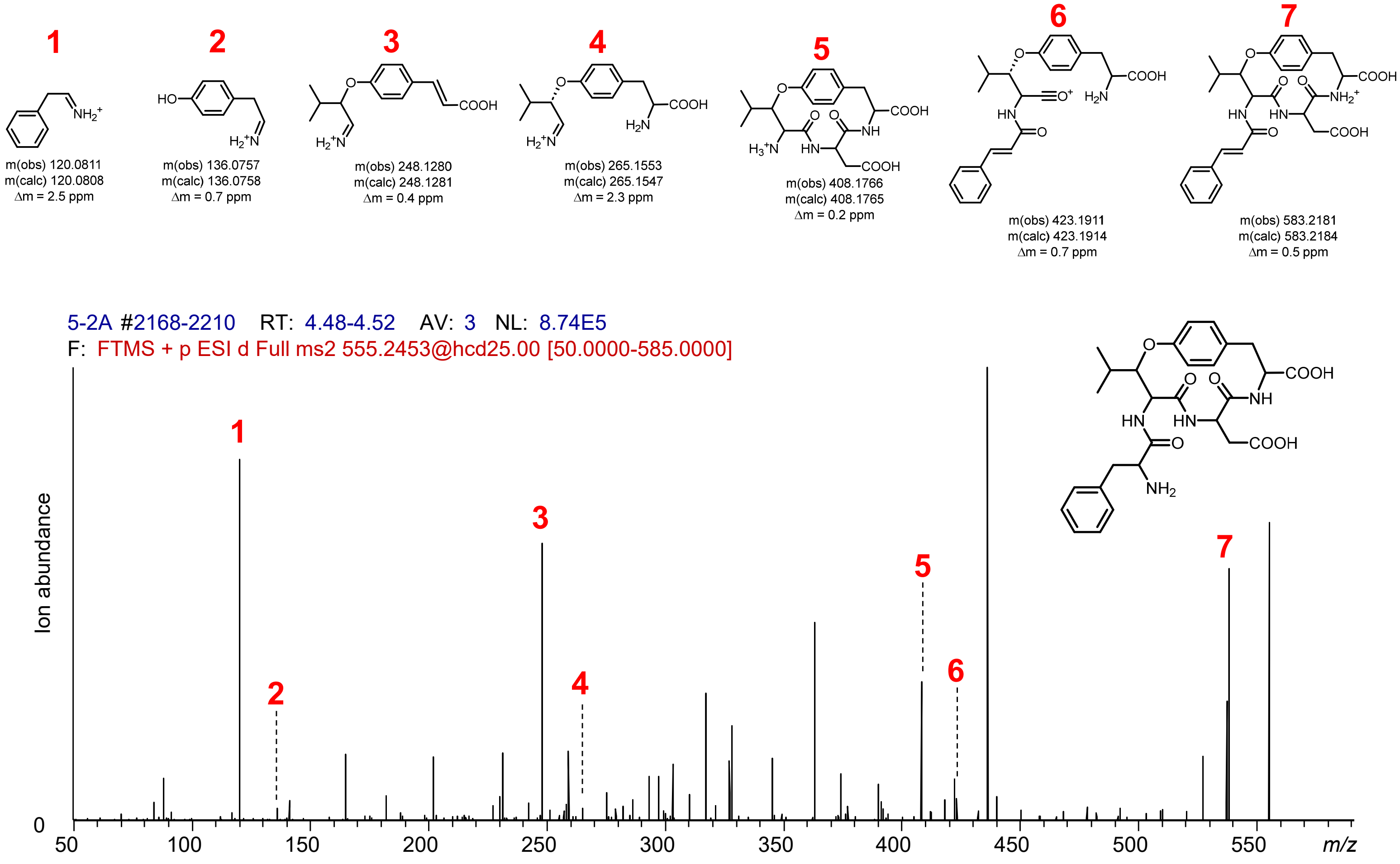


**Figure S84 | cyFLxY diversification via transient expression of SkrBURP-FLxY in *N. benthamiana*.** cyFLAY and cyFLDY

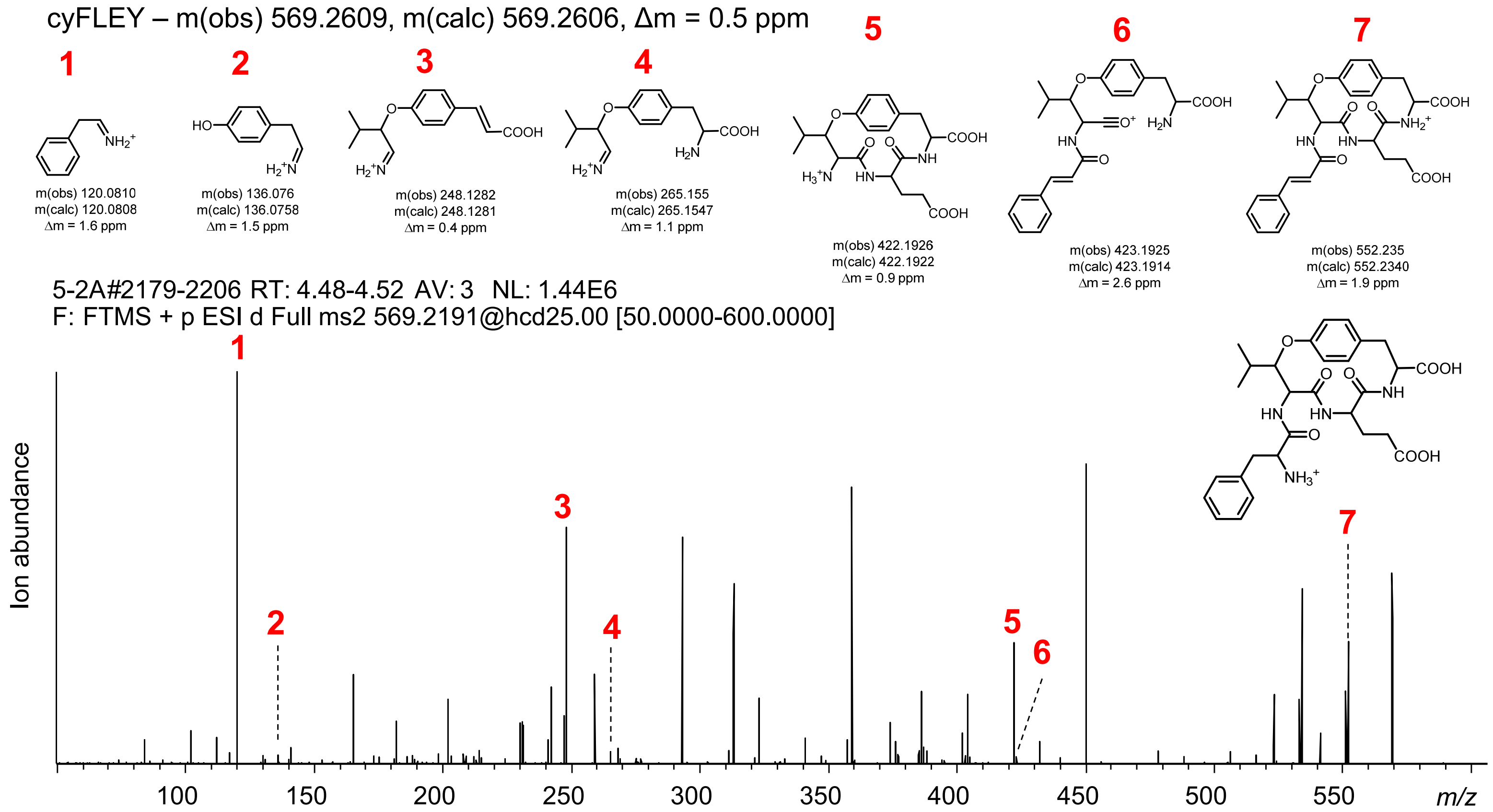


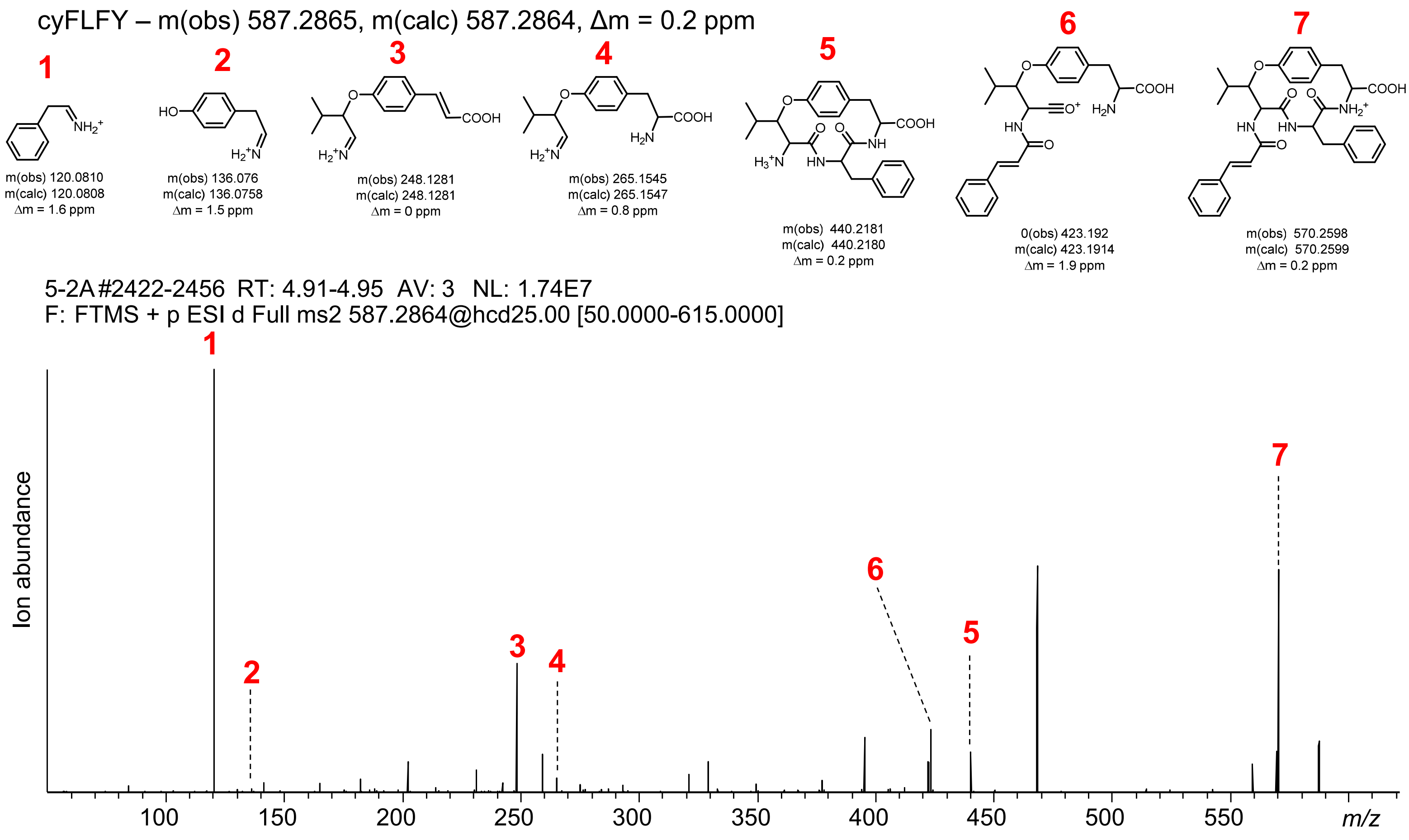


**Figure S84 | cyFLxY diversification via transient expression of SkrBURP-FLxY in *N. benthamiana*.** cyFLEY and cyFLFY cyFLGY – m(obs) 497.2394, m(calc) 497.2395, Δm = 0.2 ppm

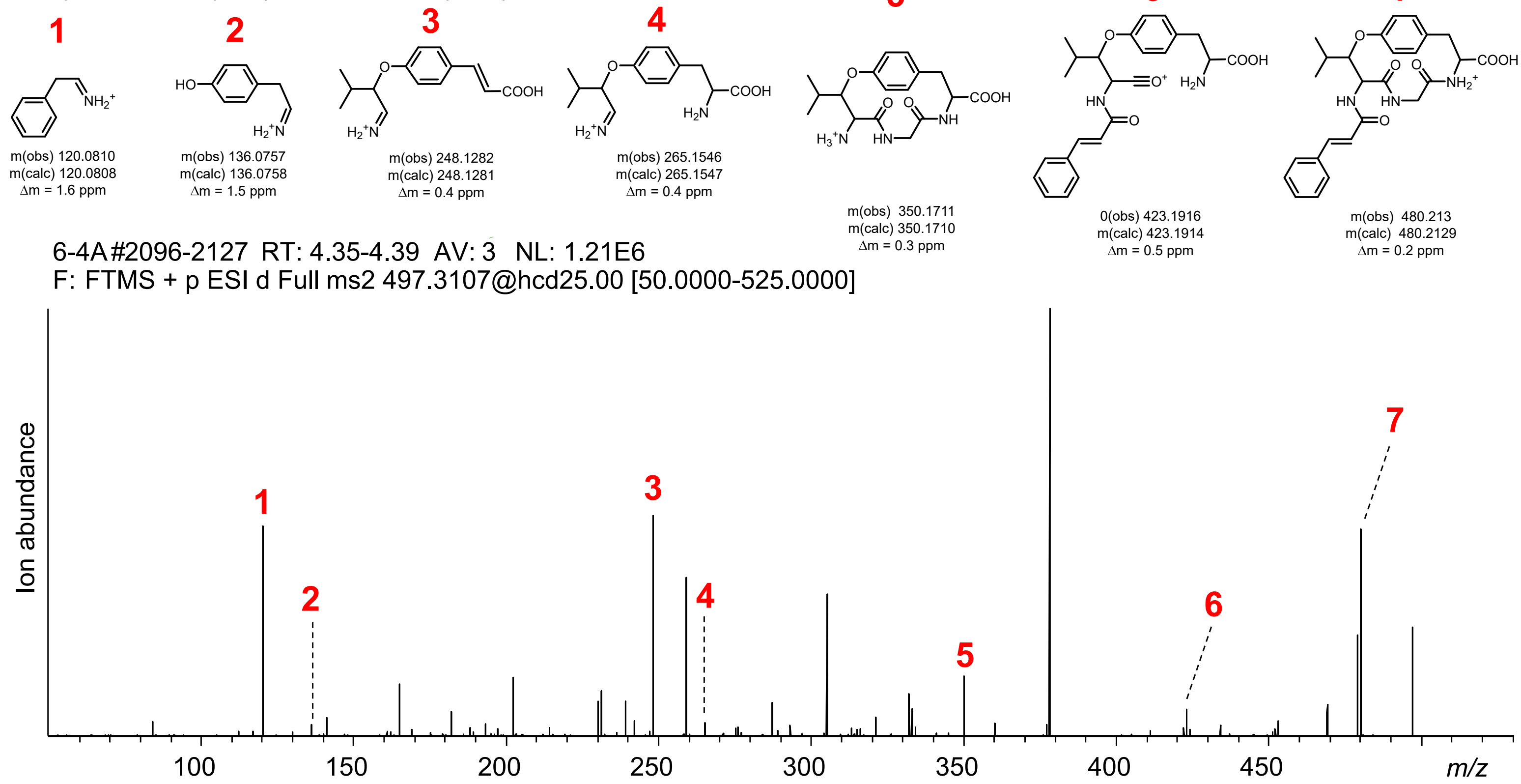


cyFLHY – m(obs) 577.2770, m(calc) 577.2769, Δm = 0.2 ppm

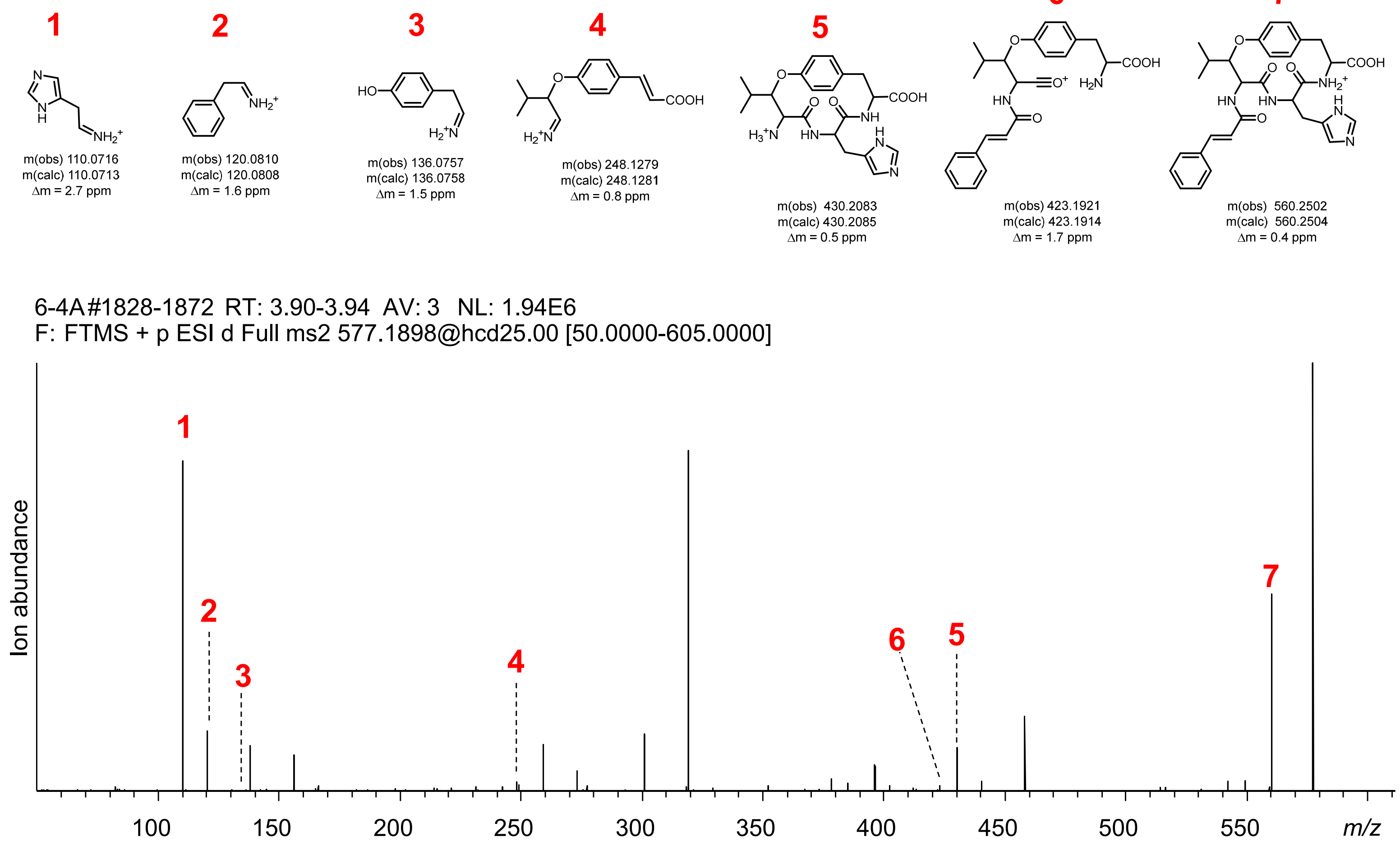


**Figure S84 | cyFLxY diversification via transient expression of SkrBURP-FLxY in *N. benthamiana*.** cyFLGY and cyFLHY cyFLIY – m(obs) 553.3018, m(calc) 553.3021, Δm = 0.5 ppm

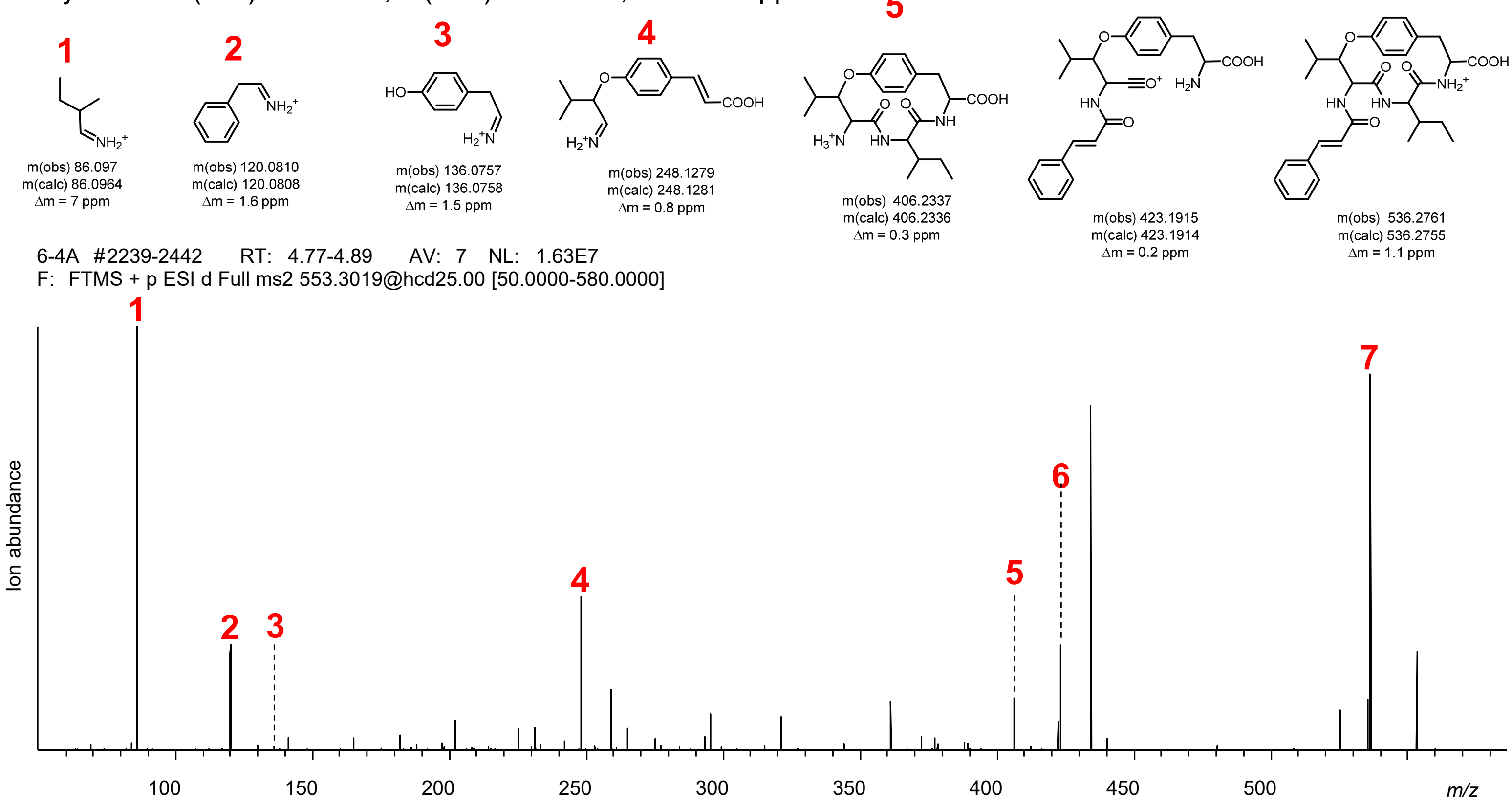


cyFLKY – m(obs) 568.3131, m(calc) 568.3130, Δm = 0.2 ppm

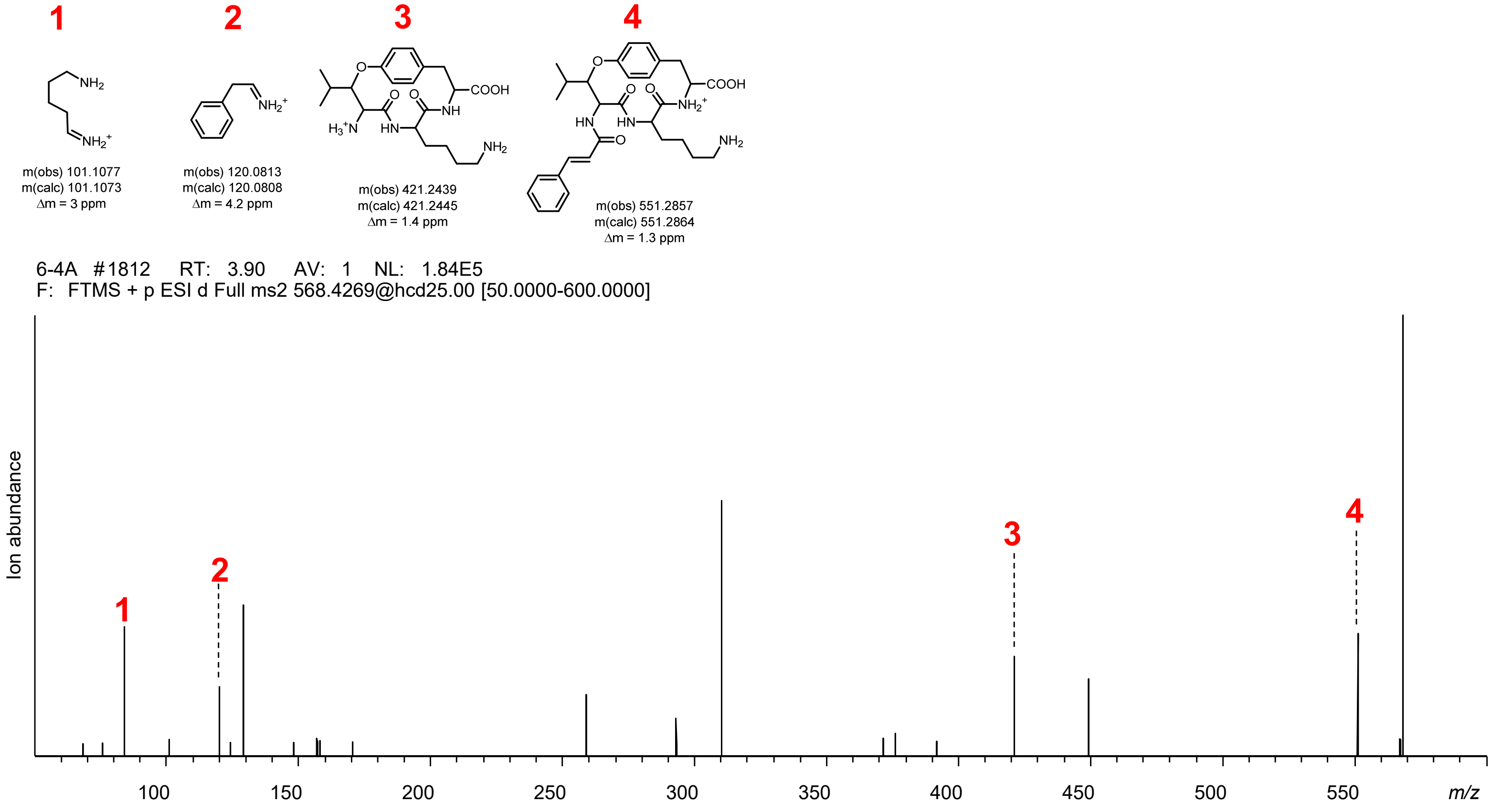


**Figure S84 | cyFLxY diversification via transient expression of SkrBURP-FLxY in *N. benthamiana*.** cyFLIY and cyFLKY cyFLLY – m(obs) 553.3018, m(calc) 553.3021, Δm = 0.5 ppm

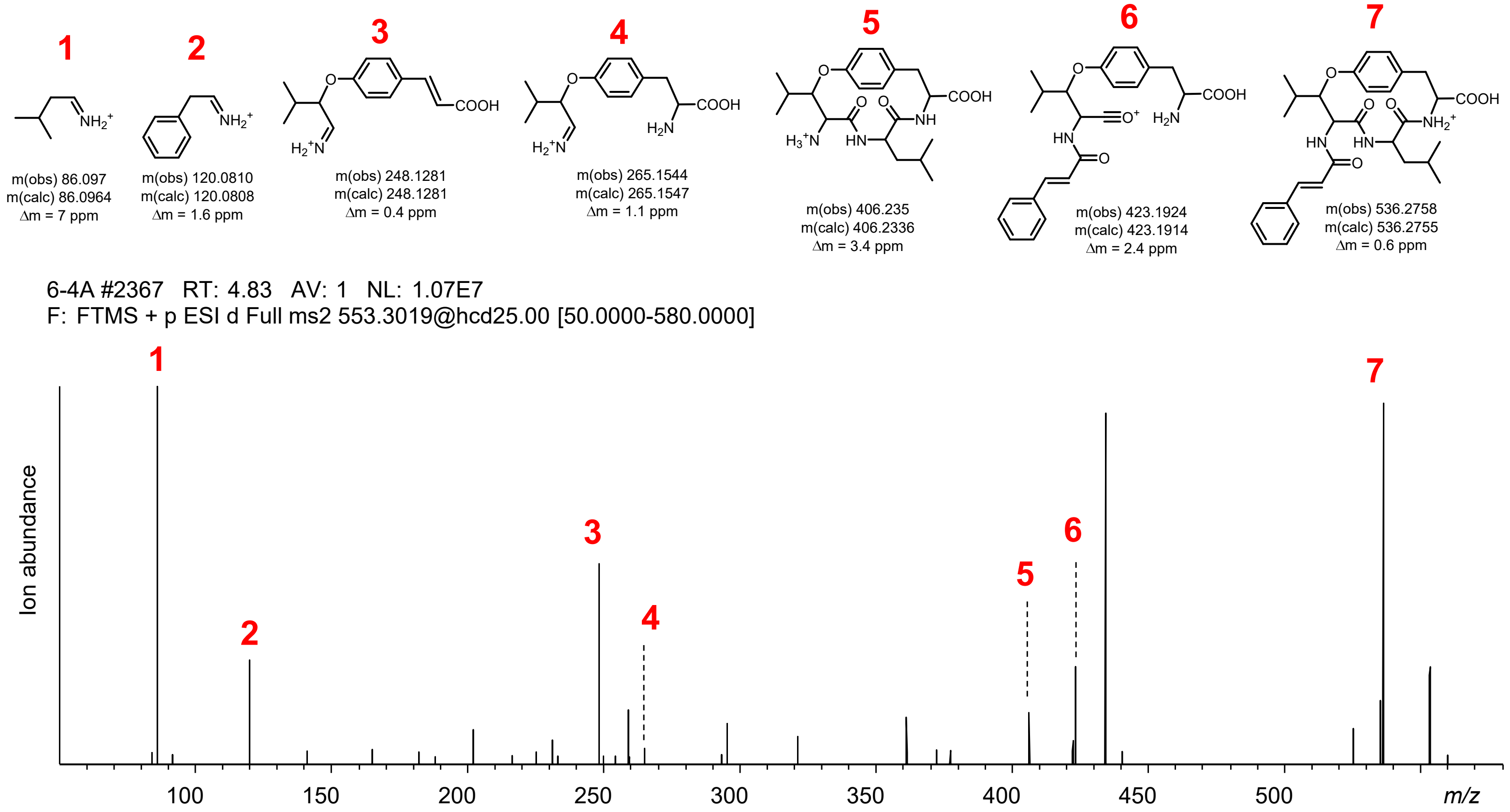


cyFLNY – m(obs) 554.2609, m(calc) 554.2609, Δm = 0 ppm

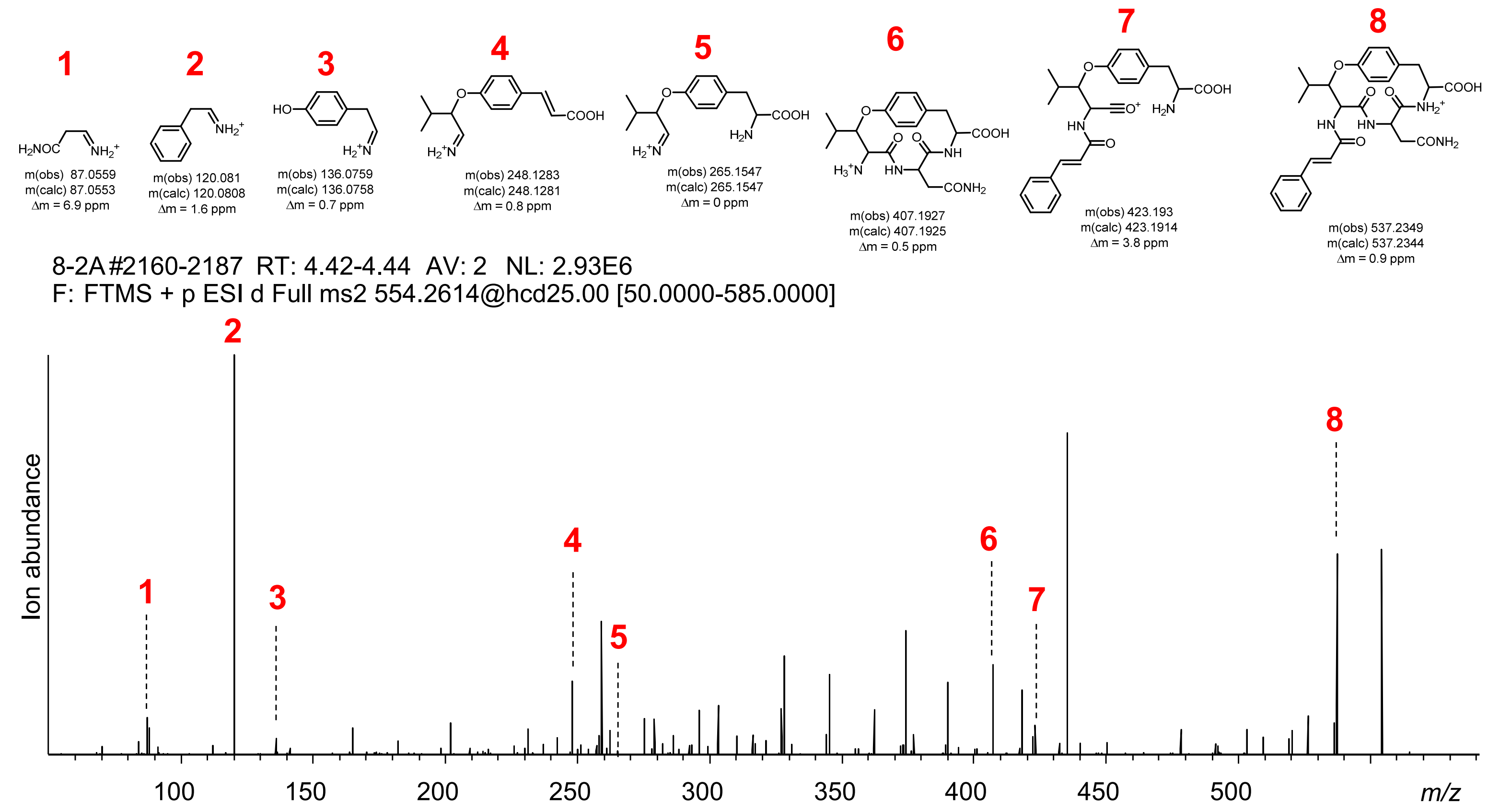


**Figure S84 | cyFLxY diversification via transient expression of SkrBURP-FLxY in *N. benthamiana*.** cyFLLY and cyFLNY cyFLPY – m(obs) 537.2711, m(calc) 537.2708, Δm = 0.6 ppm

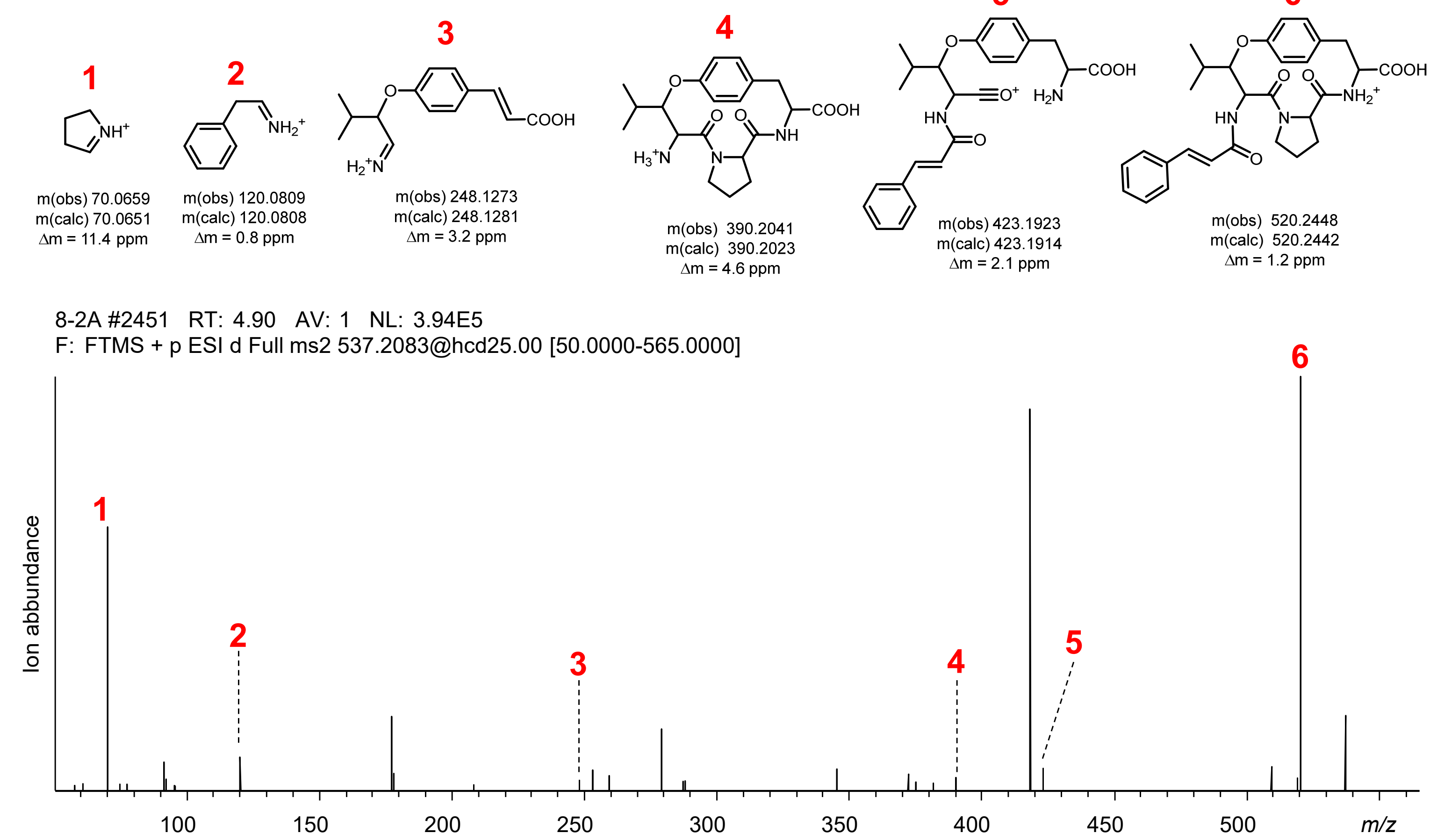


cyFLQY – m(obs) 568.2764, m(calc) 568.2766, Δm = 0.4 ppm

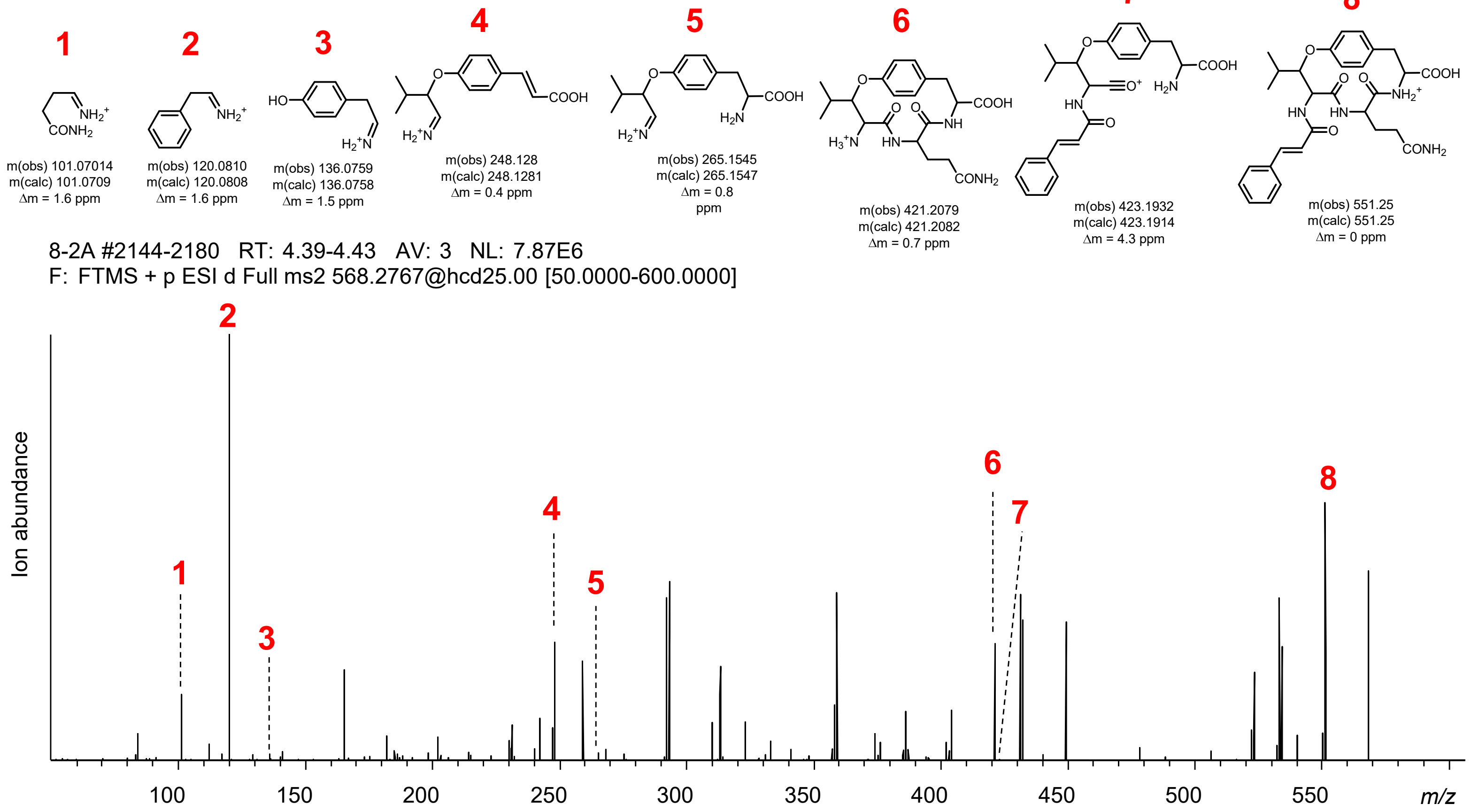


**Figure S84 | cyFLxY diversification via transient expression of SkrBURP-FLxY in *N. benthamiana*.** cyFLPY and cyFLQY cyFLRY – m(obs) 596.3192, m(calc) 596.3191, Δm = 0.2 ppm

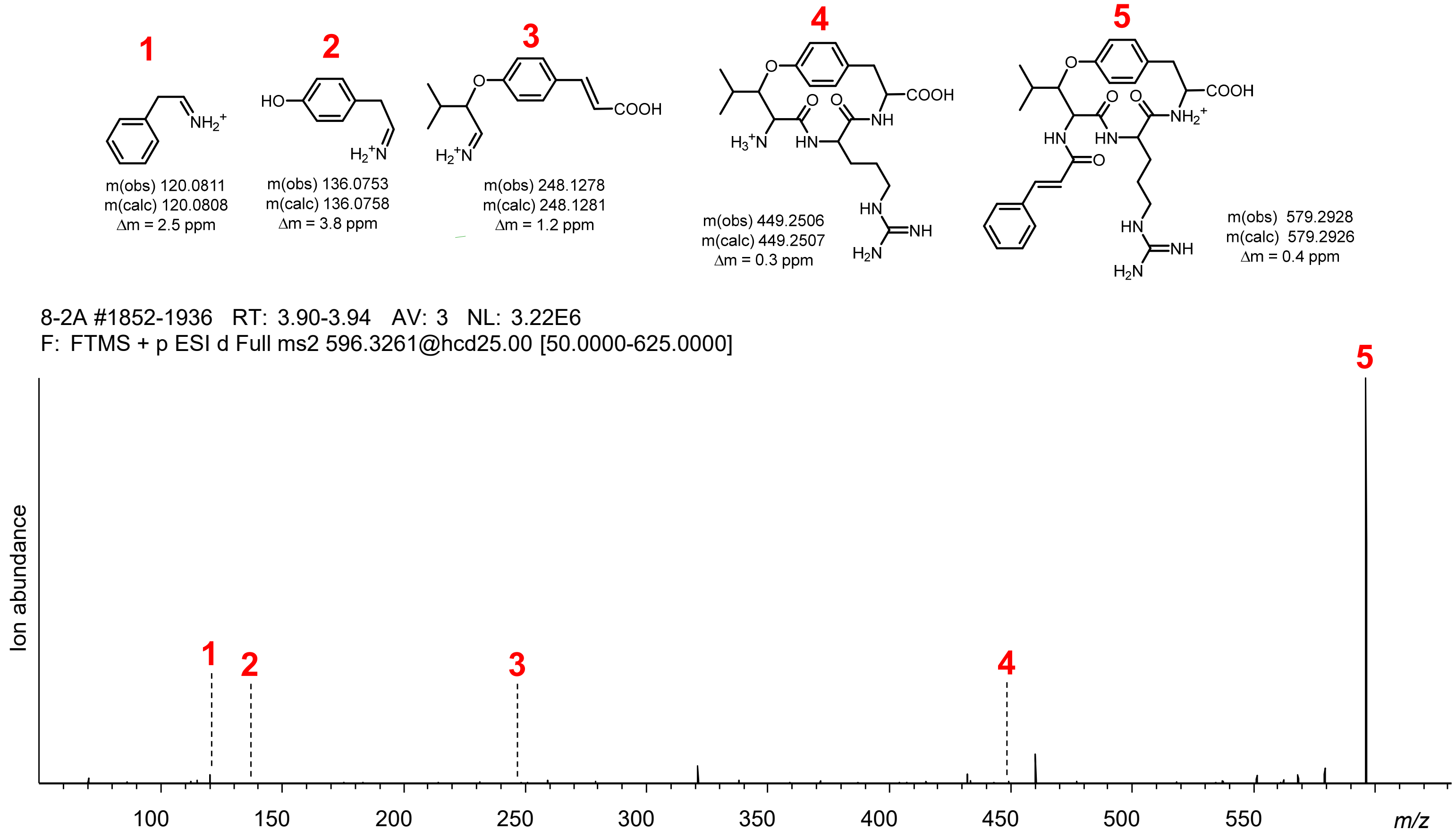


cyFLSY – m(obs) 527.2502, m(calc) 527.2500, Δm = 0.4 ppm

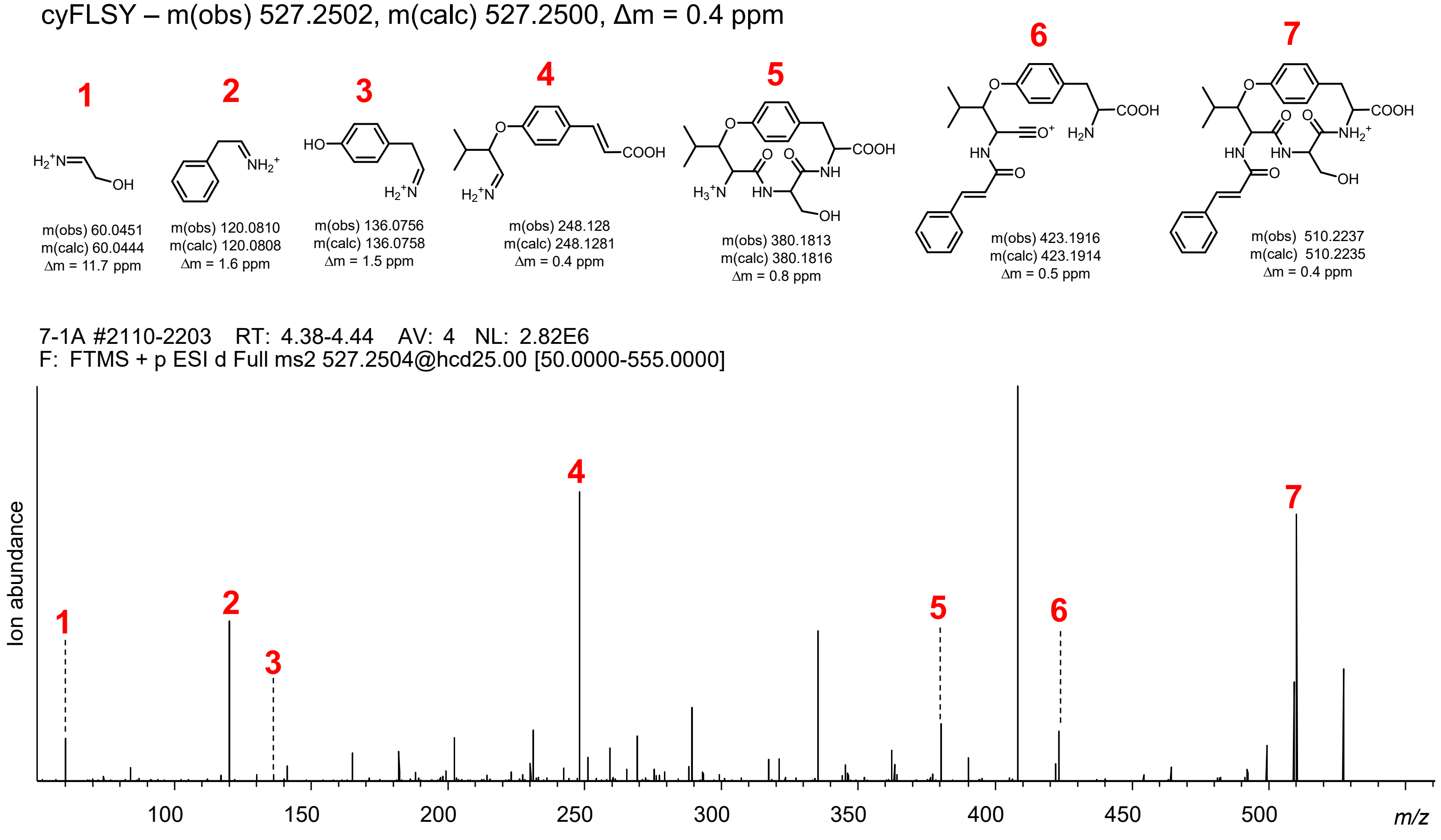


**Figure S84 | cyFLxY diversification via transient expression of SkrBURP-FLxY in *N. benthamiana*.** cyFLRY and cyFLSY cyFLTY – m(obs) 541.2654, m(calc) 541.2657, Δm = 0.6 ppm

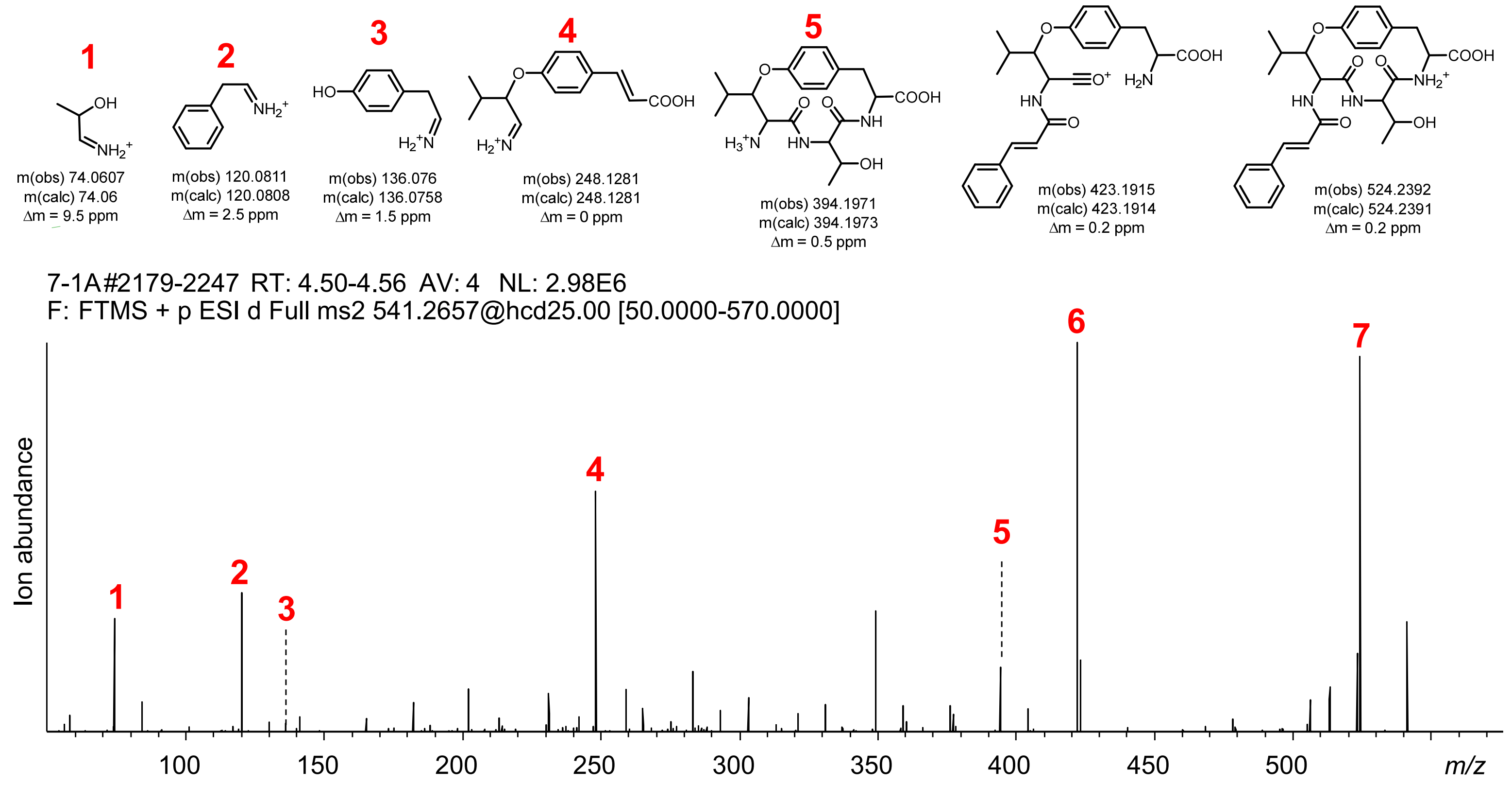


cyFLVY – m(obs) 541.2654, m(calc) 541.2657, Δm = 0.6 ppm

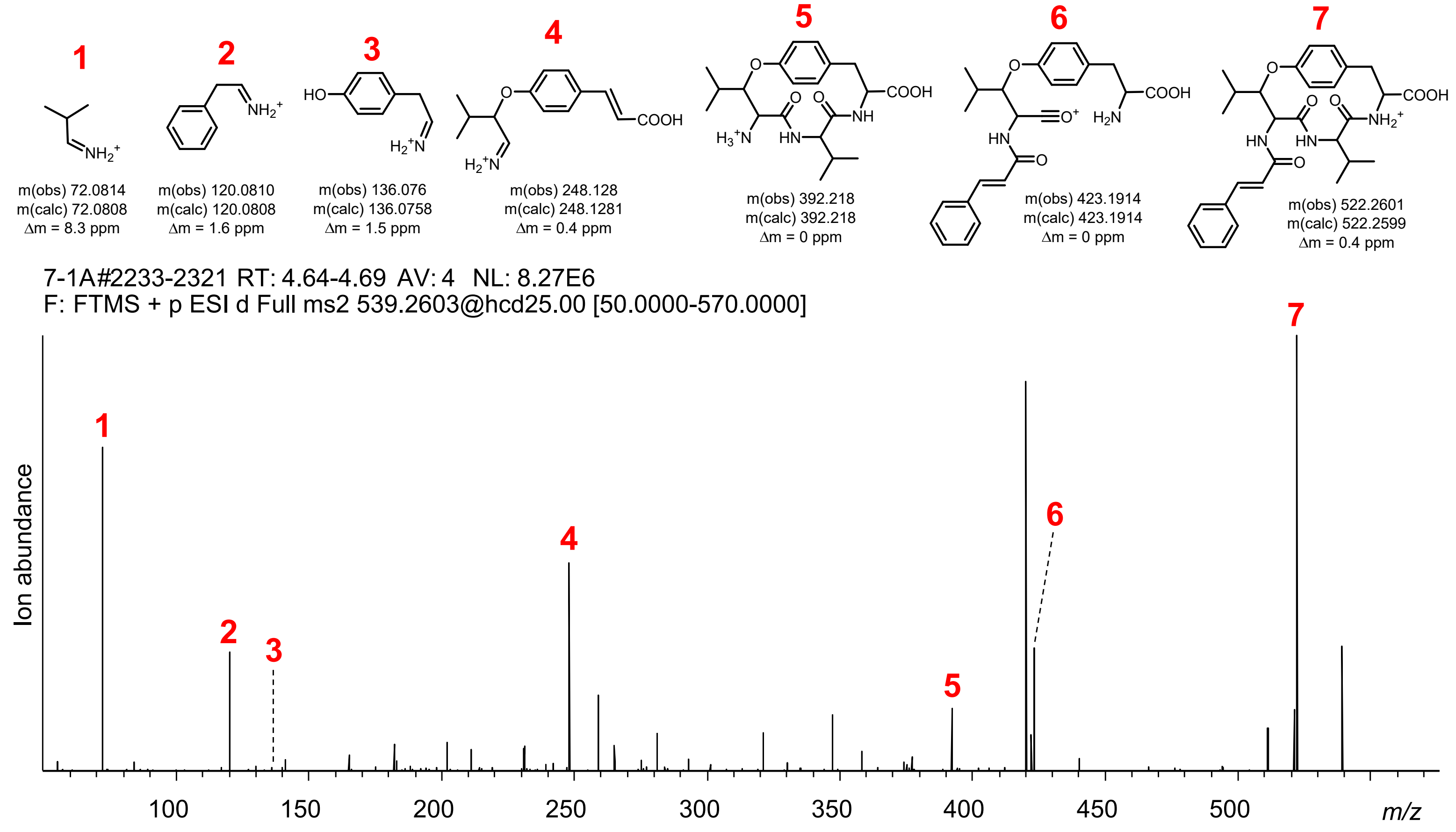


**Figure S84 | cyFLxY diversification via transient expression of SkrBURP-FLxY in *N. benthamiana*.** cyFLTY and cyFLVY cyFLWY – m(obs) 626.2967, m(calc) 626.2973, Δm = 1 ppm

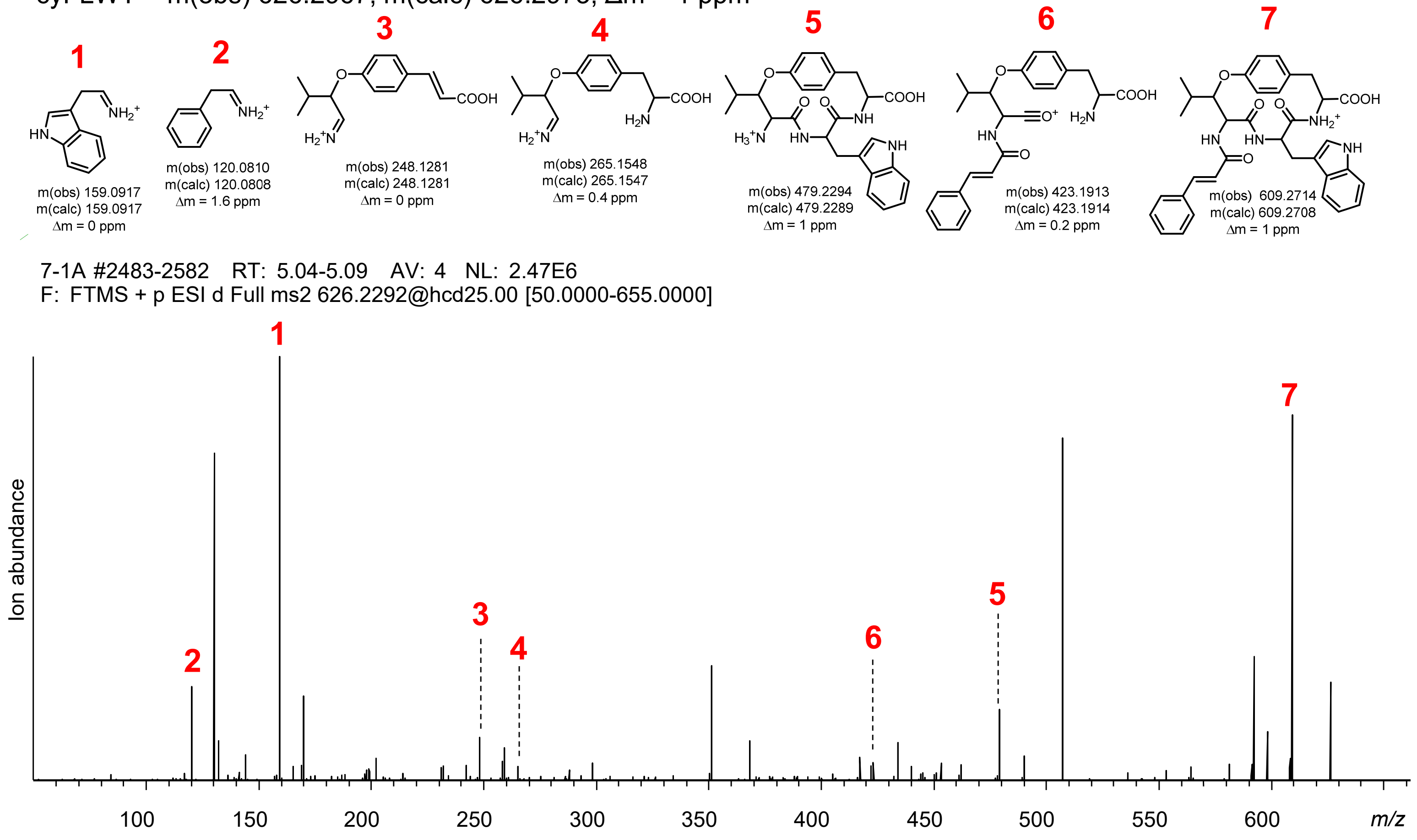


**Figure S84 | cyFLxY diversification via transient expression of SkrBURP-FLxY in *N. benthamiana*.** cyFLWY.

cyLLAY – m(obs) 477.2705, m(calc) 477.2708, Δm = 0.6 ppm

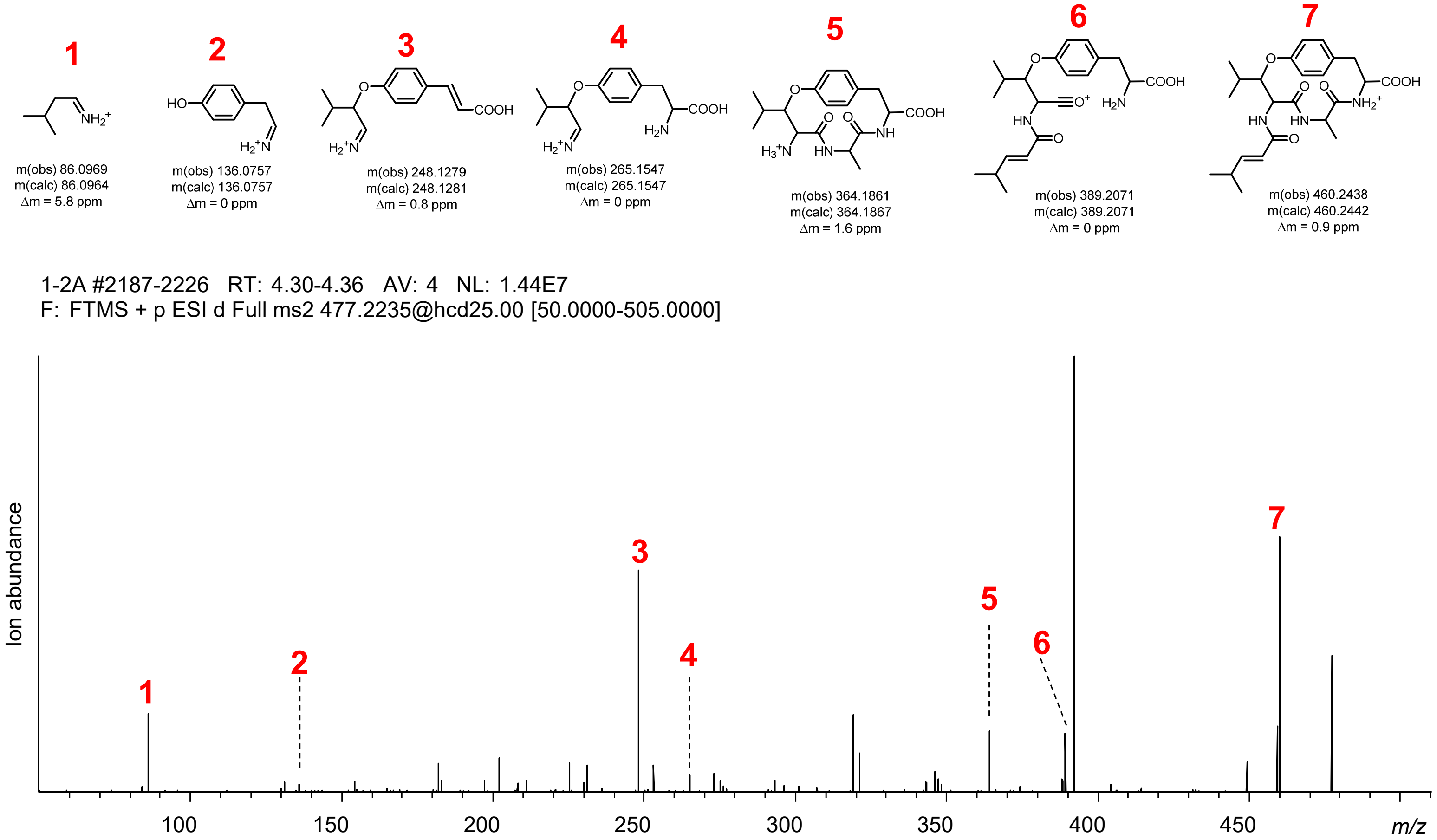


cyLLDY – m(obs) 521.2604, m(calc) 521.2606, Δm = 0.4 ppm

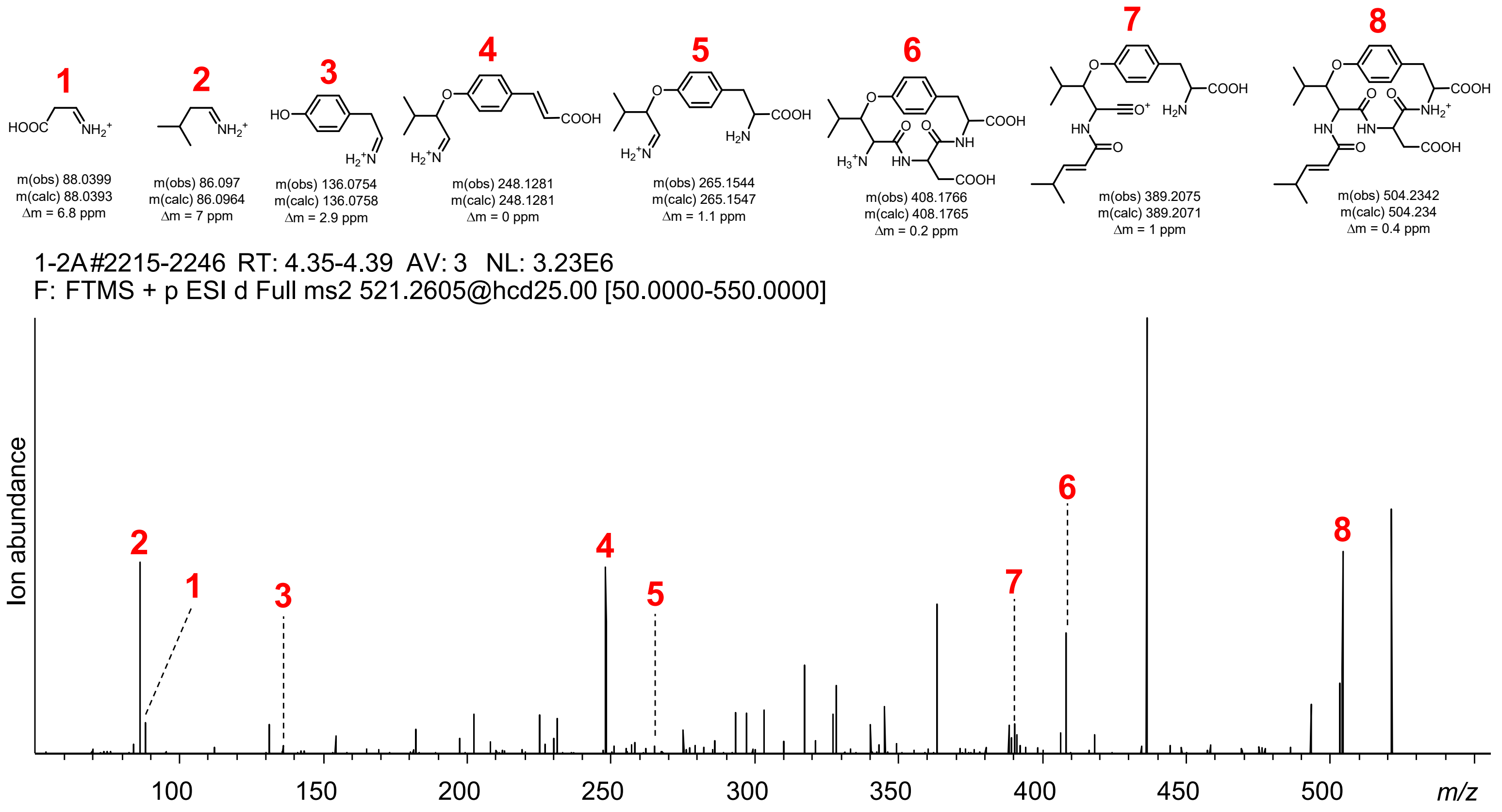


**Figure S85 | cyLLxY diversification via transient expression of SkrBURP-LLxY in *N. benthamiana*.** cyLLAY and cyLLDY.

cyLLEY – m(obs) 535.2761, m(calc) 535.2762, Δm = 0.2 ppm

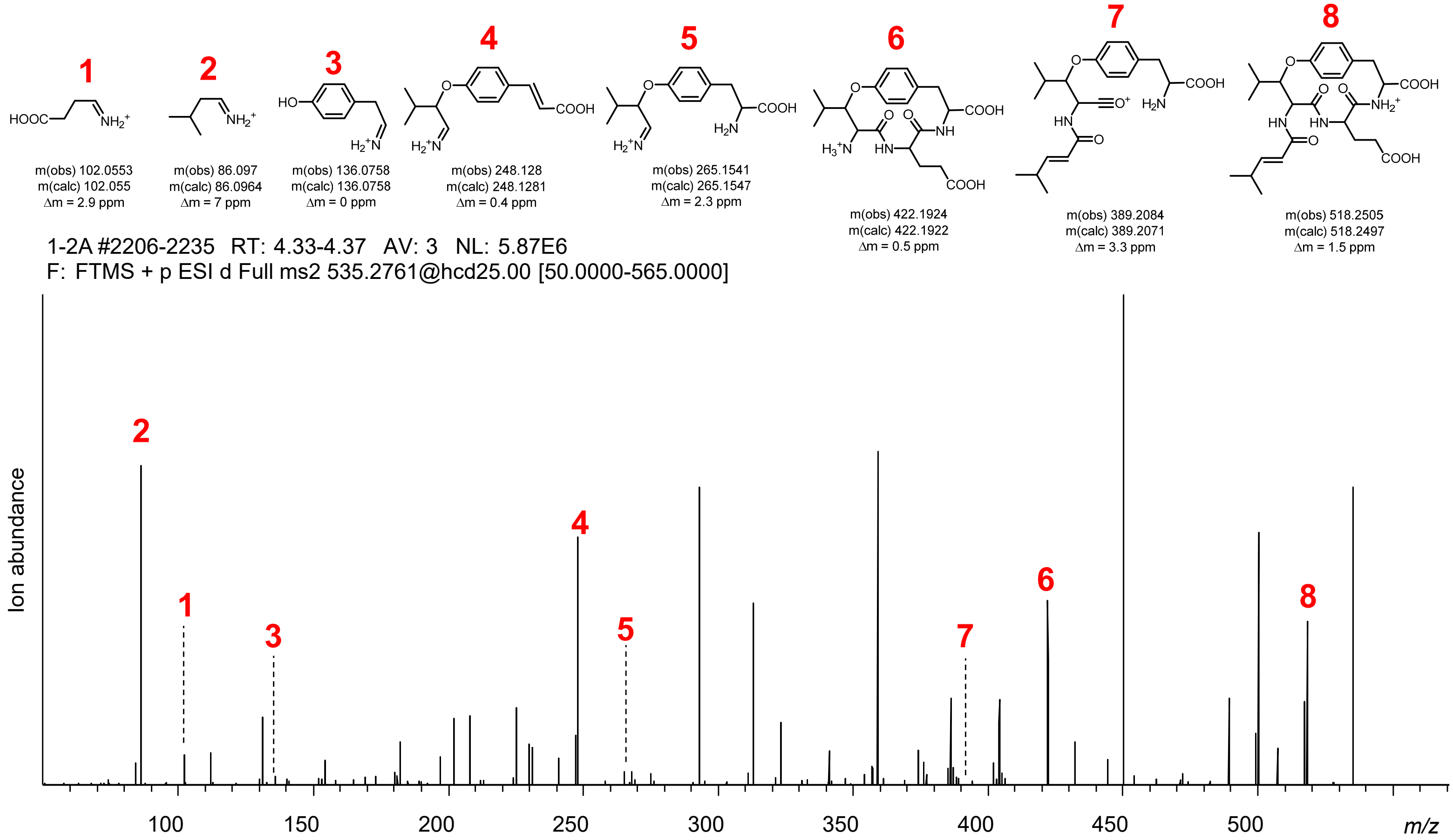


cyLLFY – m(obs) 553.3018, m(calc) 553.3021, Δm = 0.5 ppm

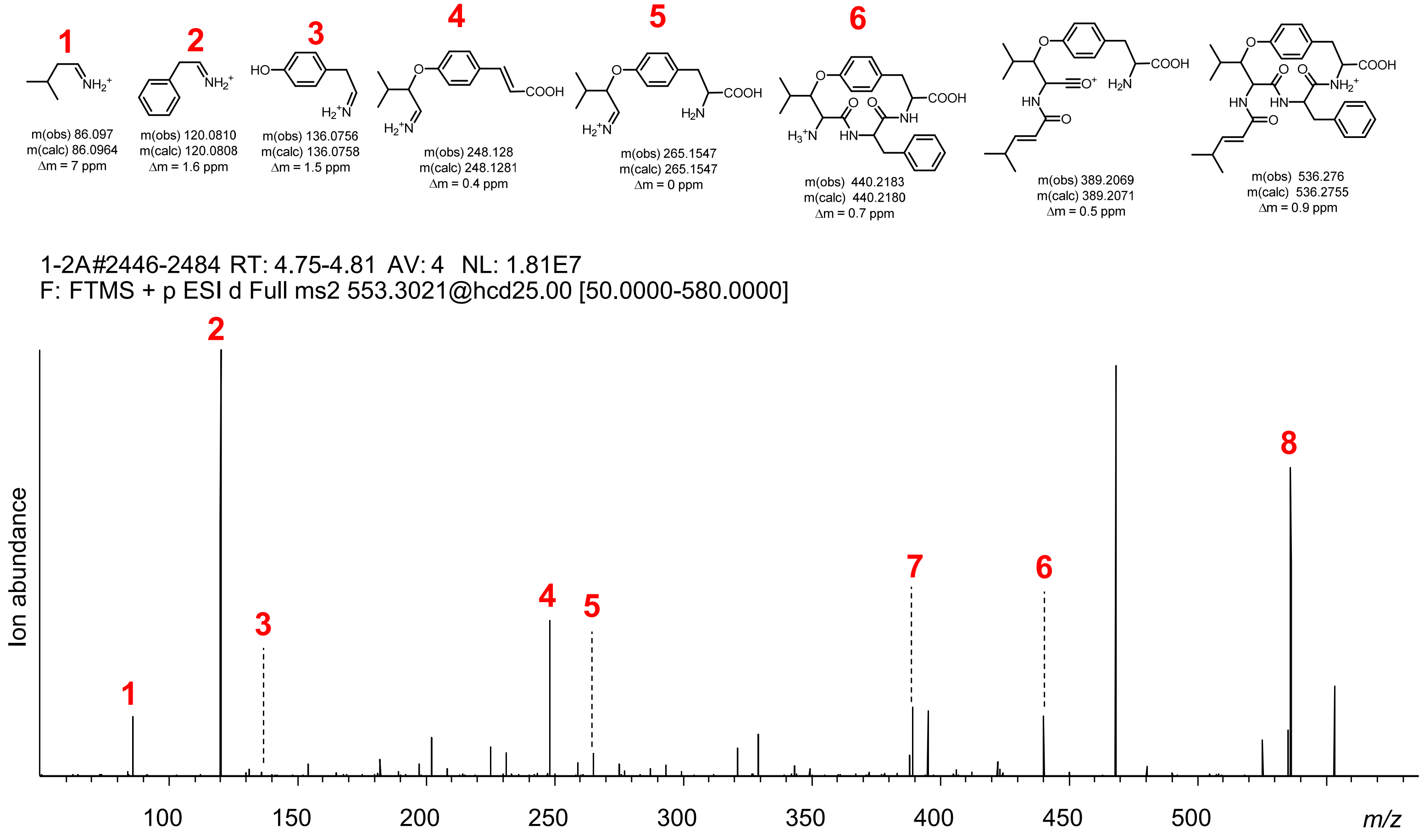


**Figure S85 | cyLLxY diversification via transient expression of SkrBURP-LLxY in *N. benthamiana*.** cyLLEY and cyLLFY cyLLGY – m(obs) 463.2547, m(calc) 463.2551, Δm = 0.9 ppm

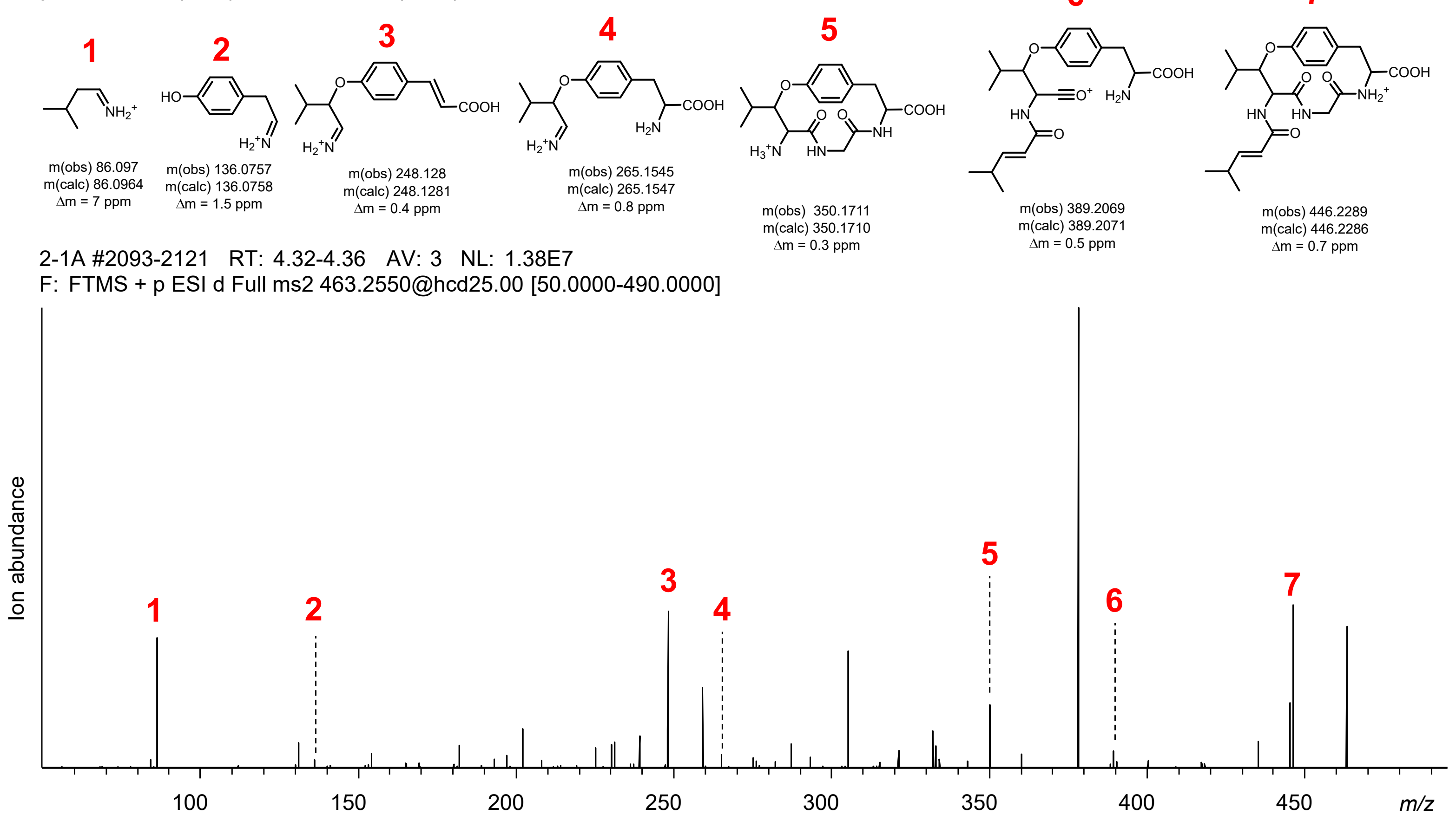


cyLLHY – m(obs) 543.2924, m(calc) 543.2926, Δm = 0.4 ppm

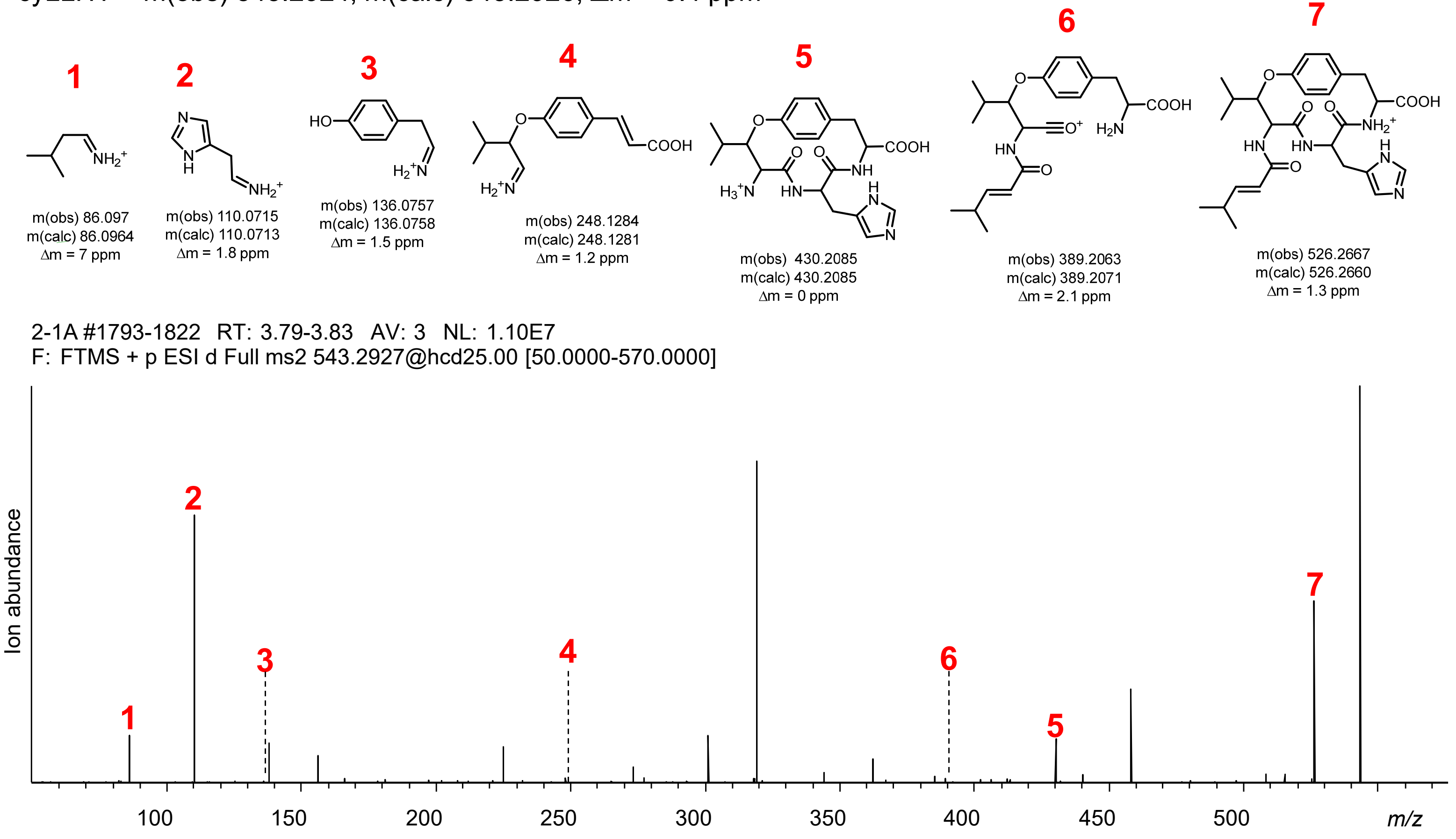


**Figure S85 | cyLLxY diversification via transient expression of SkrBURP-LLxY in *N. benthamiana*.** cyLLGY and cyLLHY.

cyLLIY – m(obs) 519.3174, m(calc) 519.3177, Δm = 0.6 ppm

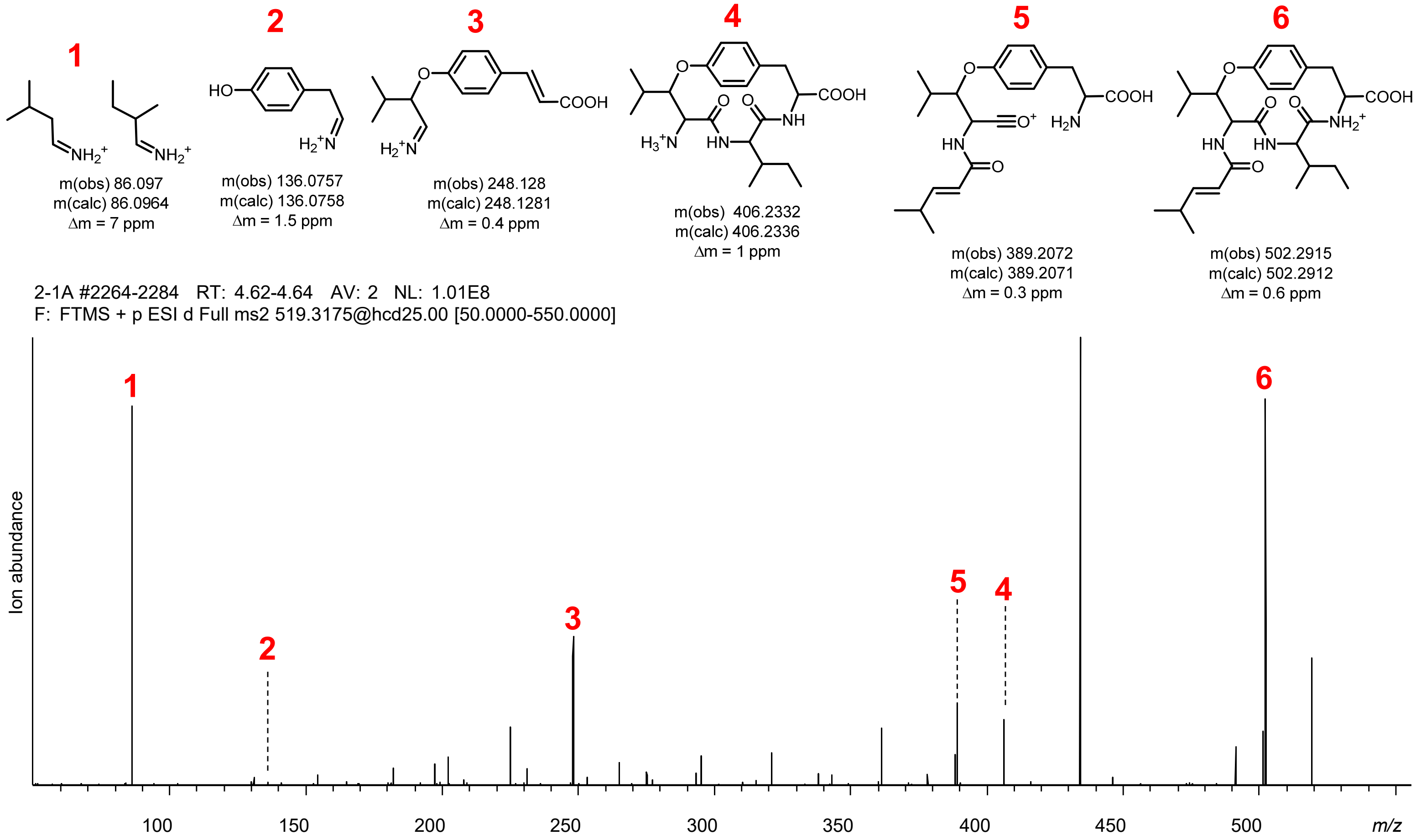


cyLLKY – m(obs) 534.3291, m(calc) 534.3286, Δm = 0.9 ppm

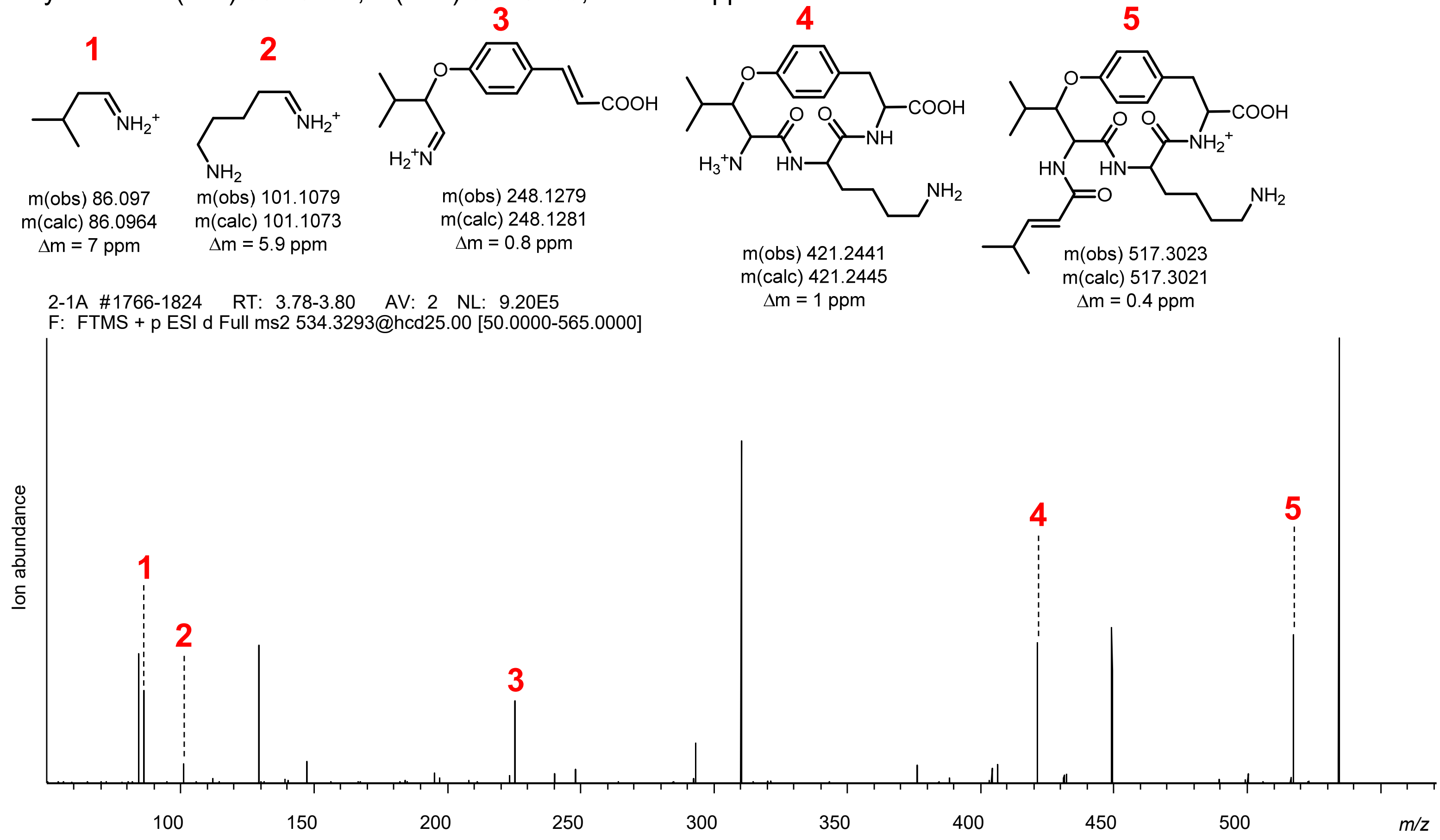


**Figure S85 | cyLLxY diversification via transient expression of SkrBURP-LLxY in *N. benthamiana*.** cyLLIY and cyLLKY cyLLLY – m(obs) 519.3173, m(calc) 519.3177, Δm = 0.8 ppm

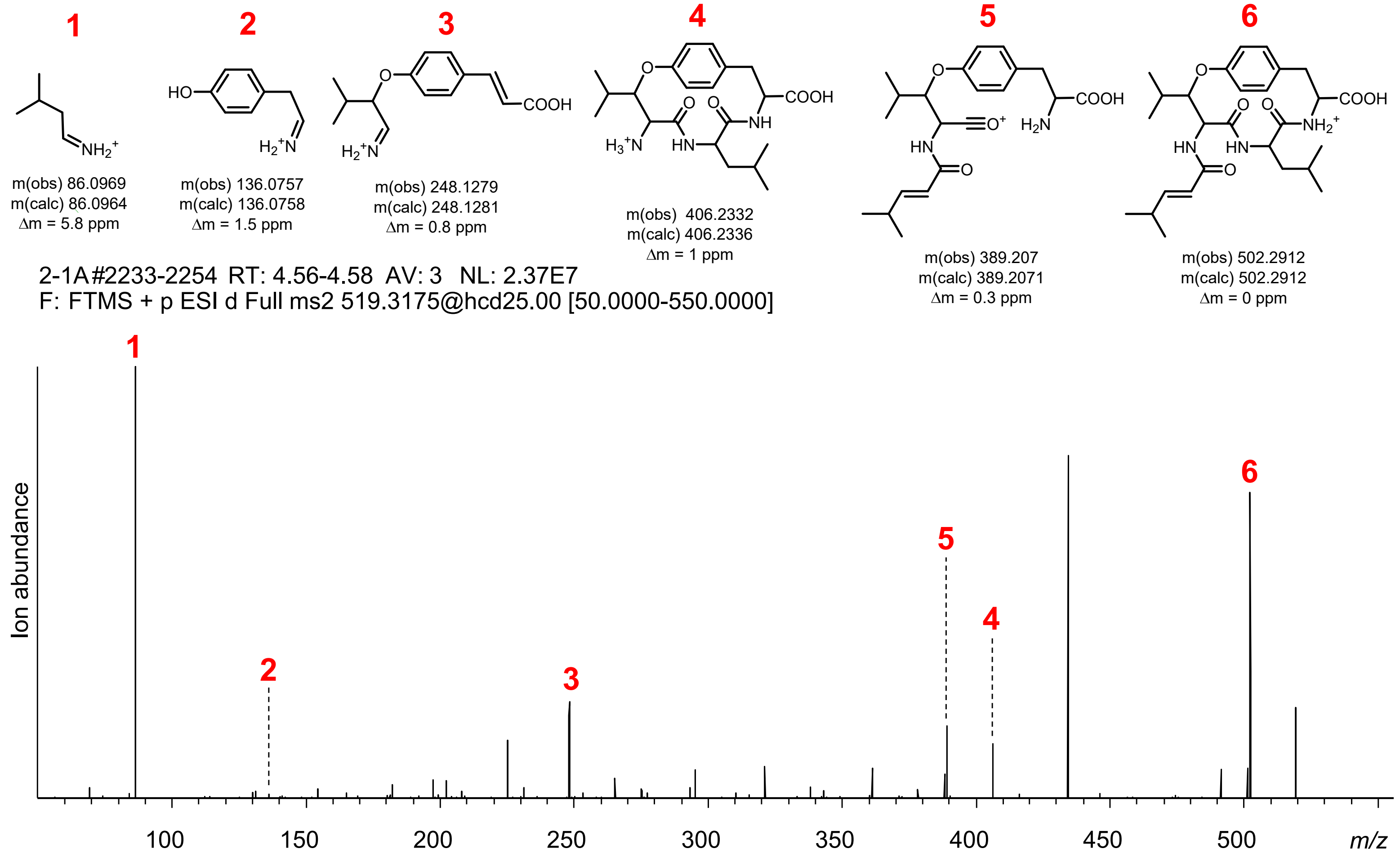


cyLLNY – m(obs) 520.2764, m(calc) 520.2766, Δm = 0.4 ppm

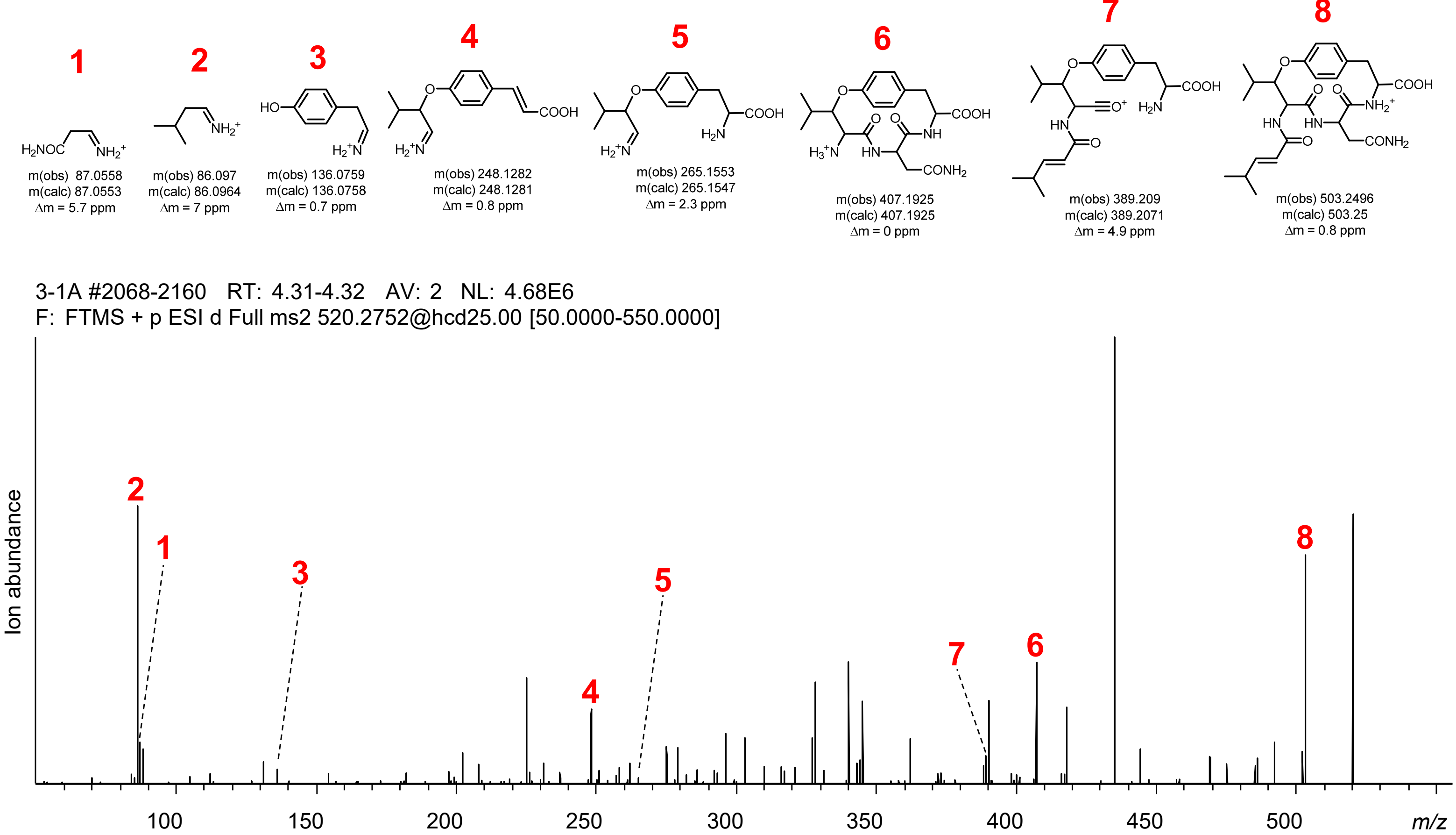


**Figure S85 | cyLLxY diversification via transient expression of SkrBURP-LLxY in *N. benthamiana*.** cyLLLY and cyLLNY cyLLPY – m(obs) 503.2866, m(calc) 503.2864, Δm = 0.4 ppm

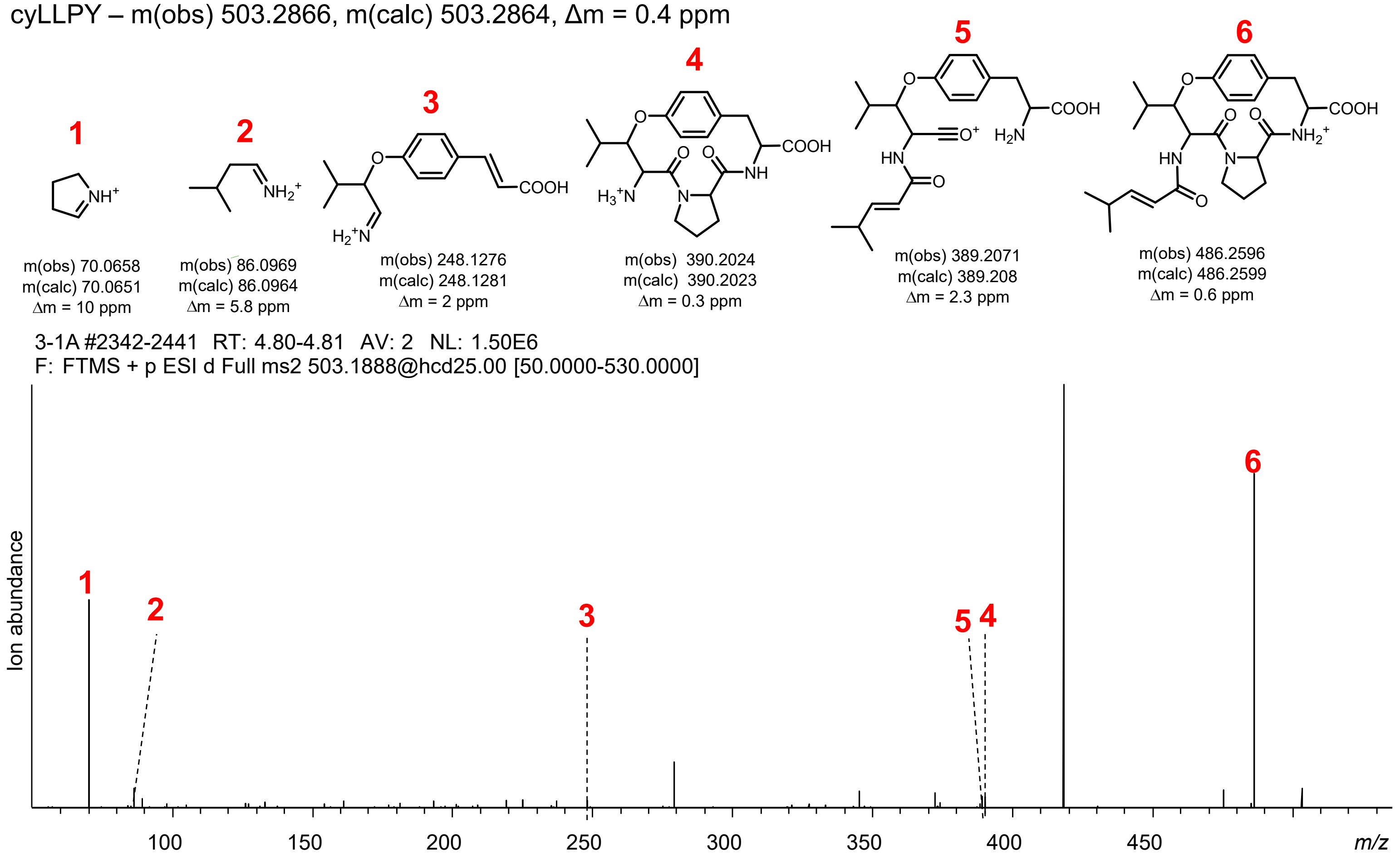


cyLLQY – m(obs) 534.2917, m(calc) 534.2922, Δm = 0.9 ppm

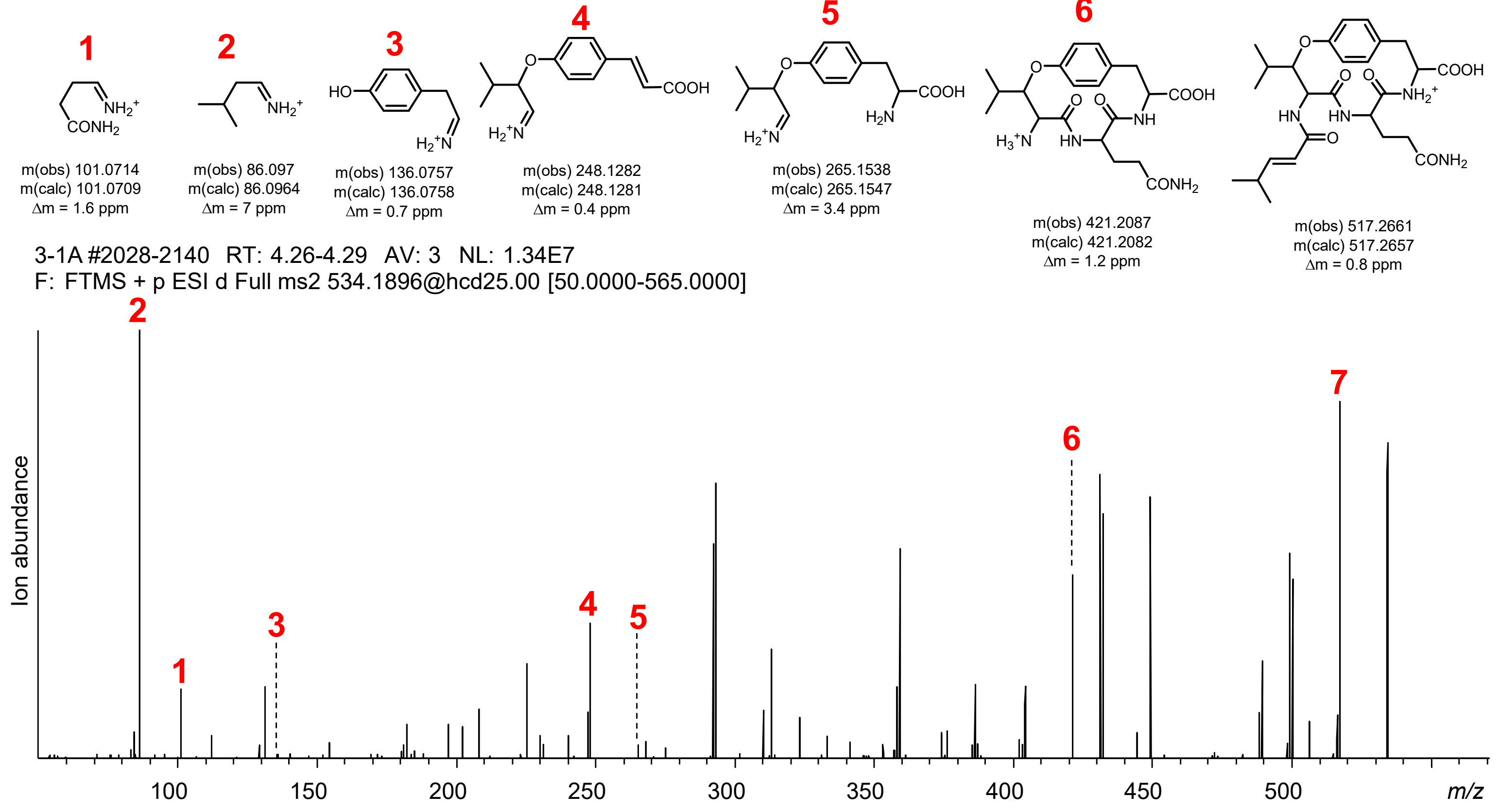


**Figure S85 | cyLLxY diversification via transient expression of SkrBURP-LLxY in *N. benthamiana*.** cyLLPY and cyLLQY cyLLRY – m(obs) 562.335, m(calc) 562.3348, Δm = 0.4 ppm

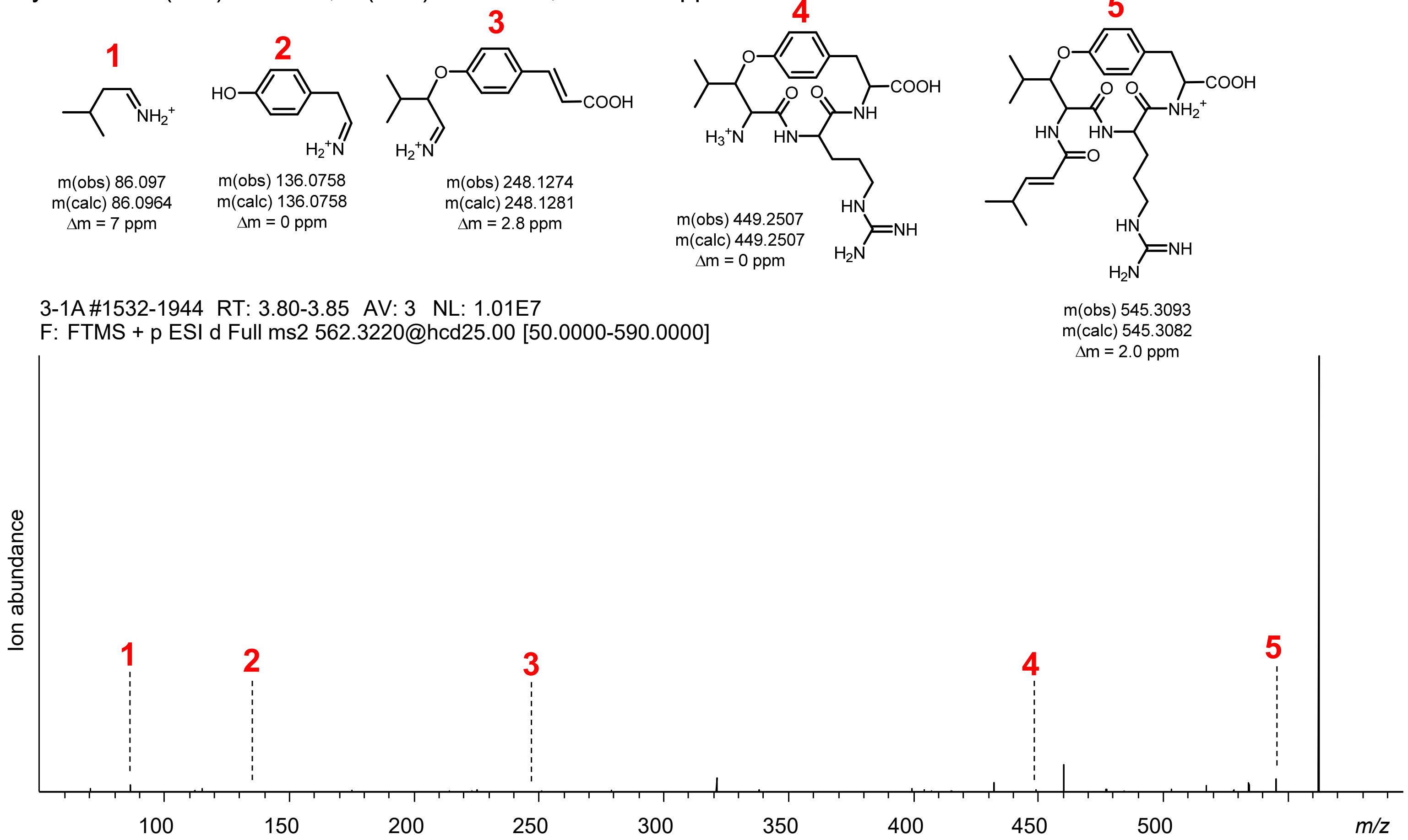


cyLLSY – m(obs) 493.2653, m(calc) 493.2657, Δm = 0.8 ppm

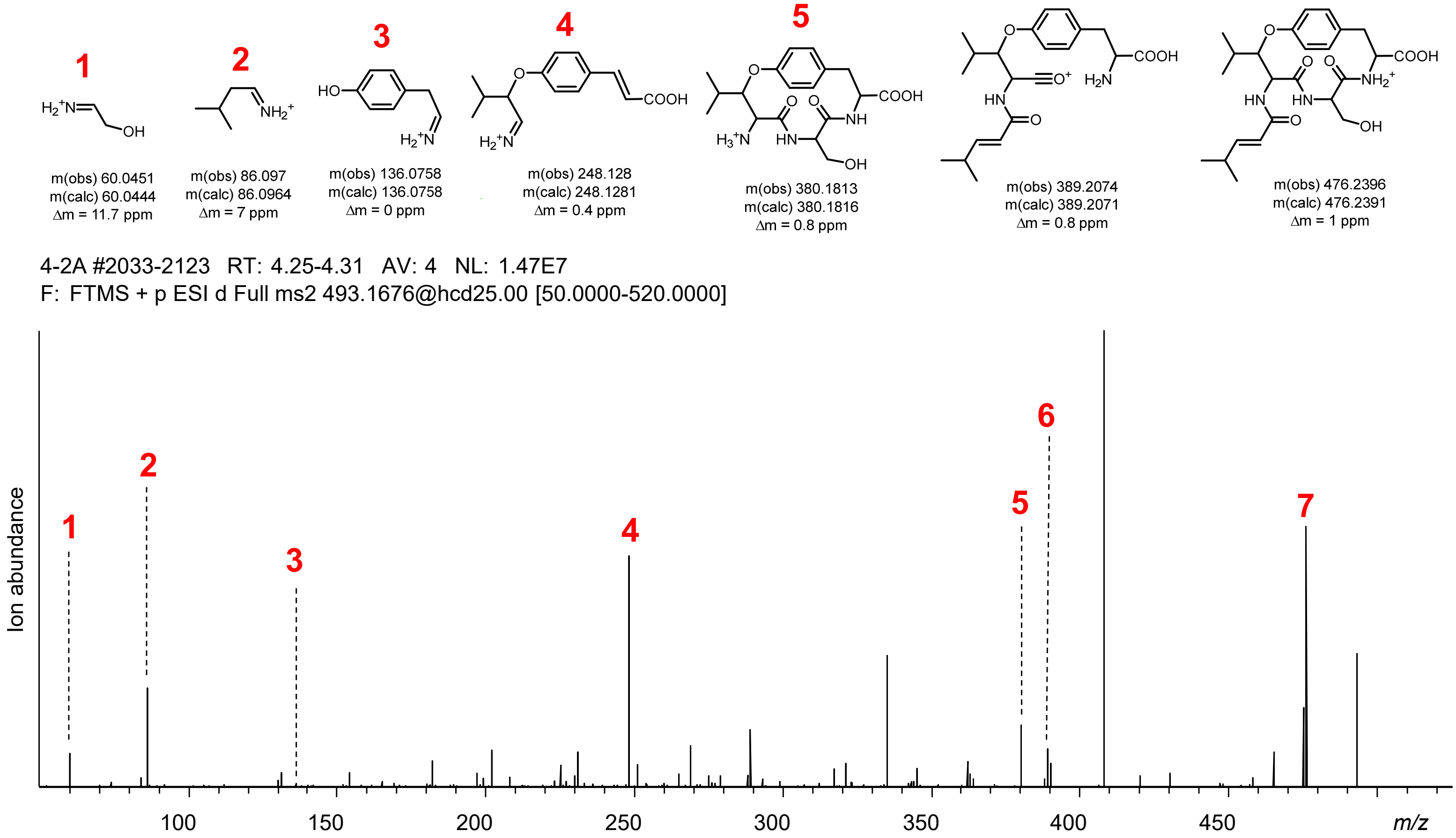


**Figure S85 | cyLLxY diversification via transient expression of SkrBURP-LLxY in *N. benthamiana.*** cyLLRY and cyLLSY

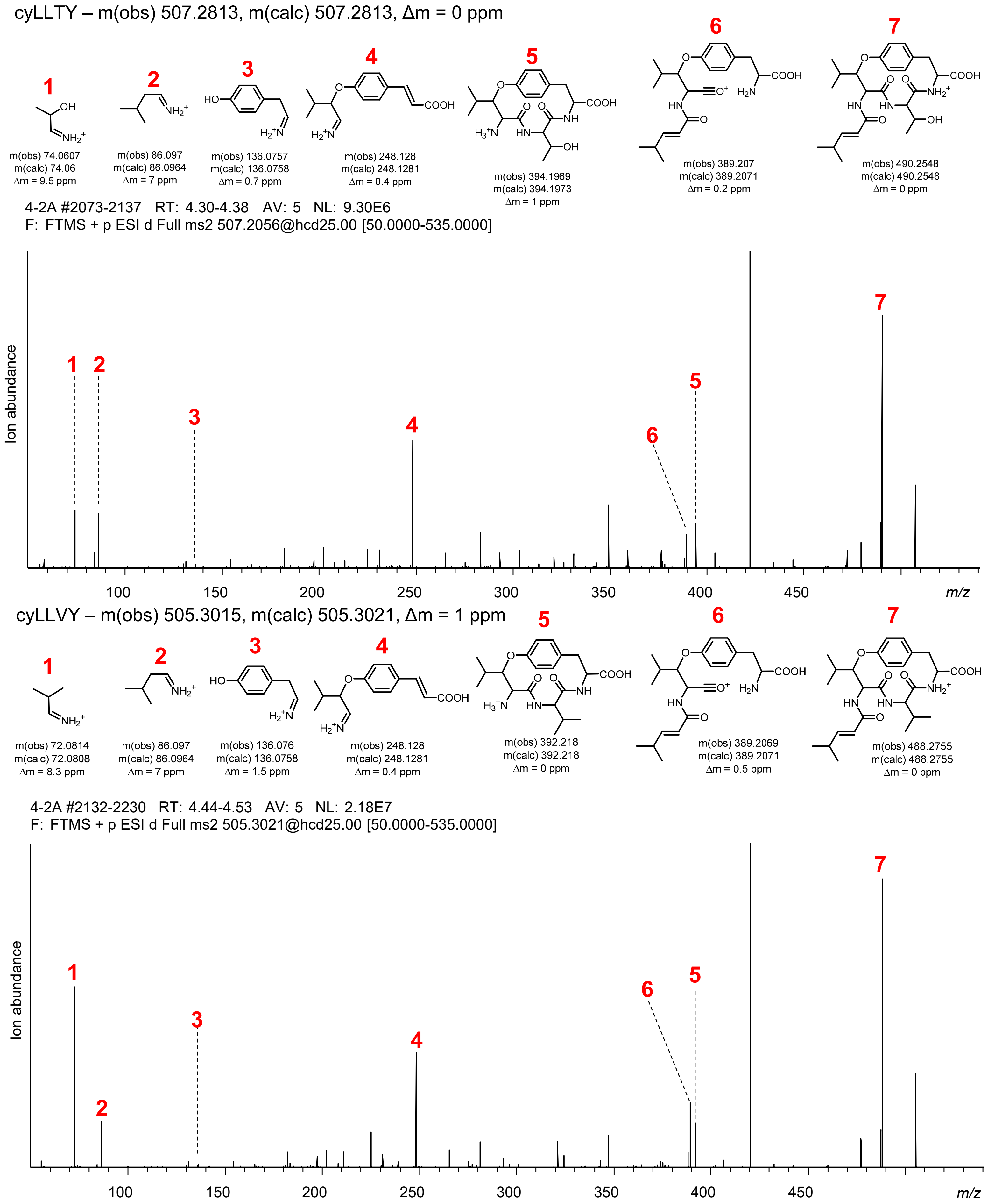


**Figure S85 | cyLLxY diversification via transient expression of SkrBURP-LLxY in *N. benthamiana*.** cyLLTY and cyLLVY cyLLWY – m(obs) 592.3125, m(calc) 592.3130, Δm = 0.8 ppm

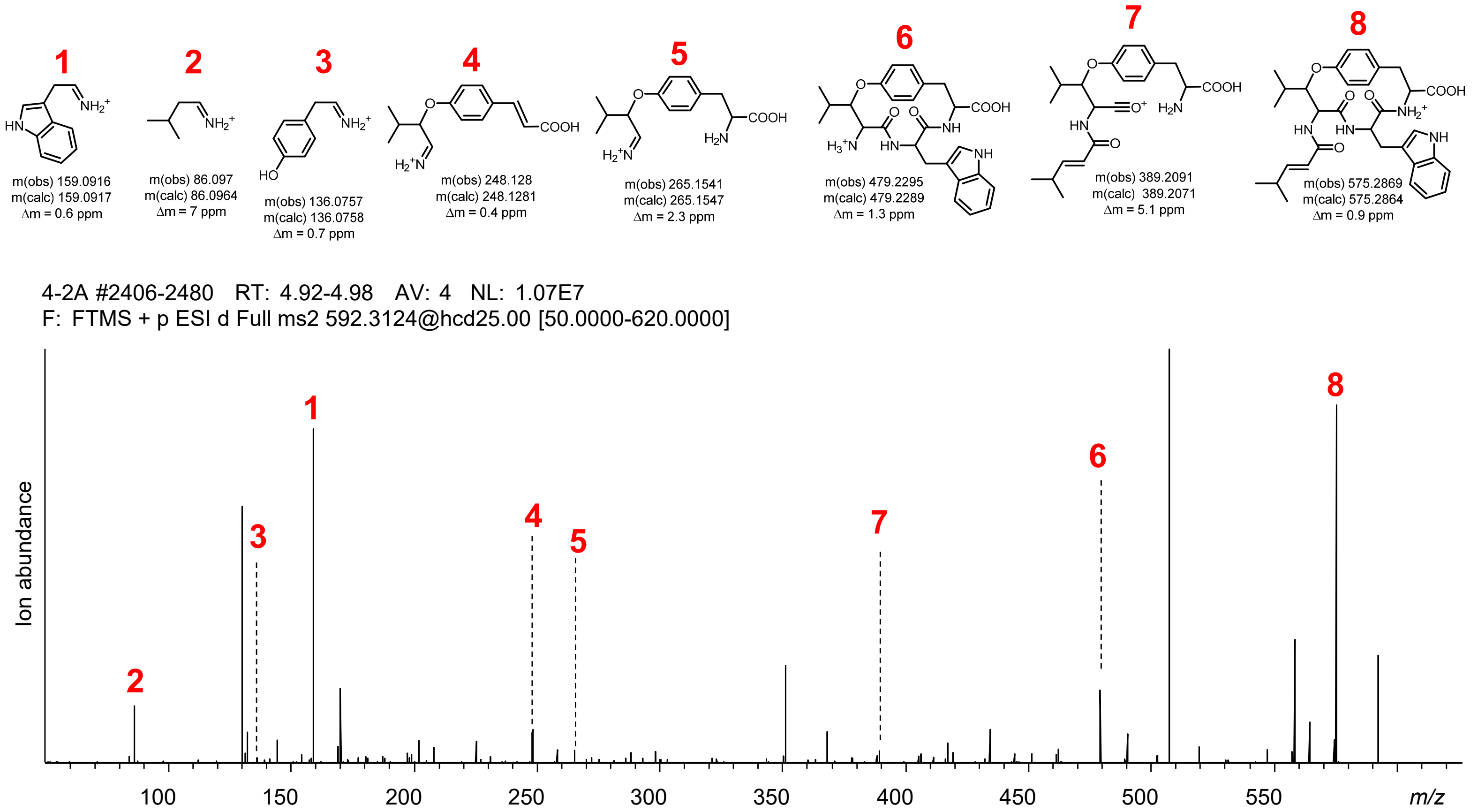


cyLLYY – m(obs) 505.3015, m(calc) 505.3021, Δm = 1 ppm

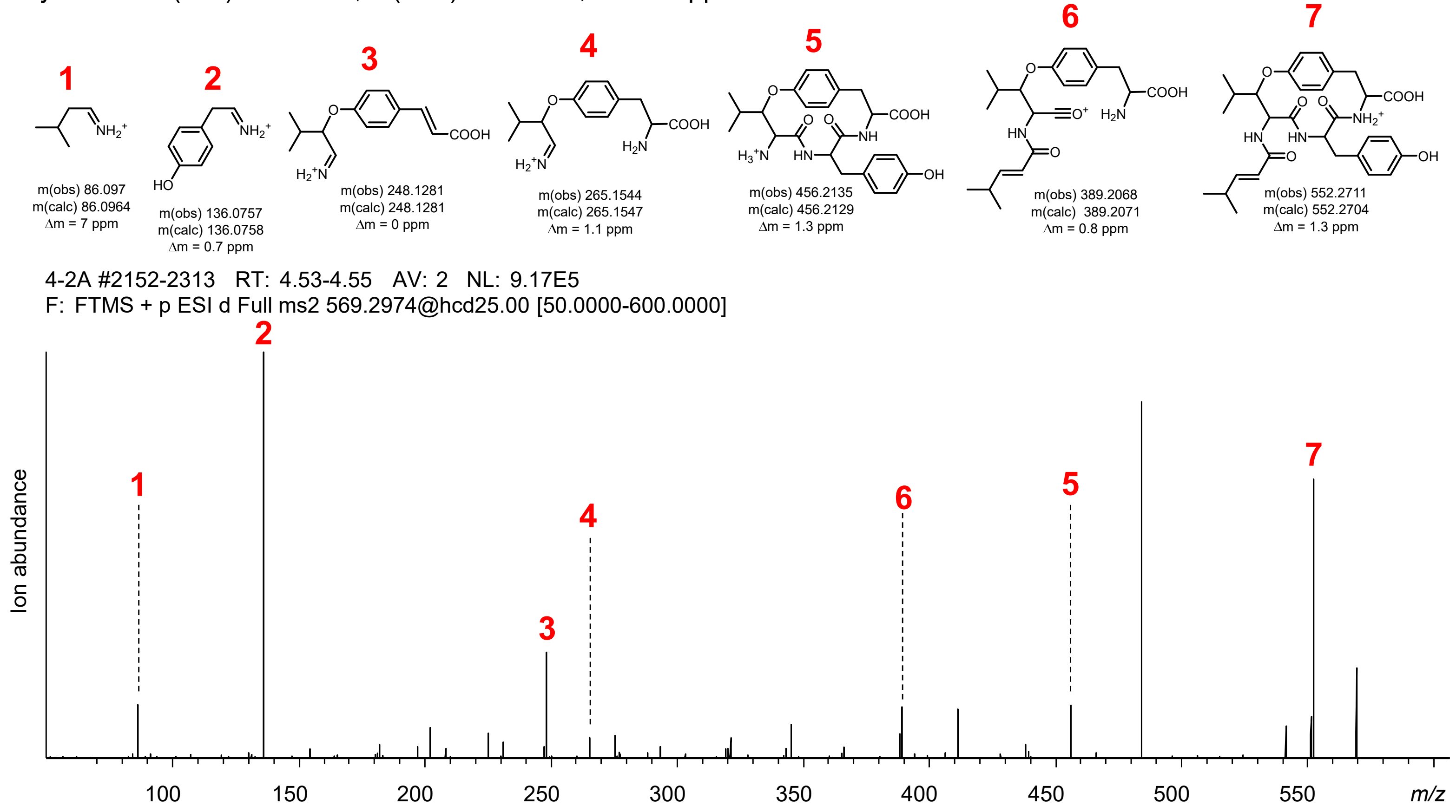


**Figure S85 | cyLLxY diversification via transient expression of SkrBURP-LLxY in *N. benthamiana*.** cyLLWY and cyLLYY

Sanjoinine A (FLAY) – m(obs) 493.2809, m(calc) 493.2809, Δm = 1 ppm

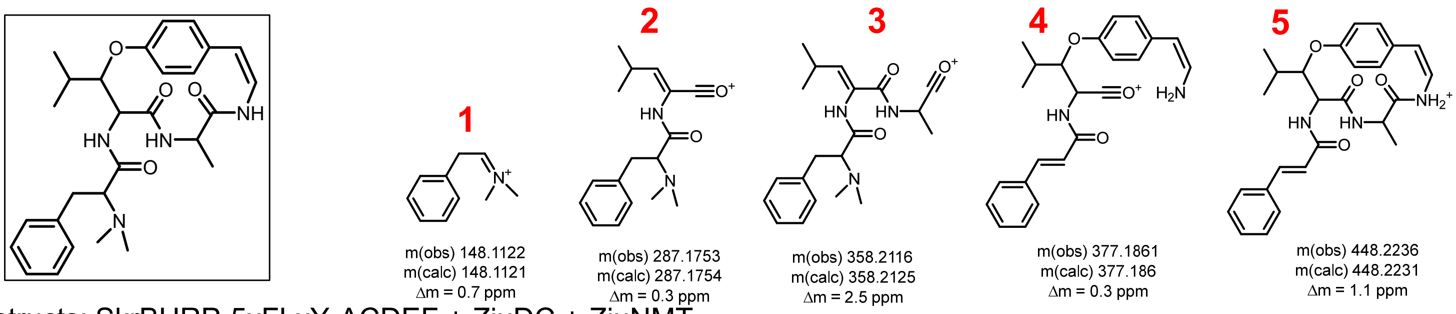


Constructs: SkrBURP-5xFLxY-ACDEF + ZjuDC + ZjuNMT

5+DC+NMT-B #2646-2695 RT: 5.02-5.04 AV: 2 NL: 4.88E6
F: FTMS + p ESI d Full ms2 493.2188@hcd25.00 [50.0000-520.0000]

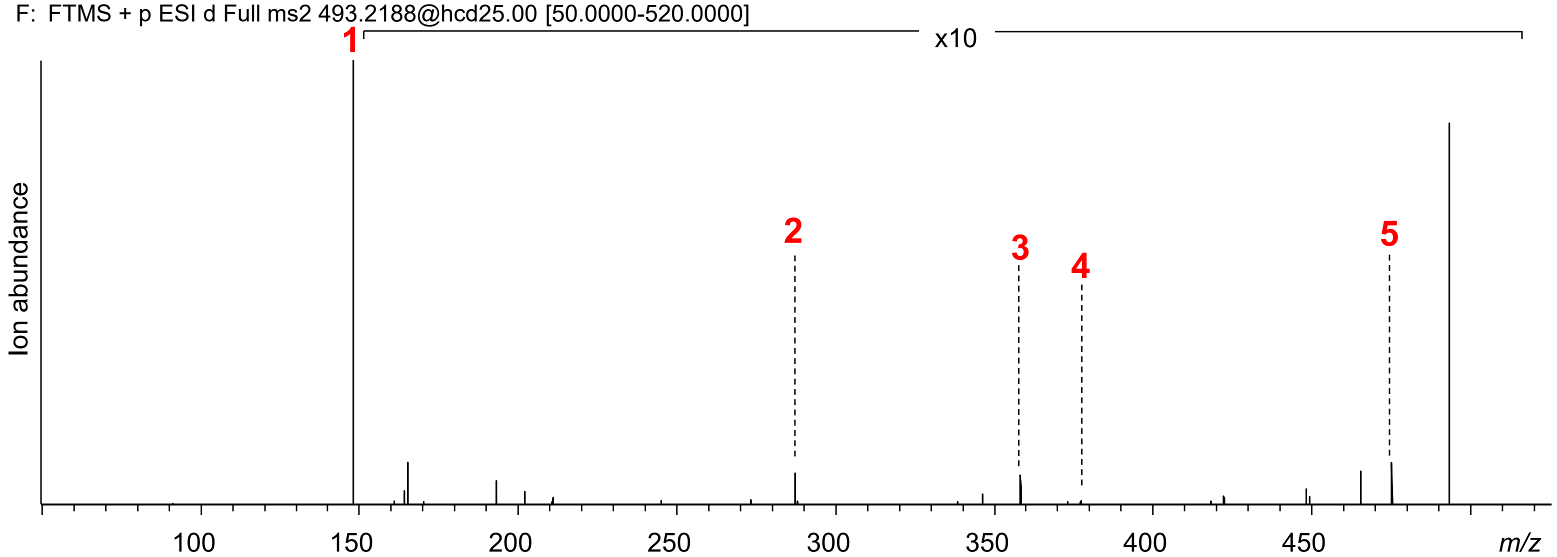


Sanjoinine A (FLFY) – m(obs) 569.3129, m(calc) 569.3122, Δm = 1.2 ppm

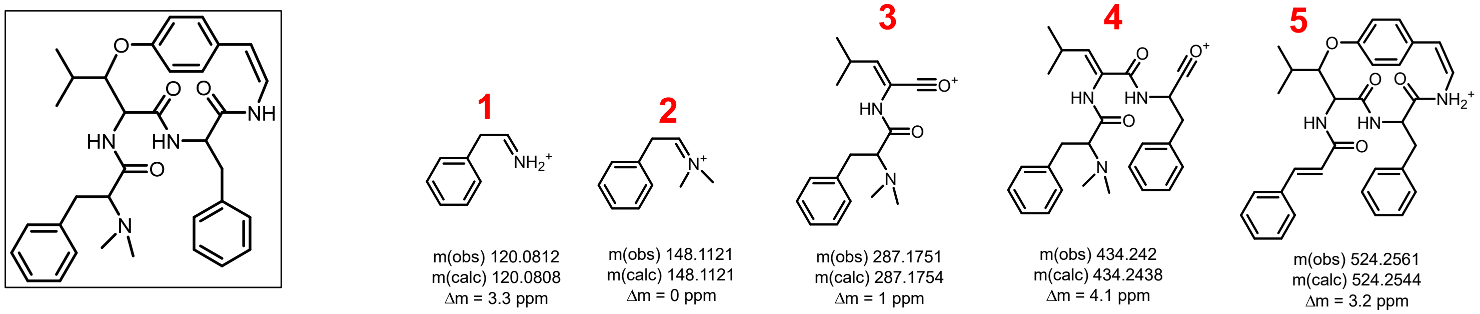


Constructs: SkrBURP-5xFLxY-ACDEF + ZjuDC + ZjuNMT

5+DC+NMT-B #2993-3029 RT: 5.61-5.65 AV: 3 NL: 5.62E6
F: FTMS + p ESI d Full ms2 569.2618@hcd25.00 [50.0000-600.0000]

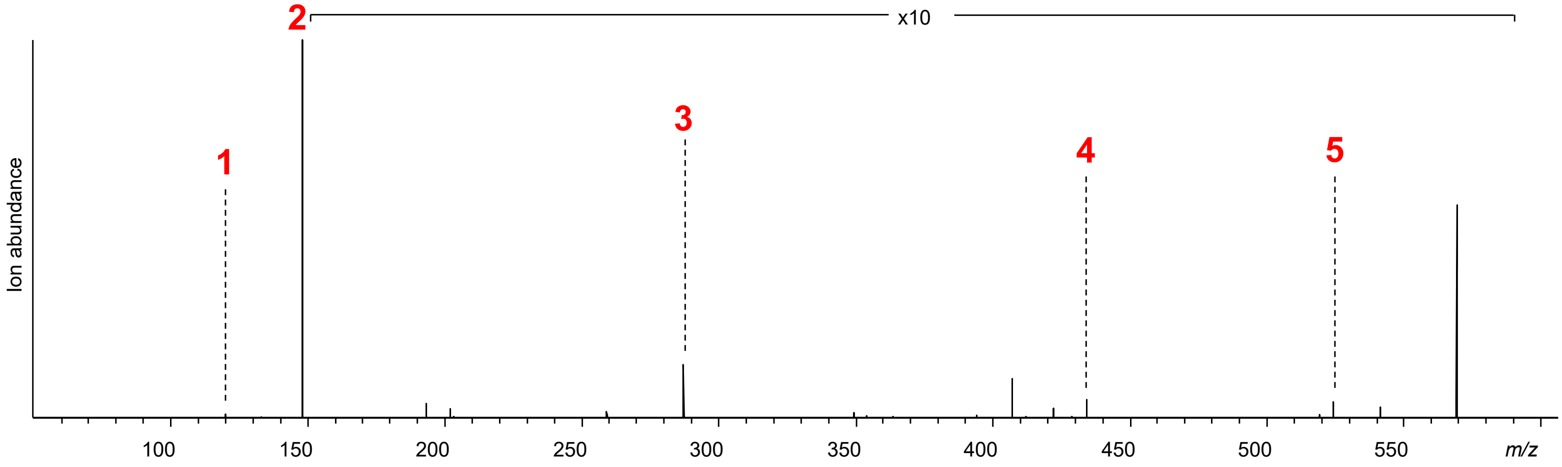


**Figure S86 | Sanjoinine A diversification via transient expression of SkrBURP-FLxY, ZjuDC and ZjuNMT in *N. benthamiana*.** Sanjoinine A (FLAY and FLFY)

Sanjoinine A (FLIY) – m(obs) 535.3284, m(calc) 535.3279, Δm = 0.9 ppm

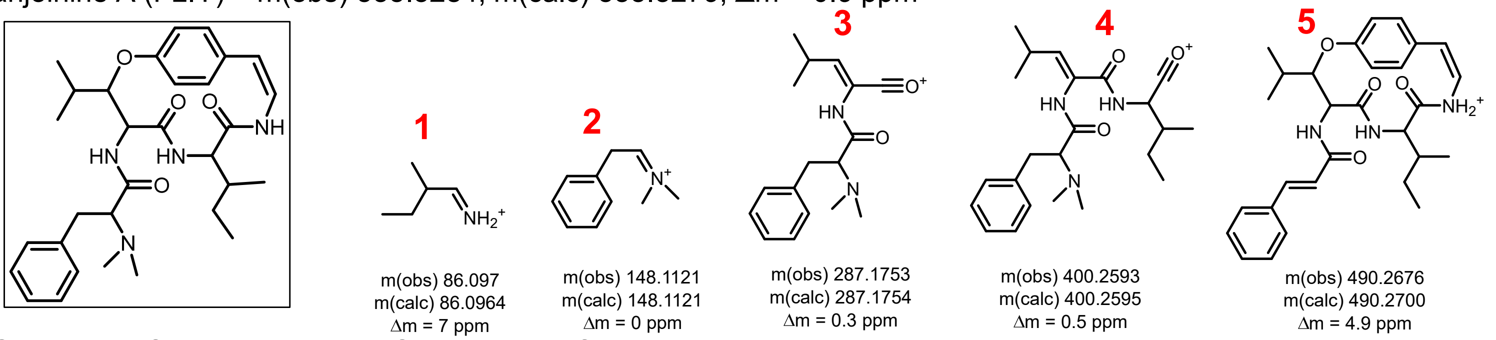


Constructs: SkrBURP-5xFLxY-GHIKL + ZjuDC + ZjuNMT

6+DC+NMT-B #2911-2932 RT: 5.45-5.46 AV: 2 NL: 9.22E6
F: FTMS + p ESI d Full ms2 535.3273@hcd25.00 [50.0000-565.0000]

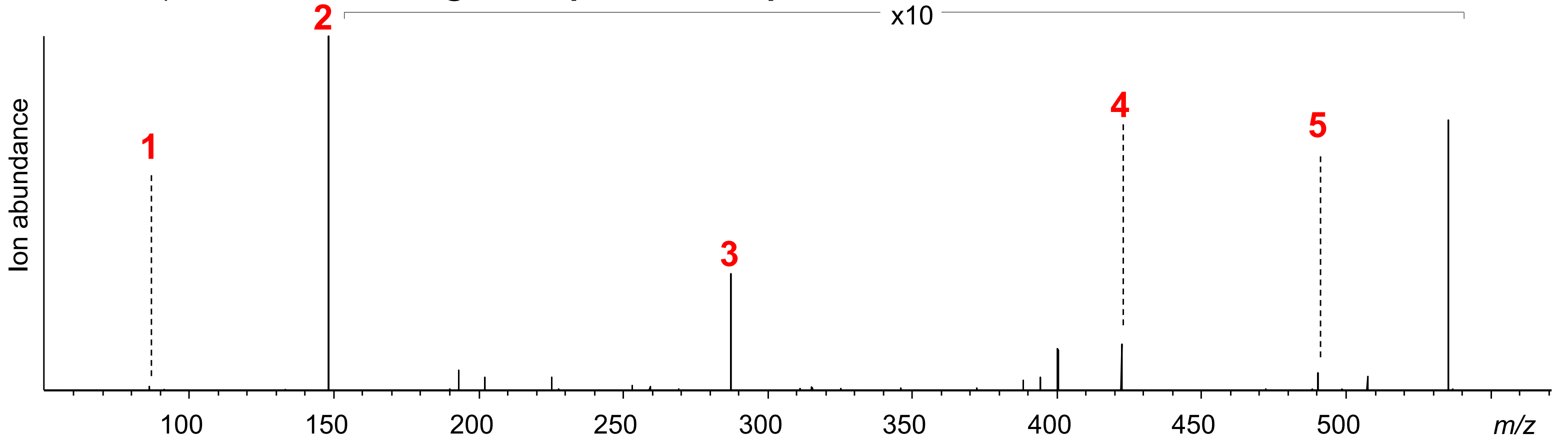


Sanjoinine A (FLLY) – m(obs) 535.328, m(calc) 535.3279, Δm = 0.2 ppm

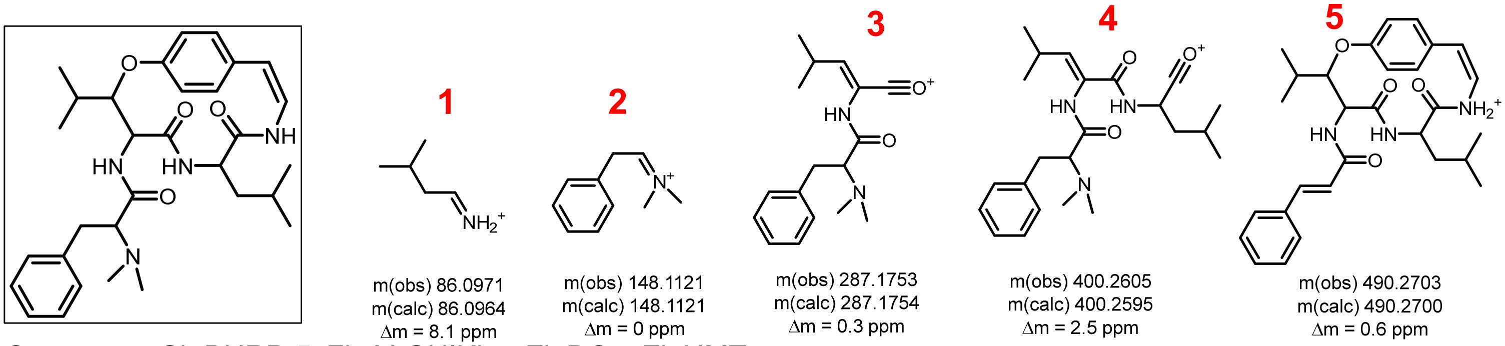


Constructs: SkrBURP-5xFLxY-GHIKL + ZjuDC + ZjuNMT

6+DC+NMT-B #2961-2987 RT: 5.53-5.57 AV: 3 NL: 1.32E7
F: FTMS + p ESI d Full ms2 535.3273@hcd25.00 [50.0000-565.0000]

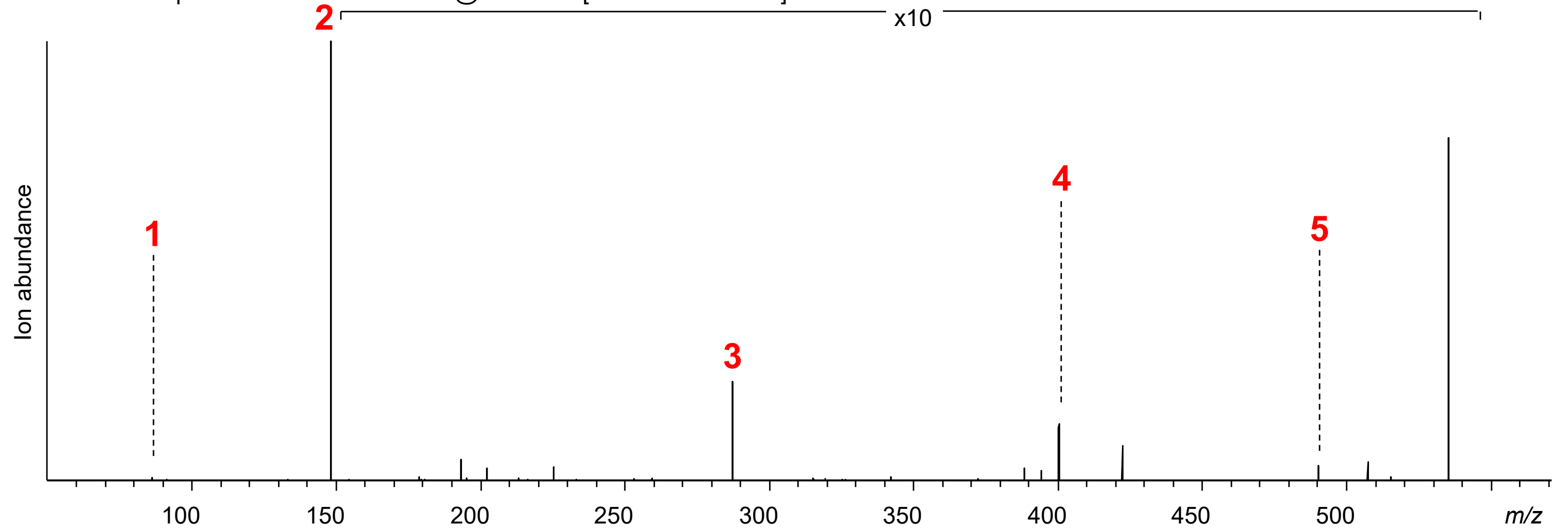


**Figure S86 | Sanjoinine A diversification via transient expression of SkrBURP-FLxY, ZjuDC and ZjuNMT in *N. benthamiana*..** Sanjoinine A (FLIY and FLLY)

Sanjoinine A (FLSY) – m(obs) 509.2765, m(calc) 509.2758, Δm = 1.4 ppm

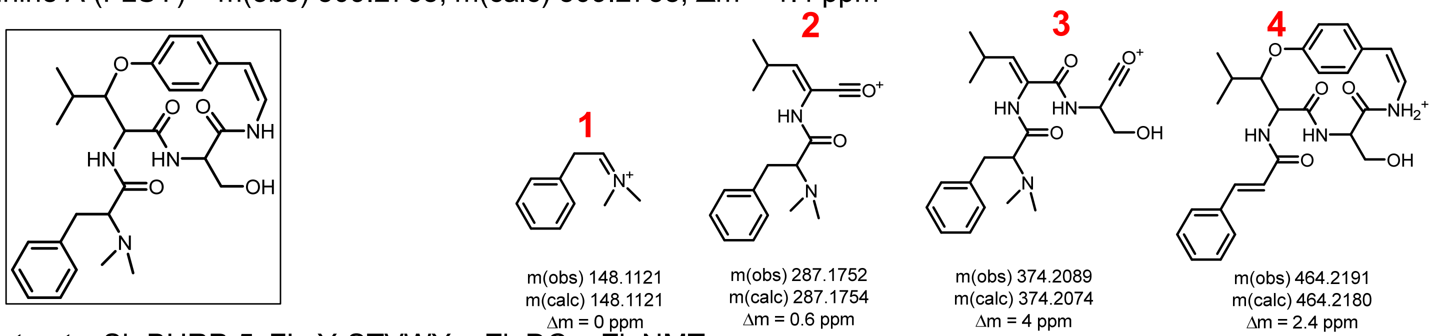


Constructs: SkrBURP-5xFLxY-STVWY + ZjuDC + ZjuNMT

7+DC+NMT-A #2492-2521 RT: 4.78-4.79 AV: 2 NL: 5.60E6
F: FTMS + p ESI d Full ms2 509.2361@hcd25.00 [50.0000-535.0000]

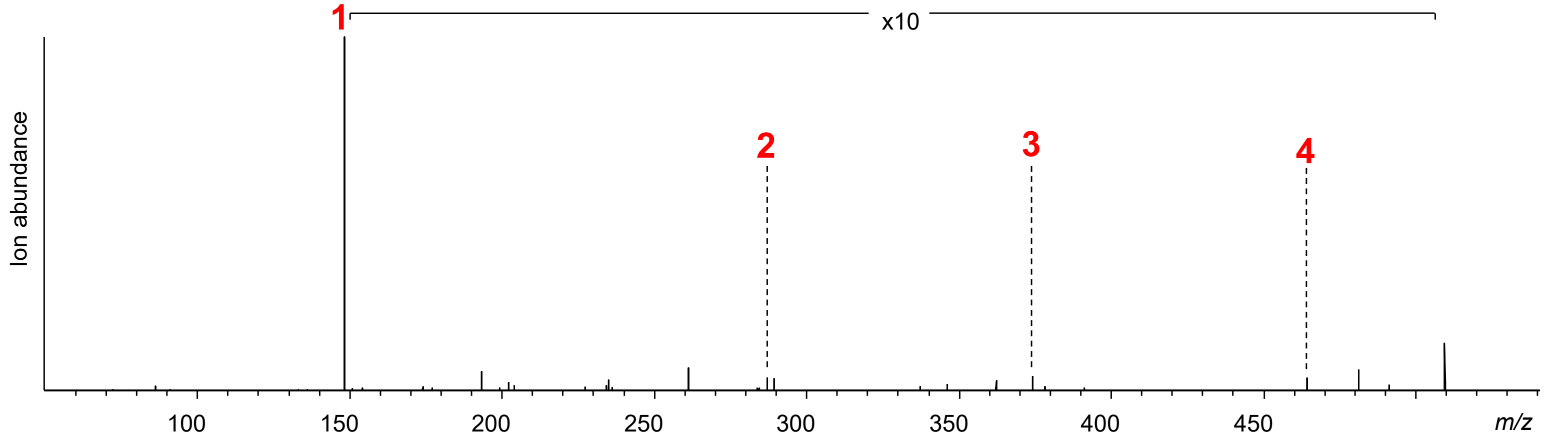


Sanjoinine A (FLTY) – m(obs) 535.328, m(calc) 535.3279, Δm = 0.2 ppm

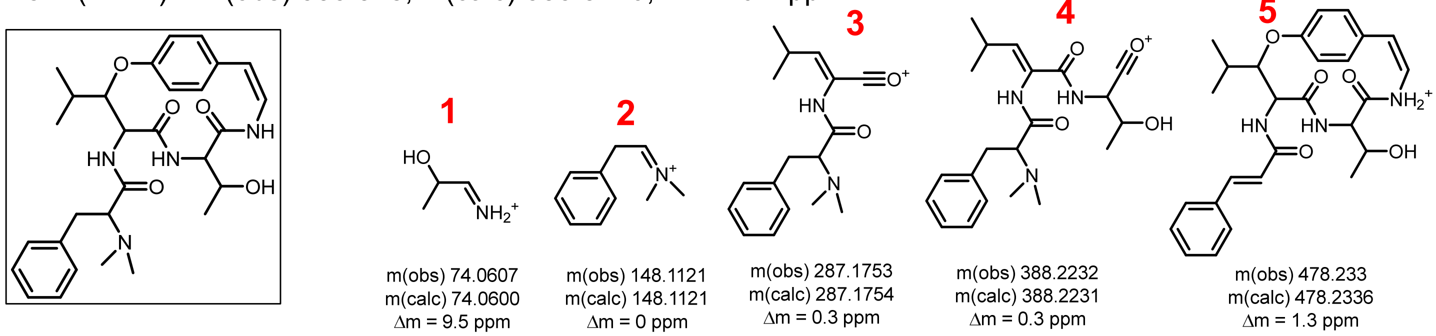


Constructs: SkrBURP-5xFLxY-STVWY + ZjuDC + ZjuNMT

7+DC+NMT-A #2593-2618 RT: 4.94-4.96 AV: 2 NL: 2.88E7
F: FTMS + p ESI d Full ms2 523.2153@hcd25.00 [50.0000-550.0000]

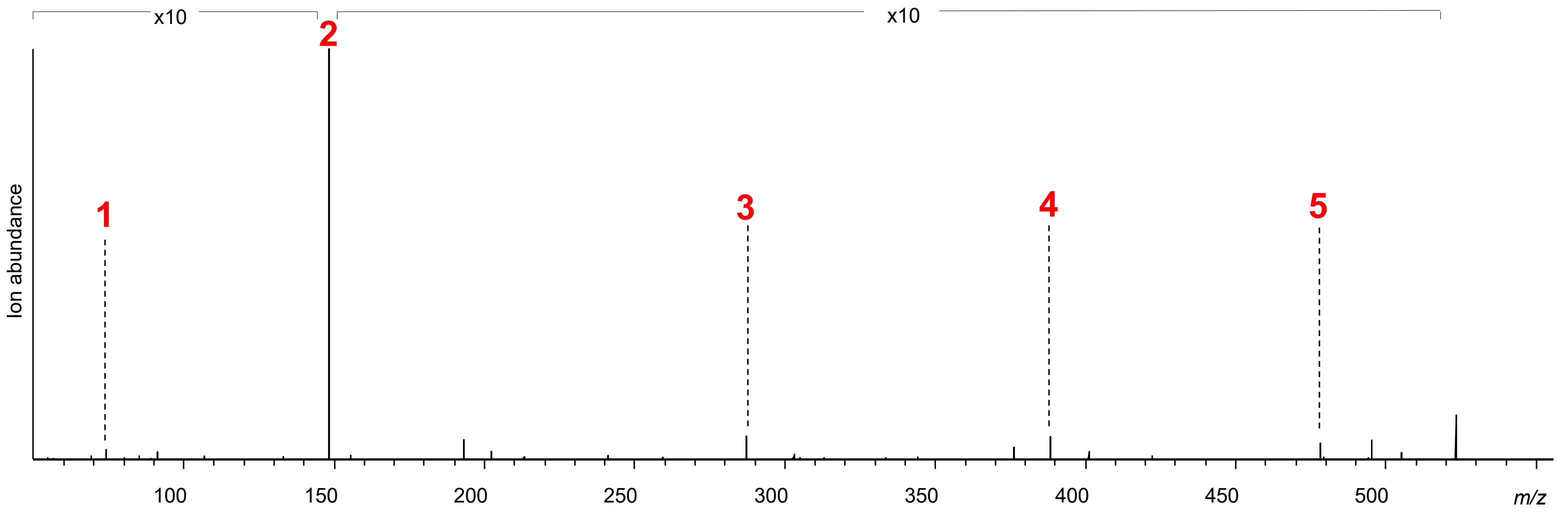


**Figure S86 | Sanjoinine A diversification via transient expression of SkrBURP-FLxY, ZjuDC and ZjuNMT in *N. benthamiana*.** Sanjoinine A (FLSY and FLTY)

Sanjoinine A (FLVY) – m(obs) 521.3123, m(calc) 521.312, Δm = 0.6 ppm

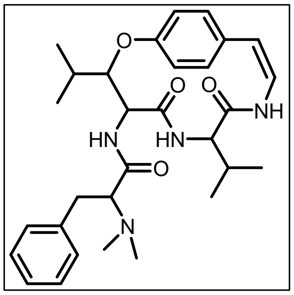

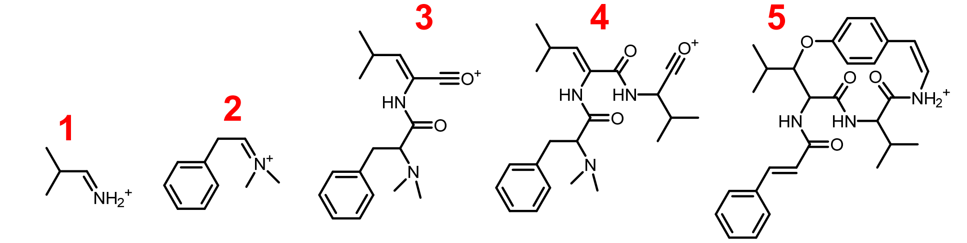


| 1 | 2 | 3 | 4 | 5 |
|---|---|---|---|---|
| m(obs) 72.0814 | m(obs) 148.1121 | m(obs) 287.1753 | m(obs) 386.2437 | m(obs) 476.2539 |
| m(calc) 72.081 | m(calc) 148.1121 | m(calc) 287.1754 | m(calc) 386.244 | m(calc) 476.254 |
| Δm = 5.5 ppm | Δm = 0 ppm | Δm = 0.3 ppm | Δm = 0.8 ppm | Δm = 0.2 ppm |

Constructs: SkrBURP-5xFLxY-STVWY + ZjuDC + ZjuNMT

7+DC+NMT-B#2782-2810 RT: 5.24-5.28 AV: 3 NL: 1.97E7
F: FTMS + p ESI d Full ms2 521.3126@hcd25.00 [50.0000-550.0000]

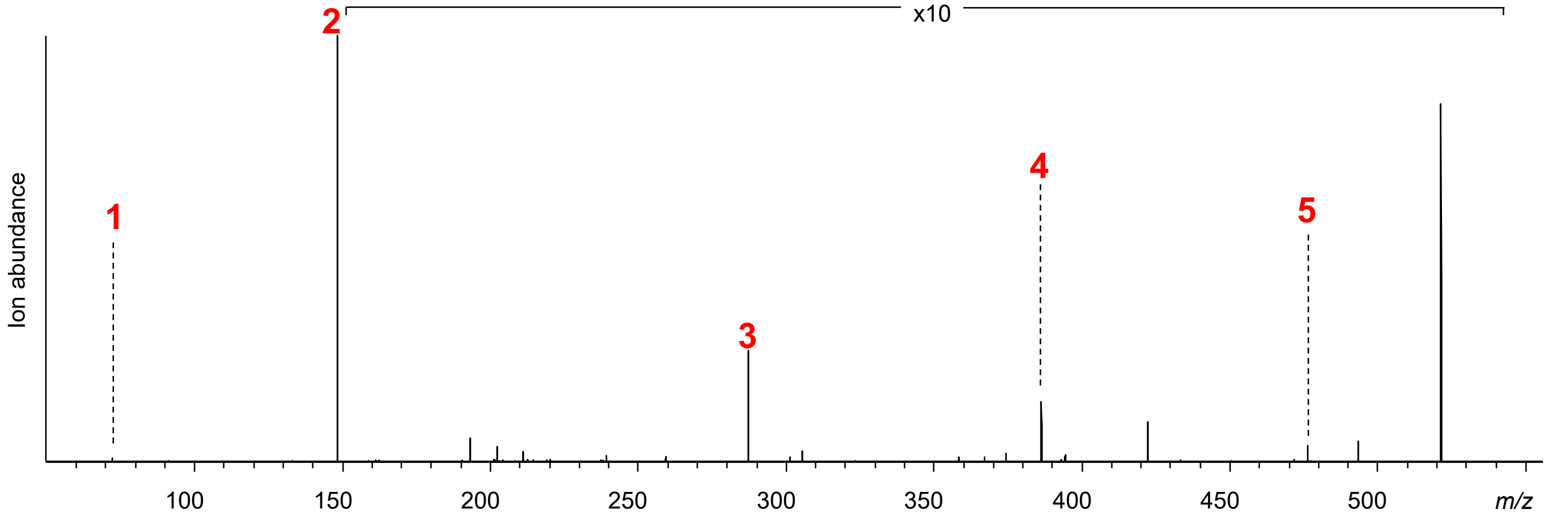


**Figure S86 | Sanjoinine A diversification via transient expression of SkrBURP-FLxY, ZjuDC and ZjuNMT in *N. benthamiana*.** Sanjoinine A (FLVY)

Adouetine X (LLIY) – m(obs) 501.3435, m(calc) 501.344, Δm = 1 ppm

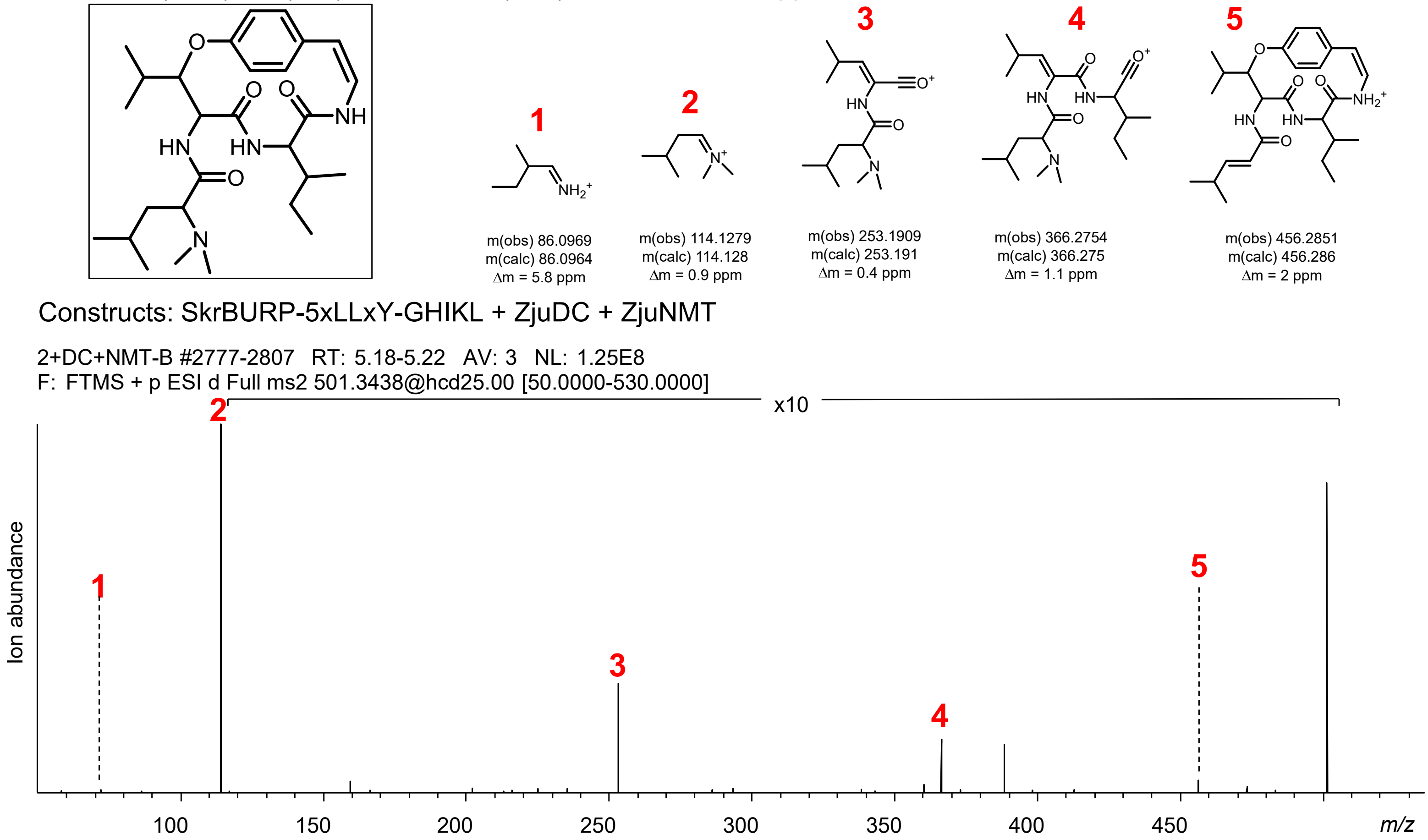


Adouetine X (LLLY) – m(obs) 501.3434, m(calc) 501.344, Δm = 1.2 ppm

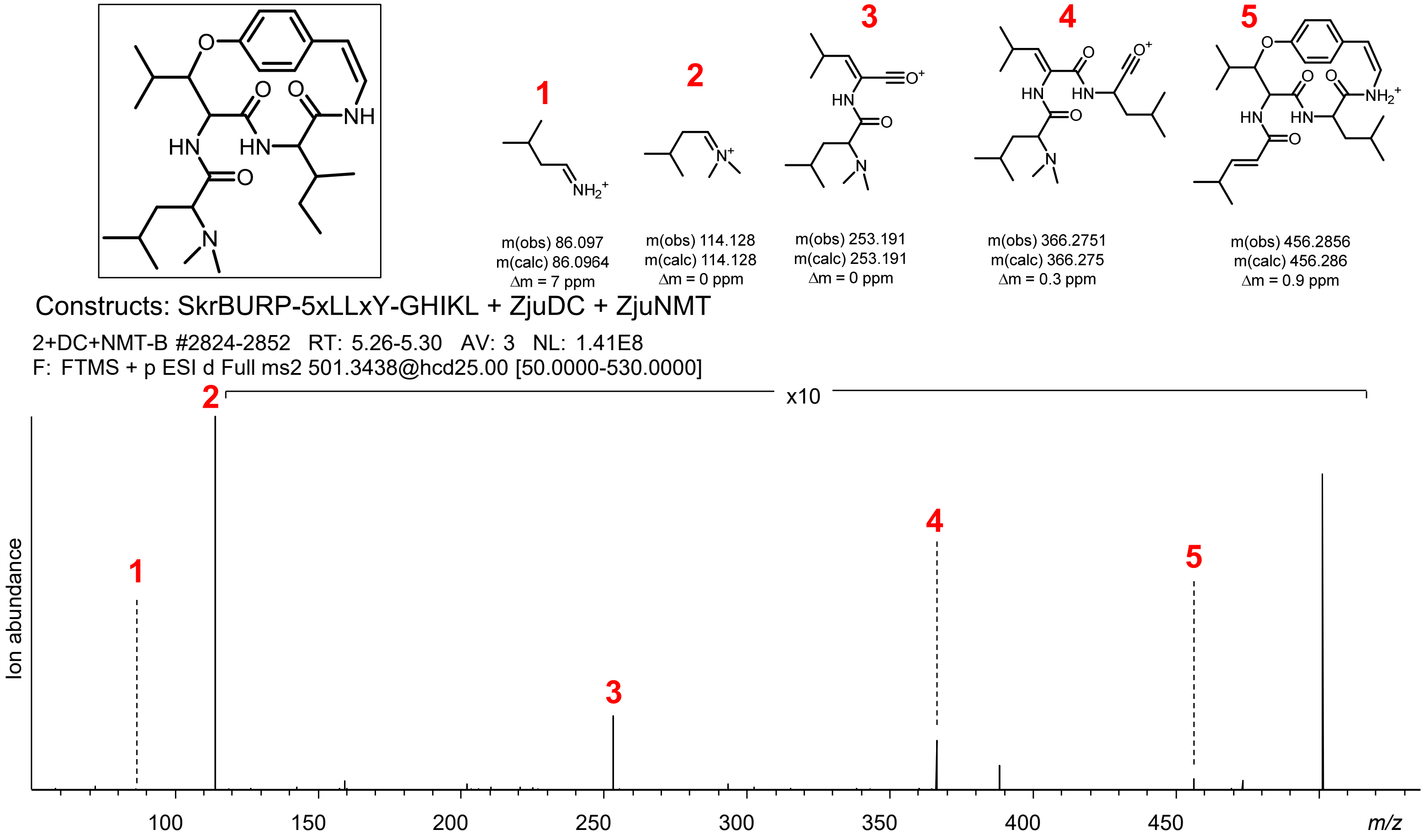


**Figure S87 | Adouetine X diversification via transient expression of SkrBURP-LLxY, ZjuDC and ZjuNMT.** Adouetine X (LLIY and LLLY)

Adouetine X (LLTY) – m(obs) 489.3074, m(calc) 489.307 Δm = 0.8 ppm

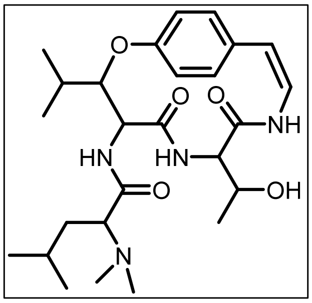

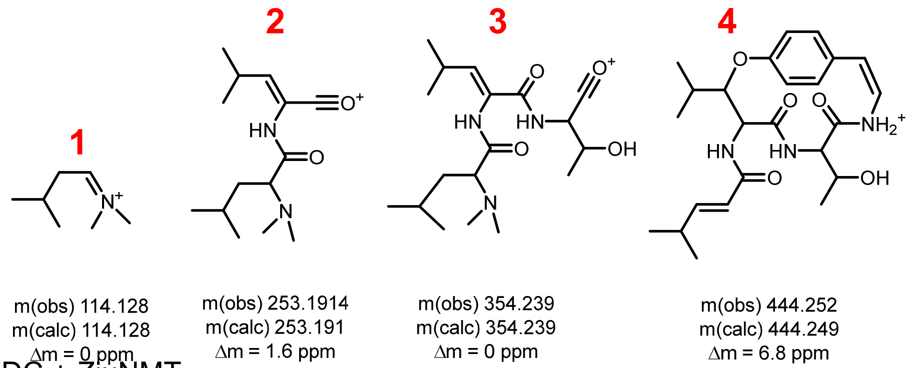


Constructs: SkrBURP-5xLLxY-STVWY + ZjuDC + ZjuNMT

4+DC+NMT-A#2231 RT: 4.60 AV: 1 NL: 3.82E6
F: FTMS + p ESI d Full ms2 489.2621@hcd25.00 [50.0000-515.0000]

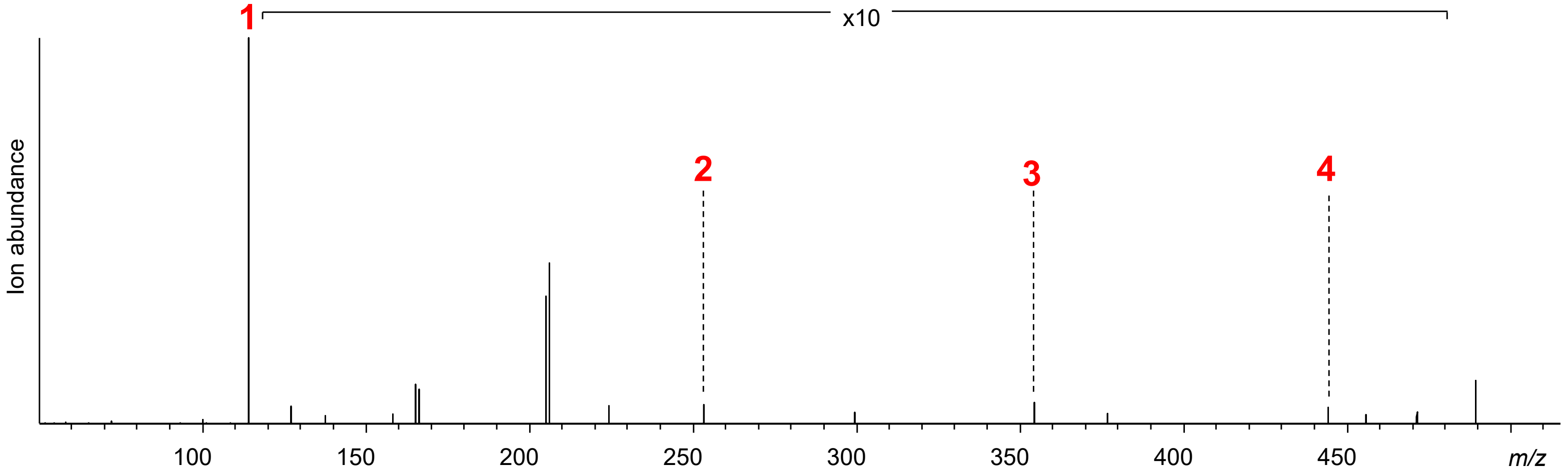


Adouetine X (LLVY) – m(obs) 487.3279, m(calc) 487.328, Δm = 0.2 ppm

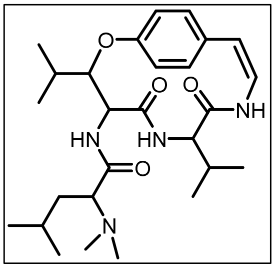

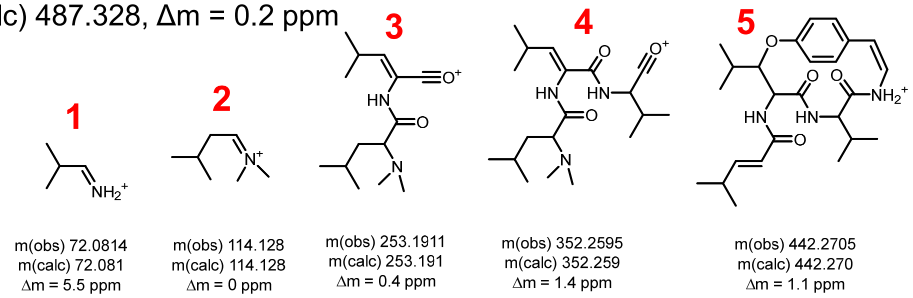


Constructs: SkrBURP-5xLLxY-STVWY + ZjuDC + ZjuNMT
4+DC+NMT-A #2567-2629 RT: 4.94-5.00 AV: 4 NL: 5.66E7
F: FTMS + p ESI d Full ms2 487.2538@hcd25.00 [50.0000-515.0000]

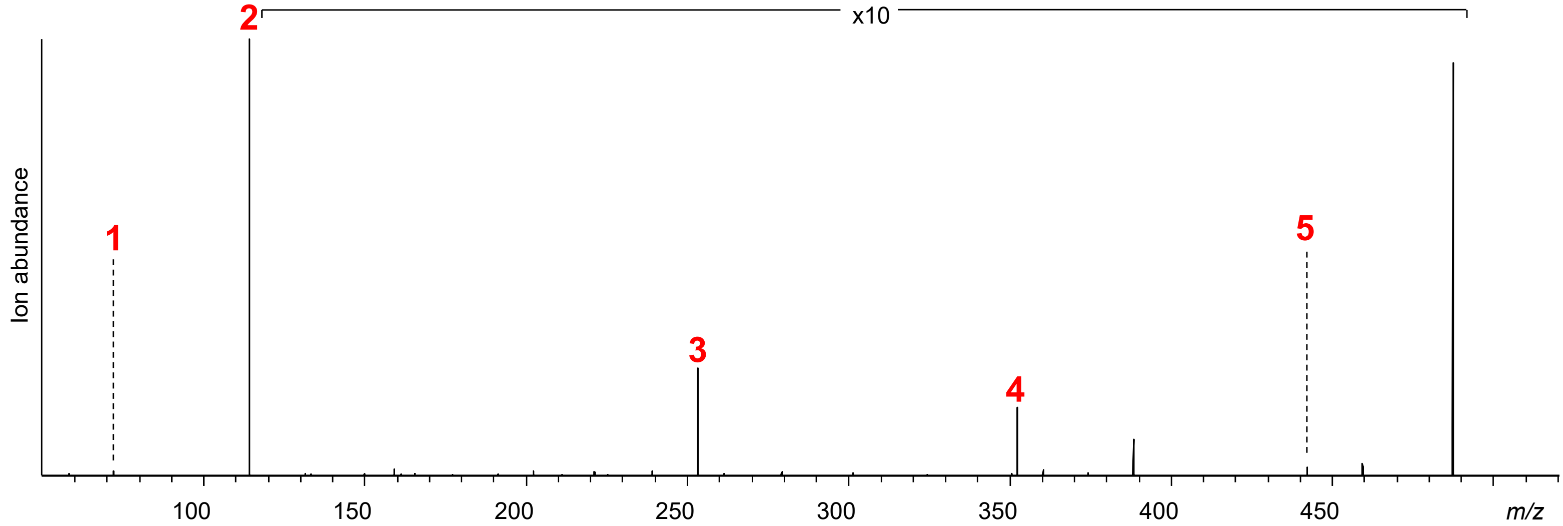


**Figure S87 | Adouetine X diversification via transient expression of SkrBURP-LLxY, ZjuDC and ZjuNMT.** Adouetine X (LLTY and LLVY)

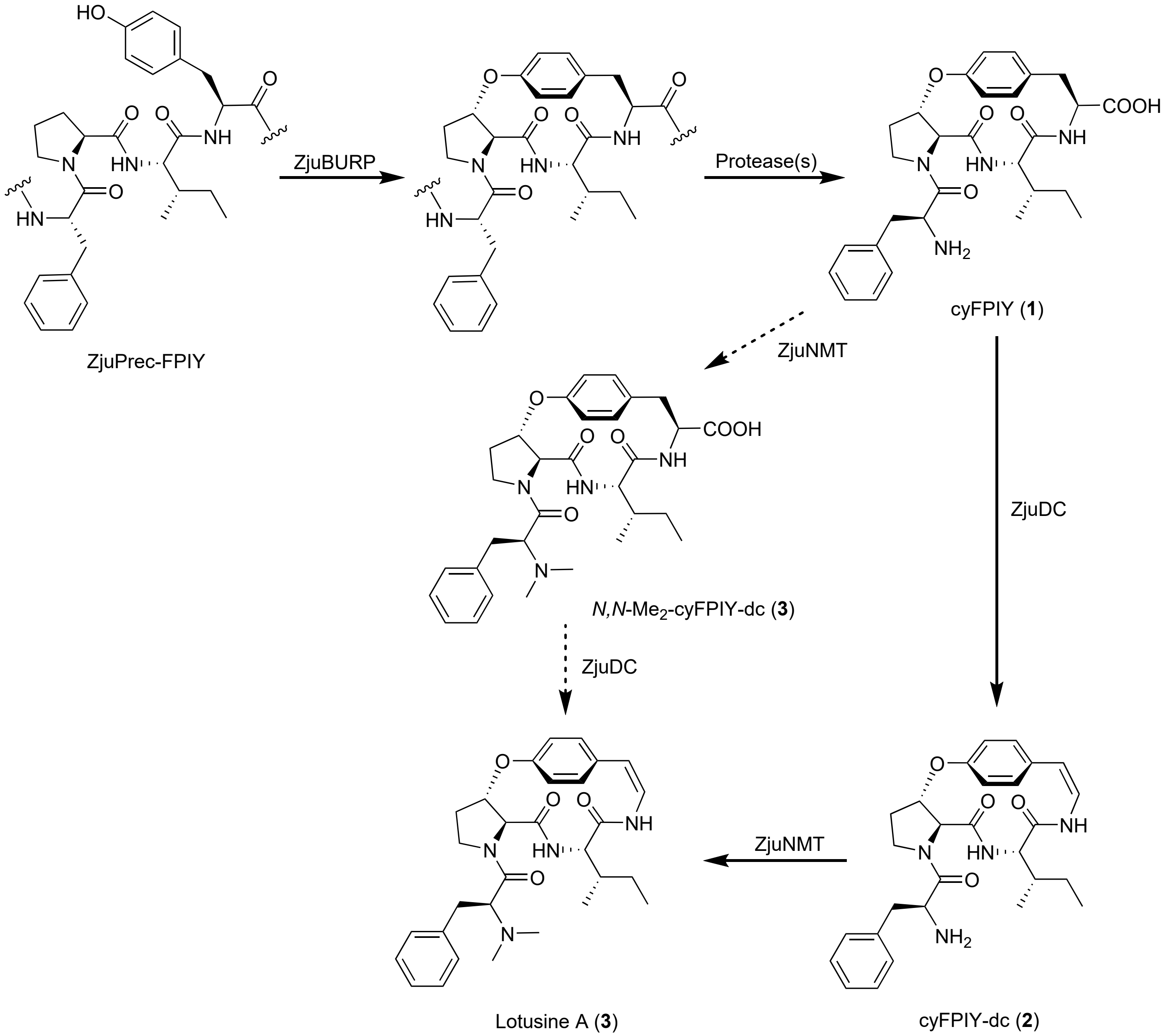


**Figure S88 | Biosynthetic pathway of lotusine A in *Ziziphus jujuba*.** Dashed arrows indicates alternative pathway through putative intermediate (**4**).

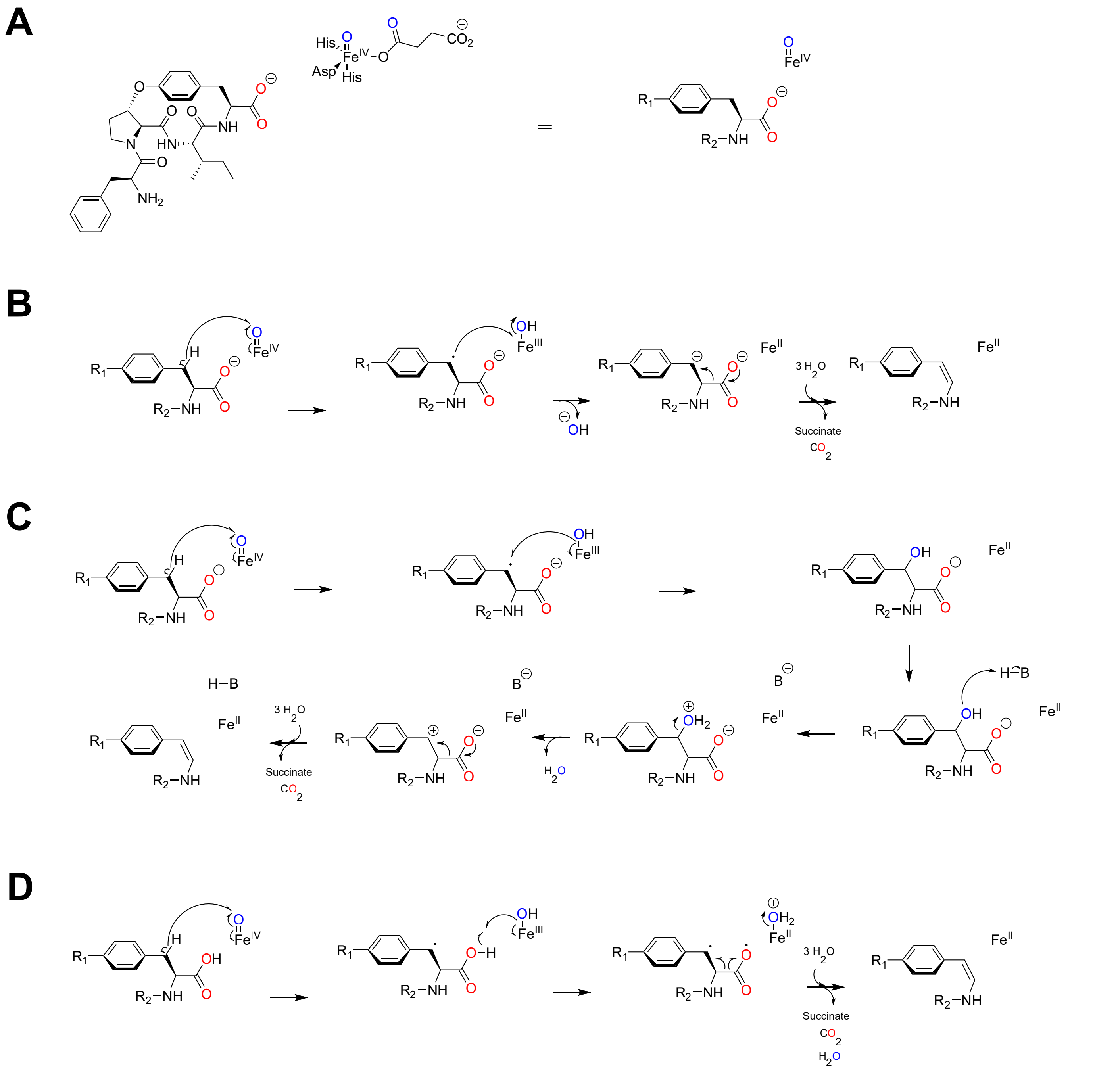


**Figure S89 | Proposed mechanisms for ZjuDC based on intermediates.** (**A**) Simplified reaction scheme after Fe(II), $O_2$, and 2-oxoglutarate react producing a loss of $CO_2$ and formation of succinate[8]. Molecular oxygen is blue, the oxygen from substrate cyclic peptide is red. (**B**) Single electron transfer and carbocation intermediate (**C**) Hydroxylation and carbocation intermediate. (**D**) Diradical formation and homolytic cleavage.
